# Emergence of neuronal diversity during vertebrate brain development

**DOI:** 10.1101/839860

**Authors:** Bushra Raj, Jeffrey A. Farrell, Aaron McKenna, Jessica L. Leslie, Alexander F. Schier

## Abstract

Neurogenesis in the vertebrate brain comprises many steps ranging from the proliferation of progenitors to the differentiation and maturation of neurons. Although these processes are highly regulated, the landscape of transcriptional changes and progenitor identities underlying brain development are poorly characterized. Here, we describe the first developmental single-cell RNA-seq catalog of more than 200,000 zebrafish brain cells encompassing 12 stages from 12 hours post-fertilization to 15 days post-fertilization. We characterize known and novel gene markers for more than 800 clusters across these timepoints. Our results capture the temporal dynamics of multiple neurogenic waves from embryo to larva that expand neuronal diversity from ∼20 cell types at 12 hpf to ∼100 cell types at 15 dpf. We find that most embryonic neural progenitor states are transient and transcriptionally distinct from long-lasting neural progenitors of post-embryonic stages. Furthermore, we reconstruct cell specification trajectories for the retina and hypothalamus, and identify gene expression cascades and novel markers. Our analysis reveal that late-stage retinal neural progenitors transcriptionally overlap cell states observed in the embryo, while hypothalamic neural progenitors become progressively distinct with developmental time. These data provide the first comprehensive single-cell transcriptomic time course for vertebrate brain development and suggest distinct neurogenic regulatory paradigms between different stages and tissues.

## INTRODUCTION

The vertebrate brain develops from a limited pool of embryonic neural progenitor cells that cycle through rounds of proliferation, diversification, and terminal differentiation into an extensive catalogue of distinct neuronal and glial cell types. A central goal in developmental neurobiology is to investigate how neuronal complexity arises through molecular specification and commitment by studying the origins and fates of cells during development. Critical insights into these processes have been gained via classic approaches using genetic markers, perturbations and fate mapping ^1-7^. However, several outstanding questions pertaining to brain organogenesis remain. For example, it is poorly understood how embryonic neural progenitors are molecularly related to post-embryonic neural progenitors. Do embryonic cell states exist over long periods of time or are they transient? What are the transcriptional differences between these progenitor states and do they vary across different regions of the brain? Furthermore, transcriptional programs that are activated and/or suppressed as neural progenitors become fate-restricted and terminally divide are largely unknown.

To complement previous studies and address the above questions, we set out to obtain global views of neurogenesis, cell type heterogeneity, cell specification trajectories and cell lineage relationships in a developing vertebrate brain. We describe the first extensive characterization of a developmental compendium of the zebrafish brain obtained by profiling cells with single-cell RNA-seq (scRNA-seq) across 12 stages ranging from 12 hours post fertilization (hpf) to 15 days post fertilization (dpf). We also improved lineage barcode capture of our previously described scGESTALT CRISPR lineage recorder ^8^. The transgenic line and accompanying brain lineage trees for 15 dpf larvae complement the transcriptional atlas and are available for use and exploration by the community. Using the cell type atlas, we describe the expansion of neuronal diversity from 12 hpf to 15 dpf based on known and novel marker genes, loss of transient embryonic neural progenitor states, and the long-term maintenance of distinct larval neural progenitor states. Furthermore, we reconstruct cell specification trajectories of the zebrafish retina and hypothalamus revealing gene expression cascades that underlie development of profiled cell types in these tissues. Finally, we find that neural progenitor differentiation paradigms are distinct between the retina and hypothalamus, thus highlighting how neuronal complexity in various regions can be generated using different strategies. Collectively, our data reveal molecular and cellular changes that accompany brain development at an unprecedented scale and resolution, and lay the foundation for detailed analyses of how neuronal diversity emerges in vertebrates.

## RESULTS

### Building a developmental atlas of the zebrafish brain with single-cell transcriptomics

To determine the spectrum of heterogeneous cell states and cell types during brain development, we profiled 223,037 cells across 12 stages of zebrafish embryonic and larval growth using the 10X Chromium scRNA-seq platform. Samples spanned from 12 hpf (shortly after gastrulation), when the embryo is undergoing early developmental patterning, to 15 dpf when larvae are mature, exhibit complex behaviors, and are expected to exhibit substantial cell type diversity (Figure 1a). To enrich for brain cell types, we dissected the heads of animals from 12 hpf to 3 dpf, and the brains and eyes from 5 dpf to 15 dpf (Figure 1b). A t-distributed Stochastic Neighbor Embedding (t-SNE) visualization of the full dataset generated a developmental atlas of transcriptionally distinct cell populations in the head and brain of zebrafish (Figure 1c). Plotting expression of known cell type markers identified clusters corresponding to neural progenitors (*sox19a*), neurons (*elavl3, gad2, slc17a6b*), eye cells (*foxg1b, lim2.4, pmela, ca14, gnat1, opn1mw1*), radial glia (*mfge8a, s100b*), neural crest (*sox10*), oligodendrocytes (*mbpa*), blood cells (*cahz, etv2, cd74a*), cartilage (*matn4, col9a2*), pharyngeal arches (*pmp22a, prrx1b, barx1*), sensory placodes (*dlx3b, six1b*), and epidermal cells (*epcam, cldni*), among others (Figure 1d). The atlas also revealed groups of embryonic clusters that are transcriptionally distinct from larval clusters (e.g. placodes, neural progenitors; see below), suggesting that many embryonic cell states/types are transient.

**Figure 1.**
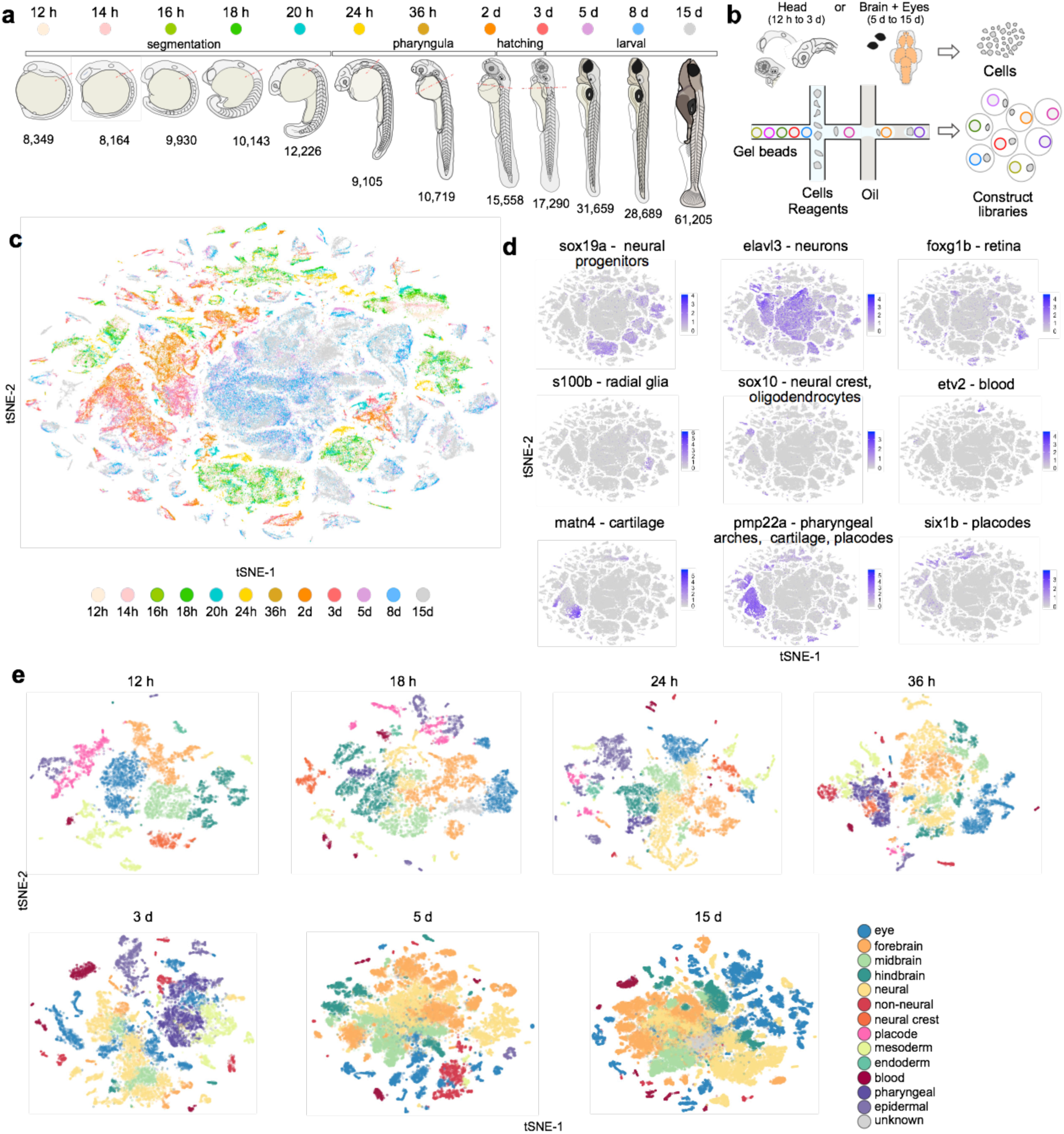
Developmental compendium of zebrafish brain cell types. **a.** Schematic of the developmental stages profiled. Red hatched line represents head regions that were selected for enrichment of brain cells in early development. Samples from 5 to 15 dpf were dissected to obtain brain and eye specifically. **b.** Schematic of scRNA-seq using 10X Genomics platform. **c.** tSNE plot of the full dataset (223,037 cells). Cells are color coded by stage. **d.** Gene expression of cell type markers. Cells are colored by mean gene expression level. **e.** Cell type heterogeneity within each stage. Clusters at each stage were assigned to a region or tissue type based on known markers and color coded to reflect their classification. tSNE implementations: Barnes-Hut (**e** 12h to 3d), Fourier transform (**c, d, e** [5d and 15d]

To obtain higher cell type resolution, data from each stage was analyzed individually using Louvain clustering (Figure 1e and Sup Fig.1). This identified a total of 815 cell clusters across all 12 timepoints (Sup Table). To classify each cluster, we compared enriched gene markers with existing gene expression annotations in the ZFIN database and literature, as described previously ^8^. As expected, cell type complexity increases with developmental time.

To enable direct comparison of cell types across our time course, we subsetted the 12 hpf dataset to only comprise neural populations and blood cells. This resulted in an initial set of 21 clusters at 12 hpf (Figure 2a) that diversified into 98 clusters by 15 dpf (Figure 2c). Notably, most clusters could be uniquely identified using a minimal group of 2-3 enriched gene markers (Figure 2b, 2d). For example, at 12 hpf, the optic vesicle is identified by expression of *rx2* and *rx3*; hindbrain rhombomeres 5/6 by *hoxb3a* and *eng2b*; and ventral diencephalon by *nkx2.4a* and *dbx1a*. Similarly, at 15 dpf, the cerebellar granule cells are marked by expression of *oprd1b* and *zic2a*; optic tectum by *pax7a* and *tal1*; and a new retinal cell type by *kidins220a, foxg1b* (exclusively detected in retinal cells) and *tbx3a*. We did not find unique gene combinations for cycling progenitors, committed progenitors and newly born neurons, as many of these subtypes had similar expression patterns of generic neuronal or progenitor marker genes, such as *elavl3* and *tubb5* in neurons, and *rpl5a* and *npm1a* in progenitors (Figure 2d, grey box).

**Figure 2.**
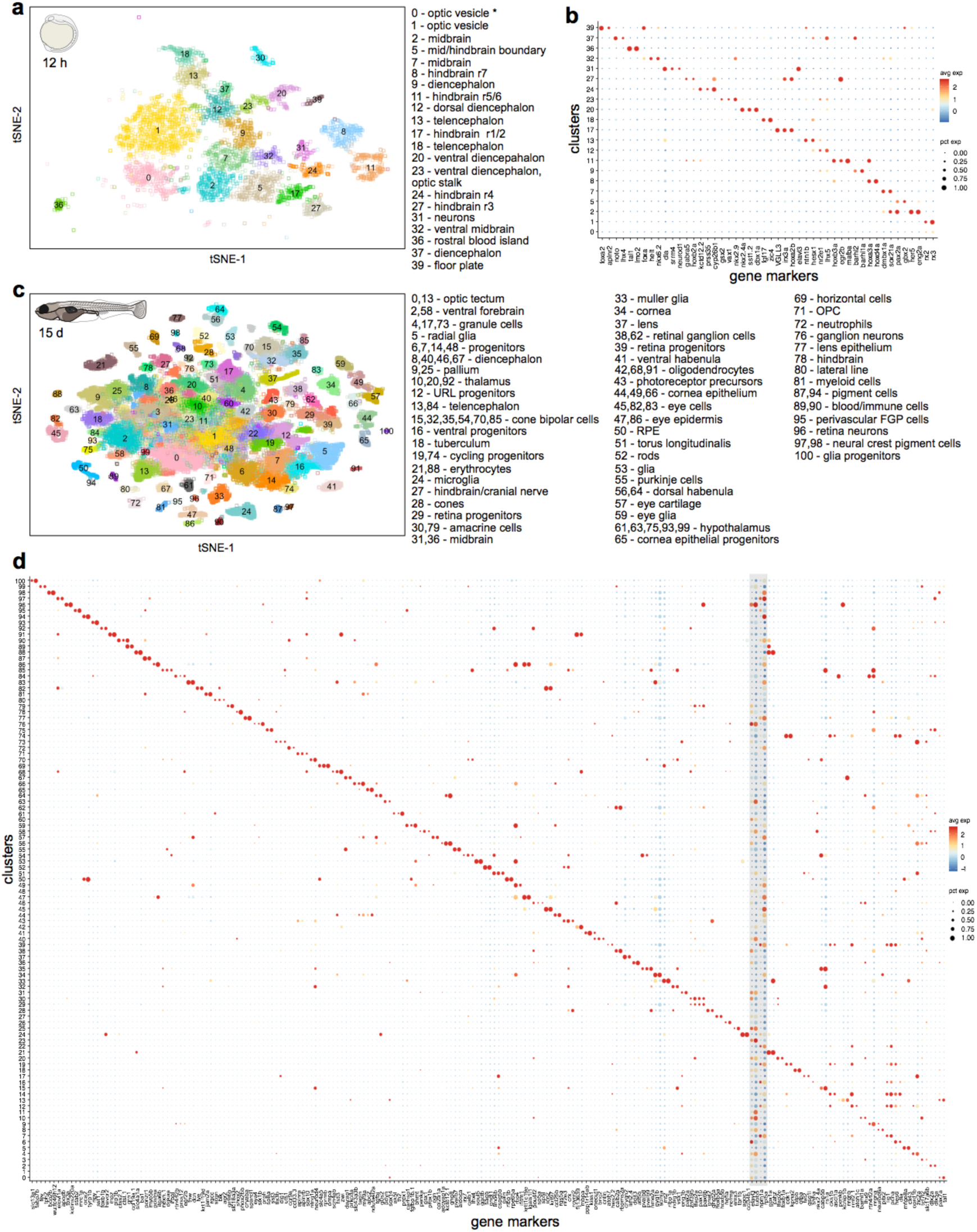
Cell type diversification from 12 hpf to 15 dpf. **a.** tSNE plot of 12 hpf dataset. Only clusters corresponding to neural and blood cell types are shown. Inferred identities of each cluster are described. **b.** Dot plot of gene expression pattern of select marker genes (columns) for each cluster (row). Dot size indicates the percentage of cells expressing the marker; color represents the average scaled expression level. **c.** tSNE plot of 15 dpf dataset. Inferred identities of each cluster are described. **d.** Dot plot of gene expression patterns of select marker genes for each cluster. Layout is same as panel **b**. Grey box represents generic neuronal and progenitor genes. tSNE implementations: Barnes-Hut (**a**), Fourier transform (**c**)

At 12 hpf, the early demarcation of multiple brain regions is already apparent and by 15 dpf these regions expand and diversify further. For example, the optic vesicle at 12 hpf is defined by one cluster and is the origin of 18 retinal neuron clusters and the retinal pigment epithelium at 15 dpf. Similarly, a single cluster of ventral diencephalon cells (expressing *shha, nkx2.4a, nkx2.1, rx3*) at 12 hpf develops into 7 major hypothalamus cell types at 15 dpf. An exception to this diversification is the loss of rhombomeres (r1-r7) in the hindbrain ^9^.

To derive lineage relationships of cell types identified in our brain development atlas, we performed lineage recording experiments with scGESTALT, which enables simultaneous cell type and cell lineage identification by combining scRNA-seq with CRISPR-Cas9 barcode editing ^8,10^. We optimized the lineage recording cassette to enable higher recovery of barcodes (see Methods). With this improved system, we barcoded early embryonic lineage relationships by injecting Cas9 and target guide RNAs into single-cell embryos (Sup. Fig 2a; see Methods). We recovered high quality barcodes from 5,794 cells total (recovery rate 30-75% compared to 6-28% in previous scGESTALT version ^8^) from four 15 dpf larval brains. Edited barcodes showed no overlap between animals, displayed a diverse spectrum of repair products that spanned single and multiple sites, and were of varying clone sizes (Sup Fig. 2). Using the recovered barcodes and associated transcriptomes, we reconstructed lineage trees of 15 dpf zebrafish representing cell type relationships in early brain development (Sup Fig. 3-4). As observed previously ^8^, cells sharing identical barcodes were generally locally enriched within compartments of the brain and eye (Sup Fig.5), agreeing with classic fate mapping experiments ^4^. Furthermore, these barcodes were distinct from those marking cell types of non-neural origins (e.g. blood, cornea, immune, etc.). These lineage trees accompany our transcriptional cell type atlas, and are available to explore at https://scgestalt.mckennalab.org/.

In summary, we generated the first developmental zebrafish brain cell type atlas across 12 stages spanning early and late brain organogenesis. Next, we mined our transcriptional atlas further and investigated global hierarchies and regulatory strategies in neural development and their associations with cell fate decisions.

### Neurogenic waves during brain development

During development, cell composition shifts from predominantly progenitor populations to more differentiated cell types ^11^. To better characterize how differentiation varies during neuronal development, we first asked if our dataset captures the two neurogenic waves before and after 2 dpf defined through histological analyses ^12-14^. At 12 hpf (early-stage primary neurogenesis), the brain is enriched in *sox19a* expressing progenitors, while neuronal differentiation (marked by *elavl3* expression) is robustly detected in only three small populations: olfactory placode, trigeminal placode and telencephalon (Figure 3a). With increasing developmental time, we observe a progressive decrease in *sox19a*^*+*^ cells with a concomitant increase in *elavl3*^+^ neurons. Notably, during late embryogenesis, neuronal populations expand substantially from 12 *elavl3*-enriched clusters at 20-24 hpf to 25 clusters at 36 hpf. This burst coincides with the presumed timing of late-stage primary neurogenesis in zebrafish ^12^. Furthermore, clusters of excitatory (*slc17a6a*^+^, *slc17a6b*^+^) and inhibitory (*gad1b*^+^, *gad2*^+^) neurons are detected by 20 hpf, and are enriched in neuronal markers (e.g. *ywhag2, gap43, snap25a, scg2b, elavl4*) characteristic of nascent neurons (Sup Fig. 6, Sup Table). By 5 dpf a second large expansion of neuronal populations, corresponding to the secondary neurogenic wave ^12^ occurs. This timepoint comprises cell types identified as early as 36 hpf, as well as subtypes only observed during the second wave, such as *nrgnb+ prkcda+* neurons in the forebrain, and photoreceptor cells, cone bipolar cells and the newly identified *kidins220a+* neuron subtype in the retina (49 neuronal subtypes at 5 dpf vs 25 subtypes at 36 hpf). Collectively, our dataset captures both waves of neurogenesis and reveals the temporal diversification of neurons in multiple brain structures.

**Figure 3.**
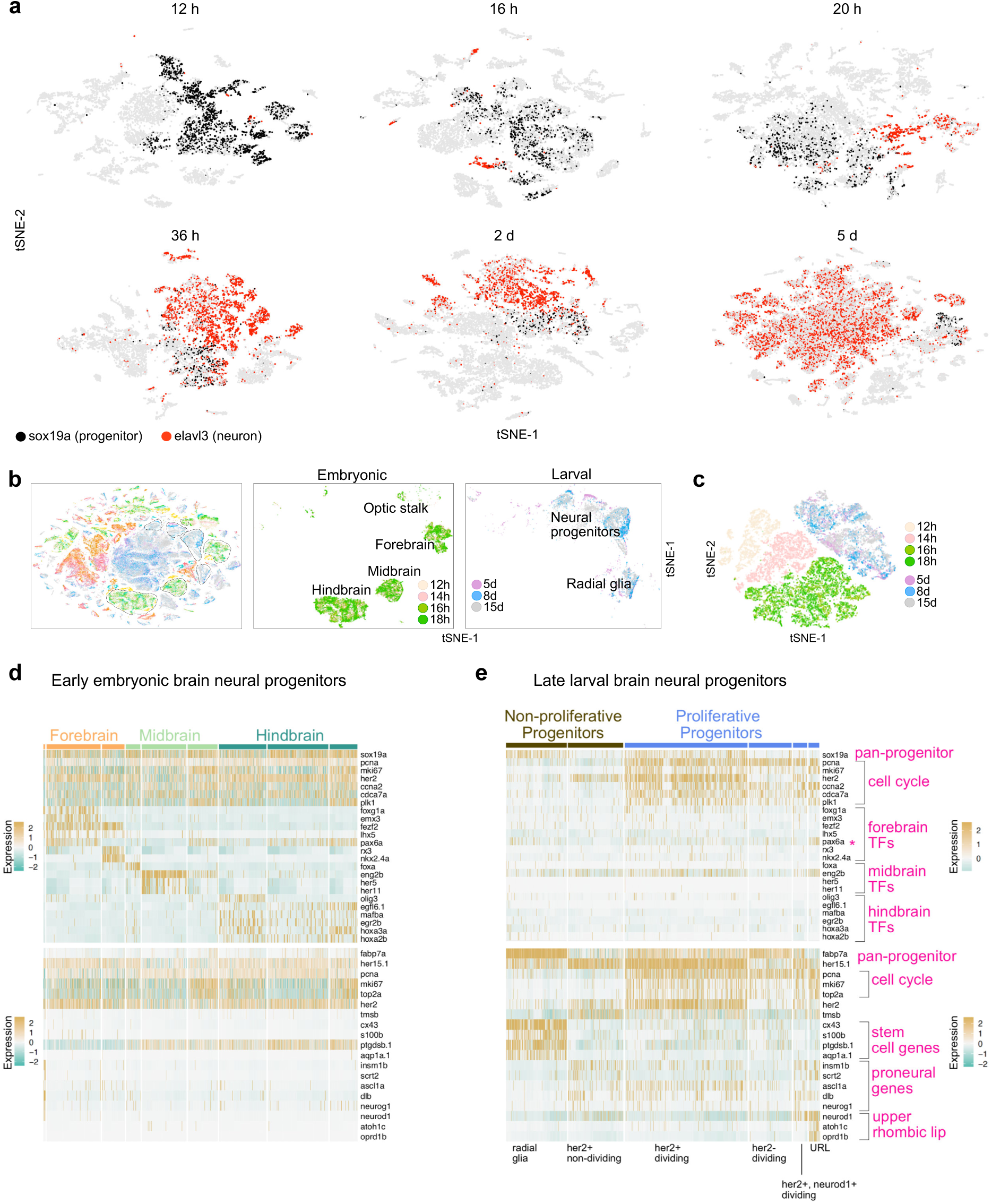
Temporal diversification of neurons and progenitors. **a.** tSNE plots highlight cells expressing sox19a (neural progenitor marker, black) or elavl3 (neuronal marker, red) across various developmental stages. **b.** tSNE plots of cells classified as embryonic (center panel) and larval (right panel) progenitors. Cells were subsetted from the whole dataset atlas (hatched lines, left panel). Cells are color coded by stage. **c.** tSNE plot of embryonic and larval progenitors. All progenitor cells were analyzed together after subsetting from the whole dataset. Developmental time is a strong source of variation, especially for embryonic progenitors. Larval progenitor clusters contain cells from all stages. **d.** and **e**. Heatmaps of select gene expression in early embryonic (**d**) and late larval (**e**) brain neural progenitors. Top panel, genes enriched in embryonic progenitors. Bottom panel, genes enriched in larval progenitors. Embryonic progenitors have a strong spatial signature (forebrain, midbrain, hindbrain) and are depleted in genes that distinguish larval progenitor subtypes (**d**). Larval progenitors segregate into non-proliferative and proliferative groups that can be resolved into additional subtypes characterized by expression of various gene combinations (**e**). TF, transcription factor tSNE implementations: Barnes-Hut (**a** [12h to 2d], **c**), Fourier transform (**a** [5d], **b**)

### Dampening of spatial and developmental signatures during embryonic to larval neural progenitor transition

We next analyzed our dataset to determine how cell states change during the transition from the embryonic to post-embryonic brain. The zebrafish brain undergoes lifetime constitutive neurogenesis due to the persistence of neural progenitor pools distributed along the brain’s axis ^11^. However, the embryonic origins and transcriptional programs that underlie their development are poorly understood. Furthermore, how the molecular identities of embryonic and post-embryonic neural progenitor cell states compare has not been well characterized. To address these questions, we asked how neural progenitor gene expression signatures globally change from embryo to larva. We defined early embryonic progenitors as brain neural cell types from 12 hpf to 18 hpf, and defined larval progenitors as brain neural progenitors from 5 dpf to 15 dpf (Figure 3b, 3c, Sup Fig.7). For both groups, we considered neural progenitors as cells that are non-differentiated precursors that may or may not be proliferating, depending on the expression of cell cycle markers (see below).

Embryonic neural progenitor cells separated into 3 major clusters (Figure 3b). We asked what is the greatest source of variation within these populations, and found that the top 3 principal components comprise genes implicated in spatial and developmental patterning ^7,9,15,16^. Cells exhibit characteristic anteroposterior and dorsoventral axial signatures (Figure 3d). For example, the telencephalon (anterior forebrain) is marked by *foxg1a* and *emx3a* expression, the midbrain by *pax2a* and *eng2a*, and the hindbrain is segmented into rhombomeres marked by distinct combinatorial patterns of *egr2b* and *hox* gene expression. Furthermore, all cells are in a highly proliferative state with strong expression of cell cycle genes such as *pcna, mki67* and *cdca7a*. Collectively, the expression signatures are reflective of a developmental state during which the embryo is orchestrating a rapid expansion of neural progenitor populations concurrent with their acquisition of positional information and overt absence of differentiation ^11,17^. In contrast, larval neural progenitors comprised two major groups: proliferating (expressing cell cycle genes *pcna* and *top2a*) and non-proliferating (depleted expression of cell cycle markers) progenitors (Figure 3e). Indeed, the top 3 principal components in the larval progenitors comprise genes that mark stem cells (PC1, PC3) and differentiation (PC2). The non-proliferating group is subdivided into radial glia (stem cells) and *her2+* neural progenitors expressing proneural genes *insm1b* and *scrt2*. The proliferating group is subdivided into *her2+* and *scrt2-*neural progenitor cells, *her2-*progenitors, *her2+* and *neurod1+* progenitor cells, and upper rhombic lip progenitors (localized to cerebellum) expressing *atoh1c* and *oprd1b*.

Strikingly, most larval progenitors are characterized by a reduced spatial signature (except for the cerebellar upper rhombic lip pool), such that cells are less enriched in region-specific transcription factors relative to embryonic progenitors (Figure 3e). For example, radial glia exist in multiple pools along the brain’s axis ^18^, however, they form a single cluster in our dataset (marked by expression of *fabp7a, cx43, s100b* and *aqp1a.1*), suggesting they are largely transcriptionally similar. Although some expression of region-specific transcription factors is detected in larval progenitor clusters, these signatures are not sufficiently strong to resolve clusters as they were during embryonic stages (Figure 3d, 3e). To explore the apparent dearth of spatial signatures further, we calculated pairwise correlation scores for 79 transcription factors and signaling proteins with known spatial expression patterns in the forebrain and midbrain based on histological analysis (ZFIN). These genes show stronger correlations in embryonic progenitors than in larval progenitors (Sup Fig. 8a). Since spatial signatures are encoded by a combinatorial code of genes with overlapping expression patterns, we asked whether the same subsets of genes co-vary with each of the 79 spatial markers in embryonic and larval neural progenitors, and found low overlap across both stages (4/79 genes had 40% overlap in their top 20 co-varying genes, 17/79 genes had 30% overlap, and 58/79 had <30% overlap). Additionally, when we searched for genes that strongly co-varied with these 79 spatial markers (Pearson correlation >0.4), we found 41 genes during embryonic stages, but only 1 gene during larval stages (Sup Fig. 8b). Thus, the overall spatial code between embryonic and larval progenitors are distinct, and the embryonic spatial code involves a larger collection of genes. Notably, the signatures of larval progenitors resemble juvenile neural progenitor pools ^8^, indicating developmental switches in neural progenitor identities from embryo to larva that are maintained to at least juvenile stages. Thus, embryonic states that existed in early progenitors are largely altered in late-stage progenitors; while spatial patterning signals are the greatest source of variation between embryonic neural progenitors, these signals are dampened in long-term neural progenitors.

### Cell specification trajectories in the retina

With the exception of a few model systems ^19-24^, little is known about the gene expression cascades that accompany the development of progenitors into terminally differentiated neurons. To address how different neuronal populations become molecularly specialized, we traced their gene expression trajectories from 12 hpf to 15 dpf. We first tested our approach on the subsetted retina dataset in which cell types expand from a single cluster at 12 hpf to 18 clusters at 15 dpf (Figure 2). UMAP embedding of the subsetted dataset revealed progressive paths from the embryonic state to defined cell types at 15 dpf, with the exception of one outlier cluster expressing *kidins220a*, whose progenitor state may not have been captured in our timepoints and was excluded from further analysis (Figure 4a, Sup Fig.9). Although UMAP represents continuity in the data, it does not order individual cells according to their relative developmental time (i.e. pseudotime). Therefore, we also used URD ^25^ to construct a branching specification tree that represents the developmental trajectories in the retina at a higher resolution (Figure 4b, Sup Fig.10, 11). Many of the major branching features agreed with the UMAP representation. For example, the trajectories revealed the early segregation of RPE, shared branching of photoreceptor cells, a path towards multiple cone bipolar cell subtypes, and common branchpoint between amacrine and retinal ganglion cells (RGC).

**Figure 4.**
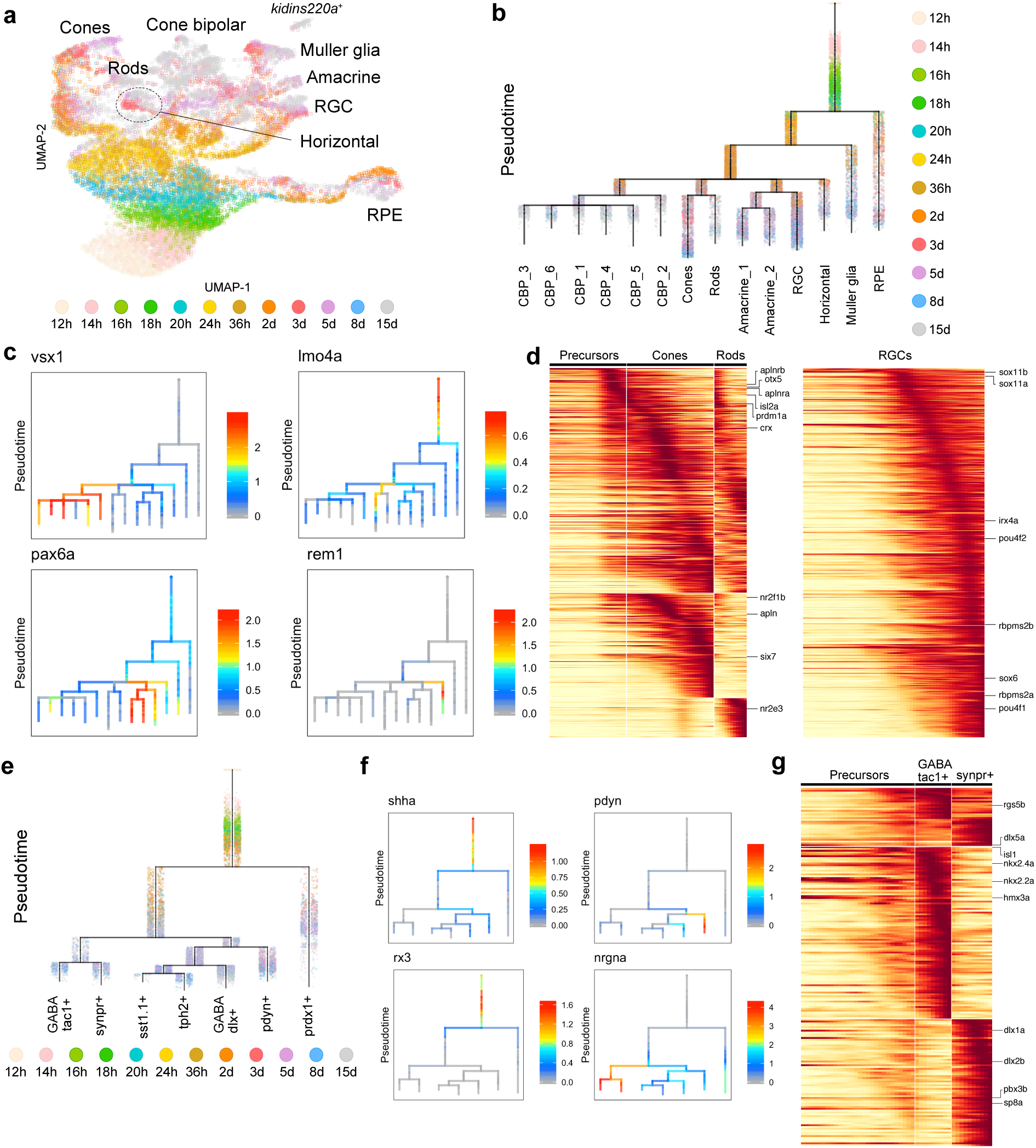
Cell specification trajectories in the retina and hypothalamus. **a.** UMAP visualization of retinal cell types. Retinal cells (based on clustering analysis) from 12 hpf to 15 dpf were subsetted from the full dataset and analyzed together. Cells are color coded by stage. **b.** Cell specification tree of zebrafish retinal development. Trajectories were generated by URD and visualized as a branching tree. Cells are color coded by stage. 12 hpf cells were assigned as the root and 15 dpf differentiated cells were assigned as tips. CBP, cone bipolar cells (6 subtypes are numbered); RGC, retinal ganglion cells; RPE, retinal pigment epithelium. **c.** Expression of select genes are shown on the retina specification tree. **d.** Heat maps of gene expression cascades of photoreceptor cell trajectories and retinal ganglion cell trajectories. Cells were selected based on high expression along trajectories leading to these cell types, compared to expression along opposing branchpoints. Red, high expression. Yellow, low expression. **e.** Cell specification tree of zebrafish hypothalamus development. Trajectories were generated by URD and visualized as a branching tree. Cells are color coded by stage. 12 hpf cells were assigned as the root and 15 dpf differentiated cells were assigned as tips. **f.** Expression of select genes are shown on the hypothalamus specification tree. **g.** Heat map of gene expression cascade of *nrgna+* cell trajectories. Red, high expression. Yellow, low expression

Plotting gene expression of known early regulators of eye development and terminal cell type markers on the URD tree supported the inferred specification branches (Fig. 4c, Sup Fig.11). For example, *pax6a* was most enriched in the amacrine and RGC branches, *vsx1* marked cone bipolar cells with *fezf2* marking one specific subtype. Notably, our analysis also revealed previously unknown markers and characteristics of horizontal and amacrine cells. Zebrafish horizontal cells are GABAergic (*gad2+, gad1b+*), but unlike mammals where these cells do not express GABA membrane uptake transporters ^26^, zebrafish cells express *slc6a1l* (likely a duplication of *slc6a1* involved in GABA uptake from the synaptic cleft) suggesting that they may be capable of uptake. Additionally, whereas *slc32a1* GABA transporter is expressed in mouse horizontal and amacrine cells ^27^, we observed restriction of *slc32a1* to amacrine cells and *slc6a1l* to horizontal cells. Finally, we detected several novel horizontal cells markers such as *ompa, rem1*, and *prkacaa*.

To discover the gene expression trajectories from precursors to different retinal cell types, we used differential gene expression approaches that characterize pseudotime-ordered molecular trajectories. This analysis revealed known and novel regulatory steps Fig. 4d, Sup Fig.12). For example, RGC specification trajectories confirmed several known differentiation regulators including *sox11a, sox11b, sox6, irx4a*, and *pou4f2* ^28^. Similarly, known regulators of photoreceptor differentiation, such as *isl2a* ^29^, *prdm1a* ^30^, *otx5* ^31^, and *crx* ^32^ are expressed early in our photoreceptor trajectories, while known regulators of cone versus rod fate, such as *six7* ^33^, *nr2f1b* ^34^, and *nr2e3* ^35^ are expressed as those trajectories diverge. Furthermore, our analysis revealed novel transcription factors within the gene expression cascades. For example, we detected *runx1t1, foxp1b, mef2aa* in the RGC pathway; *tfap2a* in horizontal cell trajectory; and *tbx3a* and *tbx2a* in amacrine cell branches. Interestingly, among signaling pathways, we found that both apelin receptors (*aplnra, aplnrb*) are expressed in photoreceptor progenitors, while their ligand (*apln*) is expressed in differentiating cones; this suggests a potential cell autonomous role for apelin signaling in photoreceptor cells in addition to its role in preventing photoreceptor degeneration via vascular remodeling ^36^.

A surprising result from this analysis was that Muller glia arise much earlier in zebrafish than expected based on studies in mouse, where these cells are the last to arise ^19,37^. We found a cluster of cells as early as 20 hpf (cluster 50) that expresses gene markers (e.g. *cahz, rlbp1a*) that are shared with the Muller glia cluster (cluster 33) at 15 dpf (Sup Table). Similarly, in our transcriptional trajectories (Fig. 4b), the Muller glia expression program is the earliest non-epithelial retinal program to diverge, commencing with the expression of several *her*-family transcription factors (*her4, her12*, and *her15*), then proceeding through a cascade of intermediate overlapping expression states such as onset of *fabp7a, s100a10b* and later connexin genes that are characteristic of Muller glia fate (Sup Fig.12). This suggests that cells early in development transition from a naive progenitor state to a Muller glia-like transcriptional state, and do so continually during larval development (cells from all timepoints can be found in the early part of the Muller glia branch).

### Cell specification trajectories in the hypothalamus

To extend our analysis, we reconstructed specification trajectories and expression cascades for hypothalamic cell types, which expand from a single ventral diencephalon cluster at 12 hpf to 7 clusters at 15 dpf (Fig.4e-g, Sup Fig. 13, 14). The earliest branchpoint denotes segregation of *prdx1*+ and *prdx1-*cells. Committed hypothalamic progenitors in the *prdx1-*trajectory give rise to neuronal precursors expressing proneural transcription factors such as *ascl1a, scrt2, insm1a* and *elavl3* (early neuron fate marker) (Sup Fig. 13). Next, the specified cell types mature over time, and are characterized by expression of neuronal maturation markers such as *tubb5, gap43, ywhag2, snap25a, scg2b* and *elavl4*. The *prdx1-*group further diverges into two major groups: *nrgna+* and *nrgna-*trajectories (Fig. 4e). The *nrgna+* branch segregated into GABAergic *tac1+, synpr-*subtype and GABAergic *tac1+, synpr+* positive subtype. The *nrgna-*branch subdivided into glutamatergic *pdyn+* neurons and a GABAergic branch that further resolved to *sst1.1+* and *tph2+* neuron subtypes. We detected expression of known regulators of hypothalamus development in the early branches such as *shha, rx3, nkx2.4b*. We also identified new candidate regulators in later branches including *nrgna* in the *synpr+* and *synpr-*trajectories; and *sox1a, sox1b* and *sox14* in the *pdyn+* trajectory (Sup Fig. 14).

### Differences in progenitor specification strategies between retina and hypothalamus

Continuous differentiation throughout the animal’s lifetime is a feature of zebrafish neurogenesis ^11^. Pseudotime analysis represents cell trajectories in relative but not absolute time; it models transcriptional progression rather than time progression ^38,39^. When we mapped developmental stage onto the URD trees (Fig. 4b, 4e), we found that real-time progression and pseudotime progression generally agreed, with some notable exceptions. For example, retinal ganglion cells (RGCs) are one of the earliest retinal neurons to be born. Our clustering analysis identified an RGC cluster as early as 36 hpf (cluster 27); interestingly plotting cells from this cluster onto the URD tree revealed that many cells were assigned late pseudotime with other differentiating RGC cells from later stages (Fig. 5a), indicating that RGCs differentiate continuously across several developmental stages. Conversely, by plotting retinal progenitors (cluster 39) from 15 dpf, we observed that most cells are assigned to early branches of the tree with retinal progenitors from prior stages (Fig. 5b). We therefore compared late (15 dpf) and early (24-36 hpf) progenitors, and found that there were only 71 differentially expressed genes between them. The majority of these genes (56) increased in all cells of the retina between these stages, while a few (15) were only upregulated in 15 dpf retinal progenitors. This suggests that retinal progenitors at 15 dpf are transcriptionally similar to ones from earlier stages of development, and that in some tissues, progenitor cell states observed in the embryo persist later in development in some cases without maturing significantly.

**Figure 5.**
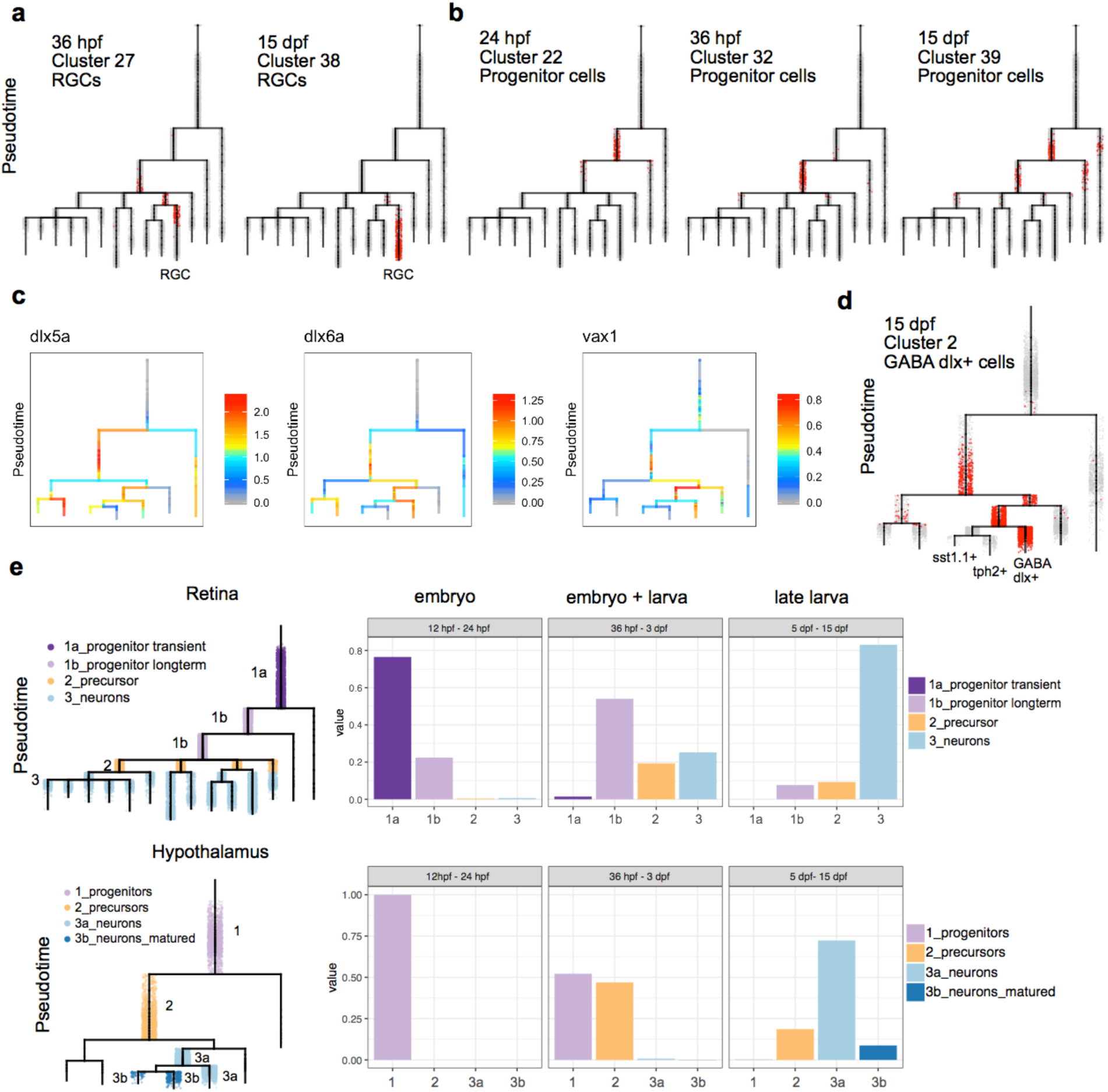
Progenitor differentiation in retina and hypothalamus. **a.** Retinal ganglion cells (RGC, red dots) from indicated stages and clusters are plotted onto the retina specification tree. A subset of 36 hpf RGCs are assigned late pseudotime and are found on the terminal RGC branch, likely representing terminally differentiated cells. **b.** Retinal progenitor cells (red dots) from indicated stages and clusters are plotted onto the retina specification tree. Subsets of long-term progenitor cells from 15 dpf are assigned early pseudotime and are found on earlier branches on the tree that also contain embryonic progenitor cells from 24-36 hpf. **c.** Expression of select genes are shown on the hypothalamus specification tree. **d.** GABAergic dlx+ cells (red dots) from 15 dpf are plotted onto the hypothalamus specification tree. Cells are found across multiple early and late branches of the tree. **e.** Retinal and hypothalamus cells were divided into progenitor (purple), precursor (orange), and neuronal (blue) transcriptional states, as shown on the URD tree. The fraction of cells in each of these transcriptional states was then determined for three developmental periods (12–24 hpf, 36 hpf – 3 dpf, and 5–15 dpf). In the retina, cells can be found in a progenitor state (light purple) across all developmental stages. In the hypothalamus, the embryonic progenitor state is transient and gone by 5 dpf, but cells can be found in a precursor state (orange) late in development.

In the hypothalamus, we also observed pre-neuronal cell states occupied by cells from numerous stages; like the retina, these likely represent undifferentiated cells that continuously differentiate over time. For instance, we observe a hypothalamic GABAergic neuronal cluster (GABA *dlx+*) at 15 dpf that is enriched in *dlx* and neuronal marker gene expression (e.g. *elavl4, scg2b*), but is not enriched in expression of mature terminal identity markers such as neuropeptides, signaling molecules and additional neurotransmitters (Fig. 5c). GABA *dlx+* 15 dpf cells are found across multiple early and late branches on our tree (Fig. 5d). This suggests that this subtype represents a heterogenous cell state pool that persists to late larval stages, and from which subsets of cells further mature into at least 2 additional neuronal subtypes (*sst1.1+* and *tph2+*); profiling cells beyond 15 dpf may reveal additional subtypes that derive from the GABA *dlx+* pool.

The data revealed distinct progenitor differentiation strategies between the retina and hypothalamus. In the retina, neural progenitor states observed in the embryo persist to late larval stages (Fig. 5e, Sup Fig. 15). In contrast, in the hypothalamus, the embryonic progenitor state (expressing *rx3, shha*) is not observed in late larval stages. Instead, non-terminally differentiated cells exist in precursor states that express a mix of progenitor (e.g. *insm1a, her4.1*) and early neuronal (e.g. *tubb5, gap43*) marker genes. Cells then differentiate into neurons and further mature by expressing terminal fate markers. These results indicate that different progenitor specification and neurogenesis strategies are used between different brain regions during the establishment of neuronal diversity.

## DISCUSSION

As the brain develops, embryonic neural progenitor pools transition through many cellular states as they become more committed, diversify into longer lasting post-embryonic neural progenitor pools, and undergo terminal differentiation. Although many regulators and transcriptional changes of this process have been identified (e.g. using specific driver lines and *in situ* expression of select genes), the global transcriptional networks mediating the sequential activation and maturation of neurogenic programs from embryo to later stages are largely unknown. To address this question, we generated a zebrafish brain developmental atlas encompassing cells from 12 hpf to 15 dpf. These data currently constitute the most extensive resource to identify novel marker genes, compare cell types, and determine cell specification and differentiation trajectories during vertebrate brain development.

Our data address how the transcriptional programs of neural progenitors vary and contribute to fate-restriction during development. Different models to explain these processes have been proposed. For example, neural progenitors of the medial and lateral mouse ganglionic eminence, which give rise to cortical interneurons, have been found to converge to a shared mitotic signature regardless of their region of origin, followed by expression of cardinal fate-specific transcription factors post-mitotically ^40^. In contrast, the spinal cord has dedicated pools of domain-specific neural progenitors that retain domain-specific signatures ^21,41,42^. Our results indicate that early embryonic neural progenitors in the brain are transcriptionally distinct from late larval neural progenitors. Gene expression profiles of neural progenitors switch from strong spatially segregated signatures in early embryos to proliferative and non-proliferative states in late larvae. We find that the greatest sources of variation between larval progenitors are stem cell versus non-stem cell, and proliferative versus non-dividing marker genes. These cell state changes might reflect developmental shifts from a migratory program during gastrulation, where strong spatial patterning cues set up regional boundaries, to a maintenance program at late stages, where progenitors are geographically confined and express dampened regional restriction signatures. Although expression of some spatially-enriched transcription factors (e.g. *pax6a, eng2a, nkx2.4a*) and signaling proteins detected in embryonic progenitors is also detected in late long-term progenitors, the overall signatures are different: these factors co-vary with different sets of genes in larva relative to embryo. This raises the question of how neural progenitor pools remain fate restricted at later stages. It is conceivable that embryo and larva share a minimal core set of regionally-restricted transcription factors that ensure spatial restriction, despite differences in their relative expression levels and downstream targets. Spatial genes that are highly expressed in the embryo may be lowly expressed in the larva, and be sufficient to maintain regionally-restricted cell states. Such signatures would be difficult to analyze via scRNA-seq, which is biased towards recovering highly expressed genes. It is also possible that restrictions at the genomic level, such as chromatin accessibility, may ensure that cells maintain the signature of their spatial origin. Fate mapping experiments of early and late neural progenitors, future studies profiling open chromatin states of neural progenitors during development, and profiling of cells with approaches that more effectively recover lowly expressed genes will provide further insight into these questions.

Our reconstruction of cell specification trajectories for cell types in the retina and hypothalamus revealed several unexpected findings. First, we detected a Muller glia transcriptional program early in development, prior to any of the retinal neuron-specific programs. This observation differs from what has been described in mouse, where these cells are born last ^19,37^. Thus, our early detection of a Muller glia transcriptional state could suggest that Muller glia specification happens earlier in zebrafish than other organisms. Alternatively, cells could enter and exit this state during early development, and truly commit to Muller glia fate only later in development. Second, our data highlights how constitutive neurogenesis in the zebrafish brain results in transcriptional cascades that are repeatedly activated in cells during late stages of our time course. Late pseudotime branches in our cell specification trees contain cells from all larval stages (5 dpf to 15 dpf) indicating that cells can be found in the same molecular state across large windows of time; thus, as several cell types acquire terminal fate identities during constitutive neurogenesis, it seems that they undergo the same sequence of molecular events regardless of their time of birth. These cascades can be further examined to determine candidate genes that may be critical for constitutive neurogenesis. Third, our results reveal that cells in the hypothalamus exist in long-term neural precursor states that are distinct from cell states observed in the embryo. In contrast, retinal cells exist in long-term progenitor states that resemble states observed in the embryo. This raises the intriguing possibility that a subset of long-term retinal progenitors may be “frozen” in an embryonic phase that could possibly underlie multi-fate potential of these cells. In contrast, we hypothesize that hypothalamus precursors may reflect more committed states prior to neuronal differentiation into specific subtypes. Collectively, these findings highlight differences in cell differentiation strategies of progenitors across different regions of the brain, and underscore the power of investigating multiple specification trajectories simultaneously.

Our analysis has laid the groundwork for characterizing transcriptional changes underlying development and diversification of a vertebrate brain. This gene expression atlas can be used to generate transgenic reporters using identified markers to select populations of interest and perform deeper analysis of cell type heterogeneity and differentiation ^43^. Furthermore, cell specification trajectories can be extended to include additional subregions of the brain to generate increasingly complex trees. Our dataset may be combined with early zebrafish embryogenesis scRNA-seq datasets ^25,44^ to trace trajectories from stages prior to gastrulation to late larval stages. Furthermore, cell specification trajectories and lineage trees can be compared vis-à-vis to investigate cell intrinsic effects on lineage decisions across many cell types and brain regions. Higher recovery of cells and enhanced time resolution in lineage trees are required for such direct comparisons. We optimized our scGESTALT recorder and library protocol to enable more dense reconstruction of lineage trees. Although our current implementation of this version was restricted to marking lineages in the early embryo, mostly prior to neuronal subtype segregations, it can be readily adapted for recording at later stages, for example by editing across multiple developmental windows ^8^. In the future, combination of single-cell transcriptomics, chromatin accessibility and lineage can be used to obtain an integrated understanding of the regulatory logic of how neuronal complexity is established.

## METHODS

### Zebrafish husbandry

All vertebrate animal work was performed at the facilities of Harvard University, Faculty of Arts & Sciences (HU/FAS). This study was approved by the Harvard University/Faculty of Arts & Sciences Standing Committee on the Use of Animals in Research & Teaching under Protocol No. 25–08. The HU/FAS animal care and use program maintains full AAALAC accreditation, is assured with OLAW (A3593-01), and is currently registered with the USDA.

### Optimization of scGESTALT lineage cassette

In our previous iteration of scGESTALT, the barcode capture rate by scRNA-seq was 6-28%. ^8^, thereby limiting the density of lineage tree reconstruction. To improve recovery we adapted a different transgenic cassette ^45^ for lineage recording. This cassette has the following modifications compared to our previous recorder: (1) The heat-shock inducible (hsp70l) promoter of the previous version is now replaced with a constitutive ubiquitous promoter (medaka beta-actin) to drive strong widespread expression of the barcode mRNA. Expression of the cassette was confirmed by fluorescence and the signal was more intense than that obtained with the heat shock promoter. Furthermore, this version eliminates the requirement to heat shock edited animals to express the barcode prior to scRNA-seq experiments. (2) We adapted the 3’ end of the DsRed open reading frame as a lineage recorder cassette with up to 8 sgRNA target sites positioned next to each other. This vastly improved expression of the construct compared to our previous version where the recording cassette was placed downstream of the DsRed open reading frame. (3) We made library preparation compatible with the 10X Genomics platform.

To generate scGESTALT.2 barcode founder fish, one-cell embryos were injected with zebrafish codon optimized Tol2 mRNA and pT2Olactb:loxP-dsR2-loxP-EGFP vector (gift from Atsushi Kawakami ^45^). Potential founder fish were screened for widespread DsRed expression and grown to adulthood. Adult founder transgenic fish were identified by outcrossing to wild type fish and screening clutches of embryos for ubiquitous DsRed expression. Single copy scGESTALT.2 F1 transgenics were identified using qPCR, as described previously ^8,10,46^.

SgRNAs specific to sites 1-8 of the scGESTALT.2 array were generated by in vitro transcription as previously described ^47^. To initiate early barcode editing, single copy scGESTALT.2 F1 male transgenic adults were crossed to wildtype female adults and one-cell embryos were injected with 1.5 nl of Cas9 protein (NEB) and sgRNAs 1-8 in salt solution (8 µM Cas9, 100 ng/µl pooled sgRNAs, 50 mM KCl, 3 mM MgCl_2_, 5 mM Tris HCl pH 8.0, 0.05% phenol red). Since editing results in loss of DsRed signal, transgenic animals were distinguished from wild type animals by amplifying the scGESTALT.2 barcode by PCR using genomic DNA from the tail fin at 15 dpf.

### Processing of samples for scRNA-seq time course

Wild type embryos (12 hpf, 14 hpf, 16 hpf, 18 hpf, 20 hpf, 24 hpf, 36 hpf) and larvae (2 dpf, 3 dpf, 5 dpf, 8 dpf) were used for scRNA-seq analysis. Samples for 15 dpf had a mix of wild type and barcode edited larvae. Embryos from 12 hpf to 36 hpf were first de-chorionated by incubating in 1 mg/ml pronase (Sigma-Aldrich) at 28 C for 6-7 min until chorions began to blister, and then washed three times in ∼200 ml of zebrafish embryo medium (5 mM NaCl, 0.17 mM KCl, 0.33 mM CaCl_2_, 0.33 mM MgSO_4_, 0.1% methylene blue) in a glass beaker. Embryos were de-yolked using two pairs of watchmaker forceps, and the heads were chopped just anterior of the spinal cord. All processing steps were done using 100 mm Petri dishes coated with Sylgard ^47^. Samples from 2 and 3 dpf were processed similarly to the embryos, except they were not de-chorionated as they had hatched out of the chorions. Larvae from 5 dpf to 15 dpf were dissected to remove whole brains and eyes as described previously ^47^. The following numbers of embryos and larvae were used for each timepoint: 12 hpf – ∼20 embryos; 14 hpf – ∼20 embryos; 16 hpf – ∼18 embryos; 18 hpf – ∼18 embryos; 20 hpf – ∼30 embryos; 24 hpf – ∼30 embryos; 36 hpf – ∼15 embryos; 2 dpf – ∼30 larvae; 3 dpf – ∼30 larvae; 5 dpf – ∼25 larvae; 8 dpf – ∼ 25 larvae; 15 dpf – ∼15 larvae. Tissues were dissociated into single cells using the Papin Dissociation Kit (Worthington) as described previously ^47^. Cells were resuspended in 50 µl to 150 µl of DPBS (Life Technologies) depending on anticipated amount of material, and counted using a hemocytometer. Samples were run on the 10X Genomics scRNA-seq platform according to the manufacturer’s instructions (Single Cell 3’ v2 kit). Libraries were processed according to the manufacturer’s instructions. Transcriptome libraries were sequenced using Nextera 75 cycle kits.

### scGESTALT.2 library prep

To generate scGESTALT.2 libraries, lineage edited 15 dpf samples post cDNA amplification and prior to fragmentation were split into two halves. One half was processed for transcriptome libraries as instructed by the manufacturer. The other half was processed for lineage libraries as follows. To enrich for scGESTALT.2 lineage barcodes, 5 µl of the whole transcriptome cDNA was PCR amplified using Phusion polymerase (NEB) and 10XPCR1_F (CTACACGACGCTCTT CCGATCT) and GP10X2_R (GTGACTGGAGTTCAGACGTGTGCTCTTCCGATCT GCTGCTTC ATCTACAAGGTGAAG). The reaction (98 C, 30 s; [98 C, 10 s; 67 C, 25 s; 72 C, 30 s] x 14-15 cycles; 72 C, 2 min) was cleaned up with 0.6X AMPure beads and eluted in 20 ul EB buffer (Omega). Finally, adapters and sample indexes were incorporated in another PCR reaction using Phusion polymerase and 10XP5Part1long (AATGATACGGCGACCACCGAGATCTACACTCTTTCC CTACACGACGCTCTTCCGATCT) and 10XP7Part2Ax (CAAGCAGAAGACGGCATACGAGAT-xxxxxxxx-GTGACTGGAGTTCAGACGTGT), where x represents index bases. These include A1: GGTTTACT; A2: TTTCATGA; A3: CAGTACTG; A4: TATGATTC. Thus, up to 4 scGESTALT.2 samples were multiplexed in a sequencing run. Libraries were sequenced using MiSeq 300 cycle kits and 20% PhiX spike-in. Sequencing parameters: Read1 250 cycles, Read2 14 cycles, Index1 8 cycles, Index2 8 cycles. Standard sequencing primers were used.

### Bioinformatic processing of raw sequencing data and cell type clustering analysis

Transcriptome sequencing data were processed using Cell Ranger 2.1.0 according to the manufacturer’s guidelines. scGESTALT.2 sequencing data were processed with a custom pipeline (https://github.com/aaronmck/SC_GESTALT) as previously described ^8^. The scGESTALT.2 barcode for each cell was matched to its corresponding cell type (tSNE cluster membership) assignment using the cell identifier introduced during transcriptome capture. Clustering analysis was performed using the Seurat v2.3.4 package ^48^ as described previously ^8^. Full analysis scripts for cell type clustering, R objects and raw sequencing data will be uploaded to GitHub and NCBI GEO.

### Construction of lineage trees from GESTALT barcodes

All unique barcodes were then encoded into an event matrix and weights file, as described previously ^8,10^, and were processed using PHYLIP mix with Camin-Sokal maximum parsimony ^49^. Individual cells were then grafted onto the leaves matching their barcode sequence. After the subtrees were attached, we repeatedly eliminated unsupported internal branching by recursively pruning parent-child nodes that had identical barcodes. Cell annotations are then added to the corresponding leaves. The resulting tree was converted to a JSON object, annotated with cluster membership, and visualized with custom tools using the D3 software framework.

### Analyzing dampened spatial correlations in progenitors

Progenitors were isolated by subsetting the data to include clusters expressing markers such as *sox19a, her* genes, *pcna, mki67, fabp7a, gfap, id1*, etc (Supplementary Table). Cells from 12 hpf – 18 hpf were considered embryonic progenitors and cells from 5 dpf – 15 dpf were considered larval progenitors. Variable genes were calculated for embryonic and larval progenitors separately using the FindVariableGenes function from Seurat v2.3.4 with parameters: *x.low.cutoff* = 0.015, *x.high.cutoff* = 3, *y.cutoff* = 0.7. Then, a list of 79 transcription factors with known spatial signatures was assembled by ***. Separately in the embryonic and larval progenitors, the pairwise Pearson correlation was calculated pairwise between all genes detected as variable in either the embryonic or larval progenitors. For several thresholds between 0.2–0.8, the number of genes that correlated more strongly than the threshold with any of the 79 spatial transcription factors (excluding self-correlation) were determined. The strongest correlations were observed in the embryonic population, and for any threshold, more genes correlated with the spatial TFs in the embryonic progenitors than the larval progenitors.

### Construction and analysis of branching transcriptional trajectories using URD

We built branching transcriptional trajectories from cells of the retina and hypothalamus to determine the molecular events that occur as cells diversify and differentiate in these tissues. First, cells from the retina and hypothalamus were isolated from each stage by determining clusters that belonged to these tissues by expression of marker genes.

#### Determination of variable genes

For URD trajectory analyses, a more restrictive set of variable genes was calculated on each subset of the data, as previously described ^25,43^ using the URD *findVariableGenes* function, with parameter *diffCV.cutoff* = 0.3. Briefly, a curve was fit that related each gene’s coefficient of variation to its mean expression level and represents the expected coefficient of variation resulting from technical noise, given a gene’s mean expression value; genes with much higher coefficients of variation likely encode biological variability and were used downstream.

#### Removal of outliers

Poorly connected outliers can disrupt diffusion map calculation and so were removed from the data. A *k*-nearest neighbor network was calculated between cells (Euclidean distance in variable genes) with 100 nearest neighbors. Cells were then removed based on either unusually high distance to their nearest neighbor or unusually high distance to their 20^th^ nearest neighbor, given their distance to their nearest neighbor using the URD function *knnOutliers* (retina: *x.max* = 40, *slope.r* = 1.05, *int.r* = 4.3, *slope.b* = 0.75, *int.b* = 11.5; hypothalamus: *x.max* = 40, *slope.r* = 1.1, *int.r* = 3, *slope.b* = 0.66, *int.b* = 11.5).

#### Removal of doublets by NMF modules

To remove putative cell doublets (i.e. where two cells are encapsulated into a single droplet and processed as one cell), which can disrupt trajectory relationships, we removed cells that expressed multiple NMF (non-negative matrix factorization) modules characteristic of different expression programs, as previously described ^50^. NMF modules were computed using a previously published NMF framework (https://github.com/YiqunW/NMF) ^25^. The analysis was performed on log-normalized read count data for a set of variable genes using the *run_nmf.py* script with the following parameters: -rep 5 -scl “false” -miter 10000 -perm True -run_perm True - tol 1e-6 -a 2 -init “random” -analyze True. Several *k* parameters were evaluated for each tissue, and k was chosen to maximize the number of modules, while minimizing the proportion of modules defined primarily by a single gene (retina, *k* = 45; hypothalamus, *k* =). Modules were used downstream that (a) had a ratio between their top-weighted and second-highest weighted gene of < 5, and (b) exhibited a strong cell-type signature, as determined by plotting on a UMAP representation and looking for spatial restriction. Pairs of modules that were appropriate for using to remove doublets (and that did not define transition states) were determined using the URD function *NMFDoubletsDefineModules* with parameters *module.thresh.high =* 0.4, and *module.thresh.low* = 0.15. Putative doublets were identified using the URD function *NMFDoubletsDetermineCells* with parameters *frac.overlap.max* = 0.03, *frac.overlap.diff.max* = 0.1, *module.expressed.thresh* = 0.33 and were then removed.

#### Choice of root and tips

Branching transcriptional trajectories in the retina and hypothalamus were constructed using URD 1.1.1 (Farrell 2018). Briefly, cells from the first stage of the time course (12 hpf) were selected as the ‘root’ or starting point for the tree. Terminal cell types comprised the clusters at 15 dpf from these tissues, with the exception of clusters that were clearly progenitor or precursors based on known gene expression (retina: 29, 39, 43). Additionally, in the retina, one cluster (96) was excluded because it did not seem that any related cell types had been recovered in previous stages.

#### Construction of branching transcriptional trajectories

A diffusion map was calculated using *destiny* ^51,52^, using 140 (retina) or 100 (hypothalamus) nearest neighbors (approximately the square root of the number of cells in the data), and with a globally-defined sigma of 14 (retina) or 8 (hypothalamus) — slightly smaller than the suggested sigma from *destiny*. Pseudotime was then computed using the simulated ‘flood’ procedure previously described ^25^, using the following parameters: *n* = 100, *minimum.cells.flooded* = 2. Biased random walks were performed to determine the cells visited from each terminal population in the data as previously described ^25^, using the following parameters: *optimal.cells.forward* = 40, *max.cells.back* = 80, *n.per.tip* = 50000, *end.visits* = 1. The branching tree was then constructed using URD’s *buildTree* function with the following parameters: *divergence.method* = “ks” (hypothalamus) or *divergence.method* = “preference” (retina), *save.all.breakpoint.info* = TRUE, *cells.per.pseudotime.bin* = 40, *bins.per.pseudotime.window* = 5, *p.thresh* = 0.0001 (hypothalamus) or, *p.thresh* = 0.01 (retina), and *min.cells.per.segment* = 10. The resulting trees were then evaluated using known marker genes and branch regulators.

#### Finding genes that vary during differentiation

Genes were selected for inclusion in gene cascades based on their differential expression relative to other cell types in the tissue. See the Supplementary Analysis for the full set of commands used. Within each tissue, cells were first compared in large populations that defined major cell types (retina: cone bipolar cells, photoreceptors, amacrine cells, retinal ganglion cells, horizontal cells, Muller glia, retinal pigmented epithelium; hypothalamus: *prdx1*+ neurons, *pdyn*+ neurons, GABAergic *dlx*+ neurons, *nrgna*+ neurons). Comparisons were performed pairwise, and genes were considered differential in a population if they were upregulated compared to at least 2 (hypothalamus) or 3 (retina) other groups. Genes were considered differentially expressed based on their expression fold-change (retina: ≥1.32-fold change, hypothalamus: ≥ 1.41-fold change) and their performance as a precision-recall classifier for the two cell populations compared (≥ 1.1-fold better than a random classifier). Additionally, the *aucprTestAlongTree* function from URD was used to select additional genes by performing pairwise comparisons, starting from a terminal cell type and comparing at each branchpoint along the way, back to the root ^25^. Genes were selected based on expression fold-change between branchpoints (hypothalamus: ≥1.74-fold upregulated; hypothalamus, populations with small cell numbers (GABAergic *dlx*+ cells): ≥1.51-fold upregulated; retina: ≥1.32-fold upregulated), their function as a precision-recall classifier between branchpoints (hypothalamus: ≥1.2-fold better than a random classifier; hypothalamus, populations with small cell numbers (GABAergic *dlx*+ cells): ≥1.15-fold better than a random classifier; retina: ≥1.1-fold better than a random classifier), their function as a precision recall classifier globally (i.e. between the entire trajectory leading to a cell type and the rest of the tissue): ≥1.03-fold better than a random classifier, and their upregulation globally (i.e. between the entire trajectory leading to a cell type and the rest of the tissue): ≥1.07-fold upregulated. Mitochondrial, ribosomal, and tandem duplicated genes were excluded. Cells were ordered according to pseudotime, split into groups of at least 25 cells that differ at least 0.005 in pseudotime, and the mean expression was determined with a 5-group moving window. A spline curve was fit to the mean expression vs. pseudotime relationship of selected genes, using the *smooth.spline* function from R’s *stats* package, with the parameter *spar* = 0.5. Genes were then sorted according to their peak expression in pseudotime, normalized to their max expression observed in the tissue, and plotted on a heatmap.

#### Analyzing progenitor populations

To determine whether retinal progenitors mature transcriptionally over time, we looked for genes that were differentially expressed between young and old progenitors. We chose cells that occupied the same region of the URD tree from either early (24 / 36 hpf) or late (15 dpf) stages. We looked for genes that were differentially expressed in 15 dpf progenitors that: (1) were 1.1-fold better as a precision-recall classifier than random, (2) changed ≥1.32-fold in expression, (3) were expressed in at least 20% of progenitors, (4) had a mean expression value ≥ 0.8, and (5) were more differentially expressed than equally sized cell populations chosen at random at least 99% of the time.

To determine whether cells were found in progenitor or precursor states long-term, we first defined progenitor and precursor states by cells’ assignment in the URD tree, cross-referenced with the expression of progenitor / precursor markers. We then determined how many cells from different stages fell into each of these different states.

## Supporting information

Supplemental Analysis

## ACKNOWLEDGEMENTS

We thank members of the Schier lab for discussion and advice, the Bauer Core Facility (Harvard) and the Molecular Biology Core Facility (Dana Farber Cancer Institute) for sequencing services, and the Harvard zebrafish facility staff for technical support. We thank M. Shafer and J. Gagnon for comments on the manuscript. This work was supported by a postdoctoral fellowship from the Canadian Institutes of Health Research and 1K99HD098298 to B.R., 1K99HD091291 to J.A.F., NIH grant R00HG010152 to A.M., NIH grants R01HD85905 and DP1HD094764 to A.F.S., and an Allen Discovery Center grant and McKnight Foundation Technological Innovations in Neuroscience Award to A.F.S.

## AUTHOR CONTRIBUTIONS

B.R. and A.F.S. conceived and designed the study. B.R., J.A.F. and A.F.S. interpreted the data and wrote the manuscript. B.R. and J.L.L. generated transgenic lines. B.R. performed scRNA-seq and scGESTALT experiments and data processing. J.A.F. performed URD trajectory analysis with assistance from B.R. B.R. and J.A.F. performed data analysis. A.M. generated lineage trees.

## REFERENCES

1. Woodworth, M. B., Girskis, K. M. & Walsh, C. A. Building a lineage from single cells: genetic techniques for cell lineage tracking. Nat. Rev. Genet. 18, 230–244 (2017).

2. Kretzschmar, K. & Watt, F. M. Lineage tracing. Cell 148, 33–45 (2012).

3. Cepko, C. Intrinsically different retinal progenitor cells produce specific types of progeny. Nat. Rev. Neurosci. 15, 615–627 (2014).

4. Woo, K. & Fraser, S. E. Order and coherence in the fate map of the zebrafish nervous system. Development 121, 2595–2609 (1995).

5. Ma, J., Shen, Z., Yu, Y.-C. & Shi, S.-H. Neural lineage tracing in the mammalian brain. Curr. Opin. Neurobiol. 50, 7–16 (2017).

6. Wamsley, B. & Fishell, G. Genetic and activity-dependent mechanisms underlying interneuron diversity. Nat. Rev. Neurosci. 18, 299–309 (2017).

7. Wilson, S. W., Brand, M. & Eisen, J. S. Patterning the zebrafish central nervous system. Results Probl Cell Differ 40, 181–215 (2002).

8. Raj, B. et al. Simultaneous single-cell profiling of lineages and cell types in the vertebrate brain. Nat Biotechnol 36, 442–450 (2018).

9. Moens, C. B. & Prince, V. E. Constructing the hindbrain: insights from the zebrafish. Dev. Dyn. 224, 1–17 (2002).

10. McKenna, A. et al. Whole-organism lineage tracing by combinatorial and cumulative genome editing. Science 353, aaf7907 (2016).

11. Schmidt, R., Strähle, U. & Scholpp, S. Neurogenesis in zebrafish - from embryo to adult. Neural Development 8, 3 (2013).

12. Mueller, T. & Wullimann, M. F. Anatomy of neurogenesis in the early zebrafish brain. Brain Res. Dev. Brain Res. 140, 137–155 (2003).

13. Korzh, V., Sleptsova, I., Liao, J., He, J. & Gong, Z. Expression of zebrafish bHLH genes ngn1 and nrd defines distinct stages of neural differentiation. Dev. Dyn. 213, 92–104 (1998).

14. Allende, M. L. & Weinberg, E. S. The expression pattern of two zebrafish achaete-scute homolog (ash) genes is altered in the embryonic brain of the cyclops mutant. Dev. Biol. 166, 509–530 (1994).

15. Wilson, S. W. & Rubenstein, J. L. Induction and dorsoventral patterning of the telencephalon. Neuron 28, 641–651 (2000).

16. Gibbs, H. C., Chang-Gonzalez, A., Hwang, W., Yeh, A. T. & Lekven, A. C. Midbrain-Hindbrain Boundary Morphogenesis: At the Intersection of Wnt and Fgf Signaling. Front. Neuroanat. 11, 64 (2017).

17. Stigloher, C., Chapouton, P., Adolf, B. & Bally-Cuif, L. Identification of neural progenitor pools by E(Spl) factors in the embryonic and adult brain. Brain Res. Bull. 75, 266–273 (2008).

18. Than-Trong, E. & Bally-Cuif, L. Radial glia and neural progenitors in the adult zebrafish central nervous system. Glia 63, 1406–1428 (2015).

19. Clark, B. S. et al. Single-Cell RNA-Seq Analysis of Retinal Development Identifies NFI Factors as Regulating Mitotic Exit and Late-Born Cell Specification. Neuron 102, 1111–1126.e5 (2019).

20. Guo, Q. & Li, J. Y. H. Defining developmental diversification of diencephalon neurons through single cell gene expression profiling. Development 146, dev174284 (2019).

21. Delile, J. et al. Single cell transcriptomics reveals spatial and temporal dynamics of gene expression in the developing mouse spinal cord. Development 146, dev173807 (2019).

22. Holguera, I. & Desplan, C. Neuronal specification in space and time. Science 362, 176–180 (2018).

23. Kim, D. W. et al. Single cell RNA-Seq analysis identifies molecular mechanisms controlling hypothalamic patterning and differentiation. bioRxiv doi: http://dx.doi.org/10.1101/657148 (2019).

24. Tambalo, M., Mitter, R. & Wilkinson, D. G. A single cell transcriptome atlas of the developing zebrafish hindbrain. bioRxiv doi: http://dx.doi.org/10.1101/745141 (2019).

25. Farrell, J. A. et al. Single-cell reconstruction of developmental trajectories during zebrafish embryogenesis. Science 360, eaar3131 (2018).

26. Deniz, S. et al. Mammalian retinal horizontal cells are unconventional GABAergic neurons. J. Neurochem. 116, 350–362 (2011).

27. Cueva, J. G. et al. Vesicular gamma-aminobutyric acid transporter expression in amacrine and horizontal cells. J. Comp. Neurol. 445, 227–237 (2002).

28. Rheaume, B. A. et al. Single cell transcriptome profiling of retinal ganglion cells identifies cellular subtypes. Nat Comms 9, 1–17 (2018).

29. Fischer, A. J., Bongini, R., Bastaki, N. & Sherwood, P. The maturation of photoreceptors in the avian retina is stimulated by thyroid hormone. Neuroscience 178, 250–260 (2011).

30. Brzezinski, J. A., Lamba, D. A. & Reh, T. A. Blimp1 controls photoreceptor versus bipolar cell fate choice during retinal development. Development 137, 619–629 (2010).

31. Viczian, A. S., Vignali, R., Zuber, M. E., Barsacchi, G. & Harris, W. A. XOtx5b and XOtx2 regulate photoreceptor and bipolar fates in the Xenopus retina. Development 130, 1281–1294 (2003).

32. Shen, Y.-C. & Raymond, P. A. Zebrafish cone-rod (crx) homeobox gene promotes retinogenesis. Dev. Biol. 269, 237–251 (2004).

33. Ogawa, Y., Shiraki, T., Kojima, D. & Fukada, Y. Homeobox transcription factor Six7 governs expression of green opsin genes in zebrafish. Proceedings of the Royal Society B: Biological Sciences 282, 20150659 (2015).

34. Satoh, S. et al. The spatial patterning of mouse cone opsin expression is regulated by bone morphogenetic protein signaling through downstream effector COUP-TF nuclear receptors. J. Neurosci. 29, 12401–12411 (2009).

35. Chen, J., Rattner, A. & Nathans, J. The rod photoreceptor-specific nuclear receptor Nr2e3 represses transcription of multiple cone-specific genes. J. Neurosci. 25, 118–129 (2005).

36. McKenzie, J. A. G. et al. Apelin is required for non-neovascular remodeling in the retina. Am. J. Pathol. 180, 399–409 (2012).

37. Centanin, L. & Wittbrodt, J. Retinal neurogenesis. Development 141, 241–244 (2014).

38. Trapnell, C. et al. The dynamics and regulators of cell fate decisions are revealed by pseudotemporal ordering of single cells. Nat Biotechnol 32, 381–386 (2014).

39. Bendall, S. C. et al. Single-cell trajectory detection uncovers progression and regulatory coordination in human B cell development. Cell 157, 714–725 (2014).

40. Mayer, C. et al. Developmental diversification of cortical inhibitory interneurons. Nature 555, 457–462 (2018).

41. Jessell, T. M. Neuronal specification in the spinal cord: inductive signals and transcriptional codes. Nat. Rev. Genet. 1, 20–29 (2000).

42. Lee, S.-K. & Pfaff, S. L. Transcriptional networks regulating neuronal identity in the developing spinal cord. Nat. Neurosci. 4, 1183–1191 (2001).

43. Pandey, S., Shekhar, K., Regev, A. & Schier, A. F. Comprehensive Identification and Spatial Mapping of Habenular Neuronal Types Using Single-Cell RNA-Seq. Curr. Biol. 28, 1052–1065.e7 (2018).

44. Wagner, D. E. et al. Single-cell mapping of gene expression landscapes and lineage in the zebrafish embryo. Science 360, 981–987 (2018).

45. Yoshinari, N., Ando, K., Kudo, A., Kinoshita, M. & Kawakami, A. Colored medaka and zebrafish: Transgenics with ubiquitous and strong transgene expression driven by the medaka β-actinpromoter. Develop. Growth Differ. 54, 818–828 (2012).

46. Pan, Y. A. et al. Zebrabow: multispectral cell labeling for cell tracing and lineage analysis in zebrafish. Development 140, 2835–2846 (2013).

47. Raj, B., Gagnon, J. A. & Schier, A. F. Large-scale reconstruction of cell lineages using single-cell readout of transcriptomes and CRISPR-Cas9 barcodes by scGESTALT. Nat Protoc 13, 2685–2713 (2018).

48. Butler, A., Hoffman, P., Smibert, P., Papalexi, E. & Satija, R. Integrating single-cell transcriptomic data across different conditions, technologies, and species. Nat Biotechnol 36, 411–420 (2018).

49. Felsenstein, J. PHYLIP - Phylogeny Inference Package (Version 3.2). Cladistics, Vol. 5 (1989), pp. 164-166 5, 164–166 (1989).

50. Siebert, S. et al. Stem cell differentiation trajectories in Hydra resolved at single-cell resolution. Science 365, eaav9314 (2019).

51. Haghverdi, L., Buettner, F. & Theis, F. J. Diffusion maps for high-dimensional single-cell analysis of differentiation data. Bioinformatics 31, 2989–2998 (2015).

52. Haghverdi, L., Büttner, M., Wolf, F. A., Buettner, F. & Theis, F. J. Diffusion pseudotime robustly reconstructs lineage branching. Nat. Methods 13, 845–848 (2016).

