## Supplemental Figures for "Emergence of neuronal diversity during vertebrate brain development"

### Sup Figure 1

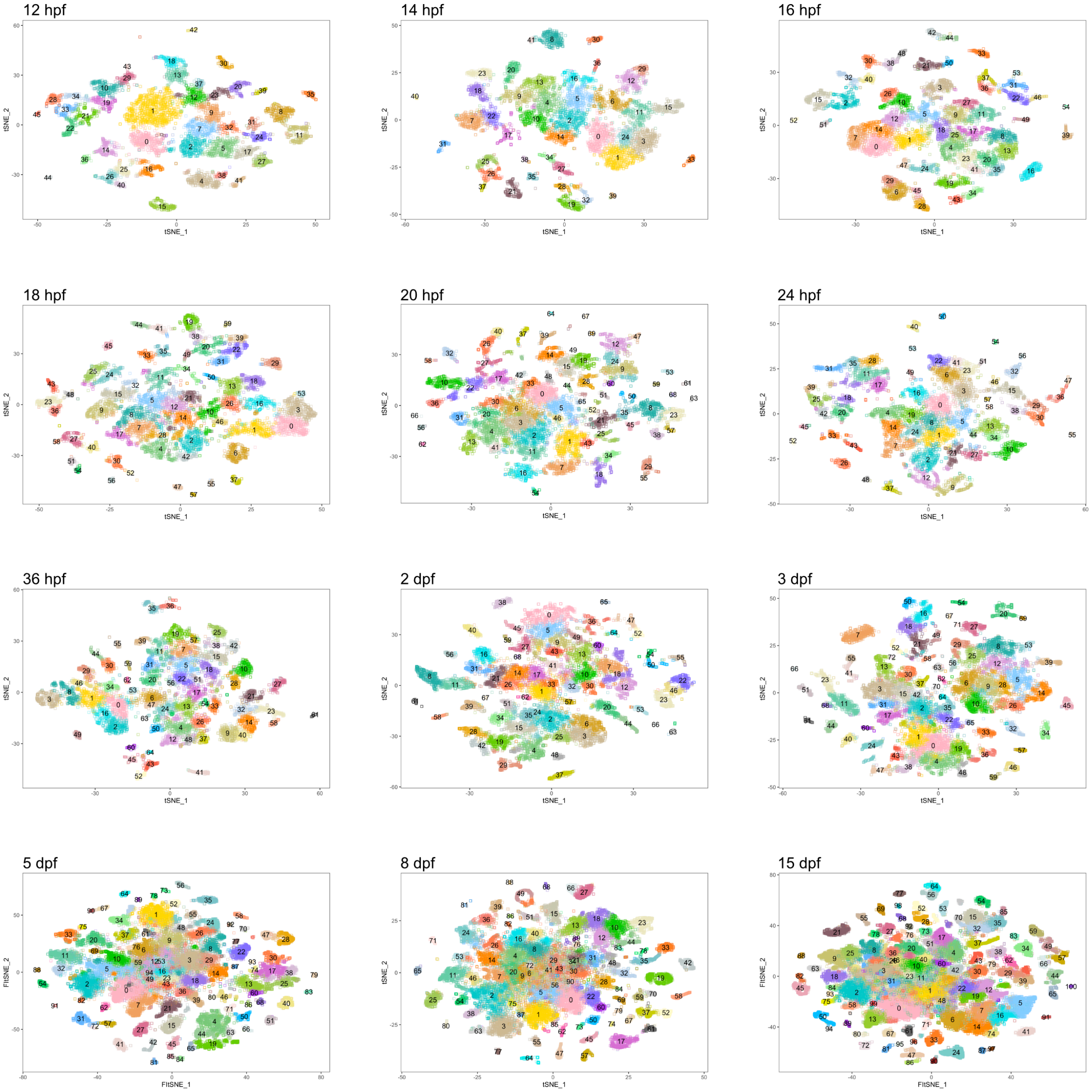


**Sup Figure 1. Zebrafish brain cell types identified at each stage of time course**

tSNE plots of cell types at each stage of the time course. Cells are color coded by stage.

tSNE implementations: Barnes-Hut (12 hpf to 3 dpf, 8 dpf), Fourier transform (5 dpf and 15 dpf)

### Sup Figure 2

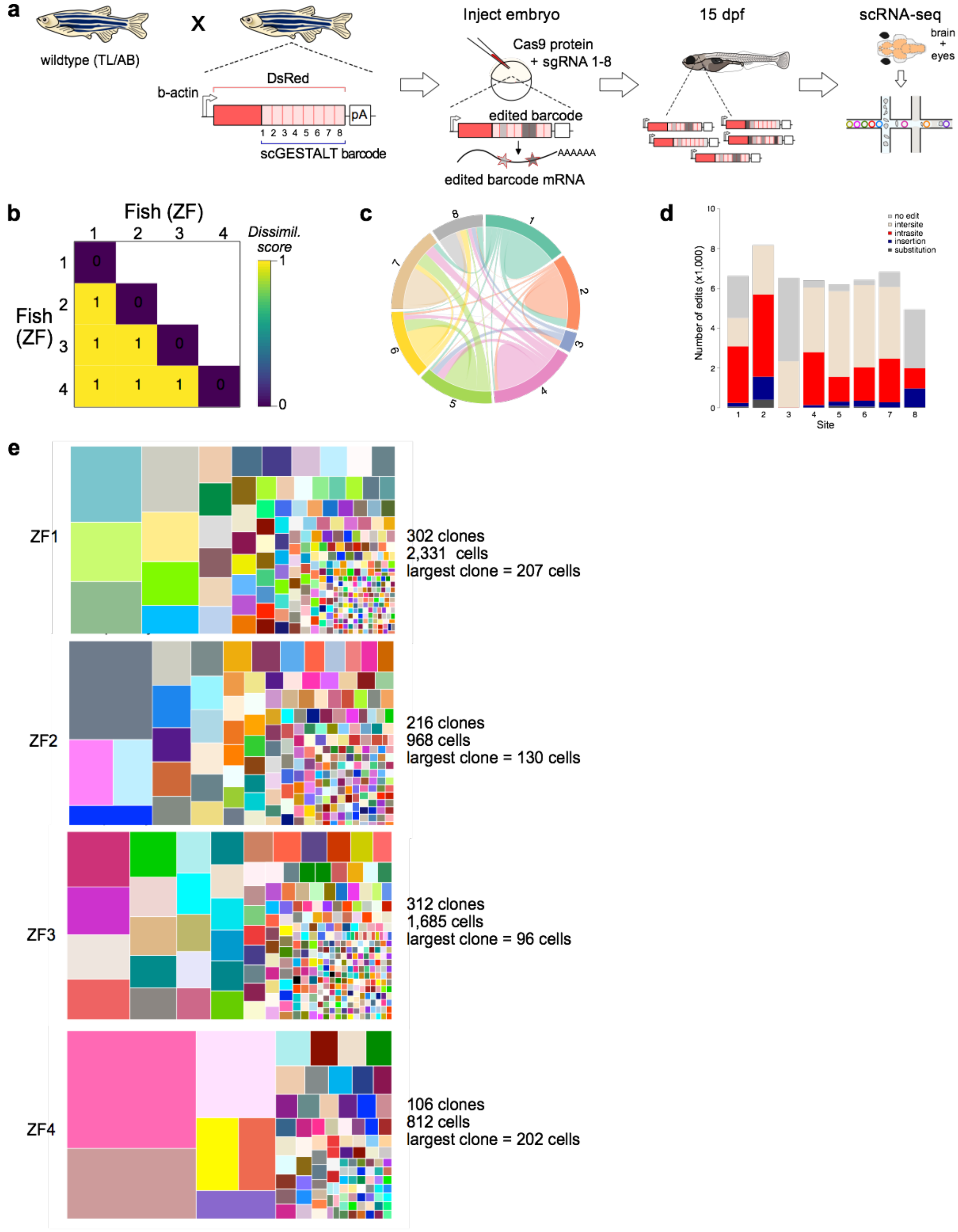


**Sup Figure 2. Optimization of scGESTALT lineage recorder for better barcode recovery**

**a.** Schematic overview of CRISPR-Cas9 lineage recording. Optimized scGESTALT comprises a barcode cassette in the 3'end of DsRed transgene (single copy) and the medaka beta-actin promoter. Embryos are injected with Cas9 protein and DsRed sgRNAs and animals are profiled at 15 dpf by scRNA-seq.

**b.** Pairwise comparisons using cosine dissimilarity of barcode edit patterns from four (ZF1-4) edited 15 dpf larval brains.
**c.** Chord diagram of the nature and frequency of deletions within and between target sites. Each colored sector represents a target site. Links between target sites represent inter-site deletions; self-links represent intra-site deletions. Link widths are proportional to the edit frequencies.
**d.** Type of edit at each target site within the barcode from edited ZF1-4 larval brains.
**e.** Size and diversity of clones from edited ZF1-4 larval brains. Each colored rectangle represents a unique clone and the area represent the size of the clone.

**Sup Fig 3**

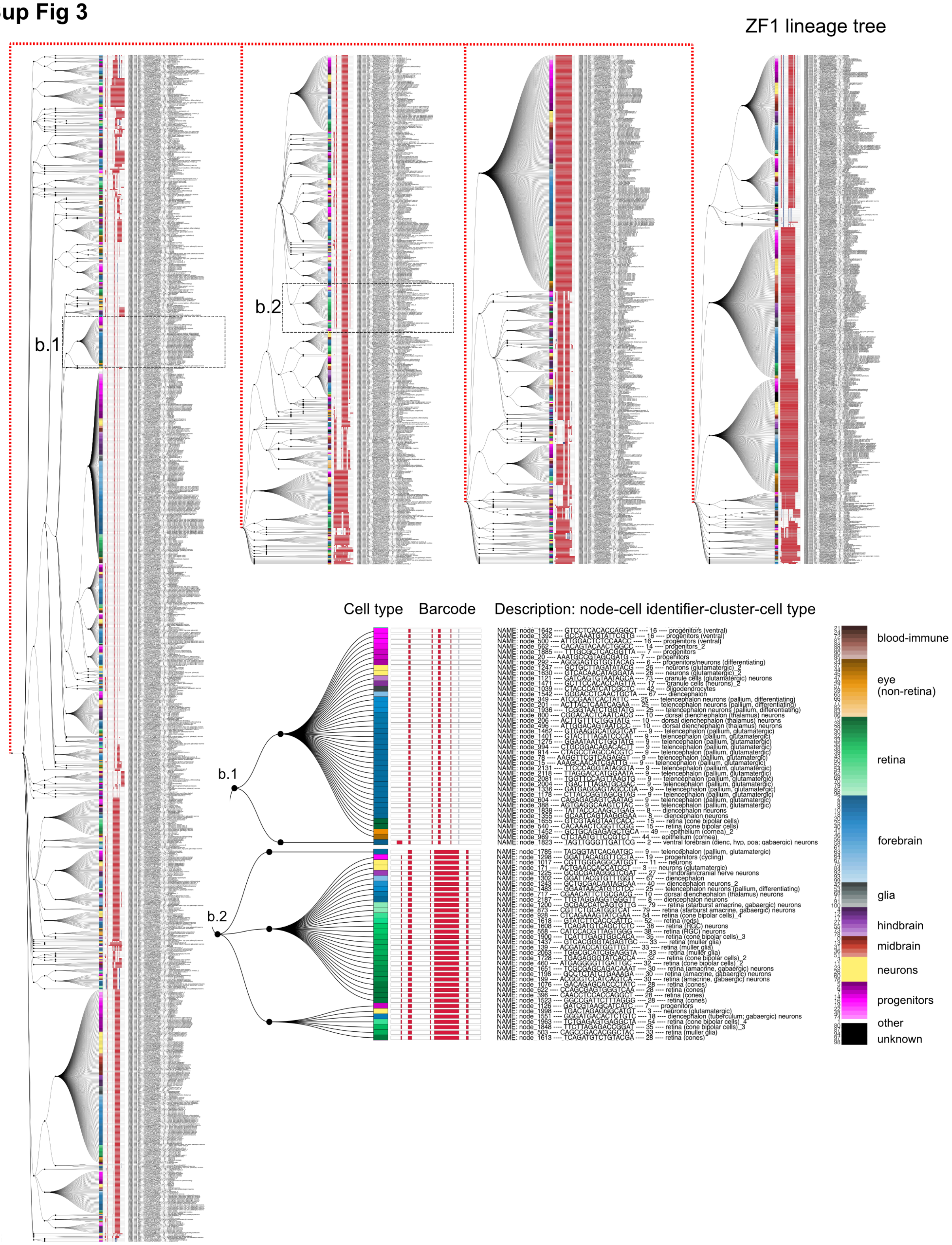


### Sup Figure 3. Reconstructed lineage tree of a 15 dpf zebrafish brain (ZF1)

A reconstructed scGESTALT brain lineage tree from one zebrafish 15 dpf brain. 302 barcodes recovered from ZF1 were assembled into a lineage tree. Barcode edits are represented as red (deletions), blue (insertions), and black (substitutions). Associated cells are color coded by cell type and region. Interactive trees are presented at: https://scgestalt.mckennalab.org/. Zoomed in views of clades b.1 and b.2 (dashed boxes) are shown. Each tip on the tree has an associated cell type assignment (color coded), a lineage barcode schematic, and a description which comprises a node number, a cell identifier (matches the transcriptome cell identifier), cluster number, and an expanded cell type information. For reasons of space, the tree is split into multiple columns and dashed lines connect subsections of the tree together.

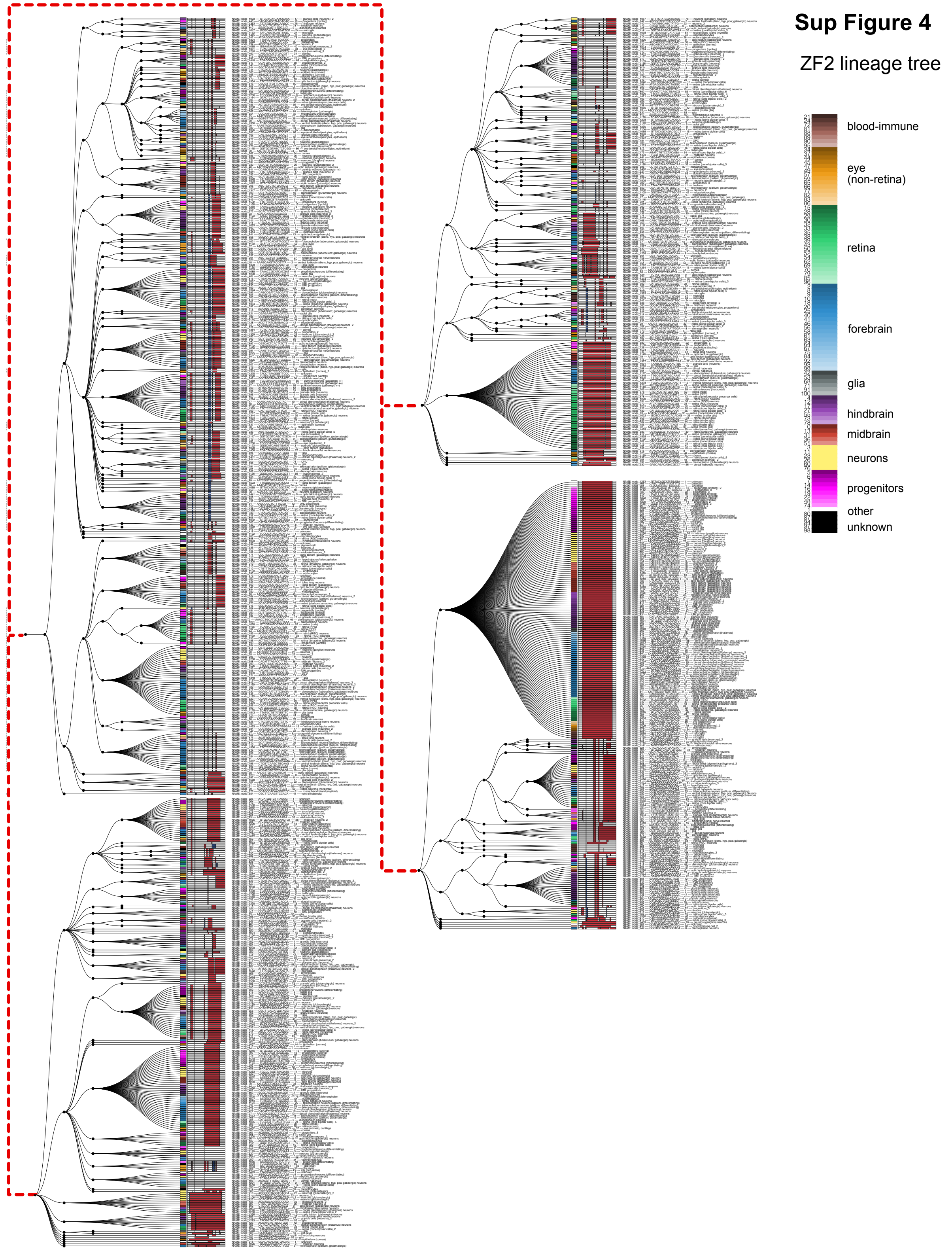


**Sup Figure 4. Reconstructed lineage tree of a 15 dpf zebrafish brain (ZF2)**
A reconstructed scGESTALT brain lineage tree from one zebrafish 15 dpf brain. 216 barcodes recovered from ZF2 were assembled into a lineage tree. Barcode edits are represented as red (deletions), blue (insertions),

and black (substitutions). Associated cells are color coded by cell type and region. Interactive trees are presented at: https://scgestalt.mckennalab.org/. For reasons of space, the tree is split into multiple columns and dashed lines connect subsections of the tree together.

### Sup Figure 5

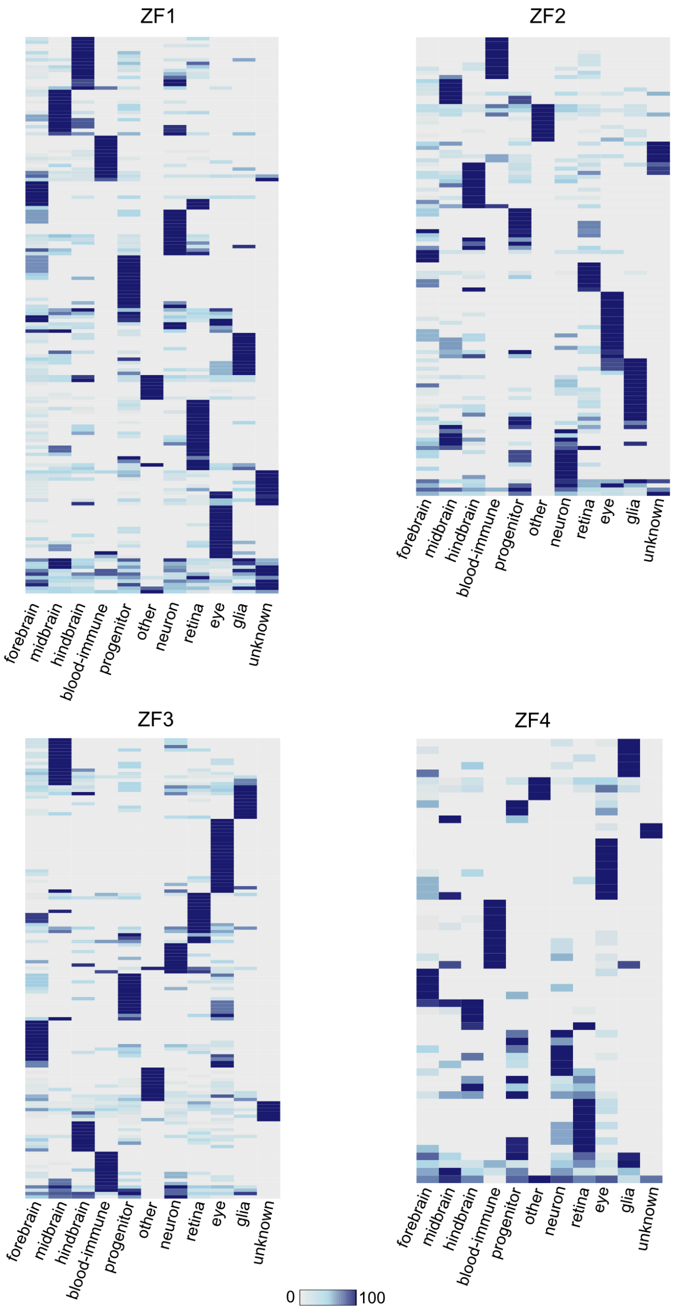


**Sup Figure 5. Barcodes are enriched within regions of the brain**
Heat map of the distribution of ZF1-ZF4 barcodes (rows) for each region of the brain (columns 1-3). Additional cell types that are not restricted to a given region or could not be classified to a given region (e.g. progenitors, nascent neurons, unknown, etc.) are also indicated. Cell types were classified as belonging to the forebrain, midbrain, hindbrain, or other areas as indicated, and the proportions of cells within each region were calculated for each barcode. Region proportions were scaled by row and colored as shown in the legend.

### Sup Figure 6

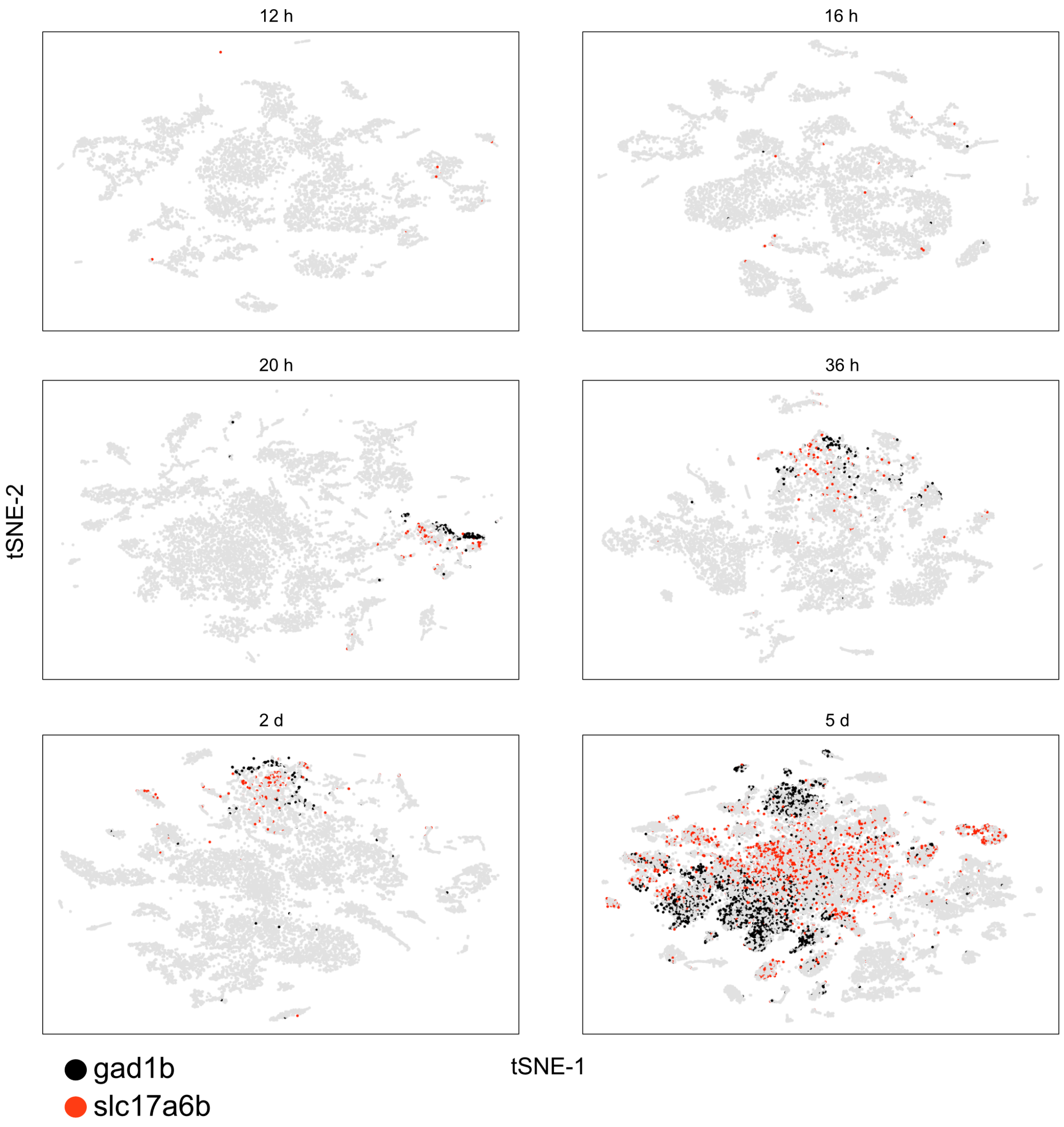


**Sup Figure 6. Detection of GABAergic and glutamatergic neuronal populations during development**

tSNE plots highlight cells expressing gad1b (GABAergic marker, black) or slc17a6b (glutamatergic marker, red) across various developmental stages.

tSNE implementations: Barnes-Hut (12h to 2d), Fourier transform (5d)

### Sup Figure 7

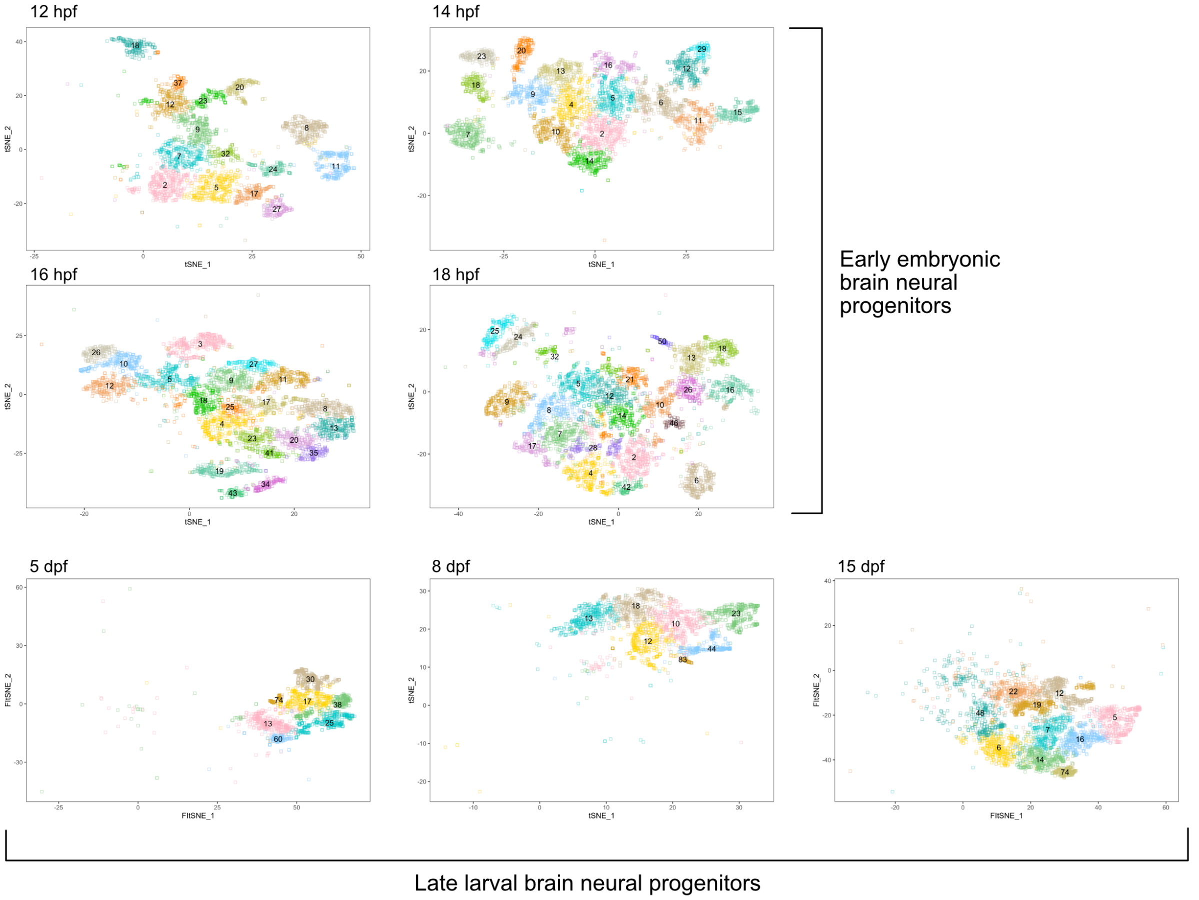


**Sup Figure 7. Embryonic and larval neural progenitor populations**

tSNE plots showing embryonic and larval progenitor clusters that were subsetted from datasets corresponding to indicated time points. Cluster numbers match plots shown in Sup Fig. 1

### Sup Figure 8

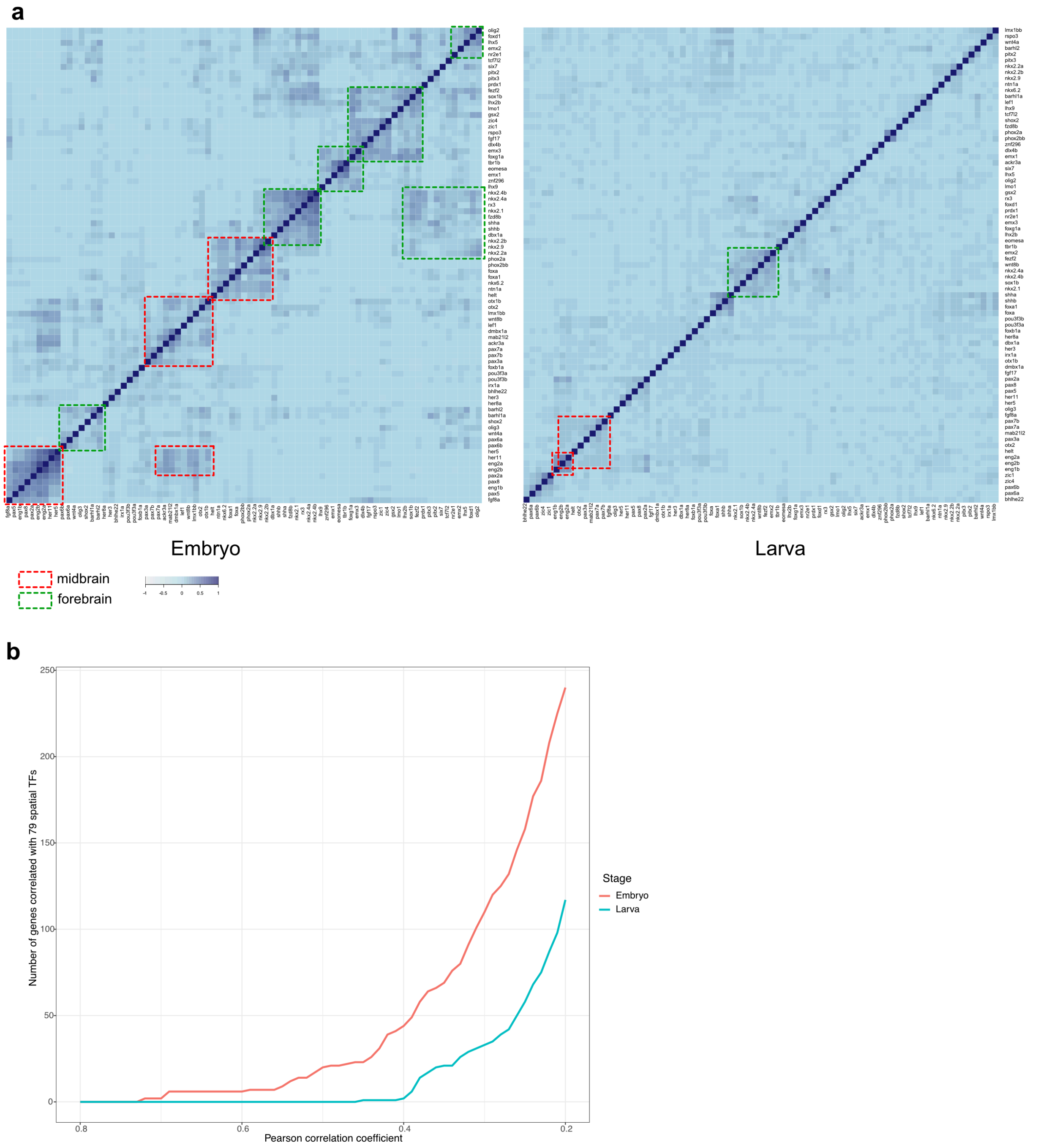


**Sup Figure 8. Dampening of spatial signatures in neural progenitors during development**

**a.** Heatmap of Pearson correlation values of 79 spatial markers in embryonic and larval neural progenitors. Spatial markers were selected based on existing literature. Groups of co-varying genes in the midbrain and forebrain are highlighted with dashed boxes.

**b.** Plot showing number of highly variable genes that co-vary with any of the selected 79 spatial markers in embryonic and larval progenitors. Co-variation was determined by Pearson correlation, with several thresholds (from stringent to relaxed) displayed along the x-axis.

**Sup Figure 9**

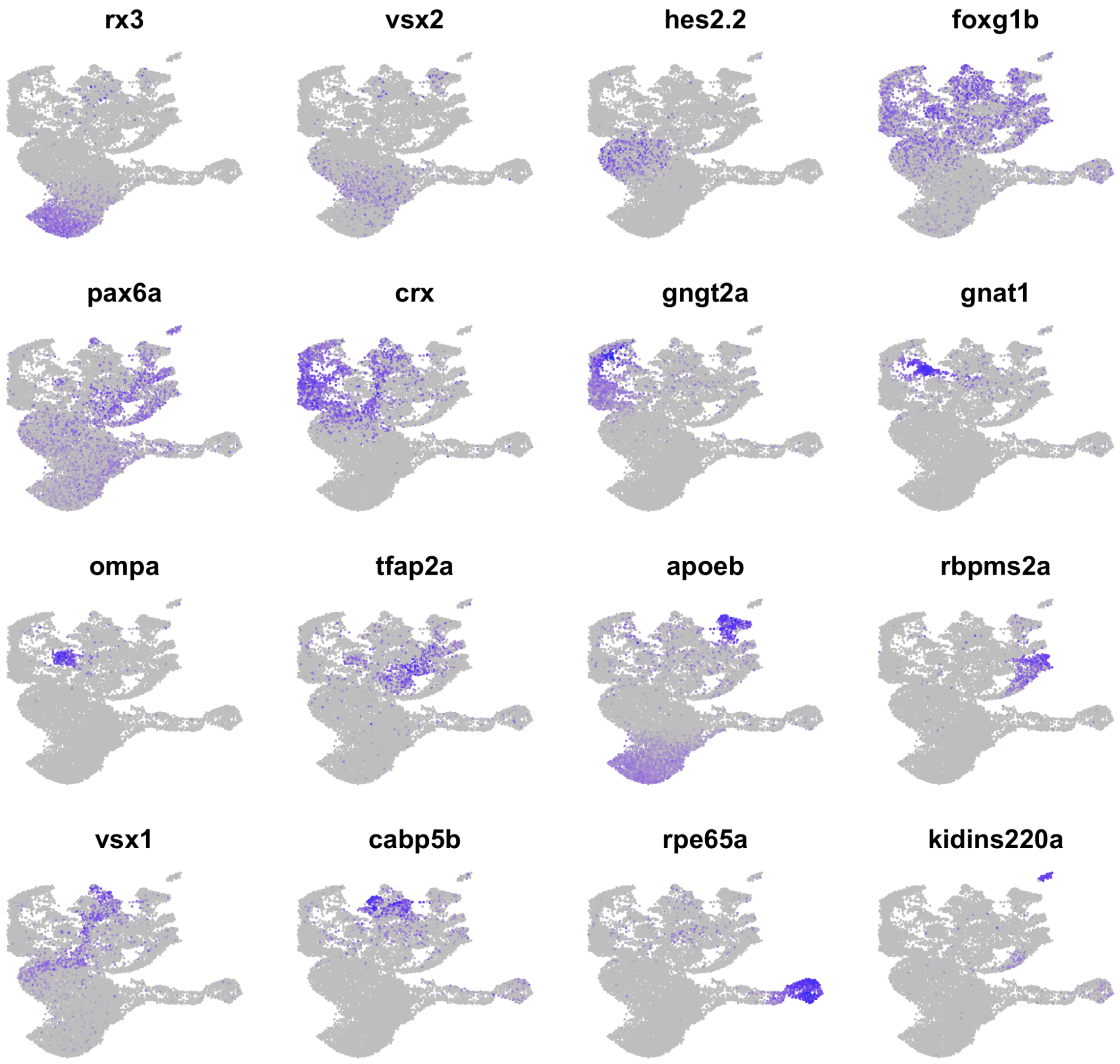


**Sup Figure 9. Expression of select retinal cell type markers**

UMAP plots highlighting expression of select genes enriched in retinal cell types. *rx3*, *vsx2*, *hes2.2* enriched in early embryonic retinal progenitors; *foxg1b* enriched in differentiated cells; *pax6a* enriched in progenitors, retinal ganglion cells (RGC) and amacrine cells; *crx* enriched in photoreceptor cells and cone bipolar cells; *gngt2a* enriched in cones; *gnat1* enriched in rods; *ompa* enriched in horizontal cells; *tfap2a* enriched in RGCs and horizontal cells; *apoeb* enriched in early progenitors and muller glia; *rbpms2a* enriched in amacrine cells; *vsx1* enriched in cone bipolar cells; *cabp5b* enriched in cone bipolar cells; *rpe65a* enriched in retinal pigment epithelium; *kidins220a* enriched in new retinal subtype.

**Sup Figure 10**

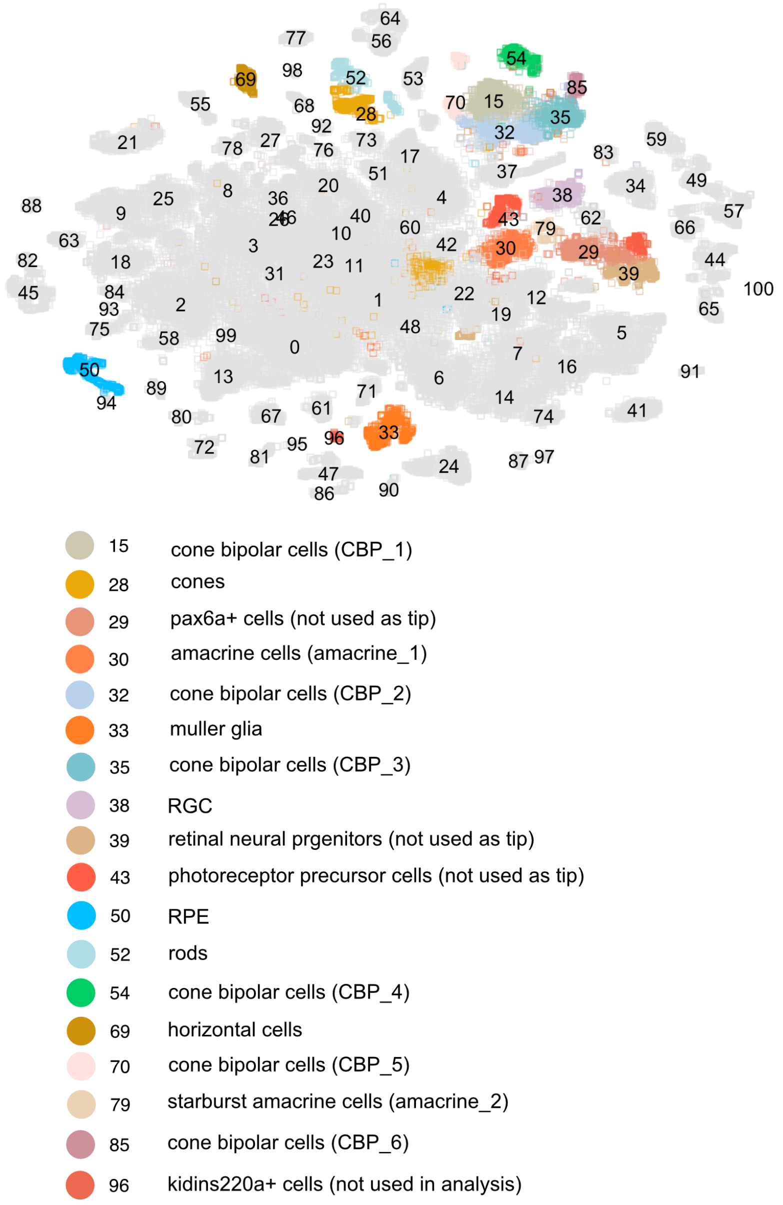


**Sup Figure 10. Retinal cell types selected as tips of specification tree**

tSNE plot of 15 dpf brain and eye cell types. Retinal cell types used in URD analysis are color coded (note cluster 96 was discarded from all analysis).

### Sup Figure 11

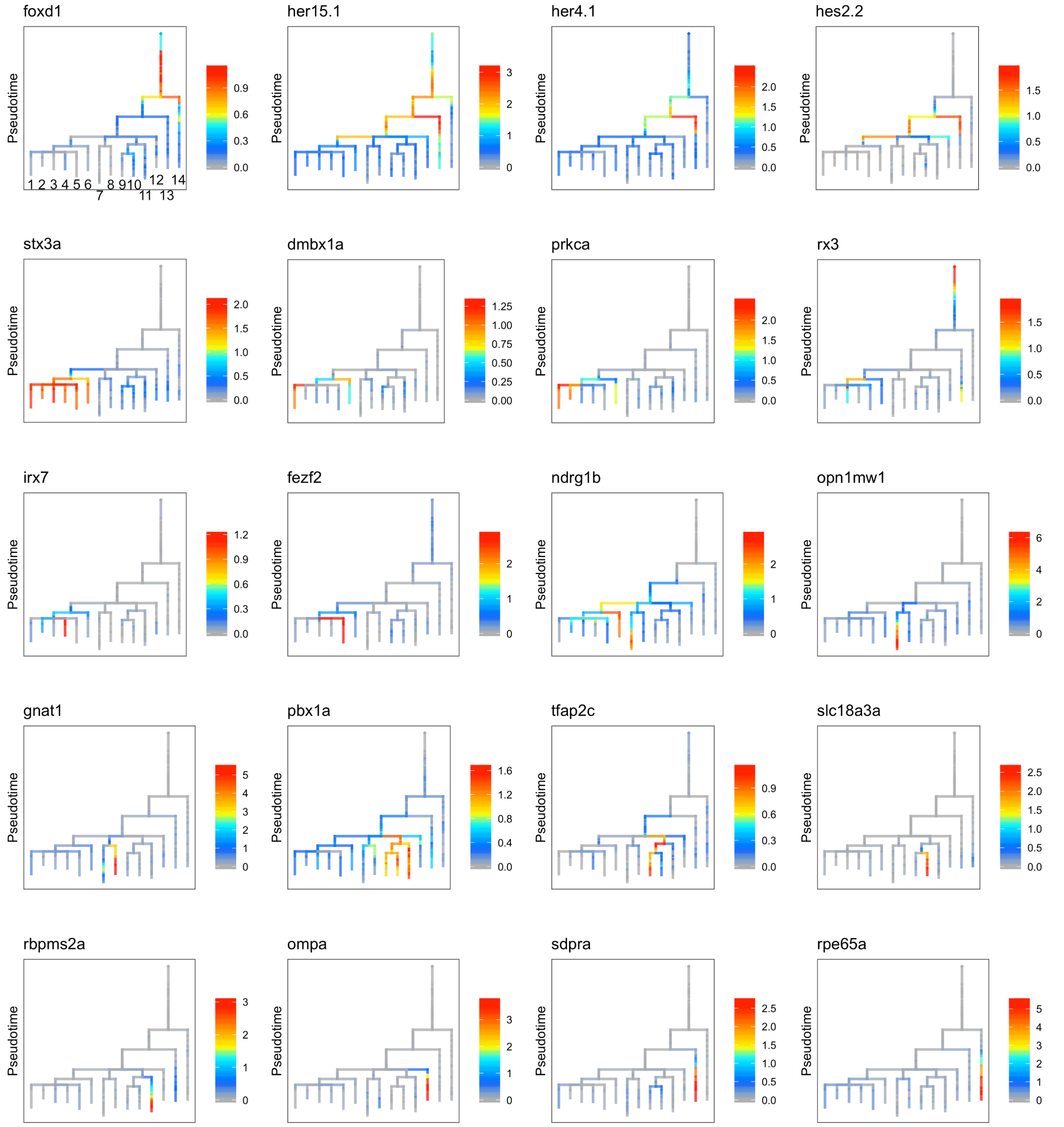


**Sup Figure 11. Gene expression along retina cell trajectories**

Expression of select genes are shown on the retina specification tree. Cell types: 1. Cone bipolar cell (CBP_3); 2. Cone bipolar cell (CBP_6); 3. Cone bipolar cell (CBP_1); 4. Cone bipolar cell (CBP_4); 5. Cone bipolar cell (CBP_5); 6. Cone bipolar cell (CBP_2); 7. Cones; 8. Rods; 9. Amacrine cells (Amacrine_1); 10. Amacrine cells (Amacrine_2); 11. Retinal ganglion cells (RGC); 12. Horizontal cells; 13. Muller glia; 14. Retinal pigment epithelium (RPE)

### Sup Figure 12

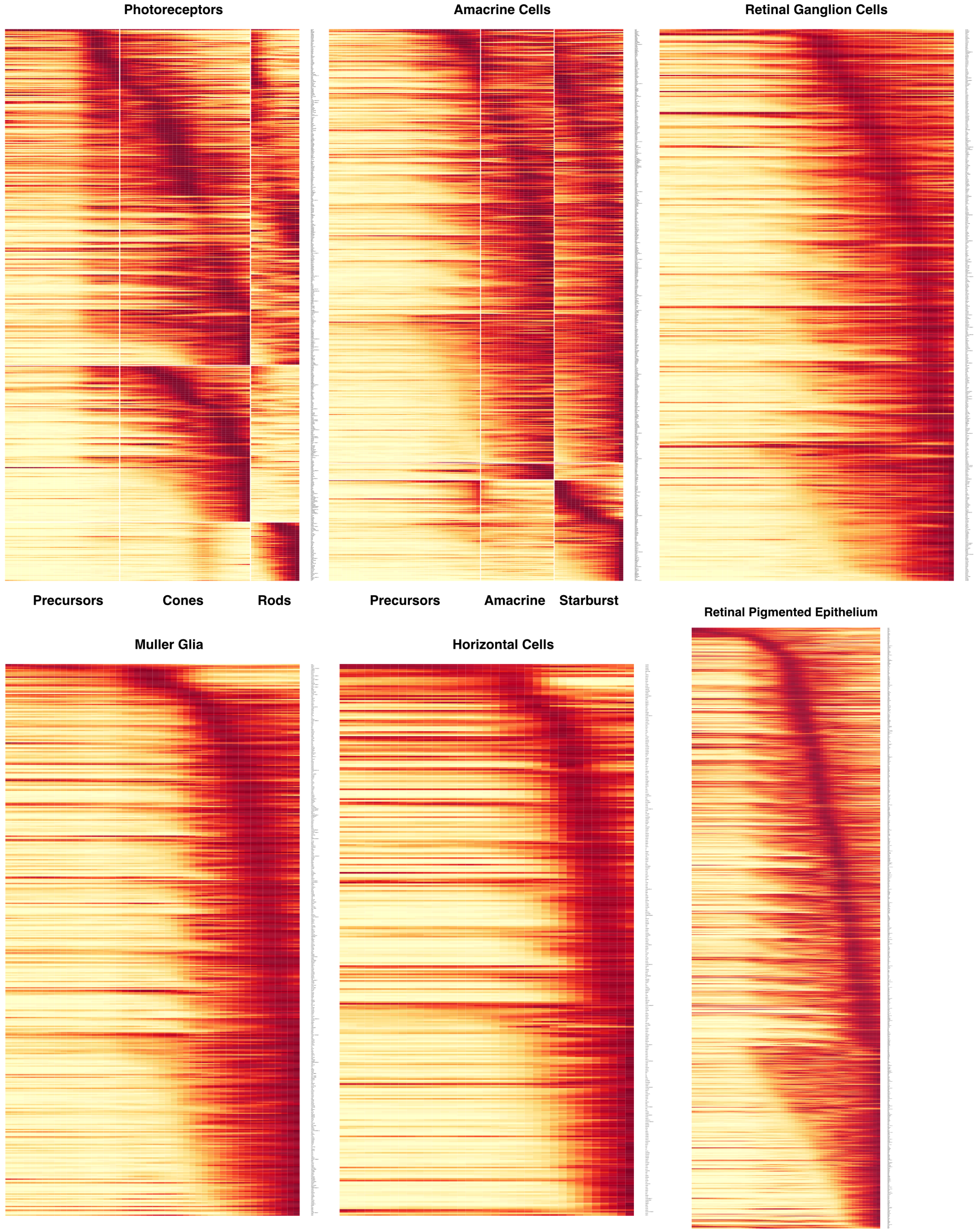


**Sup Figure 12. Gene expression cascades of retinal cell trajectories**

Heat maps of gene expression cascades of photoreceptor cell, amacrine cell, retinal ganglion cell, muller glia, horizontal cell and retinal pigment epithelium cell trajectories. Cells were selected based on high expression along trajectories leading to these cell types, compared to expression along opposing branchpoints. Red, high expression. Yellow, low expression.

### Sup Figure 13

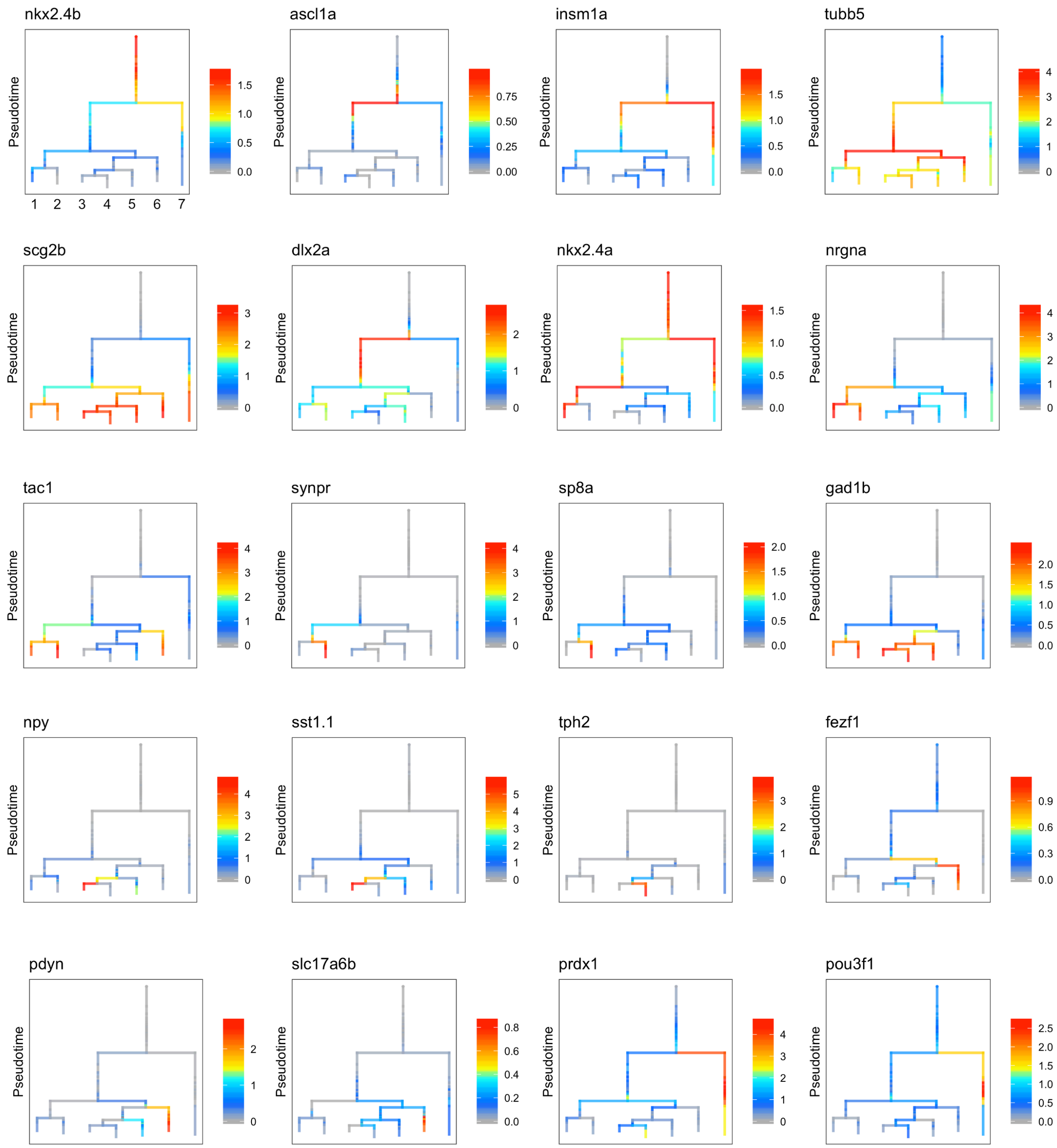


**Sup Figure 13. Gene expression along hypothalamus cell trajectories**

Expression of select genes are shown on the hypothalamus specification tree. Cell types: 1. GABA tac1+, nrgna+; 2. synpr+; nrgna+; 3. sst1.1+; 4. tph2+; 5. GABA dlx+; 6. pdyn+; 7. prdx1+

### Sup Figure 14

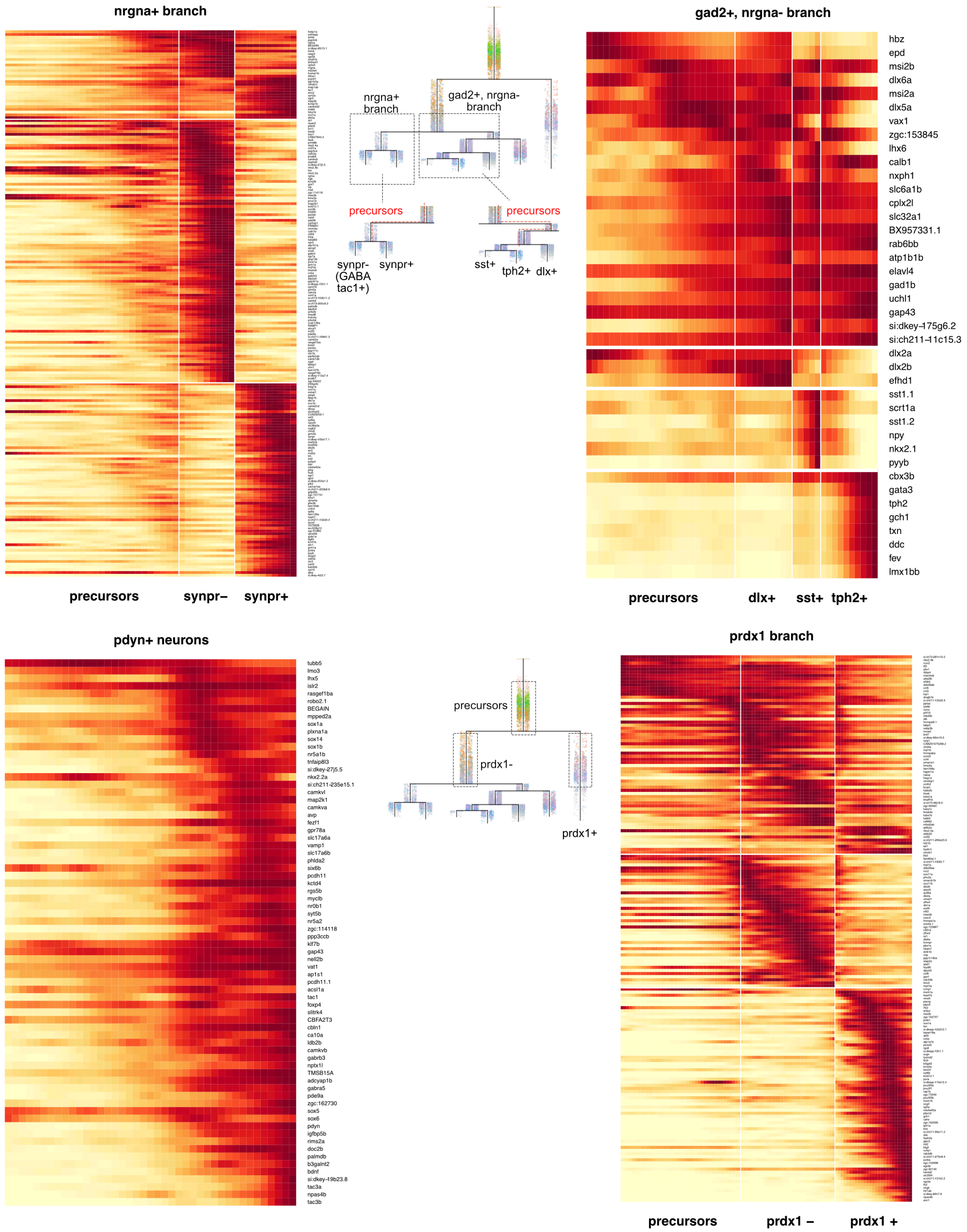


### Sup Figure 14. Gene expression cascades of hypothalamus cell trajectories

Heat maps of gene expression cascades of profiled hypothalamus cell trajectories. Cells were selected based on high expression along trajectories leading to these cell types, compared to expression along opposing branchpoints. Red, high expression. Yellow, low expression.

**Sup Figure 15**

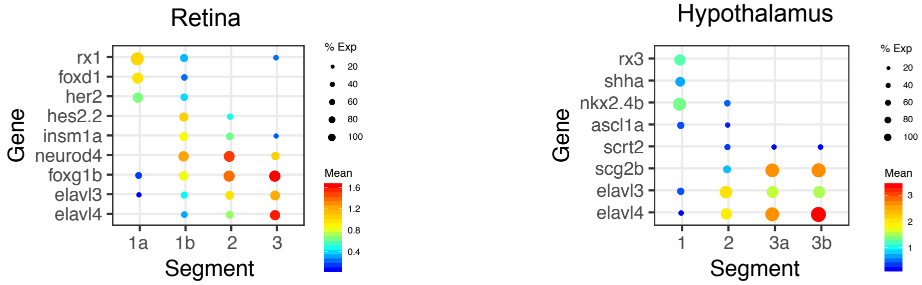


**Sup Figure 15. Gene markers of embryonic and larval progenitors in retina and hypothalamus**

Dot plot of gene expression pattern of select marker genes that were used to define progenitor and precursor states (rows) for segments (columns) of the retina (left) or hypothalamus (right) cell specification trees (see Fig. 5e). Dot size indicates the percentage of cells expressing the marker; color represents the average scaled expression level.
