## Supplemental Analysis for "Emergence of neuronal diversity during vertebrate brain development"

### Retina: 1 - URD object & doublet removal

*Jeff Farrell*

*8/22/2019*

#### Contents

Import data into URD 1
Convert Seurat object to URD . . . . . . . . . . . . . . . . . . . . . . . . . . . . . . 1
Combined individual stage clustering . . . . . . . . . . . . . . . . . . . . . . . . . . 1

Calculate highly variable genes 2

Calculate KNN graph and remove outliers 7

Remove ***kidins220a+*** population 8

Remove cell type doublets 8

### Import data into URD

#### Convert Seurat object to URD

We first loaded a Seurat object that contained just cells from the clusters that belonged to the hypothalamus from each stage.

```
suppressPackageStartupMessages(library(URD))
suppressPackageStartupMessages(library(Seurat))
```

```
base.path <- "~/urd-cluster-bushra/"

# Load Seurat object that has been cropped to hypothalamus cells
object.seurat <- readRDS(paste0(base.path, "obj/retina.new_seurat.rds"))

# Convert to URD object
suburd <- seuratToURD(object.seurat)
```

#### Combined individual stage clustering

Bushra had performed individual clusterings across each stage with different resolutions. Here, it was better to create a single identifier that included stage + cluster information to combine all those clusterings (while preventing any overlap).

```r
stages <- sort(unique(suburd@meta$stage))
clust.res.used <- paste0("res.", c("4.5", "4", "5", "5", "4.5", "5", "6",
    "6", "6", "5.5", "6", "5"))
names(clust.res.used) <- stages
suburd@group.ids$cluster <- NA
for (stage in stages) {
    suburd@group.ids[cellsInCluster(suburd, "stage", stage), "cluster"] <- paste0(stage,
        "-", suburd@group.ids[cellsInCluster(suburd, "stage", stage), clust.res.used[stage]])
}
```

#### Calculate highly variable genes

We calculated highly variable genes for each stage, used genes that were found as highly in at least two stages, but were not mitochondrial, ribosomal, heat-shock protein, or tandem duplicated genes.

```r
# Calculated on each stage separaely, final gene list was all genes
# that were 'variable' in at least two stages NB: For a couple of
# stages, the gamma fit was poor -- the library size distribution
# seemed bimodal. Have seen this before in 10X data, but not sure what
# it means.
var.genes.by.stage <- lapply(stages, function(stage) {
    findVariableGenes(suburd, cells.fit = cellsInCluster(suburd, "stage",
        stage), set.object.var.genes = F, diffCV.cutoff = 0.3, main.use = stage,
        do.plot = T)
})
```

**01-12h**

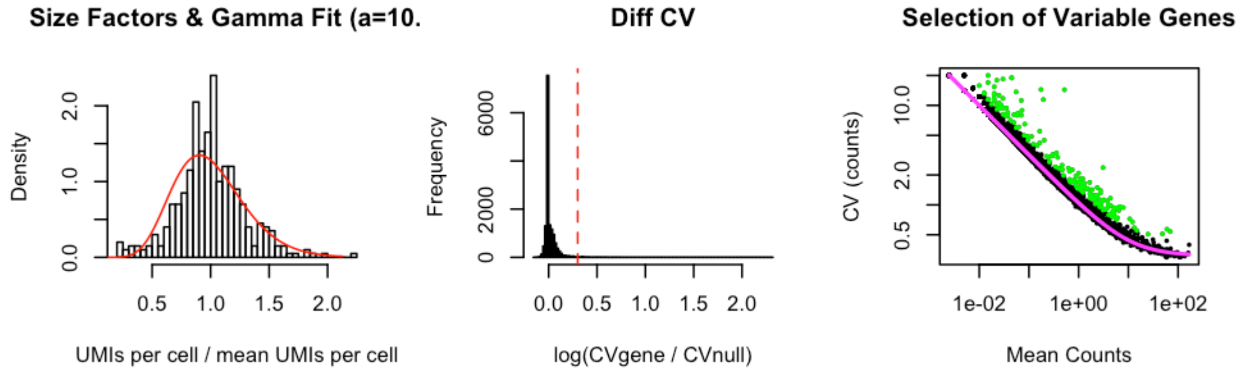

**02-14h**

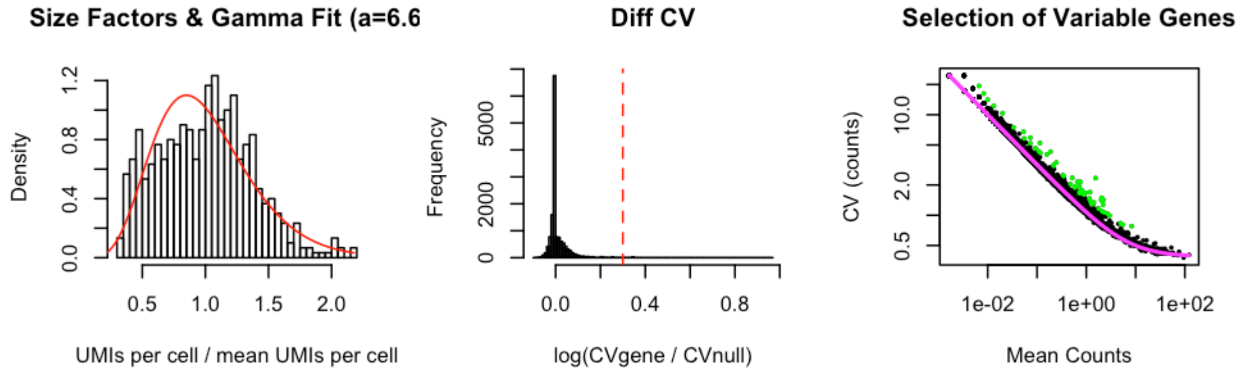

**03-16h**

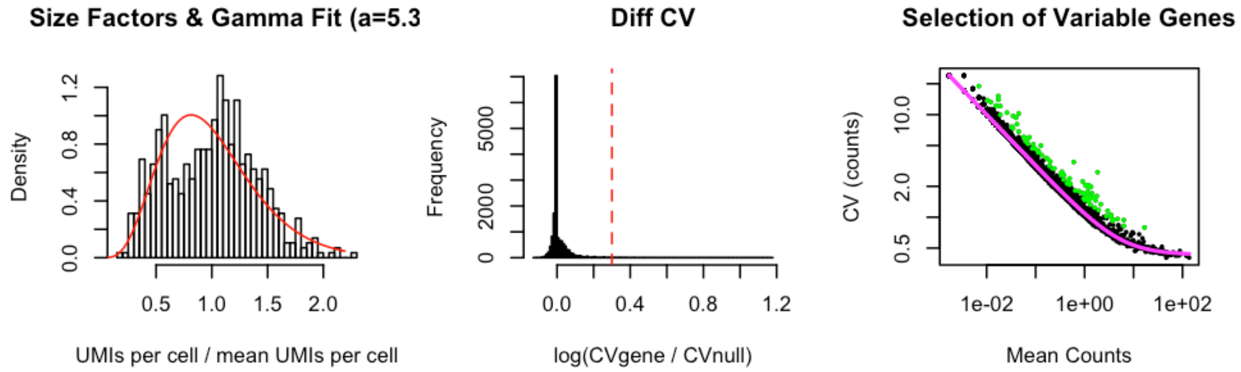

**04-18h**

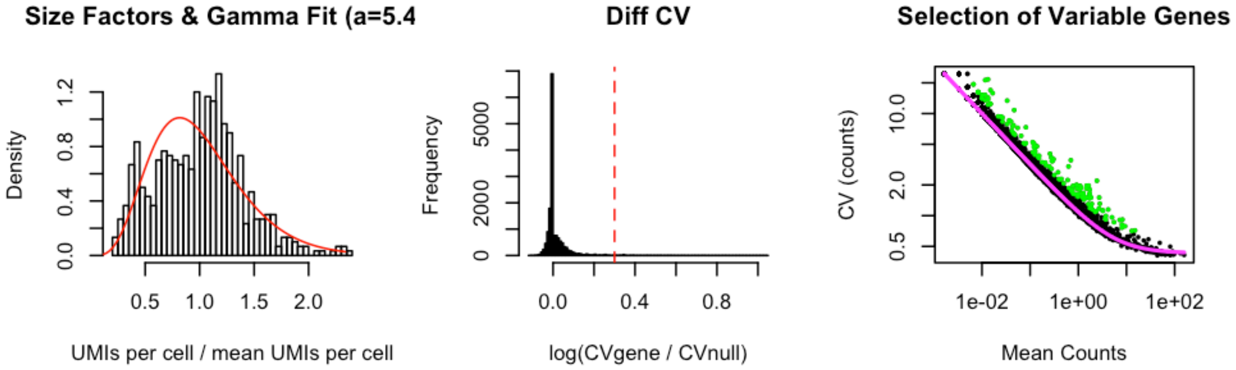

**05-20h**

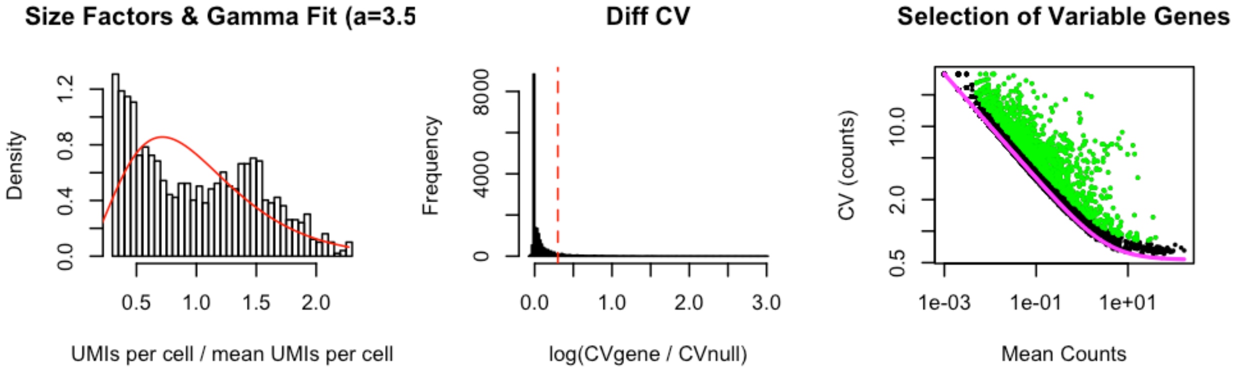

**06-24h**

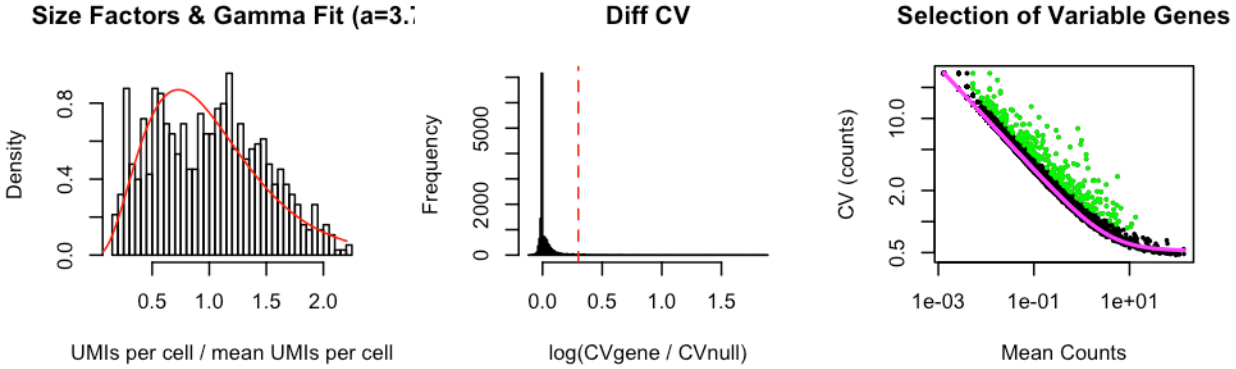

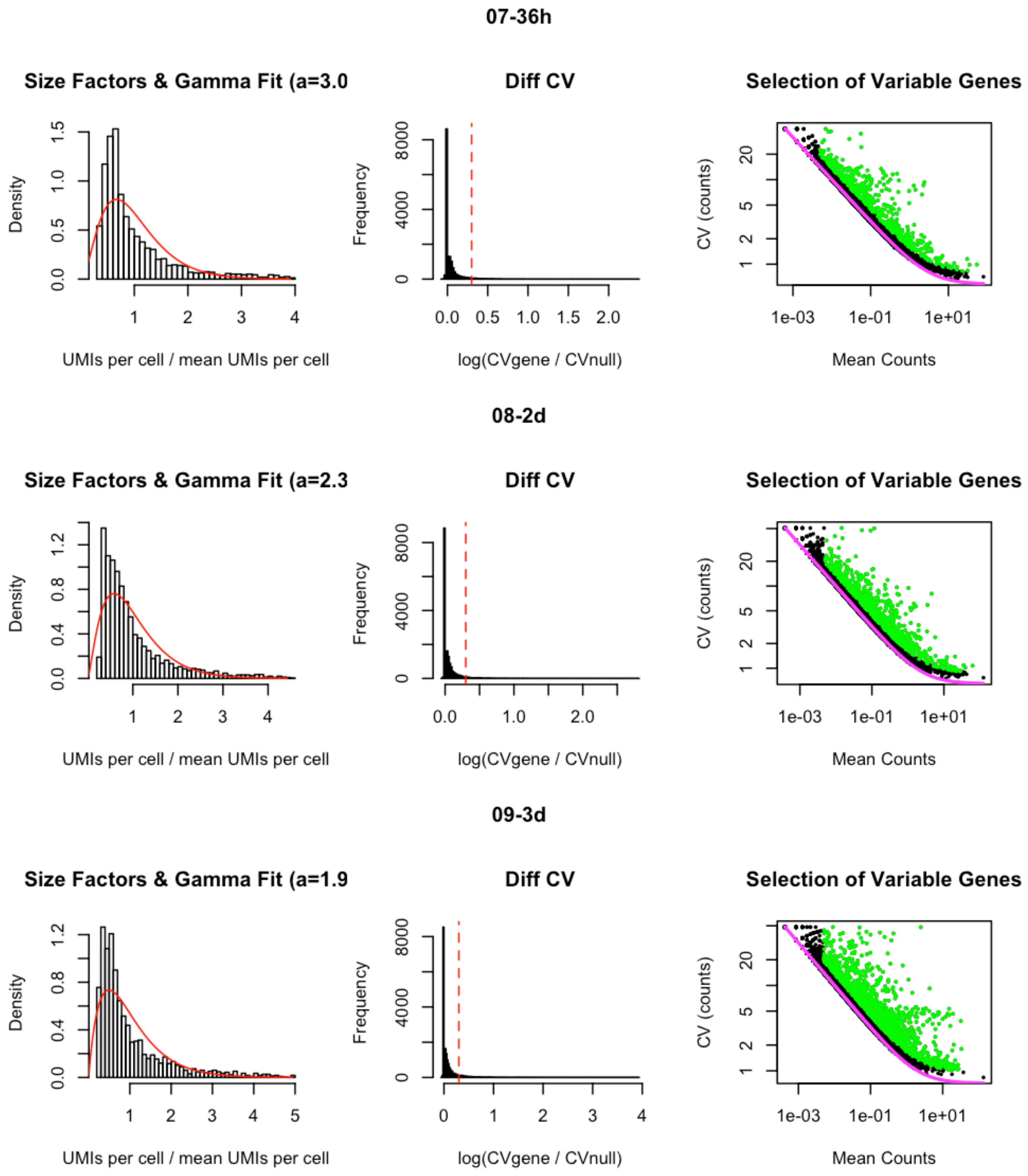
07-36h
Size Factors & Gamma Fit (a=3.0
Density
UMIs per cell / mean UMIs per cell
Diff CV
Frequency
log(CVgene / CVnull)
Selection of Variable Genes
CV (counts)
Mean Counts
08-2d
Size Factors & Gamma Fit (a=2.3
Density
UMIs per cell / mean UMIs per cell
Diff CV
Frequency
log(CVgene / CVnull)
Selection of Variable Genes
CV (counts)
Mean Counts
09-3d
Size Factors & Gamma Fit (a=1.9
Density
UMIs per cell / mean UMIs per cell
Diff CV
Frequency
log(CVgene / CVnull)
Selection of Variable Genes
CV (counts)
Mean Counts

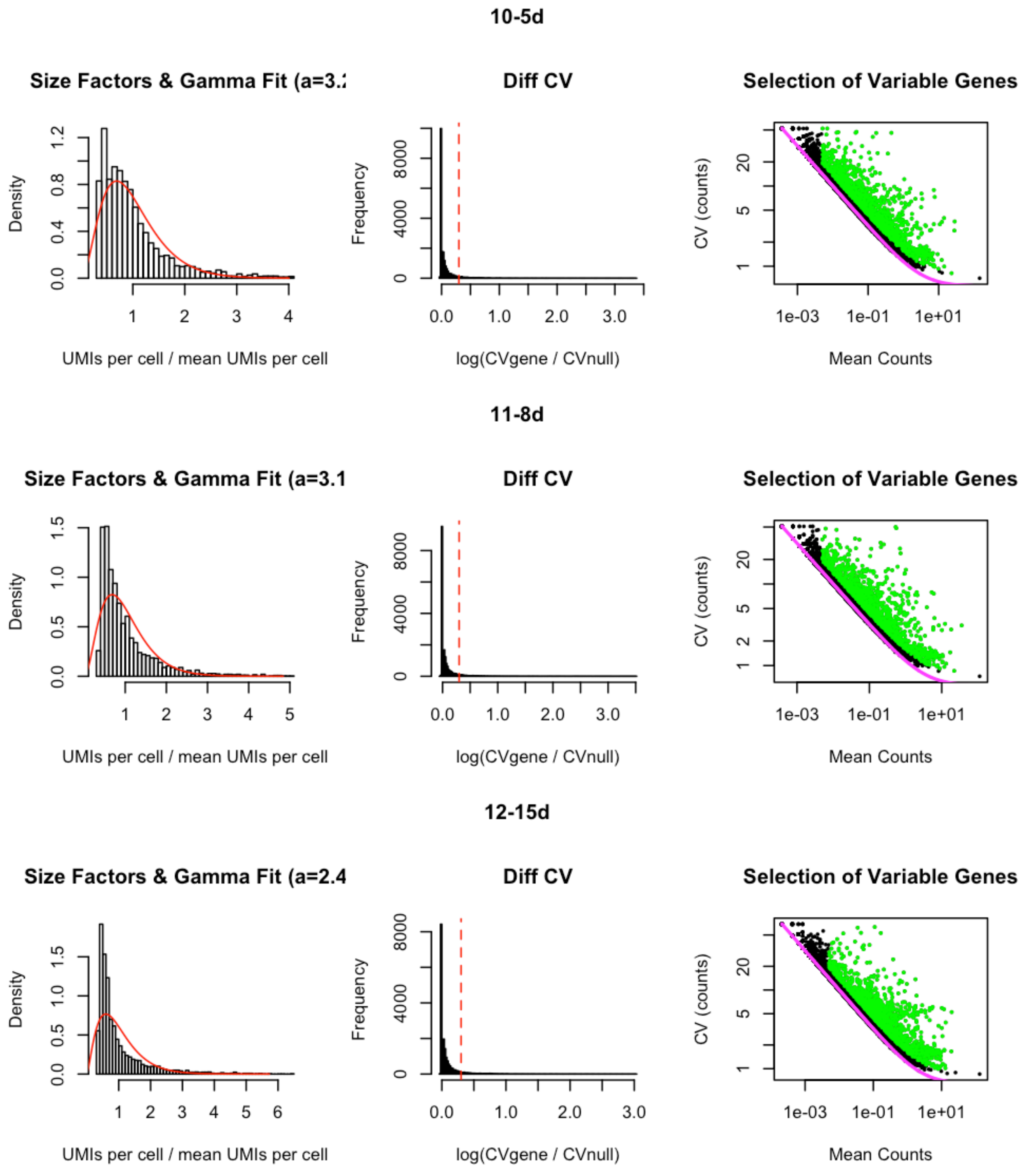

```r
names(var.genes.by.stage) <- stages
var.genes <- sort(unique(unlist(var.genes.by.stage)))
print(paste0("Length of variable genes is ", length(var.genes)))
```

```
## [1] "Length of variable genes is 2636"
```

```r
var.genes.twice <- names(which(table(unlist(var.genes.by.stage)) >= 2))
print(paste0("Length of variable genes shared across at least 2 stages is ",
    length(var.genes.twice)))
```

```
## [1] "Length of variable genes shared across at least 2 stages is 1724"
```

```
# Remove mitochondrial genes
var.mito <- grep("^mt-|^AC0", var.genes.twice, value = T)
# Remove ribosomal genes
var.ribo <- grep("^rps|^rpl", var.genes.twice, value = T)
# Remove hsp genes
var.hsp <- grep("^hsp", var.genes.twice, value = T)
# Remove genes with duplicates
var.dups <- grep("of many", var.genes.twice, value = T)
suburd@var.genes <- setdiff(var.genes.twice, c(var.mito, var.ribo, var.hsp,
    var.dups))
print(paste0("Length of final variable genes list (after removing mito, ribo, hsp genes) is ",
    length(suburd@var.genes)))
```

```
## [1] "Length of final variable genes list (after removing mito, ribo, hsp genes) is 1595"
```

To prevent downstream problems, we also removed any cells from the data that had the exact same expression of the variable genes (i.e. cells with completely duplicated coordinates in the high-dimensional space we would use for analysis downstream).

```
# Check for duplicate data points - cells with exact same expression of
# variable genes
vg.dups <- duplicated(as.data.frame(as.matrix(t(suburd@logupx.data[suburd@var.genes,
    ]))))
if (length(which(vg.dups)) > 0) {
    print(paste("Removing", length(which(vg.dups)), "cell(s) with duplicated variable gene expression."))
    not.dup.cells <- colnames(suburd@logupx.data)[!vg.dups]
    suburd <- urdSubset(suburd, not.dup.cells)
}
```

```
## [1] "Removing 6 cell(s) with duplicated variable gene expression."
```

### Calculate KNN graph and remove outliers

We then calculated a k-nearest neighbor graph and removed cells that had unusual distance to their nearest neighbor, or unusual distance to their 20th nearest neighbor (given their distance to their nearest neighbor). These sorts of outliers often cause problems or skew diffusion maps (used downstream).

```
# Calculate k-nn
suburd <- calcKNN(suburd)
```

```
# Check what the outliers are
outliers <- knnOutliers(suburd, nn.1 = 1, nn.2 = 20, x.max = 40, slope.r = 1.05,
    int.r = 4.2, slope.b = 0.75, int.b = 11.5, title = "Identifying Outliers by k-NN Distance.")
```

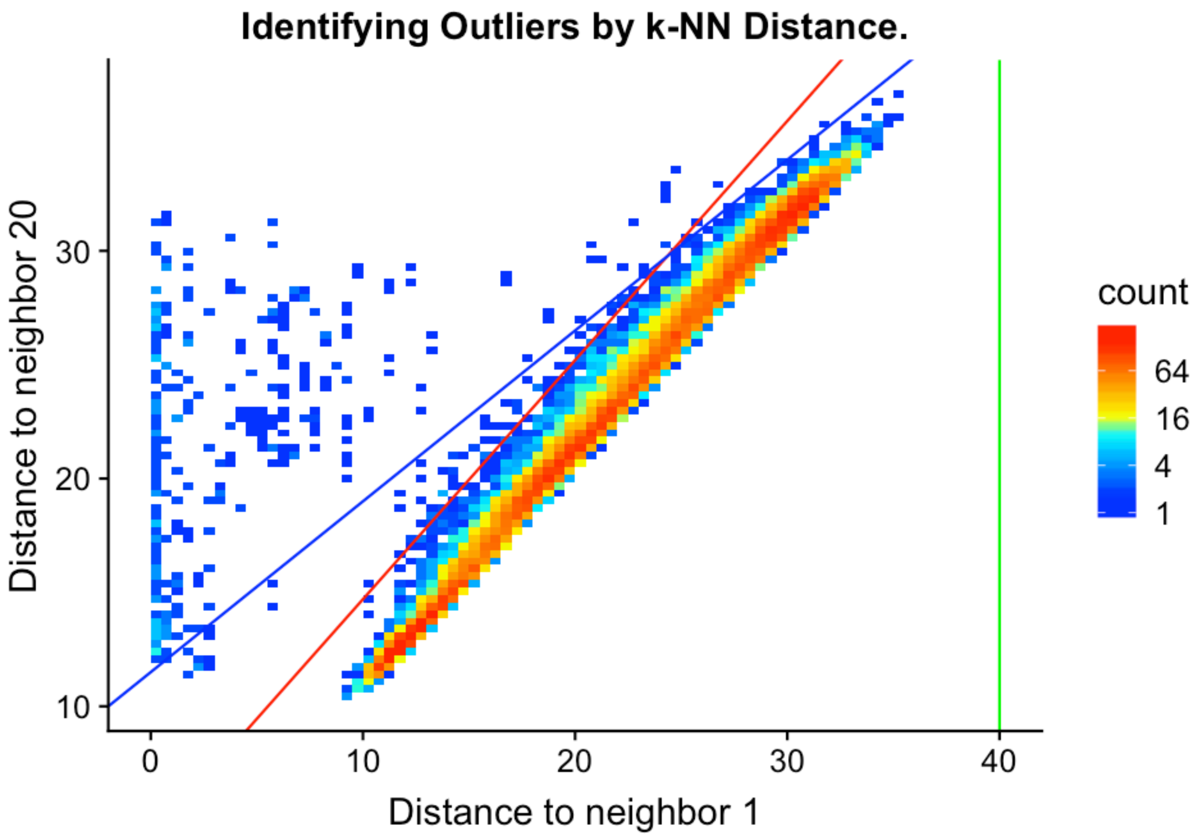

```
length(outliers)
```

```
## [1] 521
```

```
suburd <- urdSubset(suburd, cells.keep = setdiff(colnames(suburd@logupx.data),
    outliers))
```

#### Remove *kidins220a+* population

A cell cluster was observed in 15 dpf that was positive for expression of kidins220a, and foxg1b (which is exclusive to the retina). However, no similar clusters were observed in other stages, suggesting that we did not recover the progenitors of this population, so we excluded it from the URD analysis.

```
suburd <- urdSubset(suburd, cells.keep = setdiff(colnames(suburd@logupx.data),
    cellsInCluster(suburd, "cluster", "12-15d-96")))
```

#### Remove cell type doublets

##### Add UMAP projection

While not strictly required, a UMAP projection can make it easier to assess the expression of NMF modules and whether thresholds for overlap are set correctly.

```
## add UMAP command

# Load pre-calculated UMAP
umap <- readRDS(paste0(base.path, "/umap/umap_retina.rds"))

# Add projection to URD object
suburd@tsne.y <- umap[colnames(suburd@logupx.data), ]
```

##### Load NMF results and import into object

NMF results were calculated by providing `suburd@logupx.data` to an external NMF pipeline written in Python. The output results are imported here, scaled, and added to the URD object.

```
# Load the NMF results
load(paste0(base.path, "/NMF/retina/result_tbls.Robj"))

# The results object contains NMF runs for several K values.  k=45 was
# chosen for this tissue, so this extracts the results for that
# particular parameter
k.use <- "45"
nmf.cells <- result_obj[[paste0("K=", k.use)]][[1]]$C
rownames(nmf.cells) <- paste0("nmf", 1:nrow(nmf.cells))
colnames(nmf.cells) <- gsub("\\.", "-", colnames(nmf.cells))
nmf.genes <- result_obj[[paste0("K=", k.use)]][[1]]$G
colnames(nmf.genes) <- paste0("nmf", 1:nrow(nmf.cells))

# Some stages were subsampled in the original object and accidentally
# cropped out cells that had scGESTALT barcodes.  Those were added back
# in, and their expression was decomposed with original NMF gene matrix
# to give an additional NMF cell matrix for those cells.
new.nmf.c <- read.csv(paste0(base.path, "/NMF/retina/retina_new_nmfC_k45.csv"),
    row.names = 1)
rownames(new.nmf.c) <- paste0("nmf", 1:nrow(new.nmf.c))
colnames(new.nmf.c) <- gsub("\\.", "-", colnames(new.nmf.c))

# Combine old and new NMF results
nmf.cells <- cbind(nmf.cells, new.nmf.c)

# Trim NMF results to match cells in current object
nmf.cells <- nmf.cells[, colnames(suburd@logupx.data)]

# Scale NMF results 0-1
nmf.cells.scaled <- sweep(nmf.cells, 1, apply(nmf.cells, 1, max), "/")

# Add scaled NMF results to the URD object
suburd@nmf.c1 <- as(t(as.matrix(nmf.cells.scaled)), "dgCMatrix")
```

##### Select cell-type specific modules

Several NMF modules will be poor markers of cell types — these are often modules driven mostly be the expression of 1-2 genes (where the gene loading of the first gene is much greater than that of the fourth gene, for instance), or modules that don't exhibit any restriction in a tSNE or UMAP projection.

```
# Plot size parameters
plot.height = 8
plot.width = 8
dpi = 150

# Plot every module to determine which exhibit cell-type specificity
# This saves directly to the hard drive: two example plots are shown
# below.

# for (n in colnames(suburd@nmf.c1)) { png(paste0(path, '/doublets/',
# subset, '-plots/', n, '.png'), width=dpi*plot.width,
# height=dpi*plot.height) plot(plotDim(suburd, n)) dev.off() }

gridExtra::grid.arrange(grobs = list(plotDim(suburd, "nmf4", plot.title = "nmf4: strong cell-type restriction"),
    plotDim(suburd, "nmf2", plot.title = "nmf2: poor cell-type restriction")),
    ncol = 1)
```

```
## Warning: Removed 520 rows containing missing values (geom_point).

## Warning: Removed 520 rows containing missing values (geom_point).
```

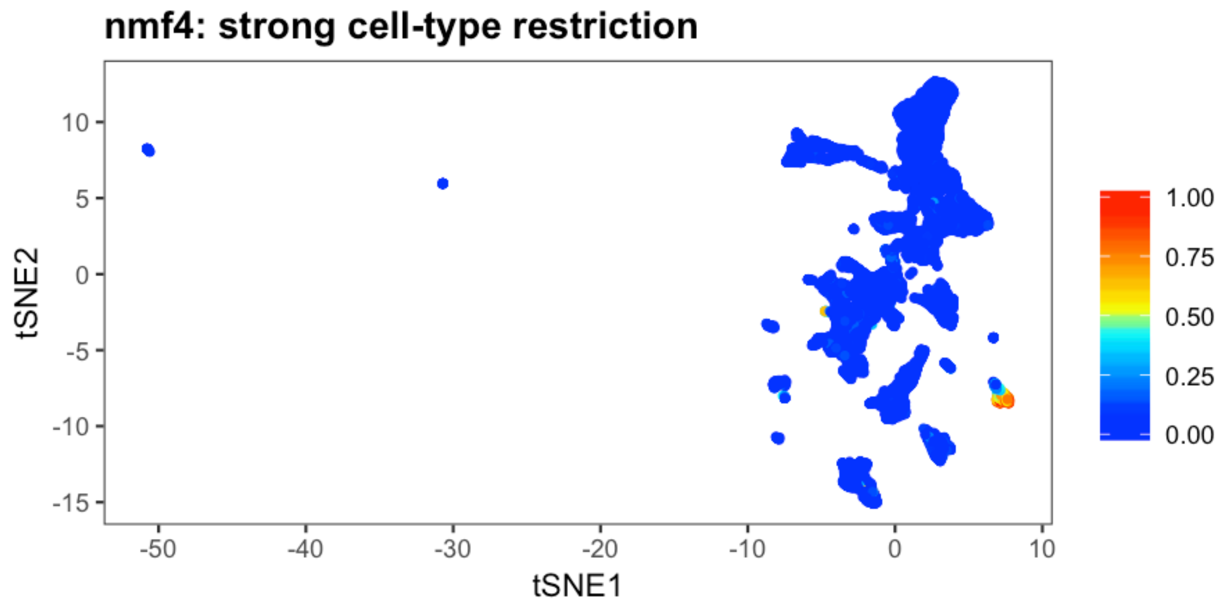

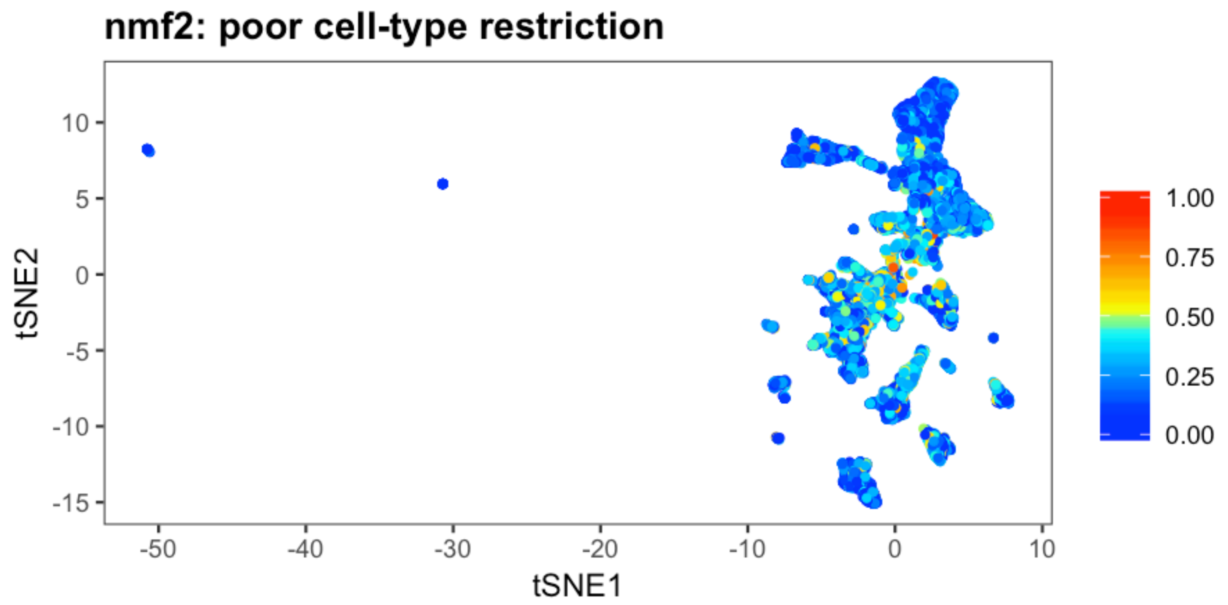

```r
# Module Gene 1 : Gene 4 Ratios
top.genes <- result_obj[[paste0("K=", k.use)]][[1]]$top30genes
top.weights <- top.genes[, grep("Weights", colnames(top.genes), value = T)]
colnames(top.weights) <- paste0("nmf", 1:nrow(nmf.cells))
top.weights.ratio <- top.weights[1, ]/top.weights[4, ]

# Which modules exhibit cell-type restriction?
modules.ok.ratio <- names(top.weights.ratio)[which(top.weights.ratio <
    5)]
restricted.modules <- paste0("nmf", c(4:5, 8:13, 15:18, 21:29, 31:35, 37:39,
    41:44))
good.modules <- intersect(modules.ok.ratio, restricted.modules)
```

Determine which module pairs to use for doublet removal

We consider NMF modules pairwise and only use those pairs that don't are non-overlapping in the data. (In other words, NMF modules that are mutually exclusive in the majority of the data.) Here, we determine thresholds for selecting those module pairs.

```
# Determine overlaps between module pairs
nmf.doublet.combos <- NMFDoubletsDefineModules(suburd, modules.use = good.modules,
    module.thresh.high = 0.4, module.thresh.low = 0.15)

# Determine thresholds for NMF modules
frac.overlap.max = 0.03
frac.overlap.diff.max = 0.1
module.expressed.thresh = 0.33

# Determine which module pairs to use for doublets
NMFDoubletsPlotModuleThresholds(nmf.doublet.combos, frac.overlap.max = frac.overlap.max,
    frac.overlap.diff.max = frac.overlap.diff.max)
```

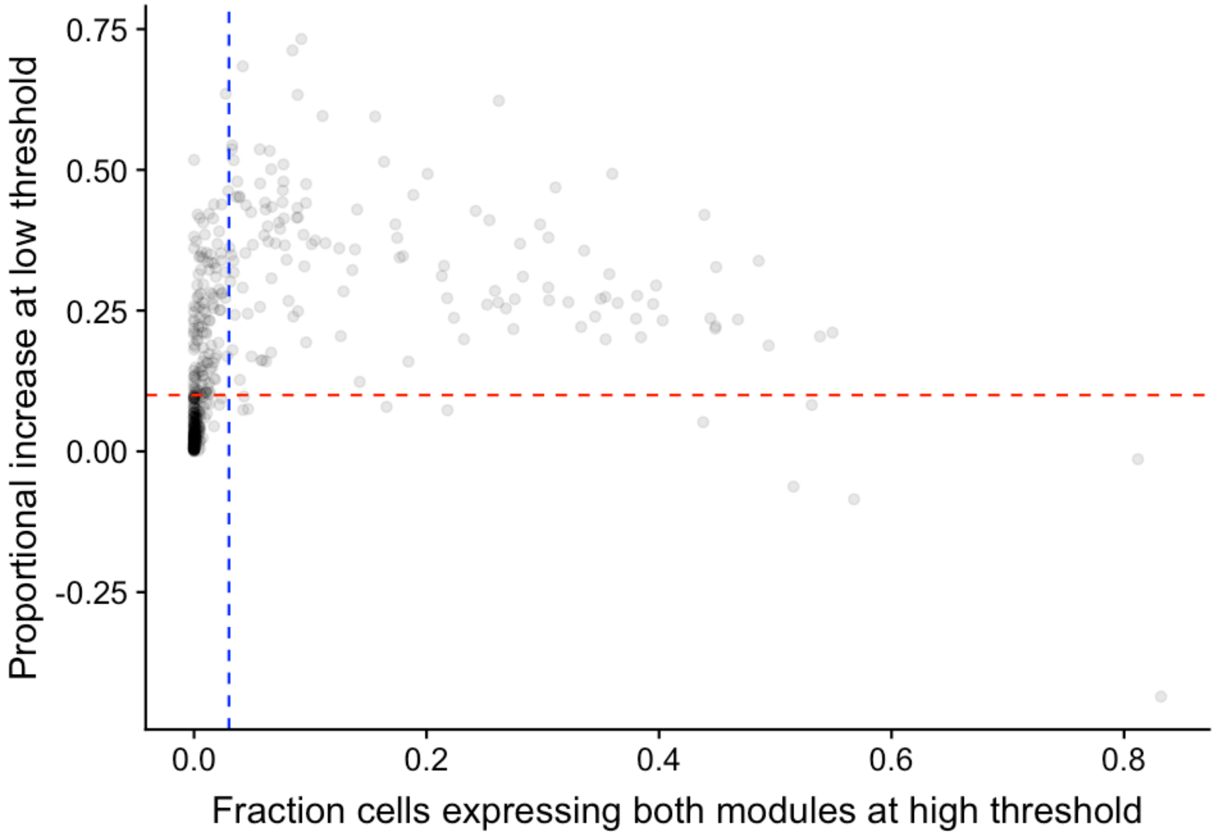

```
# These commands save plots directly to the hard-drive.

# Make plots to see how your thresholds are
NMFDoubletsPlotModuleCombos(suburd, path = paste0(path, "/doublets/", subset,
    "-doublet-combos/"), module.combos = nmf.doublet.combos, module.expressed.thresh = module.expressed.thresh,
    frac.overlap.max = frac.overlap.max, frac.overlap.diff.max = frac.overlap.diff.max,
    boundary = "pass", sort = "near", n.plots = 25)
NMFDoubletsPlotModuleCombos(suburd, path = paste0(path, "/doublets/", subset,
```

```
    "-ok-combos/"), module.combos = nmf.doublet.combos, module.expressed.thresh = module.expressed.thresh,
    frac.overlap.max = frac.overlap.max, frac.overlap.diff.max = frac.overlap.diff.max,
    boundary = "discarded", sort = "near", n.plots = 25)
```

```
# Define doublet cells
nmf.doublets <- NMFDoubletsDetermineCells(suburd, nmf.doublet.combos, module.expressed.thresh = module.expressed.
    frac.overlap.max = frac.overlap.max, frac.overlap.diff.max = frac.overlap.diff.max)  # 376 cells / 19837 cell

# Plot doublet cells on the UMAP
suburd <- groupFromCells(suburd, "nmf.doublets", cells = nmf.doublets)
plot(plotDimHighlight(suburd, clustering = "nmf.doublets", cluster = "TRUE",
    plot.title = paste0("NMF doublets: ", length(nmf.doublets), " cells"),
    point.size = 2, highlight.color = "blue"))
```

```
## Warning: Removed 520 rows containing missing values (geom_point).
```

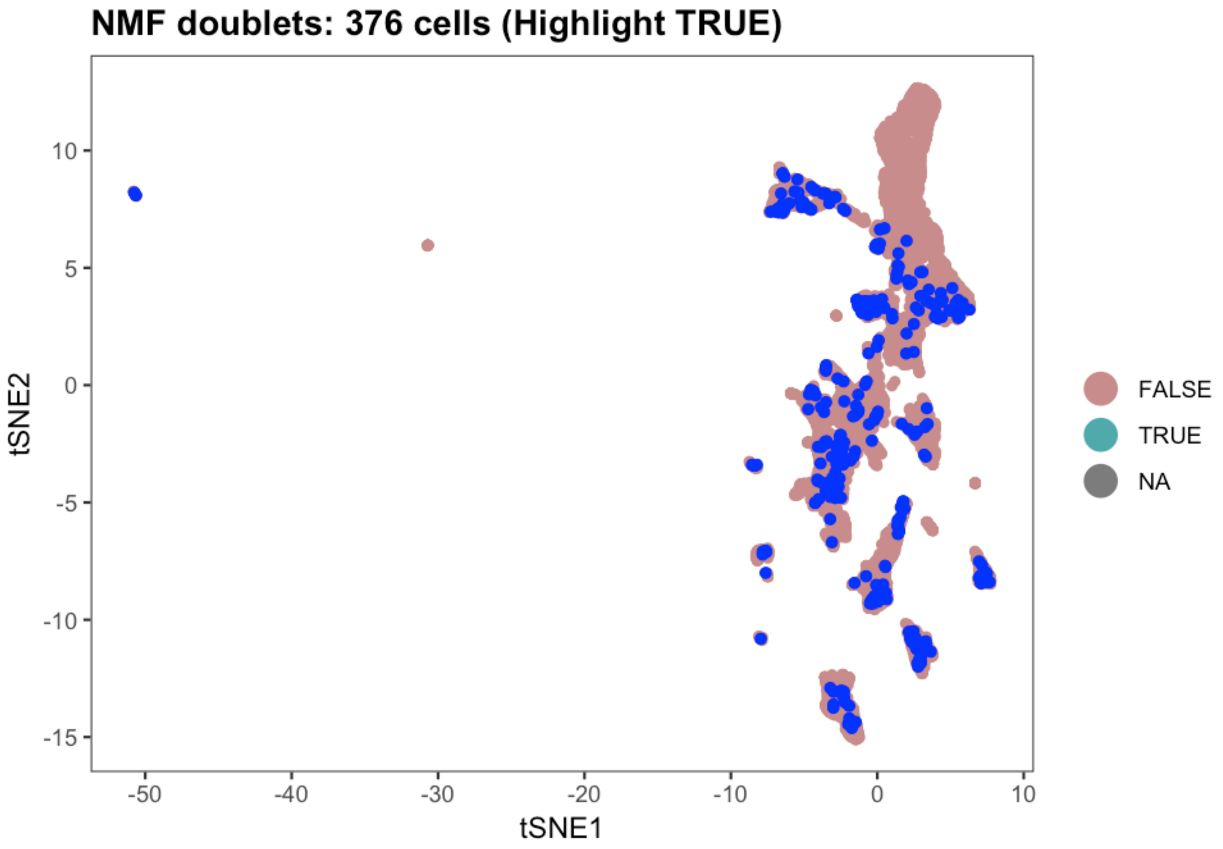

```
# Crop object to exclude doublets
suburd.cropped <- urdSubset(suburd, cells.keep = setdiff(colnames(suburd@logupx.data),
    nmf.doublets))
```

And then save the completed object for use downstream in building a tree using URD.

```
saveRDS(suburd.cropped, file = paste0(base.path, "/obj/URD_retina_ND.rds"))
```

### Retina: 2 - URD tree

*Jeff Farrell*

*9/07/2019*

Build the URD tree 12

Save the URD tree 13

#### Load data

```
suppressPackageStartupMessages(library(URD))
suppressPackageStartupMessages(library(Seurat))
```

```
base.path <- "~/urd-cluster-bushra/"

# Load procesed URD object
object <- readRDS(paste0(base.path, "obj/URD_retina_ND.rds"))
```

#### Processed on the cluster

Most of the following steps were run on a computing cluster. These individual tissue subsets can be run on a modern, well-equipped laptop. The use of a computing cluster allows multiple parameter choices to be tried in parallel, and also allows further parallelization of the random walk procedure, speeding it up. Below, we should the commands that one would run on their laptop, and then generally load the pre-processed results from the cluster that were used in the paper. If you want to parallelize your own processing on a compute cluster, the scripts we used will be available at http://github.com/farrellja/URD/cluster/

### Calculate diffusion map and pseudotime

These two steps are run in the cluster script URD-DM-PT.R.

#### Calculate diffusion map

```
# To run locally: Calculate a diffusion map projection
object <- calcDM(object, knn = 140, sigma.use = 14)
```

```
# Or: Load a pre-computed diffusion map projection
dm <- readRDS(paste0(base.path, "dm/dm_retinanewnokND_knn-140_sigma-14.rds"))
object <- importDM(object, dm)
```

```
# Plot diffusion maps
stage.colors <- c("antiquewhite", "#FFCCCC", "#99CC00", "#33CC00", "cyan3",
    "gold", "goldenrod", "darkorange", "indianred1", "plum", "deepskyblue2",
    "lightgrey")

# Plot by stage
plotDimArray(object = object, reduction.use = "dm", dims.to.plot = 1:18,
    label = "stage", plot.title = "", outer.title = "Diffusion map labeled by Stage",
    legend = T, alpha = 0.45, discrete.colors = stage.colors)
```

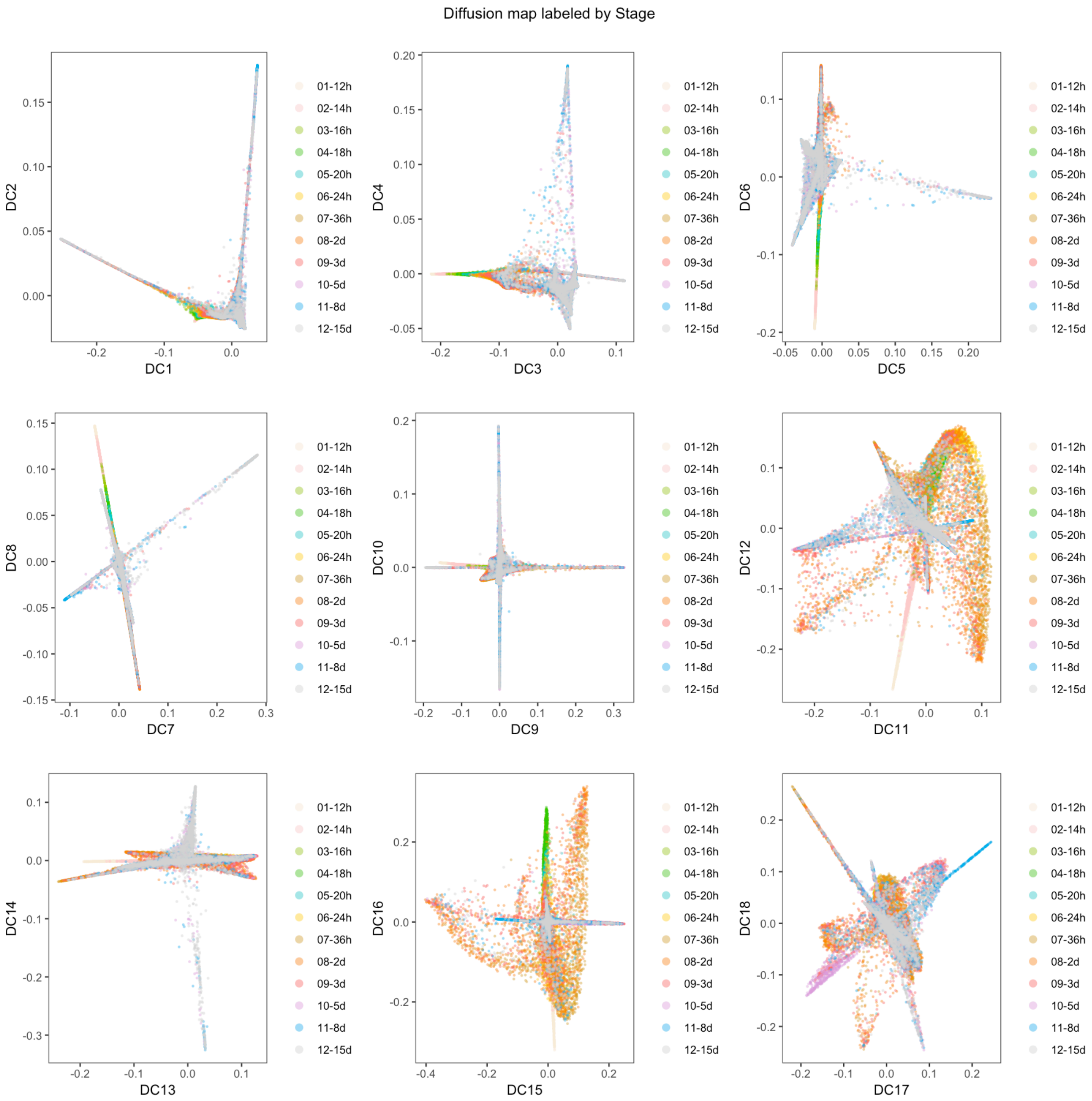

```
# Plot with final cell types labeled
object@group.ids$final.cluster <- NA
object@group.ids[cellsInCluster(object, "stage", "12-15d"), "final.cluster"] <- object@group.ids[cellsInCluster(
    "stage", "12-15d"), "res.5"]
plotDimArray(object = object, reduction.use = "dm", dims.to.plot = 1:18,
    label = "final.cluster", plot.title = "", outer.title = "Diffusion map with final clusters",
    legend = T, alpha = 0.6)
```

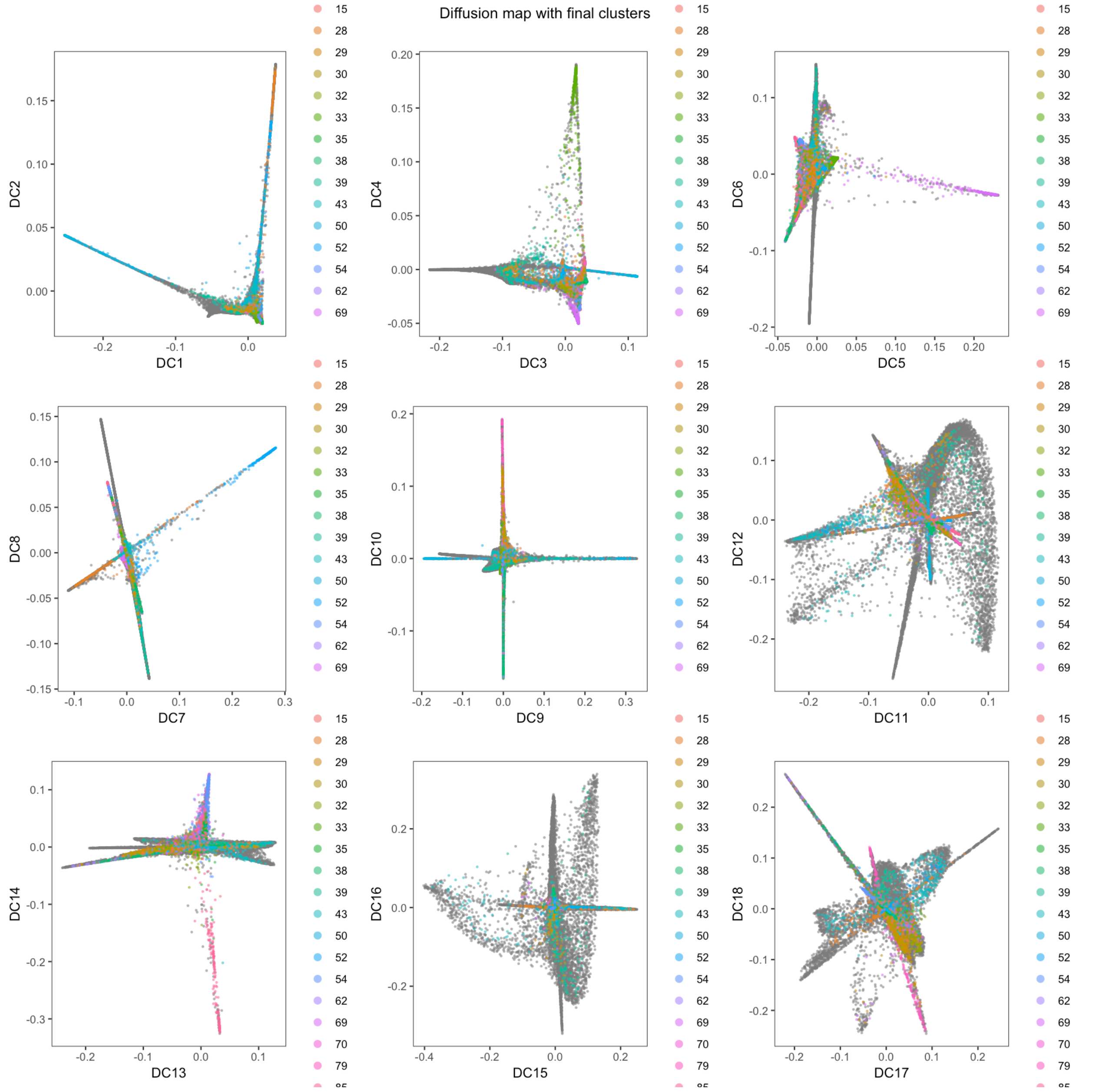

#### Calculate pseudotime

URD requires a starting point or 'root' for determining pseudotime. Here, we used all cells from the first timepoint (i.e. 12 hpf) as the root.

```
# Here, we used all cells from the first timepoint (i.e. 12 hours) as
# the root.
root.cells <- cellsInCluster(object, "stage", "01-12h")
plotDimHighlight(object, "stage", "01-12h", plot.title = "Root is 12 hpf cells")
```

```
## Warning: Removed 500 rows containing missing values (geom_point).
```

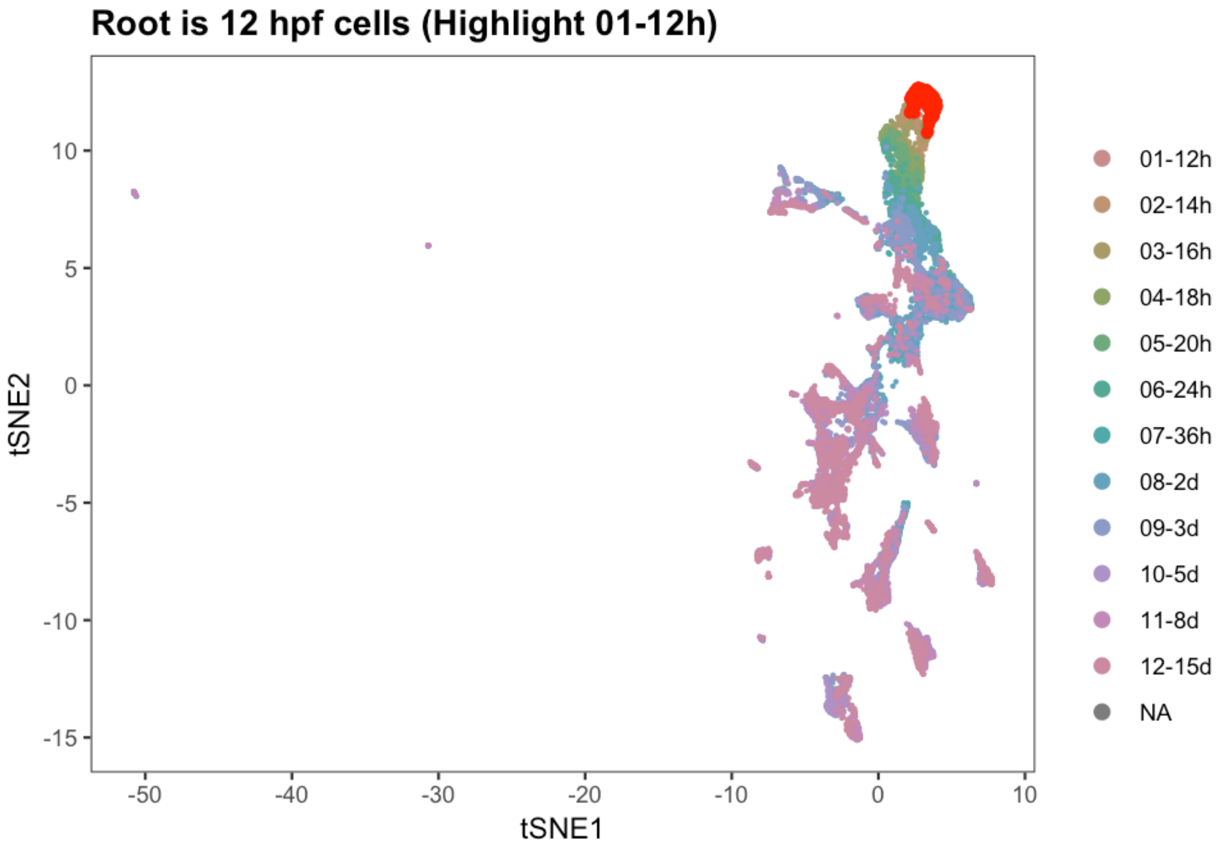

```r
# To run locally: Run graph-search simulations to determine pseudotime
flood.result <- floodPseudotime(object, root.cells = root.cells, n = 100,
    minimum.cells.flooded = 2, verbose = T)
```

```r
# Or load a pre-computed graph-search simulation result
flood.result <- readRDS(paste0(base.path, "flood/flood_retinanewnokND_knn-140_sigma-14.rds"))
```

```r
# Process the graph-search simulations to determine the pseudotime of
# each cell
object <- floodPseudotimeProcess(object, flood.result, floods.name = "pseudotime",
    max.frac.NA = 0.4, pseudotime.fun = mean, stability.div = 10)

# If enough simulations have been run, then as additional simulations
# are added, the overall change in pseudotime of cells should reach an
# asymptote. If it does not, then floodPseudotime should be run with a
# higher n.
pseudotimePlotStabilityOverall(object)
```

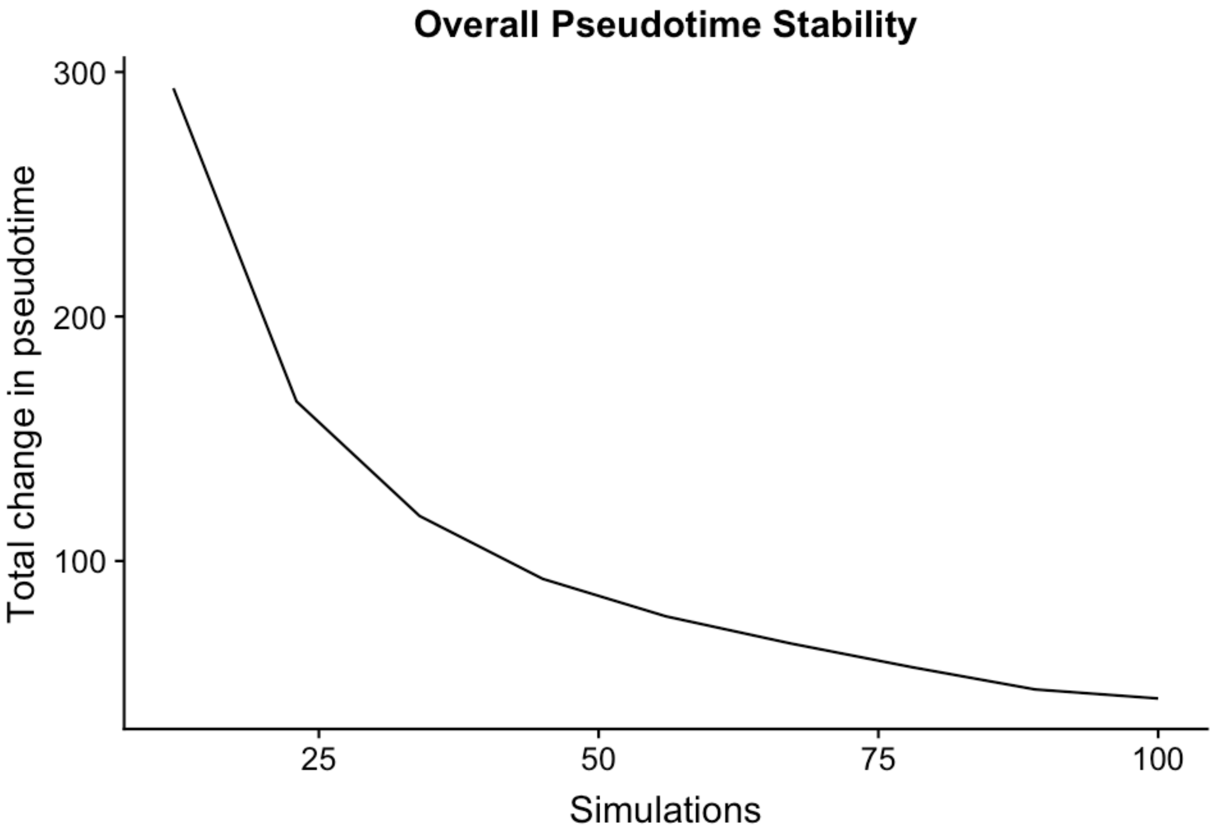

```r
plotDimArray(object = object, reduction.use = "dm", dims.to.plot = 1:18,
    label = "pseudotime", plot.title = "", outer.title = "Diffusion Map labeled by pseudotime",
    legend = F, alpha = 0.4)
```

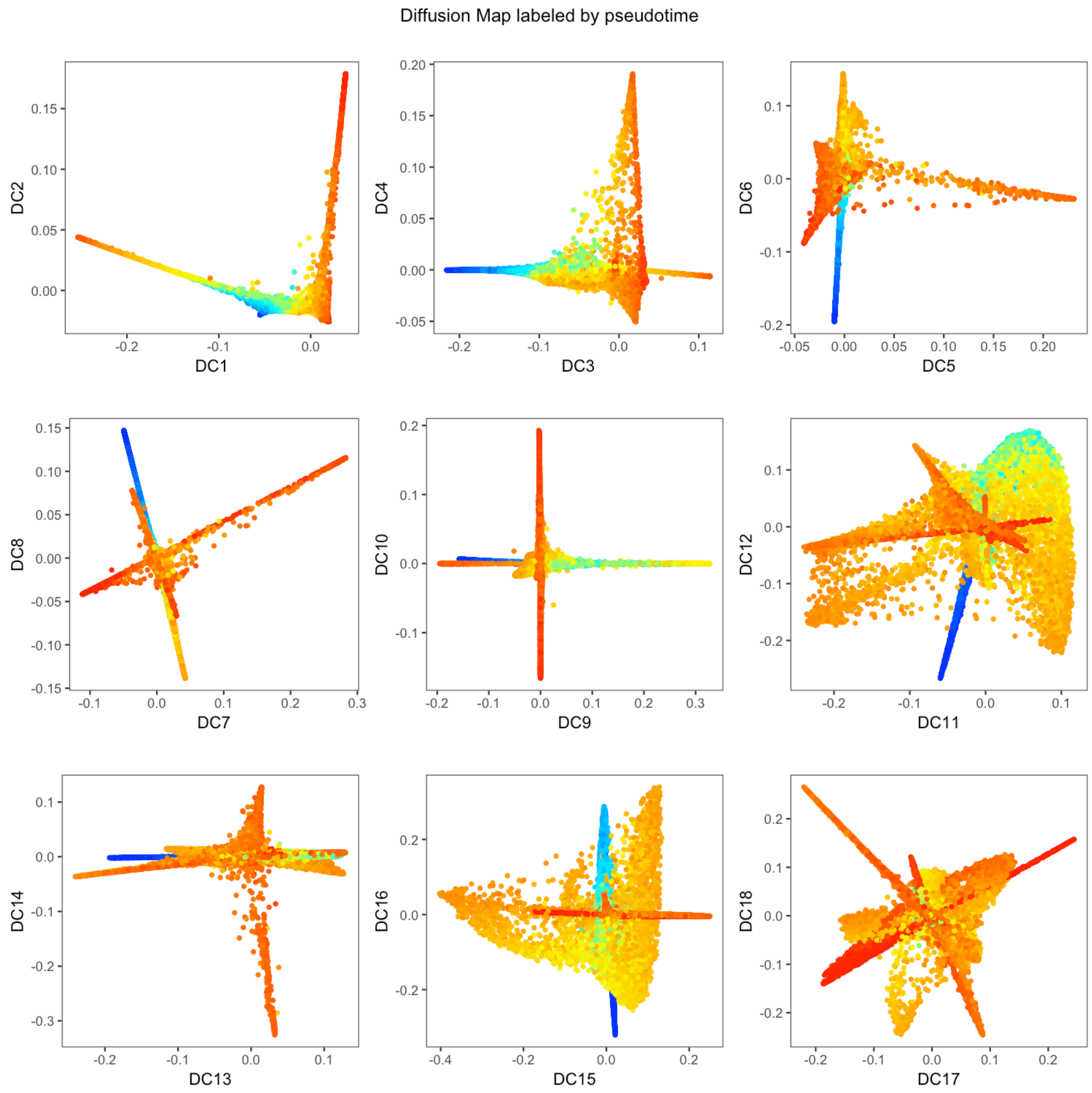

```r
plotDim(object, "pseudotime", plot.title = "UMAP projection colored by pseudotime")
```

```
## Warning: Removed 500 rows containing missing values (geom_point).
```

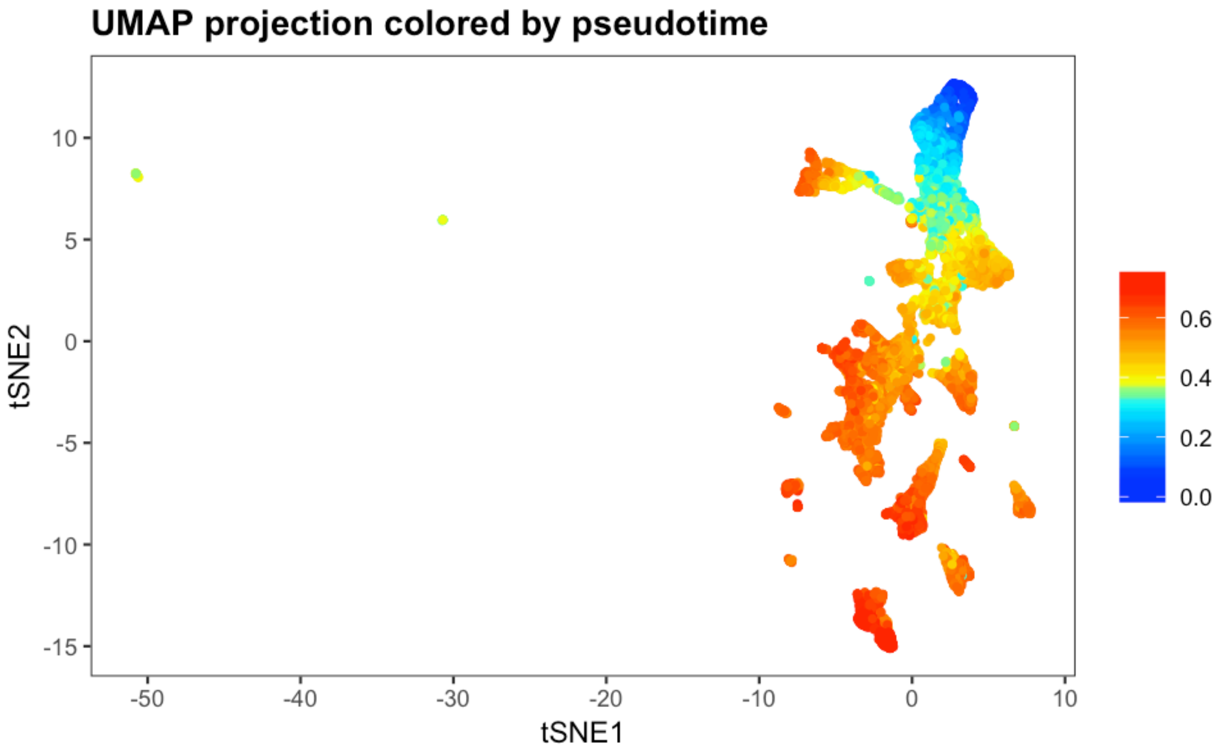

```r
plotDists(object, "pseudotime", "stage", plot.title = "Pseudotime by stage")
```

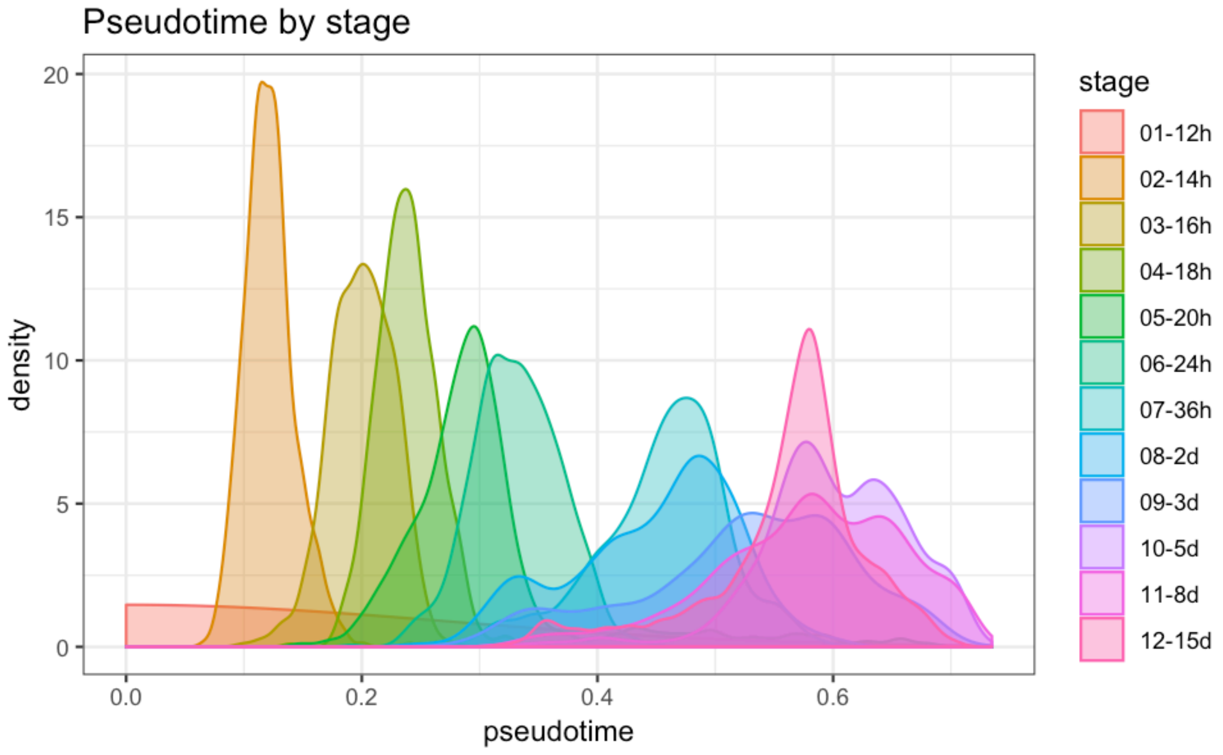

### Calculate biased transition matrix

In order to perform biased random walks, we must first bias the transition matrix to ensure that walks proceed towards the root and do not turn into other differentiated cell types. This is performed in the cluster script **URD-TM.R**.

```
# Calculate parameters for biasing the transition matrix.
diffusion.logistic <- pseudotimeDetermineLogistic(object, "pseudotime",
    optimal.cells.forward = 40, max.cells.back = 80, pseudotime.direction = "<",
    do.plot = T, print.values = T)
```

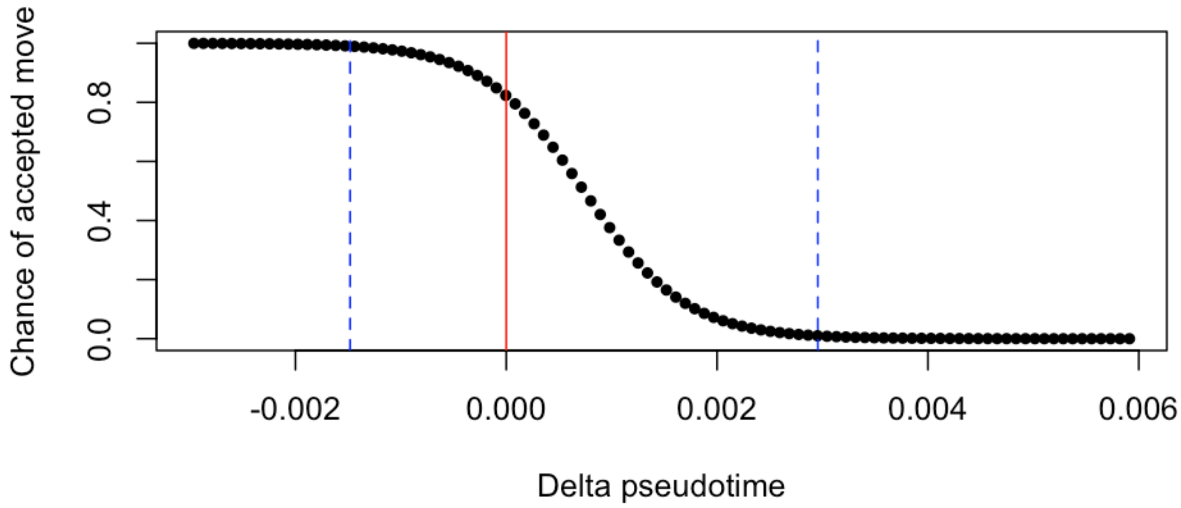

```
## [1] "Mean pseudotime back (~80 cells) 0.00295569803650678"
## [1] "Chance of accepted move to equal pseudotime is 0.822024945232085"
## [1] "Mean pseudotime forward (~40 cells) -0.00148141113789574"
```

```
# Calculate the biased matrix.
biased.tm <- pseudotimeWeightTransitionMatrix(object, pseudotime = "pseudotime",
    logistic.params = diffusion.logistic, pseudotime.direction = "<")
```

### Perform biased random walks

Then, we perform biased walks starting from each tip. Visited cells are inferred to lie along the trajectory that connects the root to each cell type. This is performed in the cluster script **URD-Walk.R**.

#### Determine tips

We used clusters from 15 dpf as the tips for performing biased random walks. Here we define the cells belonging to each of those clusters.

```r
# All clusters at 15 days
clusters.15day <- unique(object@group.ids[grep("15d", object@group.ids$stage),
    "res.5"])
# All cells at 15 days
cells.15day <- rownames(object@group.ids)[grep("15d", object@group.ids$stage)]
# Cell lists of each cluster at 15dpf
cells.15dpf.clusters <- lapply(clusters.15day, function(clust) intersect(cells.15day,
    cellsInCluster(object, "res.5", clust)))
names(cells.15dpf.clusters) <- paste0("15d-", clusters.15day)
```

We also load a .csv file that contains information about the tips. It has four columns:

- id: Cluster ID for the tip
- use: Whether this cluster should be used when building the tree
- name: The name for this tip, which will be used on 2D plots
- short.name: The 'short' name for this tip, which would be used on 3D plots (though we did not use that feature in this study).

```r
# Load CSV
tip.names <- read.csv(paste0(base.path, "tips/tip_names_retinanewnokND.csv"),
    header = F, stringsAsFactors = F, colClasses = c("character", "logical",
        "character", "character"))

# Name columns and rows
names(tip.names) <- c("id", "use", "name", "short.name")
rownames(tip.names) <- gsub("_", "-", tip.names$id)

# Sort alphabetically
tip.names <- tip.names[order(rownames(tip.names)), ]
```

These are the tips that were considered during the construction of the retina URD tree (some were excluded during tree construction later in the `buildTree` command).

```r
# Define a 'tips' clustering
object@group.ids$tip <- NA
object@group.ids$tip.id <- NA
object@group.ids$tip.name <- NA

# If the tip will be used in the tree, define its cells in the
# clustering
for (i in 1:nrow(tip.names)) {
    tip.cells <- cells.15dpf.clusters[[rownames(tip.names)[i]]]
    object@group.ids[tip.cells, "tip"] <- as.character(i)
    object@group.ids[tip.cells, "tip.id"] <- rownames(tip.names)[i]
    object@group.ids[tip.cells, "tip.name"] <- as.character(tip.names[i,
        "name"])
}

# Plot the tips
plotDim(object, "tip.name")
```

```
## Warning: Removed 500 rows containing missing values (geom_point).
```

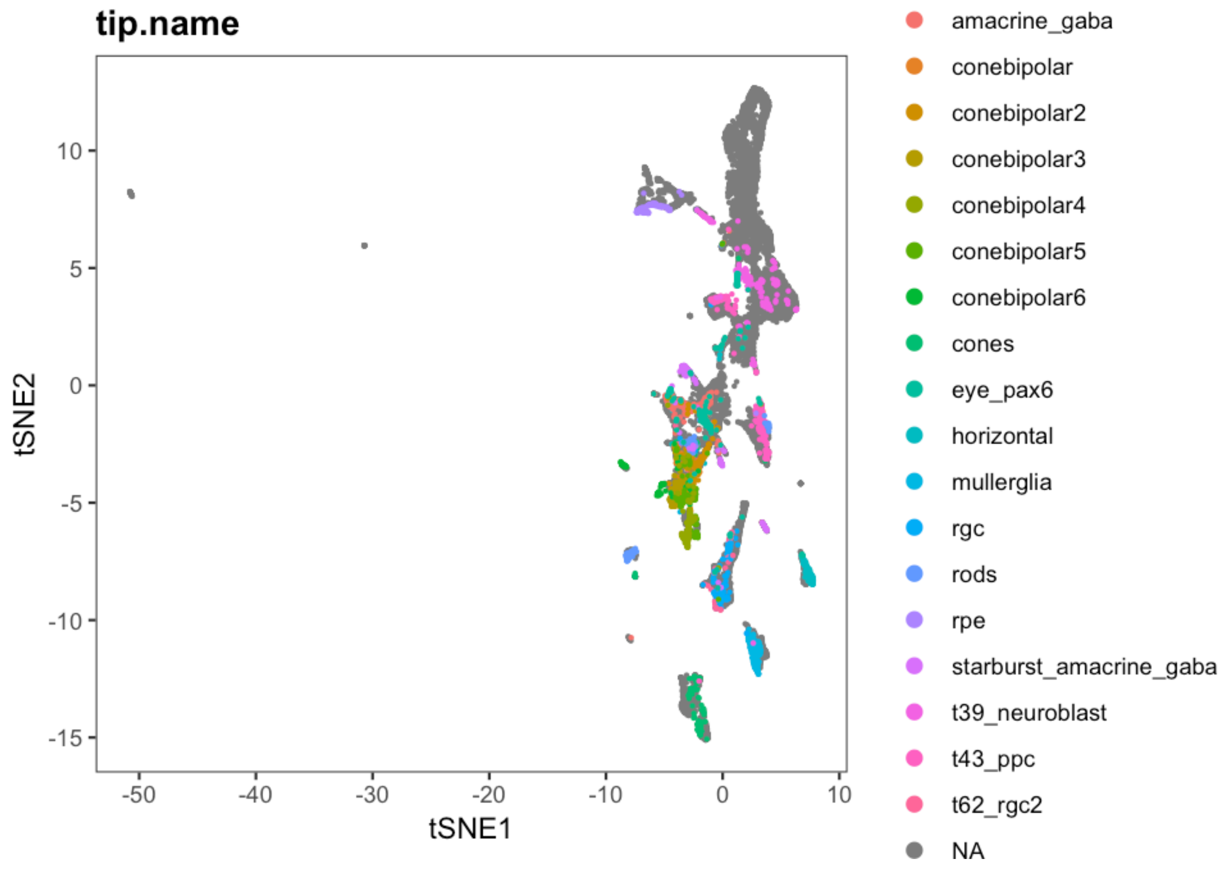

#### Perform the biased random walks

Biased random walks then need to be run starting from each tip. This can be performed on a laptop, but is an ideal candidate for parallelization on a cluster. (The walks from each tip can be run as a separate job.)

```
## IF RUNNING LOCALLY

# Loop through each cluster
walks <- lapply(rownames(tip.names), function(c) {
    # Exclude any tip cells that for whatever reason didn't end up in the
    # biased TM (e.g. maybe not assigned a pseudotime).
    tip.cells <- intersect(cells.15dpf.clusters[[c]], rownames(biased.tm))
    # Perform the random walk simulation
    this.walk <- simulateRandomWalk(start.cells = tip.cells, transition.matrix = biased.tm,
        end.cells = root.cells, n = 50000, end.visits = 1, verbose.freq = 1000,
        max.steps = 5000)
    return(this.walk)
})
names(walks) <- rownames(tip.names)

# Alternatively, this loop is automated by the function
# simulateRandomWalksFromTips
```

Alternatively, a set of pre-calculated walks can be loaded. Since the walks are a simulation (and therefore not deterministic), this is particularly crucial for reproducing results.

```
## IF LOADING PRE-CALCULATED WALKS

# Get list of files in the walks directory
walks.files <- list.files(paste0(base.path, "/walks/retinanewnokND/"),
    pattern = ".rds")

# Load the walks previously performed for each cluster
walks <- lapply(rownames(tip.names), function(c) {
    walk.file <- grep(pattern = paste0("_tip-", c, "_"), x = walks.files,
        value = T)[1]
    return(readRDS(paste0(base.path, "/walks/retinanewnokND/", walk.file)))
})
names(walks) <- rownames(tip.names)
```

##### Process the random walks

The walks are then converted to visitation frequency by importing them into the URD object.

```
for (i in 1:nrow(tip.names)) {
    # Load the individual walk visitation frequencies into the object
    object <- processRandomWalks(object, walks = walks[[i]], walks.name = i,
        n.subsample = 1, verbose = F)
}
```

#### Build the URD tree

Then, a branching tree is constructed, by joining trajectories in an agglomerative fashion when cells are highly visited by walks from multiple tips. The following steps were performed in the cluster script URD-Tree.R.

```
# Tree building is destructive, so create a copy of the object
object.tree <- object

# Load tip cells
object.tree <- loadTipCells(object.tree, "tip")

# Determine tips to use
tips.to.use <- which(tip.names$use)
```

```
# Build the tree
object.tree <- buildTree(object.tree, pseudotime = "pseudotime", divergence.method = "preference",
    cells.per.pseudotime.bin = 40, bins.per.pseudotime.window = 5, save.all.breakpoint.info = T,
    p.thresh = 0.01, verbose = F, tips.use = as.character(tips.to.use))
```

```
# Name the tips of the tree
object.tree <- nameSegments(object.tree, segments = tips.to.use, segment.names = as.character(tip.names[tips.to.u
    "name"], short.names = as.character(tip.names[tips.to.use, "short.name"])))
```

```
plotTree(object.tree, "stage", discrete.colors = stage.colors, label.segments = T)
```

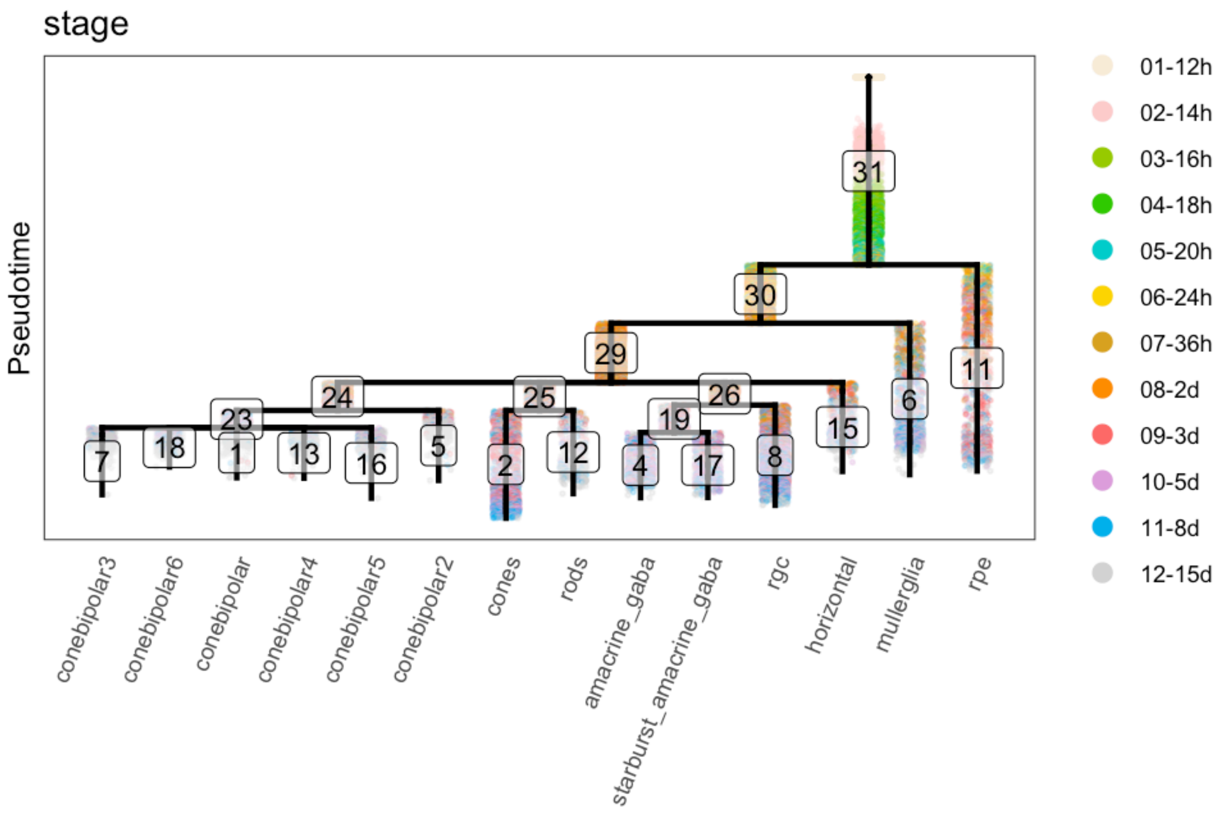

#### Save the URD tree

The tree is then saved for use in downstream analysis, and can easily be loaded for further perusal.

```
saveRDS(object.tree, file = paste0(base.path, "tree/URD-Tree-Retina.rds"))
```

### Retina: 3 - URD Cascades and Figures

*Jeff Farrell*

*10/08/2019*

### Load data

```
# Load URD
library(URD)
```

```
## Loading required package: ggplot2

## Warning: package 'ggplot2' was built under R version 3.4.4

## Loading required package: Matrix

## Warning: package 'Matrix' was built under R version 3.4.4
```

```
# Basic location
base.path <- "~/urd-cluster-bushra/"

# Load completed retina tree object
obj.path <- paste0(base.path, "tree/retinanewnokND/tree-retinanewnokND_knn-140_sigma-14_40F-80B_NO-15d-29-15d-39-
obj <- readRDS(obj.path)
```

### Plot gene expression on the tree

#### Plot tree by stage

```
stage.colors <- c("antiquewhite", "#FFCCCC", "#99CC00", "#33CC00", "cyan3", "gold",
    "goldenrod", "darkorange", "indianred1", "plum", "deepskyblue2", "lightgrey")

plotTree(obj, "stage", label.type = "group", discrete.colors = stage.colors)
```

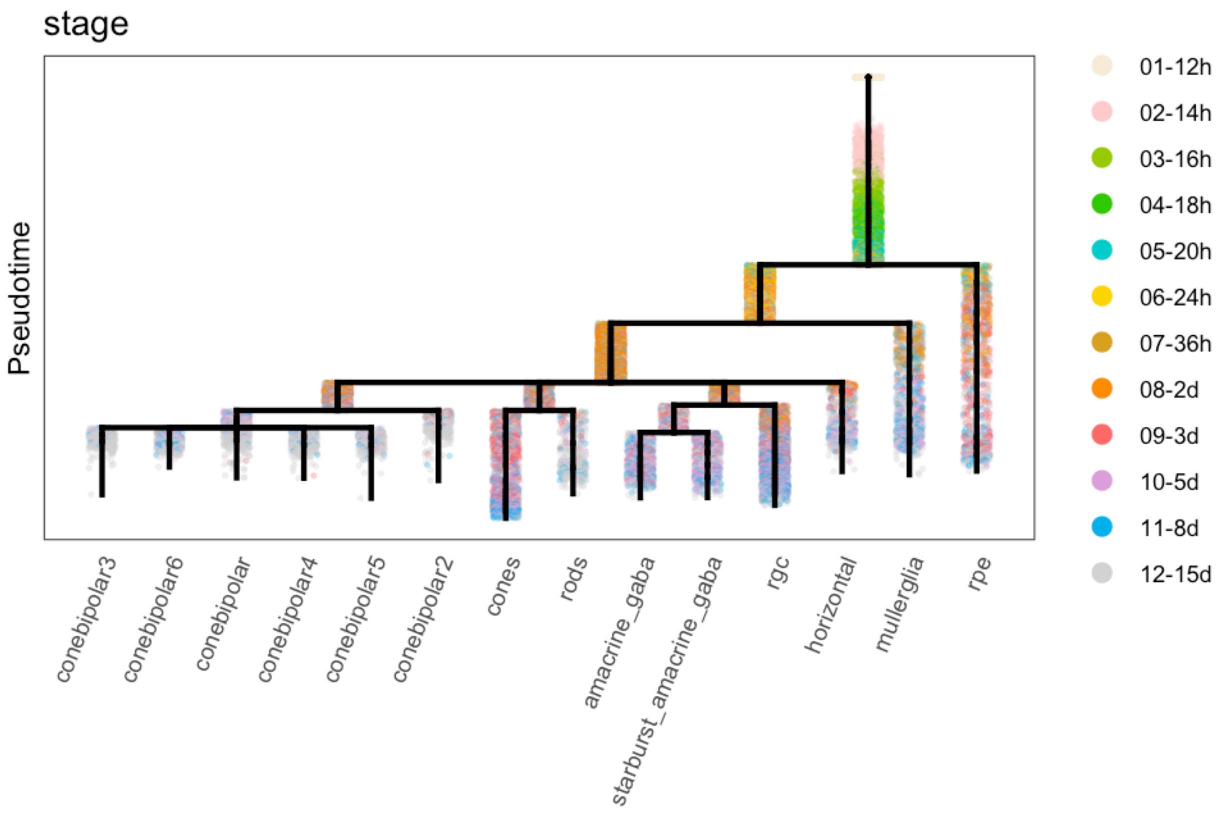

#### Plot tree with gene expression: main figures

```
gridExtra::grid.arrange(grobs = lapply(c("vsx1", "lmo4a", "pax6a", "rem1"), plotTree,
    object = obj, label.x = F, plot.cells = F), ncol = 2)
```

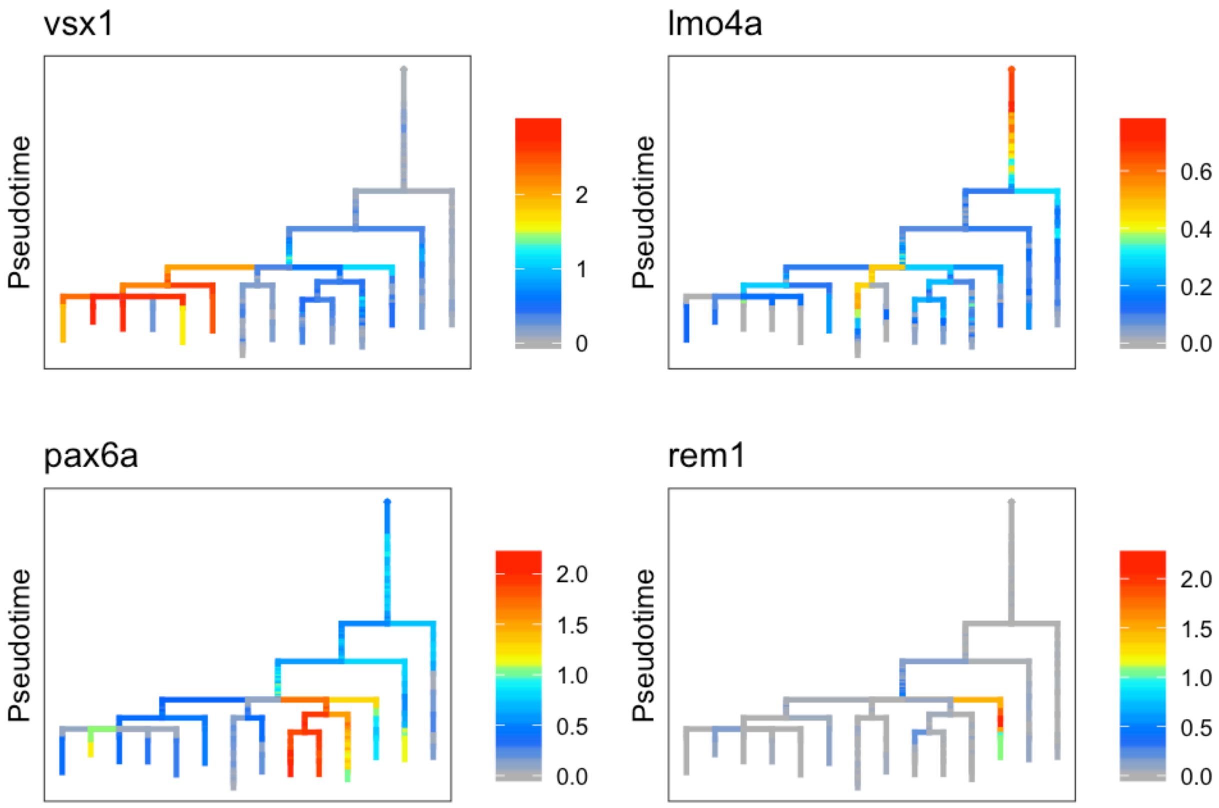

#### Plot tree with gene expression: supplemental figures

```
gridExtra::grid.arrange(grobs = lapply(c("foxd1", "her15.1", "her4.1", "hes2.2",
    "stx3a", "dmbx1a", "prkca", "rx3", "irx7", "fezf2", "ndrg1b", "opn1mw1", "gnat1",
    "pbx1a", "tfap2c", "slc18a3a", "rbpms2a", "ompa", "sdpra", "rpe65a"), plotTree,
    object = obj, label.x = F, plot.cells = F), ncol = 4)
```

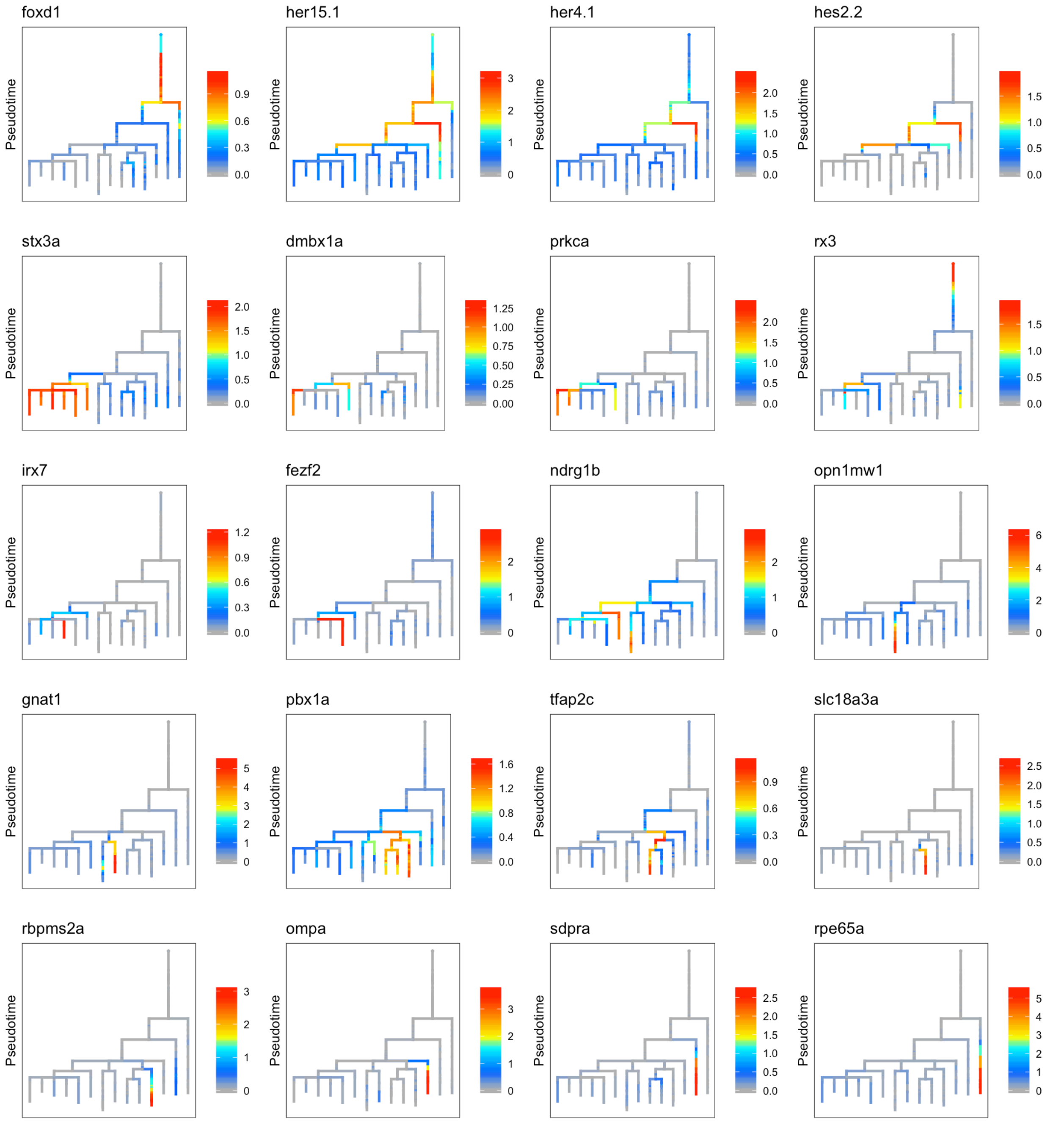

### Determine genes enriched in trajectories to particular cell types

#### Comparison between major cell types

We took each major group ("clade") of branches from the end of the tree as a single entity (i.e. cone bipolar cells, photoreceptors, amacrine cells, retinal ganglion cells, horizontal cells) and compared them against each other pairwise to look for differentially expressed genes.

```r
# Get the parent segment of each clade to consider as a group
combined.tips <- c("24", "25", "19", "8", "15")

# Get the cells in that segment and all child segments
cells.combined.tips <- lapply(combined.tips, function(t) whichCells(obj, label = "segment",
    value = segChildrenAll(obj, t, include.self = T)))
names(cells.combined.tips) <- combined.tips

# Loop through each of these clades and look for differentially expressed genes
combined.markers <- lapply(combined.tips, function(tip) {
    # Find all of the other clades
    opposing.tips <- setdiff(combined.tips, tip)
    # Perform pairwise comparisons to each other clade
    m.o <- lapply(opposing.tips, function(tip.opposing) {
        # message(paste0(Sys.time(), ': Comparing tip ', tip, ' to ', tip.opposing, '.'))
        # Find differentially expressed genes between the pair of clades
        ma <- markersAUCPR(object = obj, cells.1 = cells.combined.tips[[tip]], cells.2 = cells.combined.tips[[tip
            effect.size = 0.4, auc.factor = 1.1)
        # In order to facilitate combining all of the results later, add columns about
        # which two clades were compared and also a duplicate entry of the name of each
        # gene that's recovered.
        ma$gene <- rownames(ma)
        ma$tip1 <- tip
        ma$tip2 <- tip.opposing
        return(ma)
    })
    names(m.o) <- opposing.tips
    return(m.o)
})
names(combined.markers) <- combined.tips

# Require that genes are markers against at least 3 other clades
combined.markers.beatmult <- lapply(combined.markers, function(m) {
    names(which(table(unlist(lapply(m, rownames))) >= 3))
})

# Since genes might be a marker in a comparison to several other clades, combine
# the results into a single table, where each gene is listed only once with the
# info from the pairwise comparison where it had the strongest differential
# expression.
combined.markers.best <- lapply(1:length(combined.markers.beatmult), function(i) {
    cm <- do.call("rbind", combined.markers[[i]])
    cm <- cm[cm$gene %in% combined.markers.beatmult[[i]], ]
    cmb <- do.call("rbind", lapply(combined.markers.beatmult[[i]], function(g) {
        cmr <- cm[cm$gene == g, ]
        return(cmr[which.max(cmr$AUCPR.ratio), ])
    }))
    rownames(cmb) <- cmb$gene
    cmb <- cmb[order(cmb$AUCPR.ratio, decreasing = T), ]
    cmb$exp.global <- apply(obj@logupx.data[rownames(cmb), unlist(obj@tree$cells.in.segment)],
        1, mean.of.logs)
    cmb$exp.global.fc <- cmb$nTrans_1 - cmb$exp.global
    return(cmb)
```

```
})
names(combined.markers.best) <- combined.tips
```

##### AUCPR along tree

We also used the AUCPRTestAlongTree function to ask for genes that are differential markers of a lineage using URD's tree structure. This makes a comparison at each branchpoint from a particular cell type up to the root.

```
# Get all of the tips from the tree
tips.in.tree <- as.character(obj@tree$tips)

# Tree segments to use as root for each particular cell population.
roots <- rep("29", length(tips.in.tree))
names(roots) <- tips.in.tree
roots["11"] <- "31"
roots["6"] <- "30"
roots[c("4", "17", "8")] <- "26"

# Perform a loop of tests with each tip.
markers <- lapply(tips.in.tree, function(t) {
    this.root <- roots[t]
    # message(paste0(Sys.time(), ': Starting tip ', t, ' and root ', this.root))
    these.markers <- aucprTestAlongTree(obj, pseudotime = "pseudotime", tips = as.character(t),
        genes.use = NULL, must.beat.sibs = 0.6, report.debug = F, root = this.root,
        auc.factor = 1.1, log.effect.size = 0.4)
    these.markers$gene <- rownames(these.markers)
    these.markers$tip <- t
    return(these.markers)
})
names(markers) <- tips.in.tree
```

#### Functions for curating differential expression results

We further curated those differentially expressed genes using the following functions:

##### threshold.tree.markers

Function to threshold markers from a markersAUCPRAlongTree test with additional criteria

- markers: list of results from markersAUCPRAlongTree tests
- tip: which tip (or element of the list to pursue)
- global.fc: fold.change that gene must have along the trajectory pursued vs. rest of the data
- aucpr.ratio.all: classifier score that gene must exhibit along trajectory test vs. rest of the data
- branch.fc: fold.change that gene must have (in best case) vs. the opposing branch at any branchpoint along the trajectory.
- Returns `markers` with only a subset of rows retained.

```
threshold.tree.markers <- function(markers, tip, global.fc = 0.1, branch.fc = 0.4,
    aucpr.ratio.all = 1.03) {
    m <- markers[[tip]]
    # First off -- lose global FC < x
    bye.globalfc <- rownames(m)[m$expfc.all < global.fc]
    # Second -- get rid of branch FC < x
    bye.branchfc <- rownames(m)[m$expfc.maxBranch < branch.fc]
    # Third -- get rid of stuff essentially worse than random classification on
    # global level
    bye.badglobalaucpr <- rownames(m)[m$AUCPR.ratio.all < aucpr.ratio.all]
    bye.all <- unique(c(bye.globalfc, bye.branchfc, bye.badglobalaucpr))
    m.return <- m[setdiff(rownames(m), bye.all), ]
    return(m.return)
}
```

#### threshold.clade.markers

Function to threshold markers of particular clades (see "Combined major branch families") using additional criteria

- **markers**: result of `markersAUCPR`
- **global.fc**: fold.change that gene must have along the trajectory pursued vs. rest of the data (during testing, branches were compared pairwise. This compares one branch to all others together.)
- Returns `markers` with a subset of rows retained

```
threshold.clade.markers <- function(markers, global.fc = 0.1) {
    m <- markers
    # First off -- lose global FC < x
    bye.globalfc <- rownames(m)[m$exp.global.fc < global.fc]
    m.return <- m[setdiff(rownames(m), bye.globalfc), ]
    return(m.return)
}
```

#### divide.branches

Function to compare genes between two branches. Use this on a compiled list of markers to do a final selection of genes that are specific to one branch or another or markers of both (i.e. when making photoreceptor heatmap, use to divide into photoreceptor, cone, and rod markers)

- **object**: An URD object
- **genes**: (Character vector) Genes to test
- **clust.1**: (Character) Cluster 1
- **clust.2**: (Character) Cluster 2
- **clustering**: (Character) Clustering to pull from
- **exp.fc**: (Numeric) Minimum expression fold-change between branches to consider different
- **exp.thresh**: (Numeric) Minimum fraction of cells in order to consider gene expressed in a branch
- **exp.diff**: (Numeric) Minimum difference in fraction of cells expressing to consider gene differential
- Returns list of gene names ("specific.1" = specific to clust.1, "specific.2" = specific to clust.2, "markers" = all genes tested)

```
divide.branches <- function(object, genes, clust.1, clust.2, clustering = "segment",
    exp.fc = 0.4, exp.thresh = 0.1, exp.diff = 0.1) {
    # Double check which markers are unique to one or the other population
    mcomp <- markersAUCPR(object, clust.1 = clust.1, clust.2 = clust.2, clustering = clustering,
        effect.size = -Inf, auc.factor = 0, genes.use = genes, frac.min.diff = 0,
        frac.must.express = 0)
    specific.b <- rownames(mcomp)[abs(mcomp$exp.fc) > exp.fc & mcomp[, 4] < exp.thresh &
        mcomp[, 5] > pmin((mcomp[, 4] + exp.diff), 1)]
    specific.a <- rownames(mcomp)[abs(mcomp$exp.fc) > exp.fc & mcomp[, 5] < exp.thresh &
        mcomp[, 4] > pmin((mcomp[, 5] + exp.diff), 1)]
    r <- list(specific.a, specific.b, mcomp)
    names(r) <- c("specific.1", "specific.2", "markers")
    return(r)
}
```

### Functions for heatmap generation

These functions were used in the production of heatmaps:

#### Color scale

Generate color scale to use with heatmaps.

```
cols <- (scales::gradient_n_pal(RColorBrewer::brewer.pal(9, "YlOrRd")))(seq(0, 1,
    length.out = 50))
```

#### determine.timing

Determines order to plot genes in heatmap. "Expression" is defined as 20% higher expression than the minimum observed value. "Peak" expression is defined as 50% higher expression than minimum observed value. The two longest stretches of "peak" expression are found, and then the later one is used. The onset time of the stretch of expression that contains that peak is also determined. Genes are then ordered by the pseudotime at which they enter "peak" expression, leave "peak" expression, start "expression", and leave "expression".

- s: result from `geneSmoothFit`
- genes: genes to order; default is all genes that were fit.
- Returns `s` but with an additional list entry (`$timing`) of the order to plot genes

```
determine.timing <- function(s, genes = rownames(s$mean.expression)) {
    s$timing <- as.data.frame(do.call("rbind", lapply(genes, function(g) {
        sv <- as.numeric(s$scaled.smooth[g, ])
        pt <- as.numeric(colnames(s$scaled.smooth))
        # Figure out baseline expression & threshold for finding peaks
        min.val <- max(min(sv), 0)
        peak.val <- ((1 - min.val)/2) + min.val
        exp.val <- ((1 - min.val)/5) + min.val
        # Run-length encoding of above/below the peak-threshold
        peak.rle <- rle(sv >= peak.val)
```

```
        peak.rle <- data.frame(lengths = peak.rle$lengths, values = peak.rle$values)
        peak.rle$end <- cumsum(peak.rle$lengths)
        peak.rle$start <- head(c(0, peak.rle$end) + 1, -1)
        # Run-length encoding of above/below the expressed-threshold
        exp.rle <- rle(sv >= exp.val)
        exp.rle <- data.frame(lengths = exp.rle$lengths, values = exp.rle$values)
        exp.rle$end <- cumsum(exp.rle$lengths)
        exp.rle$start <- head(c(0, exp.rle$end) + 1, -1)
        # Take top-two longest peak RLE & select later one.  Find stretches that are
        # above peak value
        peak <- which(peak.rle$values)
        # Order by length and take 1 or 2 longest ones
        peak <- peak[order(peak.rle[peak, "lengths"], decreasing = T)][1:min(2, length(peak))]
        # Order by start and take latest one.
        peak <- peak[order(peak.rle[peak, "start"], decreasing = T)][1]
        # Identify the actual peak value within that stretch
        peak <- which.max(sv[peak.rle[peak, "start"]:peak.rle[peak, "end"]]) + peak.rle[peak,
            "start"] - 1
        # Identify the start and stop of the expressed stretch that contains the peak
        exp.start <- exp.rle[which(exp.rle$end >= peak & exp.rle$start <= peak),
            "start"]
        exp.end <- exp.rle[which(exp.rle$end >= peak & exp.rle$start <= peak), "end"]
        # Identify values of expression at start and stop
        smooth.start <- sv[exp.start]
        smooth.end <- sv[exp.end]
        # Convert to pseudotime?
        exp.start <- pt[exp.start]
        exp.end <- pt[exp.end]
        peak <- pt[peak]
        # Return a vector
        v <- c(exp.start, peak, exp.end, smooth.start, smooth.end)
        names(v) <- c("pt.start", "pt.peak", "pt.end", "exp.start", "exp.end")
        return(v)
    })))
    rownames(s$timing) <- genes

    # Decide on ordering of genes
    s$gene.order <- rownames(s$timing)[order(s$timing$pt.peak, s$timing$pt.start,
        s$timing$pt.end, s$timing$exp.end, decreasing = c(F, F, F, T), method = "radix")]

    return(s)
}
```

#### filter.heatmap.genes

Removes undesired (mitochondrial, ribosomal, tandem duplicated genes) from heatmaps for presentation purposes.

- genes: (Character vector) genes to check
- Returns `genes` with undesired genes removed.

```
filter.heatmap.genes <- function(genes) {
    mt.genes <- grep("^mt-", ignore.case = T, genes, value = T)
```

```
    many.genes <- grep("\\(1 of many\\)", ignore.case = T, genes, value = T)
    ribo.genes <- grep("^rpl|^rps", ignore.case = T, genes, value = T)
    cox.genes <- grep("^cox", ignore.case = T, genes, value = T)
    return(setdiff(genes, c(mt.genes, many.genes, ribo.genes, cox.genes)))
}
```

### Heatmaps of gene cascades

Using the genes that were determined as differentially expressed along the way to particular cell types, we generated expression cascades and plotted them as heatmaps.

#### Photoreceptors

##### Prepare cascade

```
## PHOTORECEPTORS: Seg 25 -> Cones (Seg 2) + Rods (Seg 12)

# Get markers from the two approaches:

# Lineage markers from above the combined clades
t25 <- threshold.clade.markers(combined.markers.best[["25"]], global.fc = 0.05)
# Cone markers from aucprTestAlongTree
m2 <- threshold.tree.markers(markers, "2", global.fc = 0.6)
# Rod markers from aucprTestAlongTree
m12 <- threshold.tree.markers(markers, "12", global.fc = 0.6)
pr.markers <- unique(c(rownames(t25), rownames(m2), rownames(m12)))

## Pseudotime for rods and cones is very different; for heatmaps, would like to
## normalize these, so that spline curves that consider both of them are not out
## of sync. Need to stretch pseudotime of cells in segment 12 / rods.

# Make a duplicate of the pseudotime measurement (pseudotime.212)
obj@pseudotime$pseudotime.212 <- obj@pseudotime$pseudotime
# Grab pseudotime of branchpoint
pt.start.212 <- as.numeric(obj@tree$segment.pseudotime.limits["2", "start"])
# Figure out lengths (and ratio) of the two branches in pseudotime
pt.end.212 <- as.numeric(obj@tree$segment.pseudotime.limits[c("2", "12"), "end"]) -
    pt.start.212
pt.ratio.212 <- pt.end.212[1]/pt.end.212[2]
# For cells in the shorter branch (12), subtract the starting pseudotime,
# multiply by the ratio of branch lengths, then add the starting pseudotime back
# in order to stretch the branch.
obj@pseudotime[cellsInCluster(obj, "segment", "12"), "pseudotime.212"] <- (obj@pseudotime[cellsInCluster(obj,
    "segment", "12"), "pseudotime.212"] - pt.start.212) * pt.ratio.212 + pt.start.212

# Calculate spline curves Using segments 29, 25, 2, and 12. Calculating a curve
# using only 29/25/2 for cone-specific genes, 29/25/12 for rod-specific genes,
# and 29/25/2+12 for genes that mark both. Should work now that pseudotimes are
# aligned.
```

```r
spline.2 <- geneSmoothFit(obj, pseudotime = "pseudotime.212", cells = cellsInCluster(obj,
    "segment", c("29", "25", "2")), genes = pr.markers, method = "spline", moving.window = 5,
    cells.per.window = 25, pseudotime.per.window = 0.005, spar = 0.5, verbose = F)
spline.12 <- geneSmoothFit(obj, pseudotime = "pseudotime.212", cells = cellsInCluster(obj,
    "segment", c("29", "25", "12")), genes = pr.markers, method = "spline", moving.window = 5,
    cells.per.window = 25, pseudotime.per.window = 0.005, spar = 0.5, verbose = F)
spline.212 <- geneSmoothFit(obj, pseudotime = "pseudotime.212", cells = cellsInCluster(obj,
    "segment", c("29", "25", "2", "12")), genes = pr.markers, method = "spline",
    moving.window = 5, cells.per.window = 25, pseudotime.per.window = 0.005, spar = 0.5,
    verbose = F)

# Want to plot a heatmap that shows expression in photoreceptor progenitors and
# then each branch (i.e. rods, cones) as separate columns. Going to crop each
# spline fit to the correct pseudotime range and then combine them into a single
# one that can be plotted as a three-column heatmap.

pt.2v12 <- obj@tree$segment.pseudotime.limits["2", "start"]  # pseudotime where the crop should happen
splines.pr <- list(cropSmoothFit(spline.212, pt.min = -Inf, pt.max = pt.2v12), cropSmoothFit(spline.2,
    pt.min = pt.2v12, pt.max = Inf), cropSmoothFit(spline.12, pt.min = pt.2v12, pt.max = Inf))
names(splines.pr) <- c("Photoreceptor Progenitors", "Rods", "Cones")
splines.pr.hm <- combineSmoothFit(splines.pr)  # Combine into a single one

# Calculate gene expression timing for ordering rows
spline.212 <- determine.timing(s = spline.212)
spline.2 <- determine.timing(s = spline.2)
spline.12 <- determine.timing(s = spline.12)

# Decide which markers are specific to one cell type or both
d2v12 <- divide.branches(obj, pr.markers, clust.1 = "2", clust.2 = "12", exp.fc = 0.4,
    exp.thresh = 0.2, exp.diff = 0.1)

# Generate gene ordering based on timing & specificity
order.212 <- filter.heatmap.genes(setdiff(spline.212$gene.order, c(d2v12$specific.1,
    d2v12$specific.2)))
order.2 <- filter.heatmap.genes(intersect(spline.2$gene.order, d2v12$specific.1))
order.12 <- filter.heatmap.genes(intersect(spline.12$gene.order, d2v12$specific.2))
gene.order <- c(order.212, order.2, order.12)

# Output gene table
table.save <- data.frame(gene = gene.order, marks = c(rep("both", length(order.212)),
    rep("cone", length(order.2)), rep("rod", length(order.12))), stringsAsFactors = F)
table.save$clade.AUCPR.ratio <- t25[table.save$gene, "AUCPR.ratio"]
table.save$clade.exp.fc <- t25[table.save$gene, "exp.fc"]
table.save$clade.exp.fc.global <- t25[table.save$gene, "exp.global.fc"]
table.save$cone.AUCPR.ratio.all <- m2[table.save$gene, "AUCPR.ratio.all"]
table.save$cone.AUCPR.ratio.maxBranch <- m2[table.save$gene, "AUCPR.ratio.maxBranch"]
table.save$cone.exp.fc.all <- m2[table.save$gene, "expfc.all"]
table.save$cone.exp.fc.best <- m2[table.save$gene, "expfc.maxBranch"]
table.save$rod.AUCPR.ratio.all <- m12[table.save$gene, "AUCPR.ratio.all"]
table.save$rod.AUCPR.ratio.maxBranch <- m12[table.save$gene, "AUCPR.ratio.maxBranch"]
table.save$rod.exp.fc.all <- m12[table.save$gene, "expfc.all"]
table.save$rod.exp.fc.best <- m12[table.save$gene, "expfc.maxBranch"]
write.csv(table.save, quote = F, file = paste0(base.path, "/heatmaps/retina-photoreceptor.csv"))
```

Generate heatmap: all genes

```
# Make sure any values <0 in the spline curves get set to 0 so that the heatmap
# scale doesn't get messed up.
splines.pr.hm$scaled.smooth[splines.pr.hm$scaled.smooth < 0] <- 0
# Determine where to place column separators (i.e. how many columns will each
# cell type occupy in the heatmap )
colsep <- cumsum(as.numeric(head(unlist(lapply(splines.pr, function(x) ncol(x$scaled.smooth))),
    -1)))
# Determine where to place row separators (i.e. how many common markers, and
# markers are specific to each cell type)
rowsep <- cumsum(c(length(order.212), length(order.2)))
# Open a PDF and generate the heatmap pdf(paste0(base.path,
# '/heatmaps/retina-photoreceptor.pdf'), width=6, height=10)
gplots::heatmap.2(x = as.matrix(splines.pr.hm$scaled.smooth[gene.order, ]), Rowv = F,
    Colv = F, dendrogram = "none", col = cols, trace = "none", density.info = "none",
    key = F, cexCol = 0.8, cexRow = 0.15, margins = c(8, 8), lwid = c(0.3, 4), lhei = c(0.3,
        4), labCol = NA, colsep = colsep, rowsep = rowsep, sepwidth = c(0.1, 0.2))
title(main = "Photoreceptors")
title(main = "Precursors", line = -41, adj = 0)
title(main = "Cones", line = -41, adj = 0.45)
title(main = "Rods", line = -41, adj = 0.76)
```

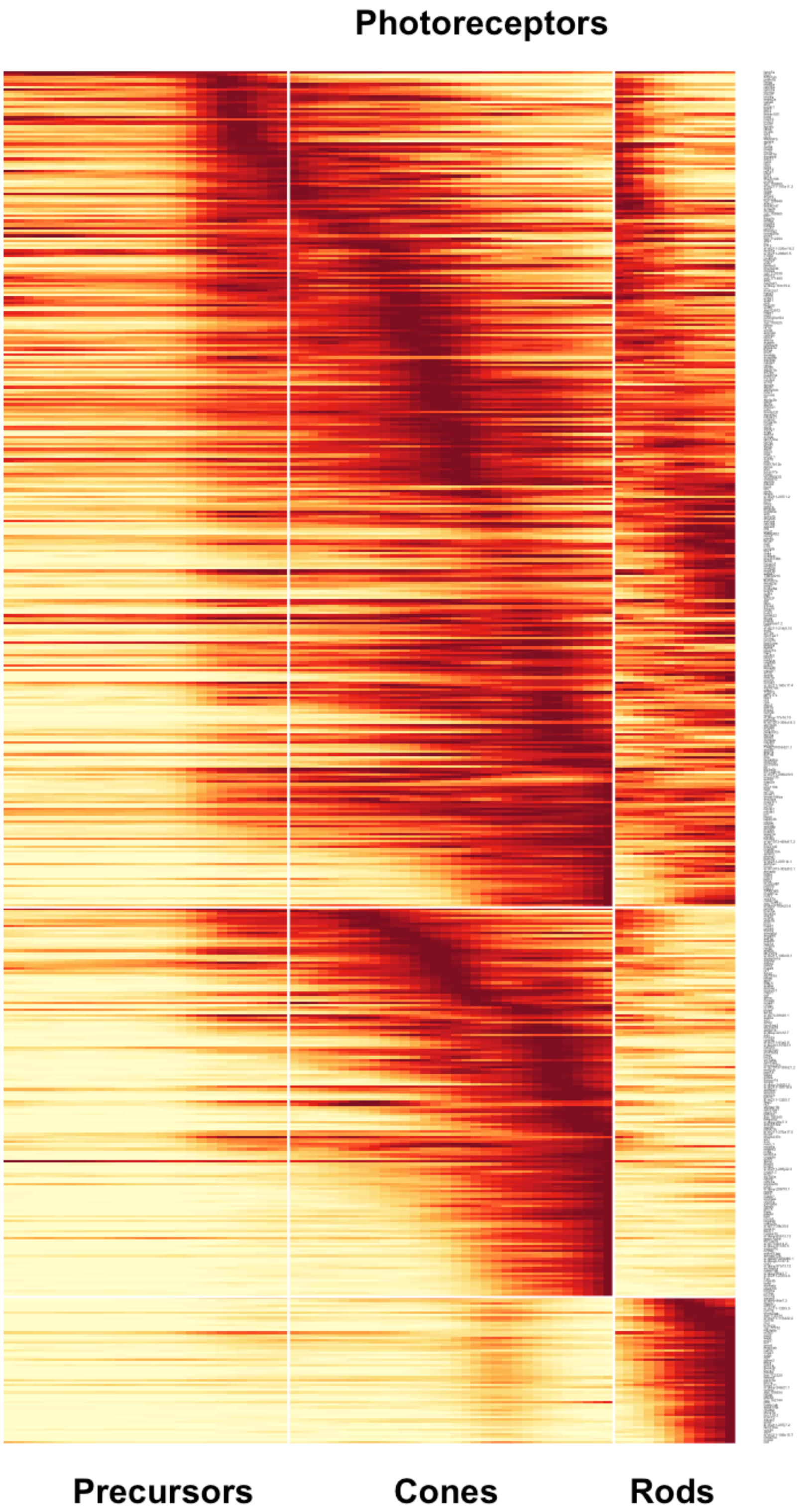

Photoreceptors
Precursors
Cones
Rods

```
# dev.off()
```

##### Generate heatmap: main figure

```
## Generate heatmap with only particular genes labeled for main figure
genes.to.plot <- c("isl2a", "prdm1a", "otx5", "crx", "six7", "nr2f1b", "nr2e3", "aplnrb",
    "aplnra", "apln")
rownames.to.plot <- gene.order
rtp <- rownames.to.plot %in% genes.to.plot
rownames.to.plot[!rtp] <- ""
rownames.to.plot[rtp] <- paste0("- ", rownames.to.plot[rtp])
# Open a PDF and generate the heatmap pdf(paste0(base.path,
# '/heatmaps/retina-photoreceptor-mainfig.pdf'), width=6, height=10)
gplots::heatmap.2(x = as.matrix(splines.pr.hm$scaled.smooth[gene.order, ]), Rowv = F,
    Colv = F, dendrogram = "none", col = cols, trace = "none", density.info = "none",
    key = F, cexCol = 0.8, cexRow = 1.8, margins = c(8, 8), lwid = c(0.3, 4), lhei = c(0.3,
        4), labCol = NA, colsep = colsep, rowsep = rowsep, sepwidth = c(0.1, 0.2),
    labRow = rownames.to.plot)
title(main = "Photoreceptors")
title(main = "Precursors", line = -41, adj = 0)
title(main = "Cones", line = -41, adj = 0.45)
title(main = "Rods", line = -41, adj = 0.76)
```

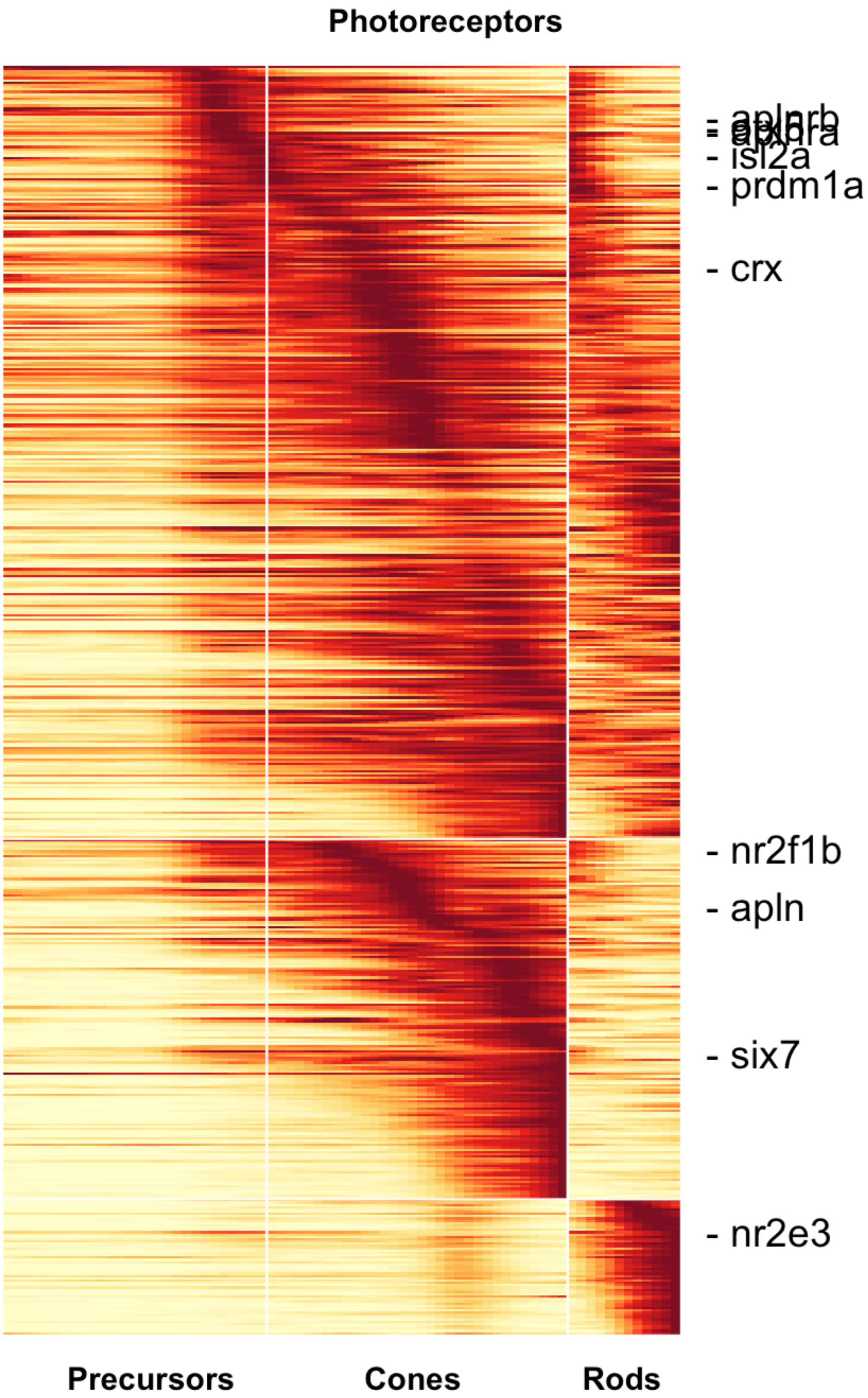
Photoreceptors
- aplnrb
- isl2a
- prdm1a
- crx
- nr2f1b
- apln
- six7
- nr2e3
Precursors
Cones
Rods

```
# dev.off()
```

#### Amacrine cells

##### Prepare cascade

```
## AMACRINE CELLS: Seg 19 -> Amarcine (Seg 4) + Starburst Amacrine (Seg 17)

# Get markers from the two approaches:

# Lineage markers from above the combined clades
t19 <- threshold.clade.markers(combined.markers.best[["19"]], global.fc = 0.05)
# Amacrine markers from aucprTestAlongTree
m4 <- threshold.tree.markers(markers, "4", global.fc = 0.6)
# Starburst amacrine markers from aucprTestAlongTree
m17 <- threshold.tree.markers(markers, "17", global.fc = 0.6)
am.markers <- unique(c(rownames(t19), rownames(m4), rownames(m17)))

## These have pretty equivalent pseudotimes, so don't need to worry about
## stretching them to match or anything.

# Calculate spline curves Using segments 29, 26, 19, and 4/17. Calculating a
# curve using only 29/26/19/4 for amacrine-specific genes, 29/26/19/4 for
# starburst-specific genes, and 29/26/19/4+17 for genes that mark both amacrine
# populations.
spline.4 <- geneSmoothFit(obj, pseudotime = "pseudotime", cells = cellsInCluster(obj,
    "segment", c("29", "26", "19", "4")), genes = am.markers, method = "spline",
    moving.window = 5, cells.per.window = 25, pseudotime.per.window = 0.005, spar = 0.5,
    verbose = F)
spline.17 <- geneSmoothFit(obj, pseudotime = "pseudotime", cells = cellsInCluster(obj,
    "segment", c("29", "26", "19", "17")), genes = am.markers, method = "spline",
    moving.window = 5, cells.per.window = 25, pseudotime.per.window = 0.005, spar = 0.5,
    verbose = F)
spline.417 <- geneSmoothFit(obj, pseudotime = "pseudotime", cells = cellsInCluster(obj,
    "segment", c("29", "26", "19", "4", "17")), genes = am.markers, method = "spline",
    moving.window = 5, cells.per.window = 25, pseudotime.per.window = 0.005, spar = 0.5,
    verbose = F)

# Want to plot a heatmap that shows expression in amacrine progenitors and then
# each branch (i.e. amacrine_gaba, starburst_amacrine) as separate columns. Going
# to crop each spline fit to the correct pseudotime range and then combine them
# into a single one that can be plotted as a three-column heatmap.

pt.4v17 <- obj@tree$segment.pseudotime.limits["4", "start"]  # pseudotime where the crop should happen
splines.am <- list(cropSmoothFit(spline.417, pt.min = -Inf, pt.max = pt.4v17), cropSmoothFit(spline.4,
    pt.min = pt.4v17, pt.max = Inf), cropSmoothFit(spline.17, pt.min = pt.4v17, pt.max = Inf))
names(splines.am) <- c("Amacrine Precursors", "Amacrine", "Starburst Amacrine")
splines.am.hm <- combineSmoothFit(splines.am)  # Combine into a single one

# Calculate gene expression timing for ordering rows
spline.417 <- determine.timing(s = spline.417)
```

```r
spline.4 <- determine.timing(s = spline.4)
spline.17 <- determine.timing(s = spline.17)

# Decide which markers are specific to one cell type or both
d4v17 <- divide.branches(obj, am.markers, clust.1 = "4", clust.2 = "17", exp.fc = 0.4,
    exp.thresh = 0.2, exp.diff = 0.1)

# Generate gene ordering based on timing & specificity
order.417 <- filter.heatmap.genes(setdiff(spline.417$gene.order, c(d4v17$specific.1,
    d4v17$specific.2)))
order.4 <- filter.heatmap.genes(intersect(spline.4$gene.order, d4v17$specific.1))
order.17 <- filter.heatmap.genes(intersect(spline.17$gene.order, d4v17$specific.2))
gene.order <- c(order.417, order.4, order.17)

# Output gene table
table.save <- data.frame(gene = gene.order, marks = c(rep("both", length(order.417)),
    rep("amacrine", length(order.4)), rep("starburst", length(order.17))), stringsAsFactors = F)
table.save$clade.AUCPR.ratio <- t19[table.save$gene, "AUCPR.ratio"]
table.save$clade.exp.fc <- t19[table.save$gene, "exp.fc"]
table.save$clade.exp.fc.global <- t19[table.save$gene, "exp.global.fc"]
table.save$am.AUCPR.ratio.all <- m4[table.save$gene, "AUCPR.ratio.all"]
table.save$am.AUCPR.ratio.maxBranch <- m4[table.save$gene, "AUCPR.ratio.maxBranch"]
table.save$am.exp.fc.all <- m4[table.save$gene, "expfc.all"]
table.save$am.exp.fc.best <- m4[table.save$gene, "expfc.maxBranch"]
table.save$star.AUCPR.ratio.all <- m17[table.save$gene, "AUCPR.ratio.all"]
table.save$star.AUCPR.ratio.maxBranch <- m17[table.save$gene, "AUCPR.ratio.maxBranch"]
table.save$star.exp.fc.all <- m17[table.save$gene, "expfc.all"]
table.save$star.exp.fc.best <- m17[table.save$gene, "expfc.maxBranch"]
write.csv(table.save, quote = F, file = paste0(base.path, "/heatmaps/retina-amacrine.csv"))
```

##### Generate heatmap: all genes

```r
# Make sure any values <0 in the spline curves get set to 0 so that the heatmap
# scale doesn't get messed up.
splines.am.hm$scaled.smooth[splines.am.hm$scaled.smooth < 0] <- 0
# Determine where to place column separators (i.e. how many columns will each
# cell type occupy in the heatmap )
colsep <- cumsum(as.numeric(head(unlist(lapply(splines.am, function(x) ncol(x$scaled.smooth))),
    -1)))
# Determine where to place row separators (i.e. how many common markers, and
# markers are specific to each cell type)
rowsep <- cumsum(c(length(order.417), length(order.4)))
# Open a PDF and generate the heatmap pdf(paste0(base.path,
# '/heatmaps/retina-amacrine.pdf'), width=6, height=10)
gplots::heatmap.2(x = as.matrix(splines.am.hm$scaled.smooth[gene.order, ]), Rowv = F,
    Colv = F, dendrogram = "none", col = cols, trace = "none", density.info = "none",
    key = F, cexCol = 0.8, cexRow = 0.15, margins = c(8, 8), lwid = c(0.3, 4), lhei = c(0.3,
        4), labCol = NA, colsep = colsep, rowsep = rowsep, sepwidth = c(0.05, 0.2))
title(main = "Amacrine Cells")
title(main = "Precursors", line = -41, adj = 0.05)
title(main = "Amacrine", line = -41, adj = 0.475)
```

```r
title(main = "Starburst", line = -41, adj = 0.75)
```

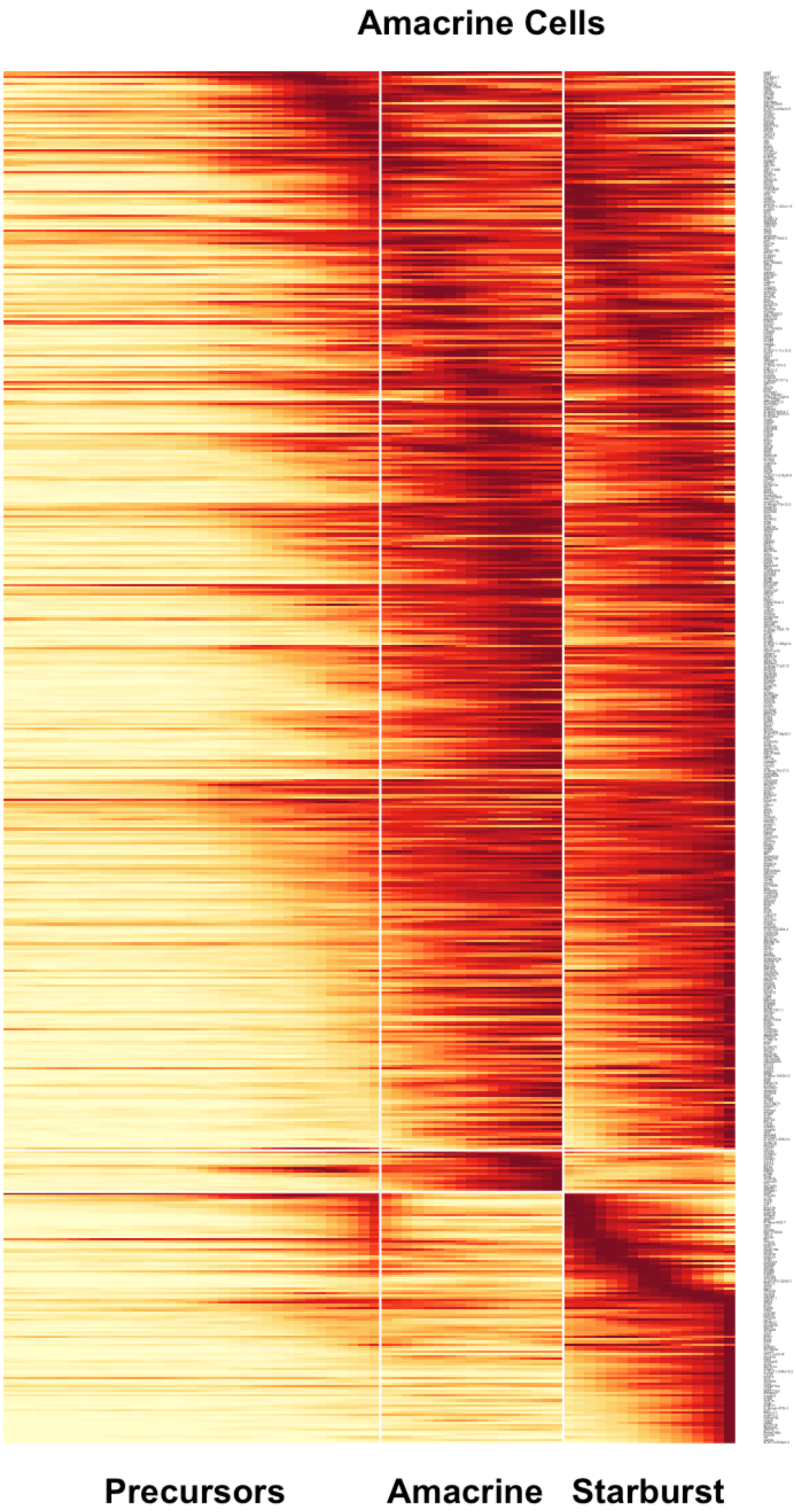
Amacrine Cells
Precursors
Amacrine
Starburst

```
# dev.off()
```

#### Retinal ganglion cells

##### Prepare cascade

```
## RGCs: Seg 8

# Get markers from the two approaches:

# Lineage markers from above the combined clades
t8 <- threshold.clade.markers(combined.markers.best[["8"]], global.fc = 0.05)
# RGC markers from aucprTestAlongTree
m8 <- threshold.tree.markers(markers, "8", global.fc = 0.6)
rgc.markers <- unique(c(rownames(t8), rownames(m8)))

# Calculate spline curves Using segments 29, 26, and 8.
spline.8 <- geneSmoothFit(obj, pseudotime = "pseudotime", cells = cellsInCluster(obj,
    "segment", c("29", "26", "8")), genes = rgc.markers, method = "spline", moving.window = 5,
    cells.per.window = 25, pseudotime.per.window = 0.005, spar = 0.5, verbose = F)

# Calculate gene expression timing for ordering rows
spline.8 <- determine.timing(s = spline.8)
order.8 <- filter.heatmap.genes(spline.8$gene.order)

# Output gene table
table.save <- data.frame(gene = order.8, stringsAsFactors = F)
table.save$clade.AUCPR.ratio <- t8[table.save$gene, "AUCPR.ratio"]
table.save$clade.exp.fc <- t8[table.save$gene, "exp.fc"]
table.save$clade.exp.fc.global <- t8[table.save$gene, "exp.global.fc"]
table.save$rgc.AUCPR.ratio.all <- m8[table.save$gene, "AUCPR.ratio.all"]
table.save$rgc.AUCPR.ratio.maxBranch <- m8[table.save$gene, "AUCPR.ratio.maxBranch"]
table.save$rgc.exp.fc.all <- m8[table.save$gene, "expfc.all"]
table.save$rgc.exp.fc.best <- m8[table.save$gene, "expfc.maxBranch"]
write.csv(table.save, quote = F, file = paste0(base.path, "/heatmaps/retina-rgc.csv"))
```

##### Generate heatmap: all genes

```
# Make sure any values <0 in the spline curves get set to 0 so that the heatmap
# scale doesn't get messed up.
spline.8$scaled.smooth[spline.8$scaled.smooth < 0] <- 0
# Open a PDF and generate the heatmap pdf(paste0(base.path,
# '/heatmaps/retina-rgc.pdf'), width=6, height=10)
gplots::heatmap.2(x = as.matrix(spline.8$scaled.smooth[order.8, ]), Rowv = F, Colv = F,
    dendrogram = "none", col = cols, trace = "none", density.info = "none", key = F,
    cexCol = 0.8, cexRow = 0.15, margins = c(8, 8), lwid = c(0.3, 4), lhei = c(0.3,
        4), labCol = NA)
title(main = "Retinal Ganglion Cells")
```

#### Retinal Ganglion Cells

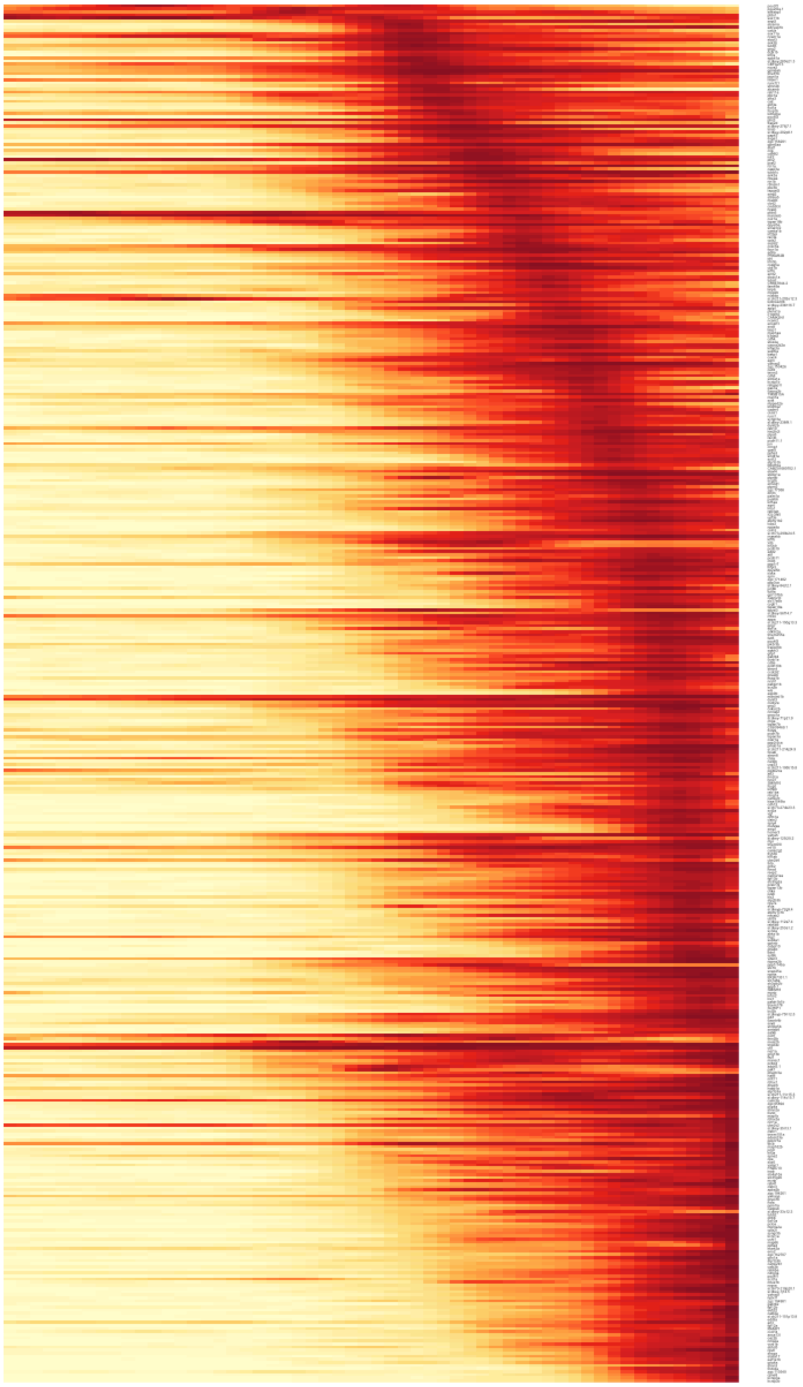

```r
# dev.off()
```

#### Generate heatmap: main figure

```r
genes.to.plot <- c("sox11a", "sox11b", "sox6", "irx4a", "pou4f2", "pou4f1", "rbpms2b",
    "rbpms2a")
rownames.to.plot <- order.8
rtp <- rownames.to.plot %in% genes.to.plot
rownames.to.plot[!rtp] <- ""
rownames.to.plot[rtp] <- paste0("- ", rownames.to.plot[rtp])
# Open a PDF and generate the heatmap pdf(paste0(base.path,
# '/heatmaps/retina-rgc-mainfig.pdf'), width=6, height=10)
gplots::heatmap.2(x = as.matrix(spline.8$scaled.smooth[order.8, ]), Rowv = F, Colv = F,
    dendrogram = "none", col = cols, trace = "none", density.info = "none", key = F,
    cexCol = 0.8, cexRow = 1.8, margins = c(8, 10), lwid = c(0.3, 4), lhei = c(0.3,
        4), labCol = NA, labRow = rownames.to.plot)
title(main = "Retinal Ganglion Cells")
```

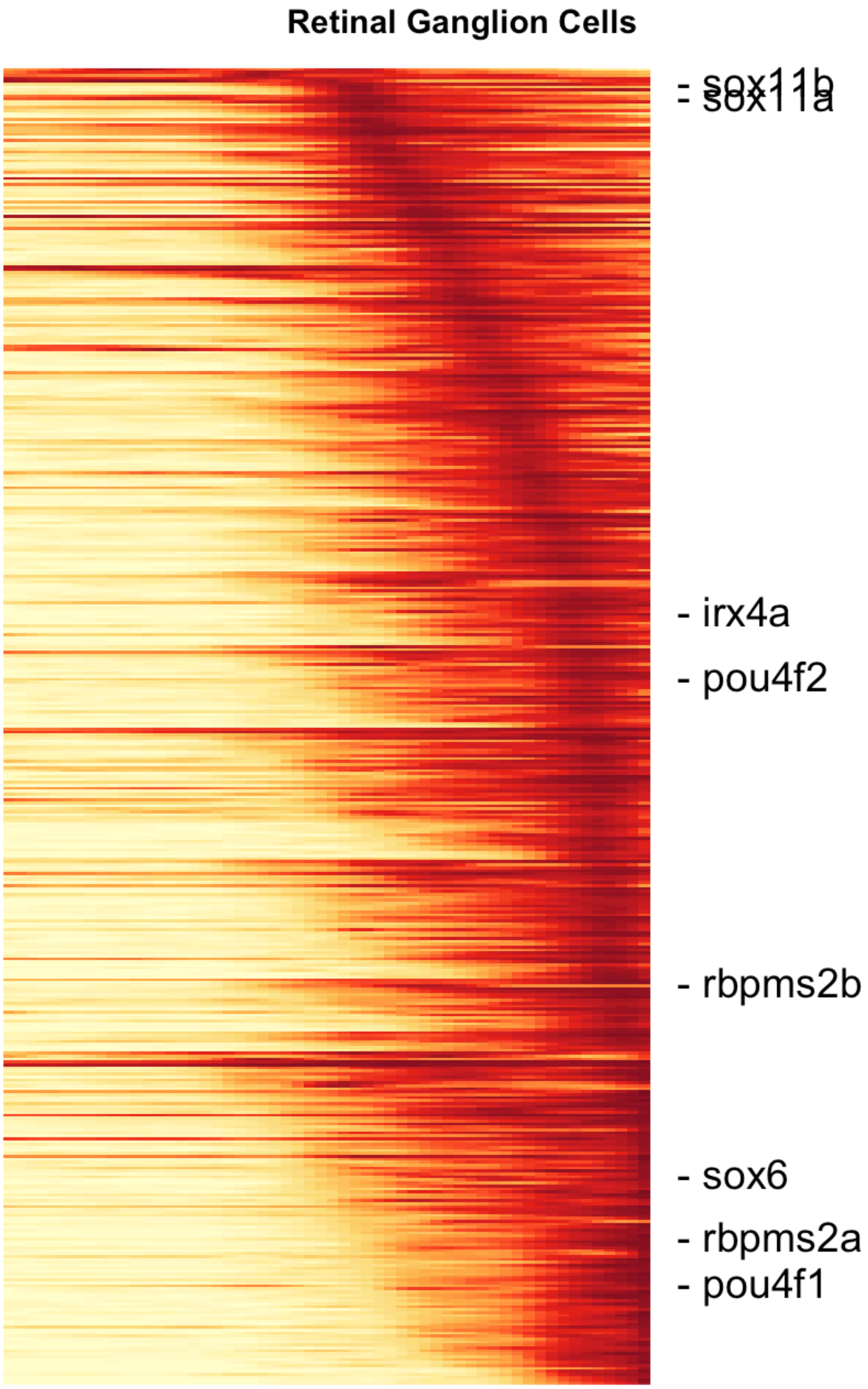
Retinal Ganglion Cells
- sox11b
- sox11a
- irx4a
- pou4f2
- rbpms2b
- sox6
- rbpms2a
- pou4f1

```
# dev.off()
```

#### Horizontal Cells

##### Prepare cascade

```
## Horizontal Cells: Seg 15

# Get markers from the two approaches:

# Lineage markers from above the combined clades
t15 <- threshold.clade.markers(combined.markers.best[["15"]], global.fc = 0.05)
# Horizontal Cell markers from aucprTestAlongTree
m15 <- threshold.tree.markers(markers, "15", global.fc = 0.6)
horiz.markers <- unique(c(rownames(t15), rownames(m15)))

# Calculate spline curves Using segments 29 and 15.
spline.15 <- geneSmoothFit(obj, pseudotime = "pseudotime", cells = cellsInCluster(obj,
    "segment", c("29", "15")), genes = horiz.markers, method = "spline", moving.window = 5,
    cells.per.window = 25, pseudotime.per.window = 0.005, spar = 0.5, verbose = F)

# Calculate gene expression timing for ordering rows
spline.15 <- determine.timing(s = spline.15)
order.15 <- filter.heatmap.genes(spline.15$gene.order)

# Output gene table
table.save <- data.frame(gene = order.15, stringsAsFactors = F)
table.save$clade.AUCPR.ratio <- t15[table.save$gene, "AUCPR.ratio"]
table.save$clade.exp.fc <- t15[table.save$gene, "exp.fc"]
table.save$clade.exp.fc.global <- t15[table.save$gene, "exp.global.fc"]
table.save$horiz.AUCPR.ratio.all <- m15[table.save$gene, "AUCPR.ratio.all"]
table.save$horiz.AUCPR.ratio.maxBranch <- m15[table.save$gene, "AUCPR.ratio.maxBranch"]
table.save$horiz.exp.fc.all <- m15[table.save$gene, "expfc.all"]
table.save$horiz.exp.fc.best <- m15[table.save$gene, "expfc.maxBranch"]
write.csv(table.save, quote = F, file = paste0(base.path, "/heatmaps/retina-horiz.csv"))
```

##### Generate heatmap: all genes

```
# Make sure any values <0 in the spline curves get set to 0 so that the heatmap
# scale doesn't get messed up.
splines.am.hm$scaled.smooth[splines.am.hm$scaled.smooth < 0] <- 0
# Determine where to place column separators (i.e. how many columns will each
# cell type occupy in the heatmap )
colsep <- cumsum(as.numeric(head(unlist(lapply(splines.am, function(x) ncol(x$scaled.smooth))),
    -1)))
# Determine where to place row separators (i.e. how many common markers, and
# markers are specific to each cell type)
rowsep <- cumsum(c(length(order.417), length(order.4)))
# Open a PDF and generate the heatmap pdf(paste0(base.path,
```

```r
# '/heatmaps/retina-amacrine.pdf'), width=6, height=10)
gplots::heatmap.2(x = as.matrix(splines.am.hm$scaled.smooth[gene.order, ]), Rowv = F,
    Colv = F, dendrogram = "none", col = cols, trace = "none", density.info = "none",
    key = F, cexCol = 0.8, cexRow = 0.15, margins = c(8, 8), lwid = c(0.3, 4), lhei = c(0.3,
        4), labCol = NA, colsep = colsep, rowsep = rowsep, sepwidth = c(0.05, 0.2))
title(main = "Amacrine Cells")
title(main = "Precursors", line = -41, adj = 0.05)
title(main = "Amacrine", line = -41, adj = 0.475)
title(main = "Starburst", line = -41, adj = 0.75)
```

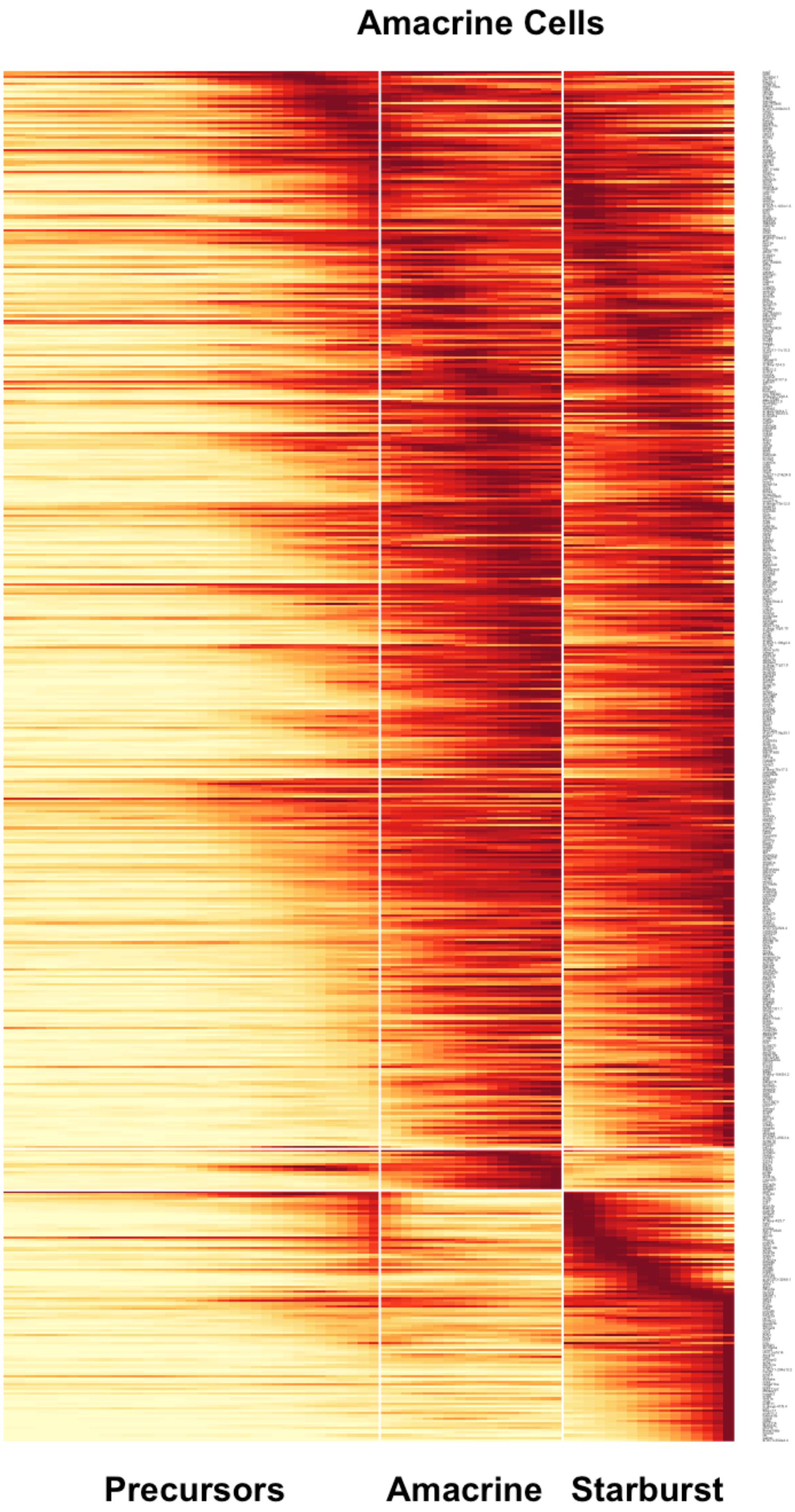
Amacrine Cells
Precursors
Amacrine
Starburst

```
# dev.off()
```

#### Muller Glia

##### Prepare cascade

```
## Muller Glia: Seg 6

# Get markers from the two approaches
m6 <- threshold.tree.markers(markers, "6", global.fc = 0.6)  # Muller Glia markers from aucprTestAlongTree
muller.markers <- rownames(m6)

# Calculate spline curves Using segments 29 and 15.
spline.6 <- geneSmoothFit(obj, pseudotime = "pseudotime", cells = cellsInCluster(obj,
    "segment", c("30", "6")), genes = muller.markers, method = "spline", moving.window = 5,
    cells.per.window = 25, pseudotime.per.window = 0.005, spar = 0.5, verbose = F)

# Calculate gene expression timing for ordering rows
spline.6 <- determine.timing(s = spline.6)
order.6 <- filter.heatmap.genes(spline.6$gene.order)

# Output gene table
table.save <- data.frame(gene = order.6, stringsAsFactors = F)
table.save$muller.AUCPR.ratio.all <- m6[table.save$gene, "AUCPR.ratio.all"]
table.save$muller.AUCPR.ratio.maxBranch <- m6[table.save$gene, "AUCPR.ratio.maxBranch"]
table.save$muller.exp.fc.all <- m6[table.save$gene, "expfc.all"]
table.save$muller.exp.fc.best <- m6[table.save$gene, "expfc.maxBranch"]
write.csv(table.save, quote = F, file = paste0(base.path, "/heatmaps/retina-muller.csv"))
```

##### Generate heatmap: all genes

```
# Make sure any values <0 in the spline curves get set to 0 so that the heatmap
# scale doesn't get messed up.
spline.6$scaled.smooth[spline.6$scaled.smooth < 0] <- 0
# Open a PDF and generate the heatmap pdf(paste0(base.path,
# '/heatmaps/retina-muller.pdf'), width=6, height=10)
gplots::heatmap.2(x = as.matrix(spline.6$scaled.smooth[order.6, ]), Rowv = F, Colv = F,
    dendrogram = "none", col = cols, trace = "none", density.info = "none", key = F,
    cexCol = 0.8, cexRow = 0.15, margins = c(8, 8), lwid = c(0.3, 4), lhei = c(0.3,
        4), labCol = NA)
title(main = "Muller Glia")
```

#### Muller Glia

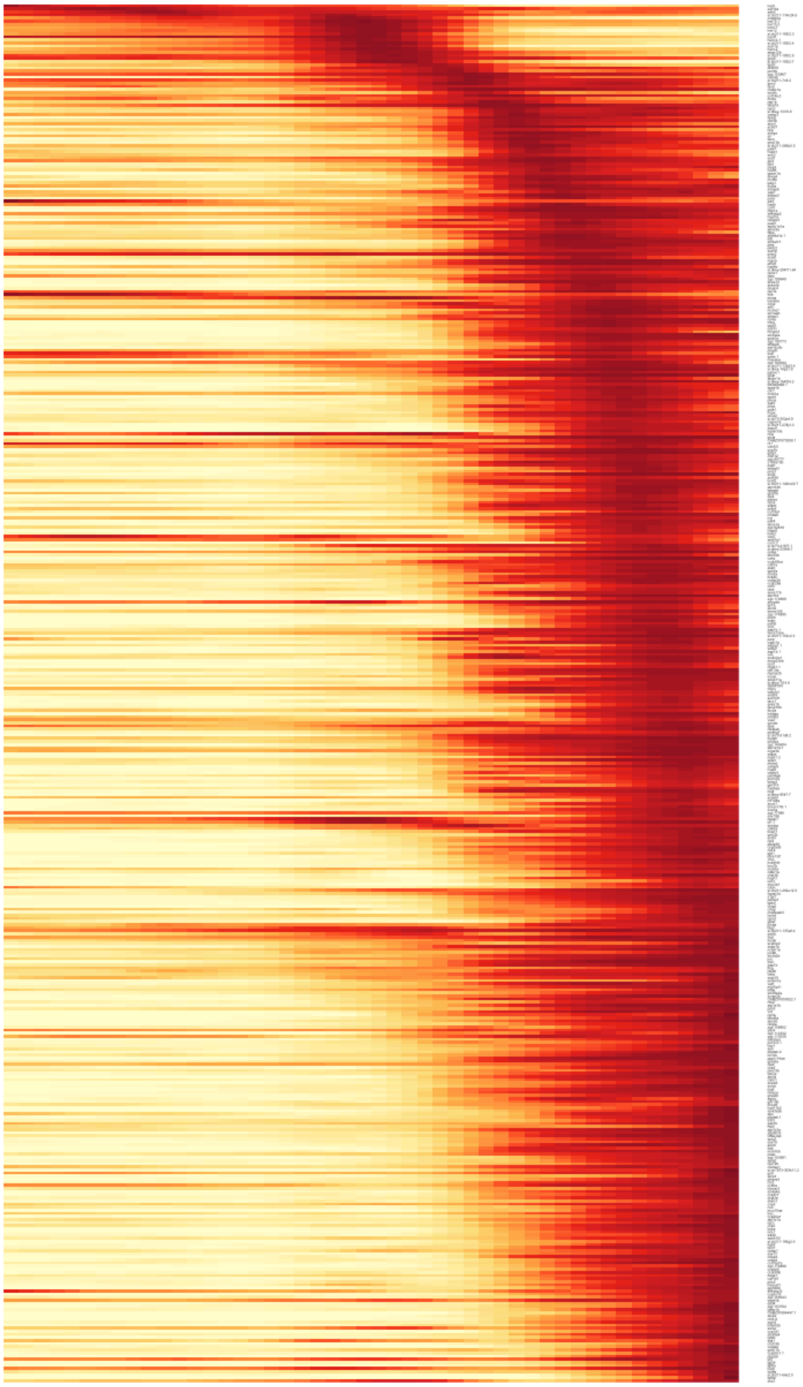

```
# dev.off()
```

#### Retinal Pigmented Epithelium

##### Prepare cascade

```
## RPE: Seg 11

# Get markers from the two approaches
m11 <- threshold.tree.markers(markers, "11", global.fc = 0.6)  # RPE markers from aucprTestAlongTree
rpe.markers <- rownames(m11)

# Just want to plot part of cells from upstream segment 31, which is very long.
# Going to use cells from segment 11 and from segment 31 with pseudotime > 0.23
cells.rpe <- unique(c(whichCells(obj, "pseudotime", c(0.23, 0.30308134)), cellsInCluster(obj,
    "segment", "11")))

# Calculate spline curves
spline.11 <- geneSmoothFit(obj, pseudotime = "pseudotime", cells = cells.rpe, genes = rpe.markers,
    method = "spline", moving.window = 5, cells.per.window = 25, pseudotime.per.window = 0.005,
    spar = 0.5, verbose = F)

# Calculate gene expression timing for ordering rows
spline.11 <- determine.timing(s = spline.11)
order.11 <- filter.heatmap.genes(spline.11$gene.order)

# Output gene table
table.save <- data.frame(gene = order.11, stringsAsFactors = F)
table.save$rpe.AUCPR.ratio.all <- m11[table.save$gene, "AUCPR.ratio.all"]
table.save$rpe.AUCPR.ratio.maxBranch <- m11[table.save$gene, "AUCPR.ratio.maxBranch"]
table.save$rpe.exp.fc.all <- m11[table.save$gene, "expfc.all"]
table.save$rpe.exp.fc.best <- m11[table.save$gene, "expfc.maxBranch"]
write.csv(table.save, quote = F, file = paste0(base.path, "/heatmaps/retina-rpe.csv"))
```

##### Generate heatmap: all genes

```
# Make sure any values <0 in the spline curves get set to 0 so that the heatmap
# scale doesn't get messed up.
spline.11$scaled.smooth[spline.11$scaled.smooth < 0] <- 0
# Open a PDF and generate the heatmap pdf(paste0(base.path,
# '/heatmaps/retina-rpe.pdf'), width=6, height=16)
gplots::heatmap.2(x = as.matrix(spline.11$scaled.smooth[order.11, ]), Rowv = F, Colv = F,
    dendrogram = "none", col = cols, trace = "none", density.info = "none", key = F,
    cexCol = 0.8, cexRow = 0.15, margins = c(8, 8), lwid = c(0.3, 4), lhei = c(0.3,
        4), labCol = NA)
title(main = "Retinal Pigmented Epithelium")
```

**Retinal Pigmented Epithelium**

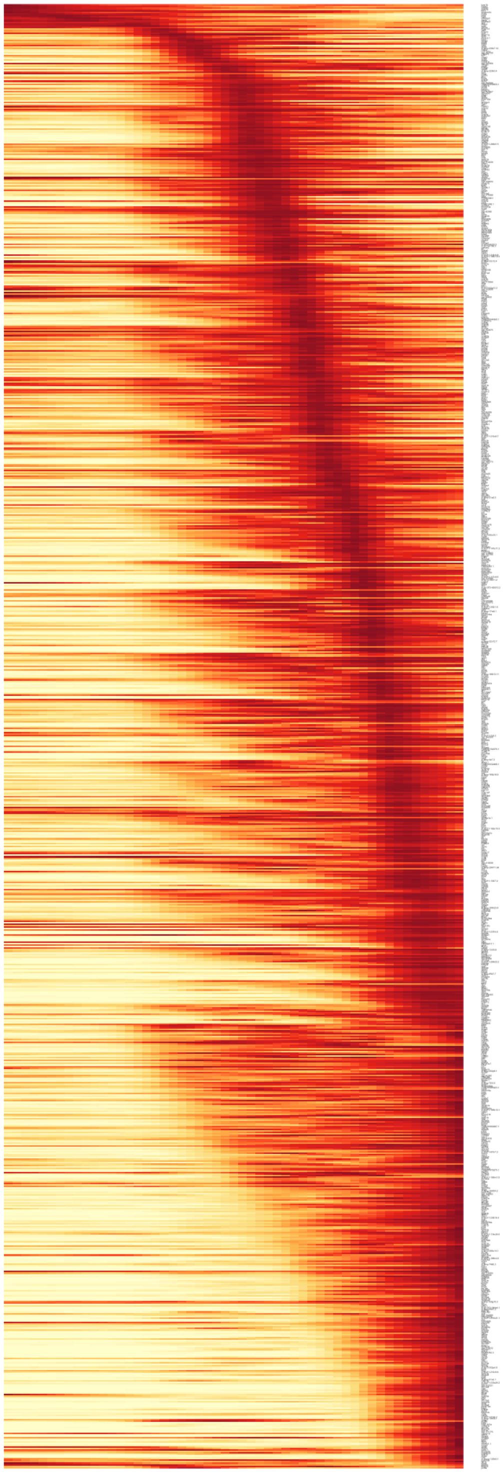

```
# dev.off()
```

#### Continuous differentiation

Retinal cell types were often found with similar molecular states across many stages of development. This reflects that pseudotime accurately represents the asynchrony introduced by continuous differentation.

##### RGC cells

```
gridExtra::grid.arrange(grobs = list(plotTreeHighlight(obj, label.name = "clus.orig",
    label.value = "7-36h_27", highlight.size = 1, title = "36 hpf Cluster 27: RGCs",
    label.x = F), plotTreeHighlight(obj, label.name = "clus.orig", label.value = "12-15d_38",
    highlight.size = 1, title = "15 dpf Cluster 38: RGCs", label.x = F)), ncol = 2)
```

```
## Warning: Removed 5 rows containing missing values (geom_point).
```

```
## Warning: Removed 17 rows containing missing values (geom_point).
```

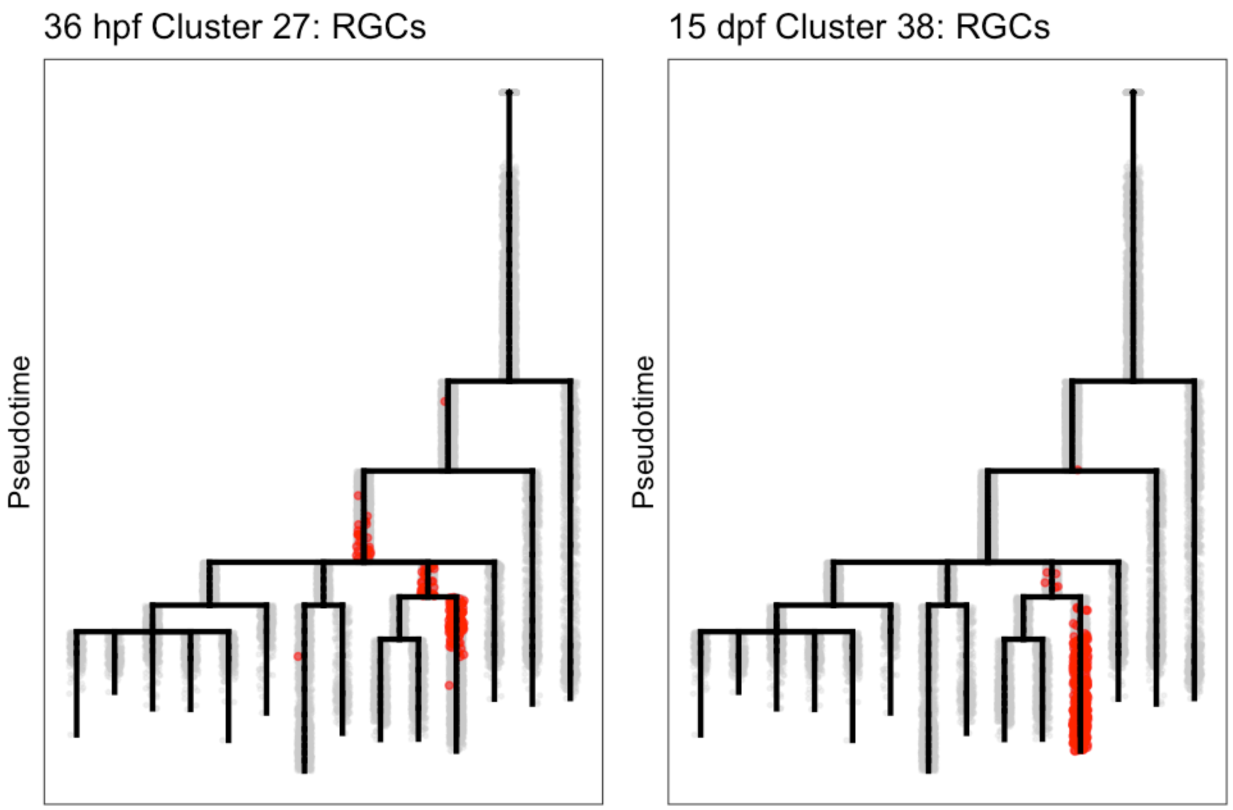

##### Progenitor cells

```r
gridExtra::grid.arrange(grobs = list(plotTreeHighlight(obj, label.name = "clus.orig",
    label.value = "6-24h_22", highlight.size = 1, title = "24 hpf Cluster 22: Progenitor Cells",
    label.x = F), plotTreeHighlight(obj, label.name = "clus.orig", label.value = "7-36h_32",
    highlight.size = 1, title = "36 hpf Cluster 32: Progenitor Cells", label.x = F),
    plotTreeHighlight(obj, label.name = "clus.orig", label.value = "12-15d_39", highlight.size = 1,
        title = "15 dpf Cluster 39: Progenitor Cells", label.x = F)), ncol = 2)
```

```
## Warning: Removed 9 rows containing missing values (geom_point).
```

```
## Warning: Removed 4 rows containing missing values (geom_point).
```

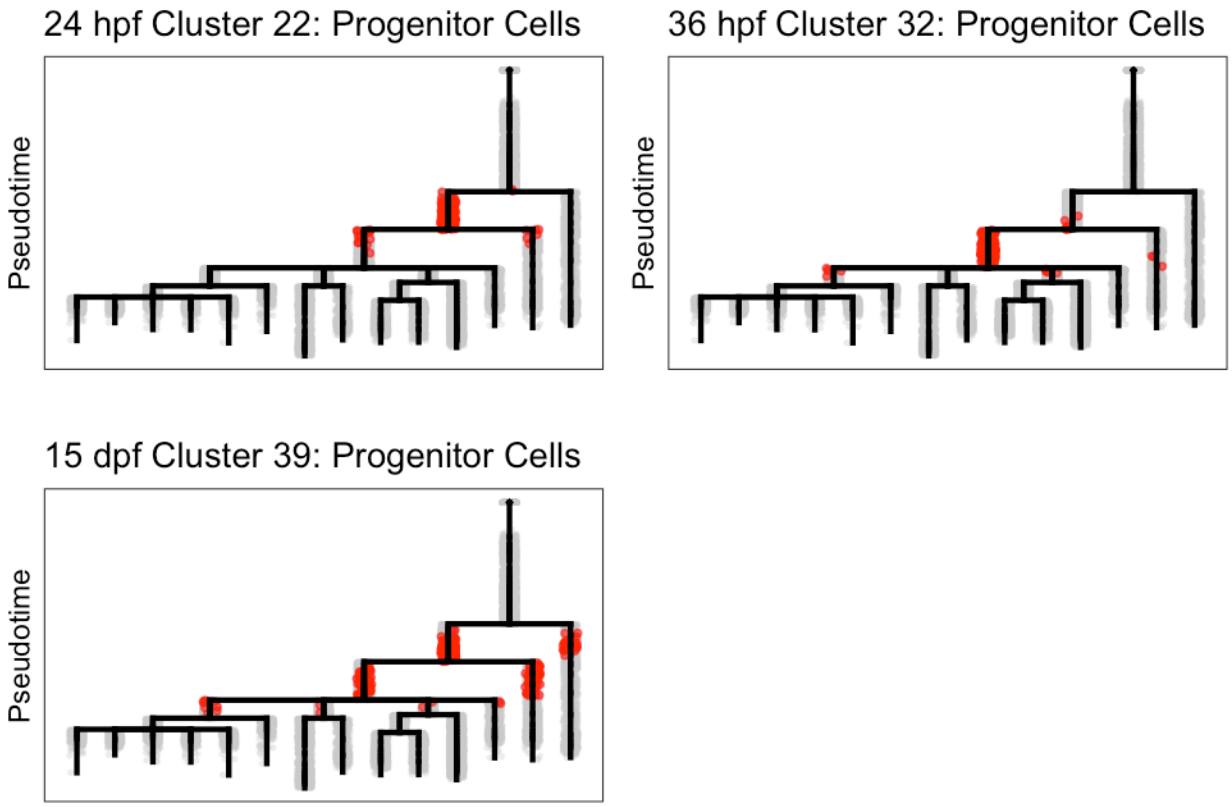

#### Progenitors over time

Retinal progenitors with similar transcriptional states are found across many different time points. We wanted to know whether there were significant transcriptional changes within those progenitors between early stages and late stages.

##### Identify populations

First we grabbed early (24 / 36 hpf) and late (15 dpf) progenitors from two sections of the tree.

```
# Progenitors / 24-36 hpf / Segment 30
prog.early.s30 <- intersect(cellsInCluster(obj, "stage", c("06-24h", "07-36h")),
    cellsInCluster(obj, "segment", "30"))
obj <- groupFromCells(obj, group.id = "prog.early.s30", cells = prog.early.s30)
plotTreeHighlight(obj, "prog.early.s30", "TRUE", highlight.size = 1, highlight.alpha = 0.5,
    title = "Progenitors / 24-36 hpf / Segment 30")
```

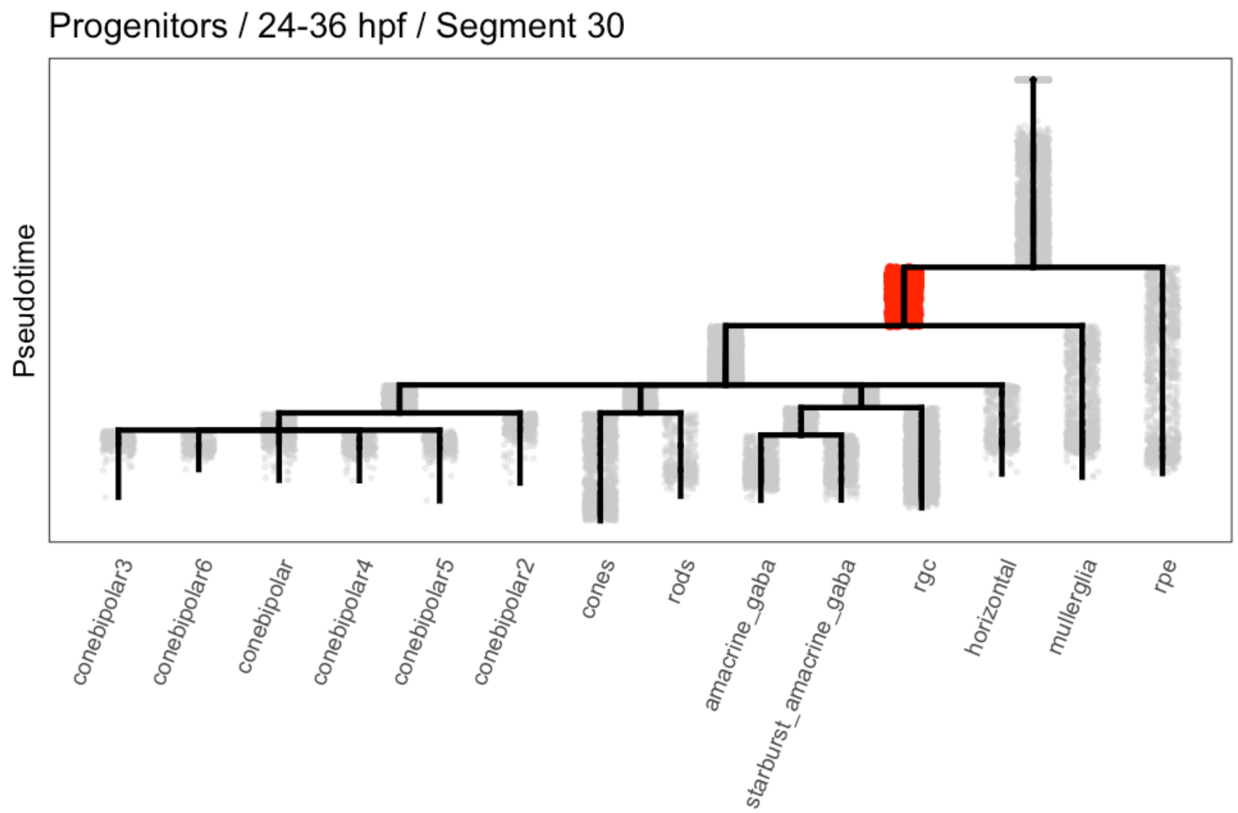

```
# Progenitors / 24-36 hpf / Segment 29
prog.early.s29 <- intersect(cellsInCluster(obj, "stage", c("06-24h", "07-36h")),
    cellsInCluster(obj, "segment", "29"))
obj <- groupFromCells(obj, group.id = "prog.early.s29", cells = prog.early.s29)
plotTreeHighlight(obj, "prog.early.s29", "TRUE", highlight.size = 1, highlight.alpha = 0.5,
    title = "Progenitors / 24-36 hpf / Segment 29")
```

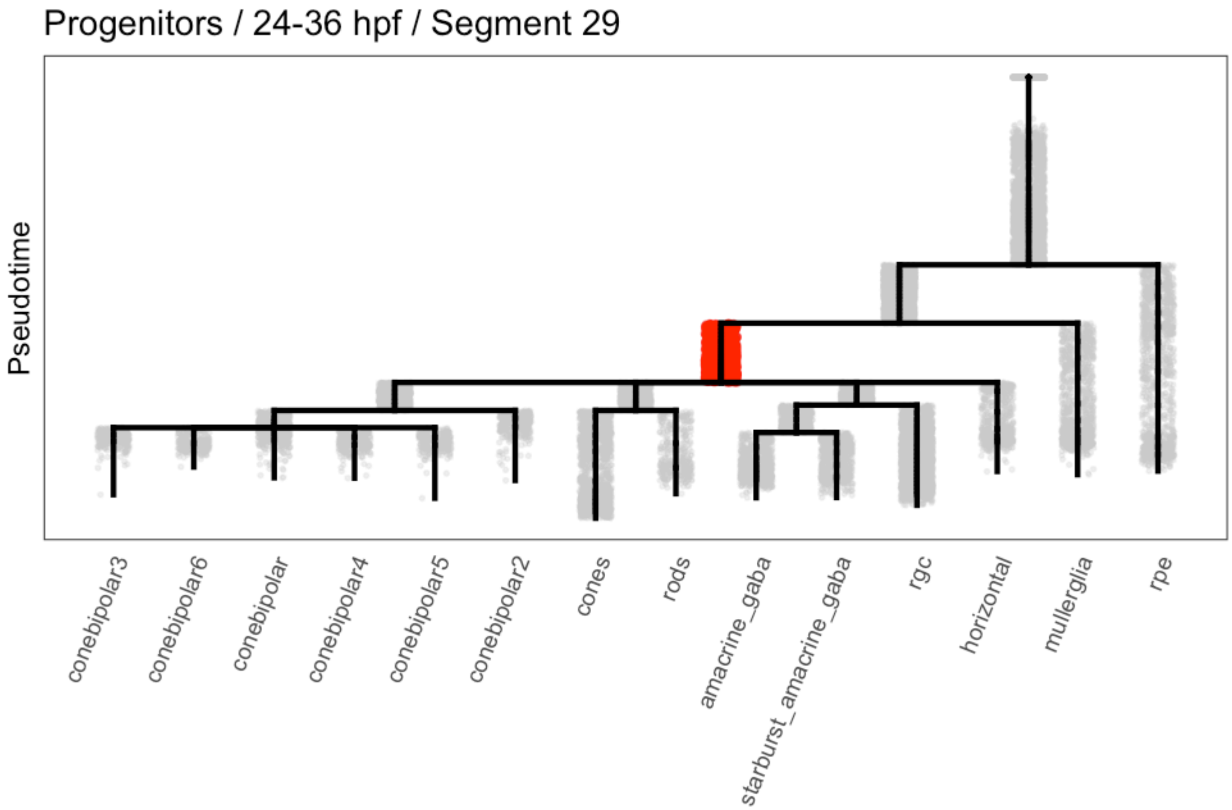

```r
# Progenitors / 15 dpf / Cluster 39 / Segment 30
prog.15d.c39.s30 <- intersect(cellsInCluster(obj, "clus.orig", "12-15d_39"), cellsInCluster(obj,
    "segment", "30"))
obj <- groupFromCells(obj, group.id = "prog.15d.c39.s30", cells = prog.15d.c39.s30)
plotTreeHighlight(obj, "prog.15d.c39.s30", "TRUE", highlight.size = 1, highlight.alpha = 0.5,
    title = "Progenitors / 15 dpf / Cluster 39 / Segment 30")
```

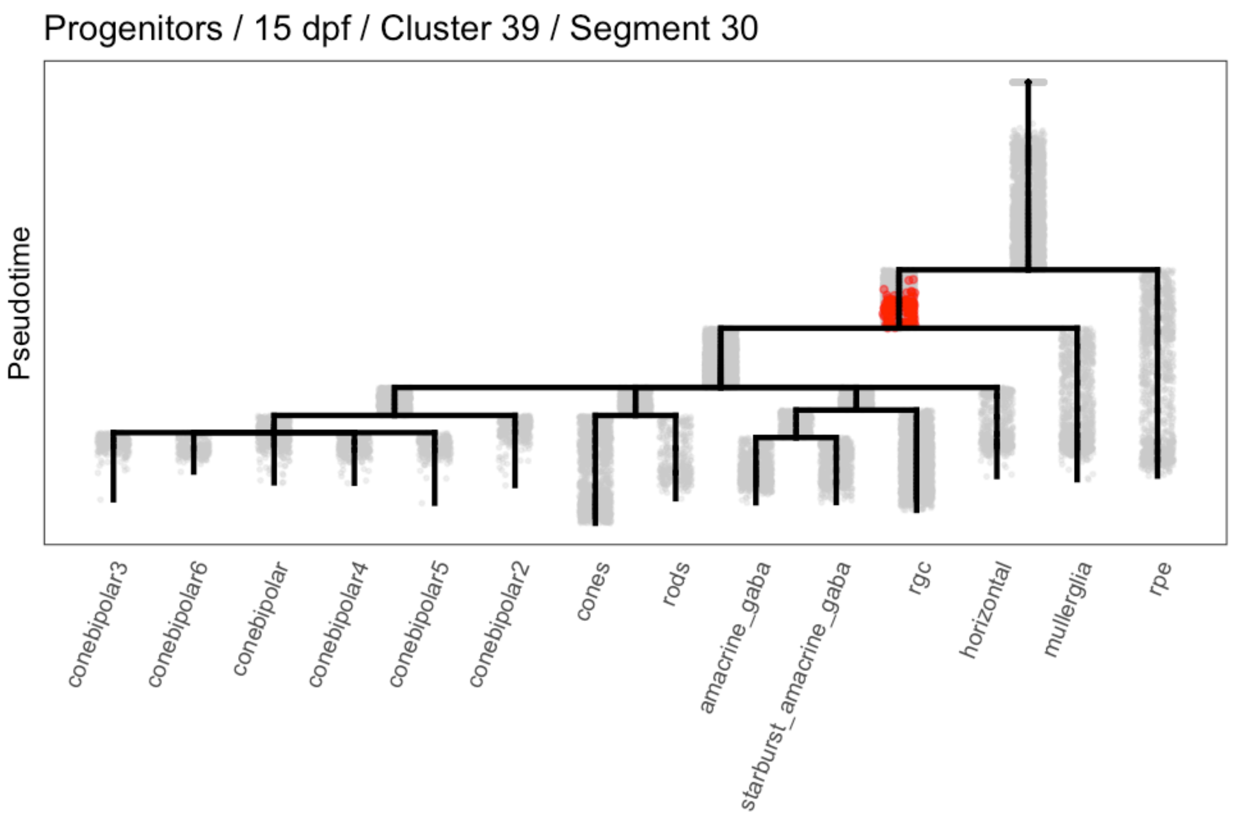

```r
# Progenitors / 15 dpf / Cluster 39 / Segment 29
prog.15d.c39.s29 <- intersect(cellsInCluster(obj, "clus.orig", "12-15d_39"), cellsInCluster(obj,
    "segment", "29"))
obj <- groupFromCells(obj, group.id = "prog.15d.c39.s29", cells = prog.15d.c39.s29)
plotTreeHighlight(obj, "prog.15d.c39.s29", "TRUE", highlight.size = 1, highlight.alpha = 0.5,
    title = "Progenitors / 15 dpf / Cluster 39 / Segment 29")
```

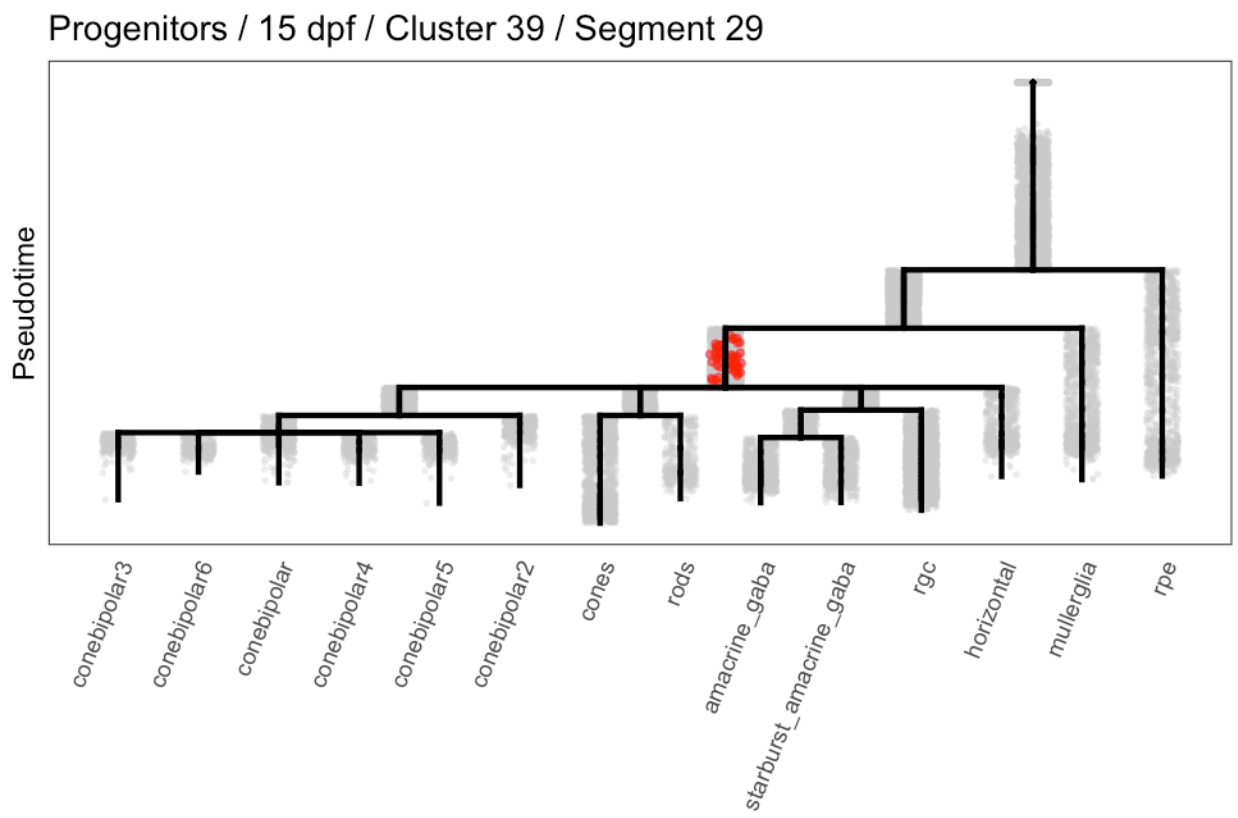

#### Differential expression between neural progenitor populations

Then we determined what was differentially expressed between them.

```
# Compare 15d ('late') vs. early - S29
markers.nb.lve.s29 <- markersAUCPR(obj, cells.1 = prog.15d.c39.s29, cells.2 = prog.early.s29,
    auc.factor = 1.1, effect.size = 0.4)

# Compare 15d ('late') vs. early - S30
markers.nb.lve.s30 <- markersAUCPR(obj, cells.1 = prog.15d.c39.s30, cells.2 = prog.early.s30,
    auc.factor = 1.1, effect.size = 0.4)
```

#### boot.fc

- **object**: An URD object
- **cells.1**: Cells from group 1 of the differential expression
- **cells.2**: Cells from group 2 of the differential expression
- **cells.segment**: All cells in the segment that can be pulled for bootstrapping
- **genes.test**: Genes to test in the bootstrapping
- **exp.fc**: Exp.fc from the original differential expression test to compare for bootstrap
- **exp.data**: Can pre-calculated un-logged expression data to pass to the function (`getUPXData`)
- **n**: (Numeric) Number of bootstrap simulations to run

- Returns list: `p` is the empirical p-value for each differential expression, `boot.fc` contains all of the test information.

```
# Function to bootstrap fold-change
boot.fc <- function(object, cells.1, cells.2, cells.segment, genes.test, exp.fc,
    exp.data = NULL, n = 1000) {

    # Pull random populations of equivalent sizes
    l1 <- length(cells.1)
    l2 <- length(cells.2)
    random.pops <- lapply(1:1000, function(i) {
        y <- sample(x = cells.segment, size = l1 + l2, replace = F)
        return(list(a = y[1:l2], b = y[(l2 + 1):(l1 + l2)]))
    })

    # Get un-logged expression data, if not provided
    if (is.null(exp.data))
        exp.data <- getUPXData(object)

    # Calculate the expression fold-change for each random population
    fc.boot <- as.data.frame(lapply(1:n, function(i) {
        exp.a <- exp.data[genes.test, random.pops[[i]][["a"]]]
        exp.b <- exp.data[genes.test, random.pops[[i]][["b"]]]
        exp.fc <- log2((rowMeans(exp.a)/rowMeans(exp.b)) + 1)
        return(exp.fc)
    }))
    names(fc.boot) <- paste0("rep", 1:n)

    # Figure out p-value (proportion of these that beat provided exp.fc)
    beat.boot <- sweep(fc.boot, 1, exp.fc, ">")
    p.boot <- rowSums(beat.boot)/n

    # Return information
    return(list(p = p.boot, boot.fc = fc.boot))
}
```

##### Empirical p-value

Because these are relatively small populations, there's a decent chance that (due to the variability and noise inherent in scRNAseq data) that choosing any two similarly sized populations would find a number of differentially expressed genes also. Thus, we used an empirically-determined p-value to limit ourselves to differentially expressed genes that probably wouldn't arise by chance. We asked that our real comparison had a greater expression fold-change than two populations from a given segment of the same size chosen at random at least 99% of the time (i.e. $p < 0.01$).

```
# Try a bootstrapping approach to determine which markers are real, vs. which
# ones would arise just from small number of compared cells.  Going to just do it
# on expression fc, so that the computation is reasonably fast.

# Isolate cells from each segment
cells.seg.29 <- cellsInCluster(obj, "segment", "29")
cells.seg.30 <- cellsInCluster(obj, "segment", "30")

# Get un-logged expression data to pass to the function
```

```
exp.data <- getUPXData(obj)

# Run the actual bootstrapping.
boot.s30.lve <- boot.fc(object, cells.1 = prog.15d.c39.s30, cells.2 = prog.early.s30,
    cells.segment = cells.seg.30, genes.test = rownames(markers.nb.lve.s30), exp.fc = markers.nb.lve.s30$exp.fc,
    exp.data = exp.data, n = 1000)

boot.s29.lve <- boot.fc(object, cells.1 = prog.15d.c39.s29, cells.2 = prog.early.s29,
    cells.segment = cells.seg.29, genes.test = rownames(markers.nb.lve.s29), exp.fc = markers.nb.lve.s29$exp.fc,
    exp.data = exp.data, n = 1000)

# Limit markers to those that pass the bootstrap test
markers.nbb.lve.s30 <- markers.nb.lve.s30[which(boot.s30.lve$p <= 0.01), ]
markers.nbb.lve.s29 <- markers.nb.lve.s29[which(boot.s29.lve$p <= 0.01), ]
```

##### Tissue-specific changes

We also then divided genes based on whether they changed in all cells between 24/36 hpf and 15 dpf, or specifically in progenitors. Genes that change in all cells could represent either (a) global transcriptional changes in the tissue, or (b) changes in ambient RNA that is included with most cells based on highly expressed genes during different stages.

```
# Add global stage information to these - genes must change more in progenitors
# than just generally.

# Figure out early/late cells
cells.early <- cellsInCluster(obj, "stage", c("06-24h", "07-36h"))
cells.late <- cellsInCluster(obj, "stage", "12-15d")

# Calculate markers across stages generally with no restrictions
markers.nbball.lve <- markersAUCPR(object = obj, cells.1 = cells.late, cells.2 = cells.early,
    effect.size = -Inf, frac.must.express = 0, auc.factor = 0, genes.use = unique(c(rownames(markers.nbb.lve.s30)
        rownames(markers.nbb.lve.s29))))

# Transfer information to NBB comparisons
markers.nbb.lve.s30$exp.fc.stage <- markers.nbball.lve[rownames(markers.nbb.lve.s30),
    "exp.fc"]
markers.nbb.lve.s30$posFrac_stage1 <- markers.nbball.lve[rownames(markers.nbb.lve.s30),
    "posFrac_1"]

markers.nbb.lve.s29$exp.fc.stage <- markers.nbball.lve[rownames(markers.nbb.lve.s29),
    "exp.fc"]
markers.nbb.lve.s29$posFrac_stage1 <- markers.nbball.lve[rownames(markers.nbb.lve.s29),
    "posFrac_1"]

# Calculate ratios (i.e. how much more does a gene change in progenitors than in
# the entire tissue)
markers.nbb.lve.s30$exp.fc.ratio <- pmin(markers.nbb.lve.s30$exp.fc, 1000) - pmin(markers.nbb.lve.s30$exp.fc.stag
    1000)
markers.nbb.lve.s29$exp.fc.ratio <- pmin(markers.nbb.lve.s29$exp.fc, 1000) - pmin(markers.nbb.lve.s29$exp.fc.stag
    1000)
```

```
markers.nbb.lve.s30$posFrac.ratio <- markers.nbb.lve.s30$posFrac_1/markers.nbb.lve.s30$posFrac_stage1
markers.nbb.lve.s29$posFrac.ratio <- markers.nbb.lve.s29$posFrac_1/markers.nbb.lve.s29$posFrac_stage1
```

##### Limit to well-expressed

We also limited ourselves to genes that had a decent level of expression. (In this case, they were detected in at least 20% of progenitor cells, and had a mean expression of at least 0.8.)

```
# All genes that change in segment 30
markers.nbbexp.lve.s30 <- markers.nbb.lve.s30[Reduce(intersect, list(which(markers.nbb.lve.s30$posFrac_1 >=
    0.2), which(markers.nbb.lve.s30$nTrans_1 >= 0.8))), ]

# All genes that change in segment 30
markers.nbbexp.lve.s29 <- markers.nbb.lve.s30[Reduce(intersect, list(which(markers.nbb.lve.s29$posFrac_1 >=
    0.2), which(markers.nbb.lve.s29$nTrans_1 >= 0.8))), ]

# Genes that change in segment 30 more than in the entire tissue
markers.nbbselect.lve.s30 <- markers.nbb.lve.s30[Reduce(intersect, list(which(markers.nbb.lve.s30$posFrac_1 >=
    0.2), which(markers.nbb.lve.s30$nTrans_1 >= 0.8), which(markers.nbb.lve.s30$exp.fc.ratio >=
    1.2), which(markers.nbb.lve.s30$posFrac.ratio >= 1.1))), ]

# Genes that change in segment 29 more than in the entire tissue
markers.nbbselect.lve.s29 <- markers.nbb.lve.s29[Reduce(intersect, list(which(markers.nbb.lve.s29$posFrac_1 >=
    0.2), which(markers.nbb.lve.s29$nTrans_1 >= 0.8), which(markers.nbb.lve.s29$exp.fc.ratio >=
    1.2), which(markers.nbb.lve.s29$posFrac.ratio >= 1.1))), ]
```

##### Result

That recovered a total of 71 genes that vary in progenitors between 24/36 hpf and 15 dpf, of which 16 change more in neural progenitors than the rest of the tissue.

```
# All genes that change in progenitors
unique(c(rownames(markers.nbbexp.lve.s29), rownames(markers.nbbexp.lve.s30)))
```

```
##  [1] "hbbe2"              "hbz"                "ba1.1"
##  [4] "rho"                "crabp1a"            "si:ch211-251b21.1"
##  [7] "hbaa1"              "pde6h"              "tsc22d3"
## [10] "si:xx-by187g17.1"   "rpe65a"             "zgc:153704"
## [13] "arr3a"              "lin7a"              "crygm1"
## [16] "gnat1"              "ptgdsb.1"           "gngt1"
## [19] "rgs16"              "cabp2a"             "junba"
## [22] "crygm2b"            "zgc:112320"         "si:dkey-183i3.5"
## [25] "krt91"              "cabp5a"             "sagb"
## [28] "crygmx"             "crygm2a"            "scinla"
## [31] "rbp4l"              "gngt2b"             "rs1a"
## [34] "mt2"                "fosab"              "cebpd"
## [37] "snap25b"            "CNDP1"              "crabp2a"
## [40] "cryba4"             "jdp2b"              "cst3"
## [43] "higd1a"             "mt-nd3"             "si:dkey-16p21.8"
## [46] "crybb1"             "crygn2"             "gadd45ba"
## [49] "gapdhs"             "eno1a"              "mif"
## [52] "ggctb"              "ckbb"               "glula"
```

```
## [55] "tsc22d1"           "sod2"              "btg2"
## [58] "sod1"              "stmn1b"            "si:dkey-238o13.4"
## [61] "fabp11a"           "mdkb"              "gstp1"
## [64] "slc3a2b"           "si:dkey-238c7.12"  "CABZ01102240.1"
## [67] "atp5ia"            "atpif1b"           "cadm3"
## [70] "si:ch73-46j18.5"
```

```r
# Genes that change in progenitors more than the rest of the tissue
unique(c(rownames(markers.nbbselect.lve.s29), rownames(markers.nbbselect.lve.s30)))
```

```
##  [1] "si:ch211-114n24.6" "rps29"             "rrm2.1"
##  [4] "si:ch211-193l2.6"  "si:dkey-238o13.4"  "crabp1a"
##  [7] "si:ch211-251b21.1" "CNDP1"             "junba"
## [10] "crabp2a"           "cryba4"            "crybb1"
## [13] "crygn2"            "fabp11a"           "si:dkey-238c7.12"
## [16] "cadm3"
```

### Long-term undifferentiated states

We wanted to look for differences in the transcriptional states of long-term progenitors between the retina and the hypothalamus. In the retina, cells persist in progenitor states, while in the hypothalamus, the progenitor state is short-lived, but cells persist in a neural precursor state.

#### Identify populations to compare

We defined progenitor / precursor / neuron populations based on their location in the tree, cross-referenced with the expression of markers of each of these types

```r
obj@group.ids$precursor.group <- NA

cells.s31 <- intersect(cellsInCluster(obj, "segment", "31"), whichCells(obj, "pseudotime",
    c(0.05, 1)))
obj@group.ids[cells.s31, "precursor.group"] <- "1a_prog_transient"

cells.prog.late <- cellsInCluster(obj, "segment", c("30", "29"))
obj@group.ids[cells.prog.late, "precursor.group"] <- "1b_prog_longterm"

cells.precursor <- intersect(cellsInCluster(obj, "segment", c("24", "25", "26", "15")),
    whichCells(obj, "pseudotime", c(0, 0.535)))
obj@group.ids[cells.precursor, "precursor.group"] <- "2_precursor"

cells.neurons <- setdiff(whichCells(obj, "pseudotime", c(0.535, 1)), cellsInCluster(obj,
    "segment", c("6", "11")))
obj@group.ids[cells.neurons, "precursor.group"] <- "3_neurons"
```

```r
# Colors for ggplot
stage.colors <- RColorBrewer::brewer.pal(12, "Paired")[c(10, 9, 7, 1)]

# Plot tree to show where the groups are
plotTree(obj, "precursor.group", discrete.colors = stage.colors)
```

```
## Warning: Removed 2530 rows containing missing values (geom_point).
```

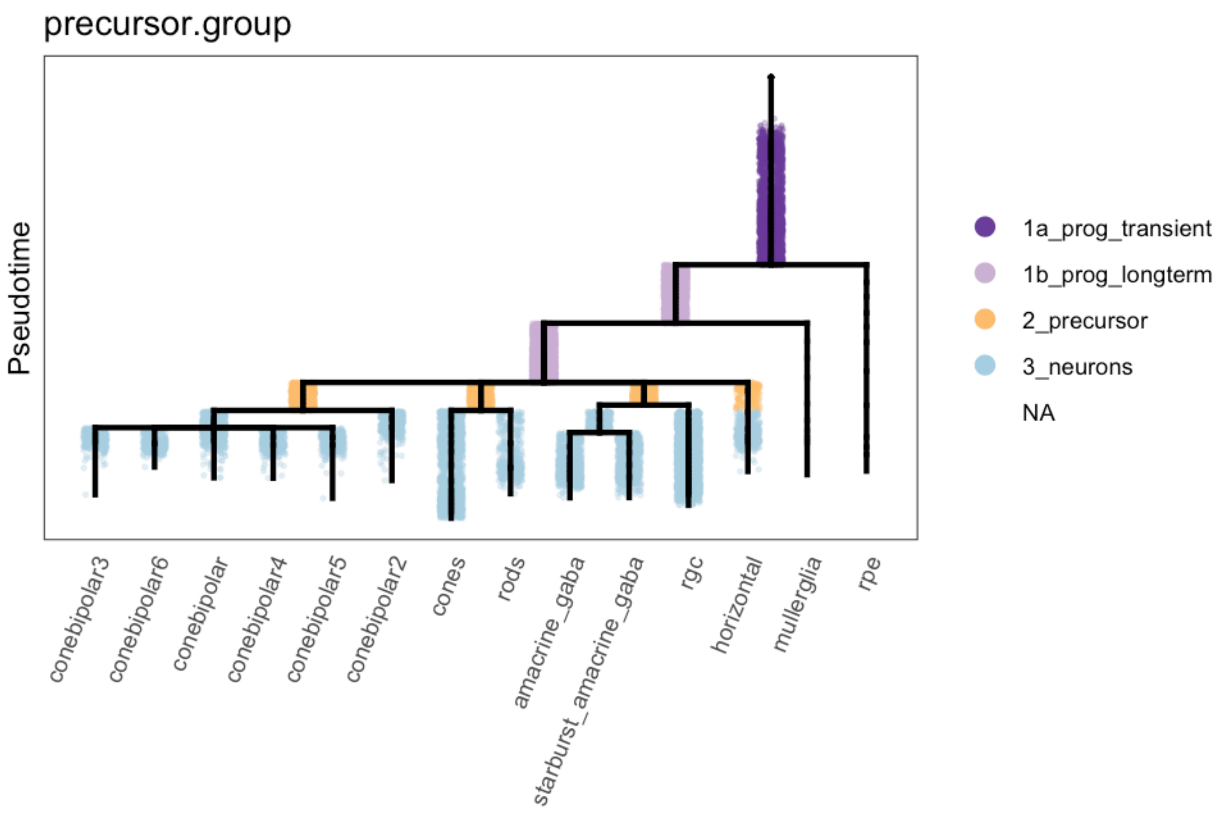

```r
# Plot genes in each group
plotDot(obj, genes = c("rx1", "foxd1", "her2", "hes2.2", "insm1a", "neurod4", "foxg1b",
    "elavl3", "elavl4"), clustering = "precursor.group", scale.by = "area") + theme_bw()
```

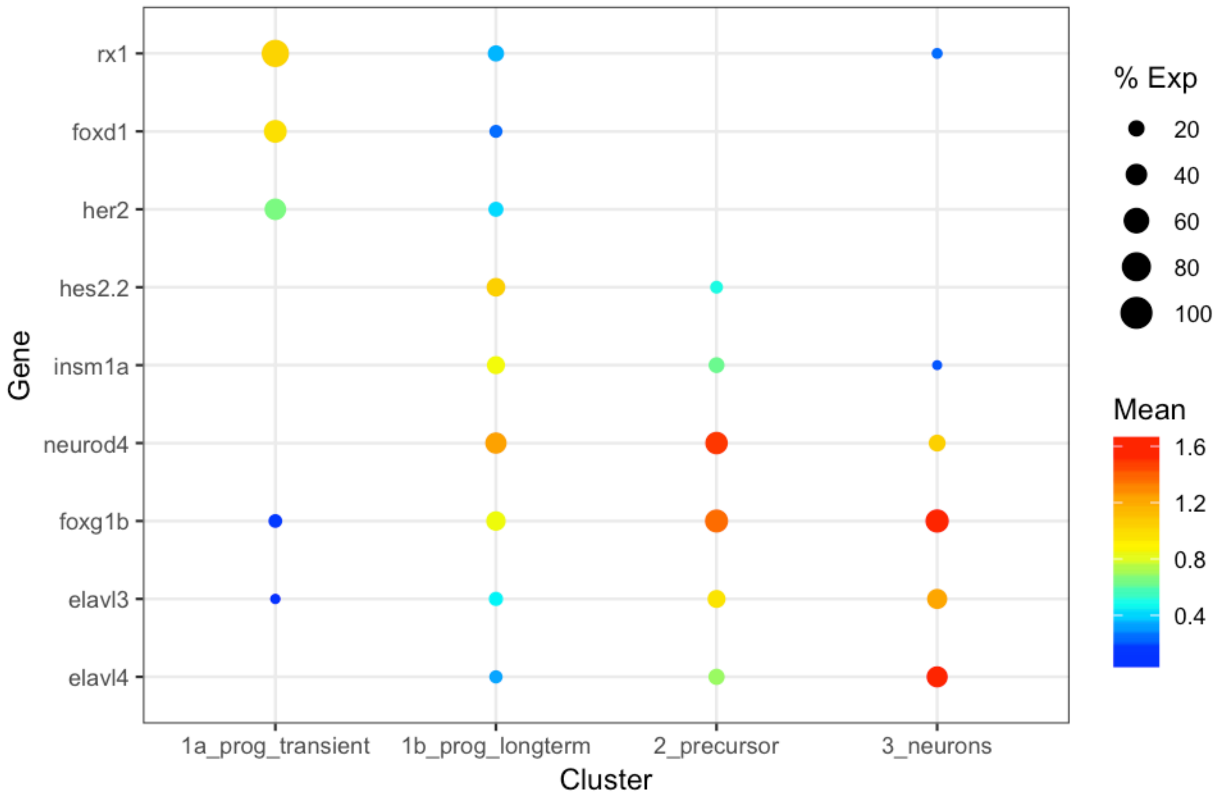

#### Determine proportion of cells in each state

We then determined the proportion of cells from different stages that fell into each of these transcriptional states.

```
# We combined stages to reduce number of plots
obj@group.ids$stage.collapsed <- plyr::mapvalues(x = obj@group.ids$stage, from = c("01-12h",
    "02-14h", "03-16h", "04-18h", "05-20h", "06-24h", "07-36h", "08-2d", "09-3d",
    "10-5d", "11-8d", "12-15d"), to = c(rep("01-12h-24h", 6), rep("02-36h-3d", 3),
    rep("03-5d-15d", 3)))

# Count number of cells from each stage group in each precursor group
stage.group.count <- plyr::count(obj@group.ids, vars = c("stage.collapsed", "precursor.group"))

# Remove cells that weren't part of a precursor group
stage.group.count <- stage.group.count[complete.cases(stage.group.count), ]

# Cast into a data frame and convert NA to 0 (no cells of that type observed)
stage.group.df <- reshape2::dcast(stage.group.count, formula = stage.collapsed ~
    precursor.group)
```

```
## Using freq as value column: use value.var to override.
```

```
stage.group.df[is.na(stage.group.df)] <- 0

# Normalize by the number of precursors from each stage group
```

```r
stage.group.df[, 2:5] <- sweep(stage.group.df[, 2:5], 1, rowSums(stage.group.df[,
    2:5]), "/")

# Melt for ggplot
stage.group.df.melt <- reshape2::melt(stage.group.df, id.vars = "stage.collapsed")

# Plot proportions
ggplot(stage.group.df.melt, aes(x = variable, y = value, group = stage.collapsed,
    fill = variable)) + geom_bar(stat = "identity") + facet_wrap(~stage.collapsed) +
    theme_bw() + scale_fill_manual(values = stage.colors) + theme(axis.title.x = element_blank(),
    axis.text.x = element_blank(), axis.ticks.x = element_blank())
```

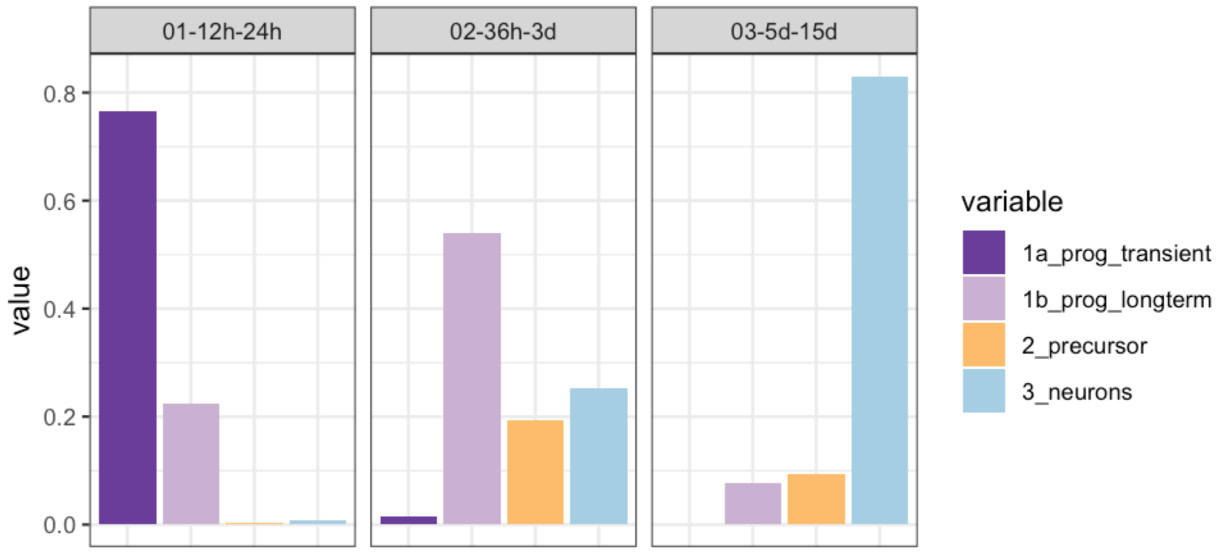

### Hypothalamus: 1 - URD object & doublet removal

*Jeff Farrell*

*8/22/2019*

#### Contents

Import data into URD 1
    Convert Seurat object to URD . . . 1
    Combined individual stage clustering . . . 1

Calculate highly variable genes 2

Calculate KNN graph and remove outliers 7

Remove cell type doublets 8

#### Import data into URD

##### Convert Seurat object to URD

We first loaded a Seurat object that contained just cells from the clusters that belonged to the hypothalamus from each stage.

```
suppressPackageStartupMessages(library(URD))
suppressPackageStartupMessages(library(Seurat))
```

```
base.path <- "~/urd-cluster-bushra/"

# Load Seurat object that has been cropped to hypothalamus cells
object.seurat <- readRDS(paste0(base.path, "obj/hypo_seurat.rds"))

# Convert to URD object
suburd <- seuratToURD(object.seurat)
```

##### Combined individual stage clustering

Bushra had performed individual clusterings across each stage with different resolutions. Here, it was better to create a single identifier that included stage + cluster information to combine all those clusterings (while preventing any overlap).

```
stages <- sort(unique(suburd@meta$stage))
clust.res.used <- paste0("res.", c("4.5", "4", "5", "5", "4.5", "5", "6",
    "6", "6", "5.5", "6", "5"))
names(clust.res.used) <- stages
suburd@group.ids$cluster <- NA
for (stage in stages) {
    suburd@group.ids[cellsInCluster(suburd, "stage", stage), "cluster"] <- paste0(stage,
        "-", suburd@group.ids[cellsInCluster(suburd, "stage", stage), clust.res.used[stage]])
}
```

### Calculate highly variable genes

We calculated highly variable genes for each stage, used genes that were found as highly in at least two stages, but were not mitochondrial, ribosomal, heat-shock protein, or tandem duplicated genes.

```
# Calculated on each stage separaely, final gene list was all genes
# that were 'variable' in at least two stages NB: For a couple of
# stages, the gamma fit was poor -- the library size distribution
# seemed bimodal. Have seen this before in 10X data, but not sure what
# it means.
var.genes.by.stage <- lapply(stages, function(stage) {
    findVariableGenes(suburd, cells.fit = cellsInCluster(suburd, "stage",
        stage), set.object.var.genes = F, diffCV.cutoff = 0.3, main.use = stage,
        do.plot = T)
})
```

**01-12h**

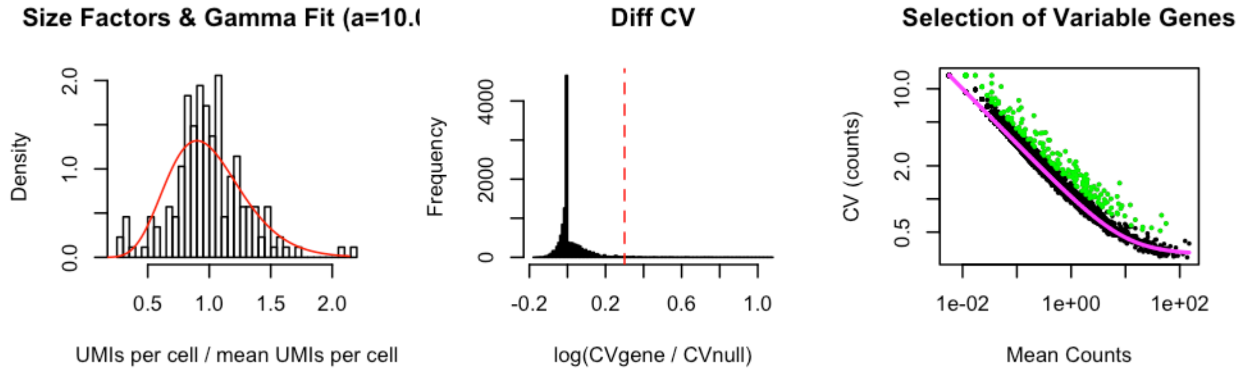

**02-14h**

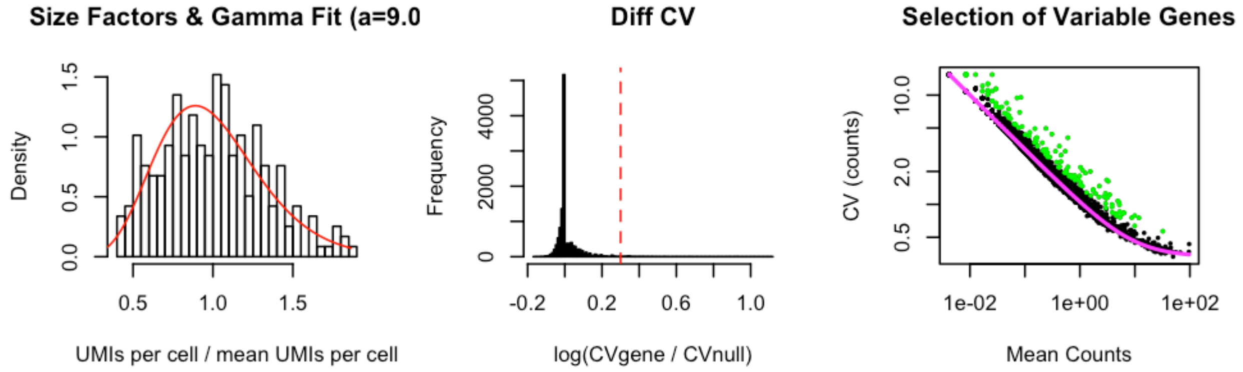

**03-16h**

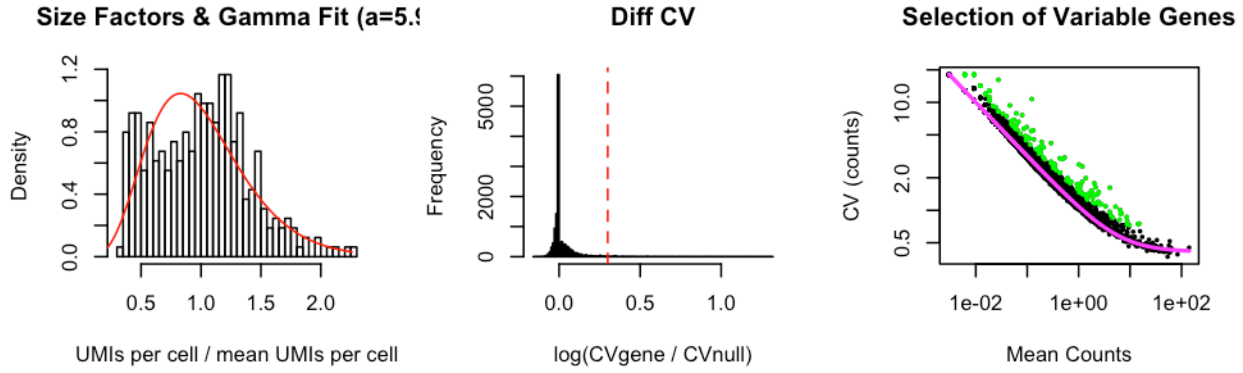

**04-18h**

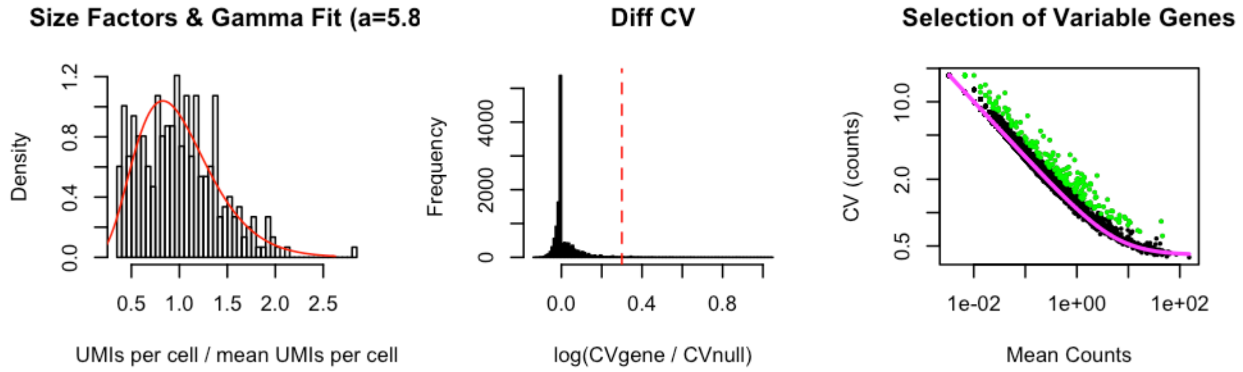

**05-20h**

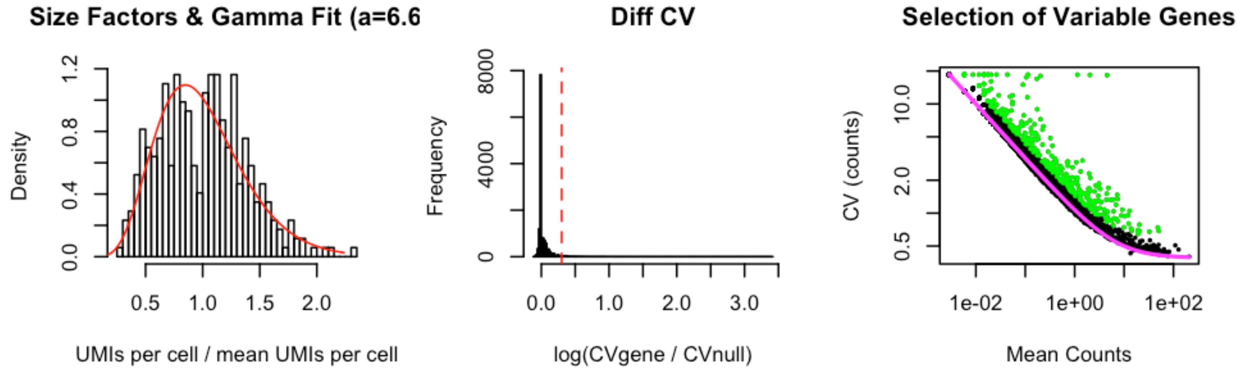

**06-24h**

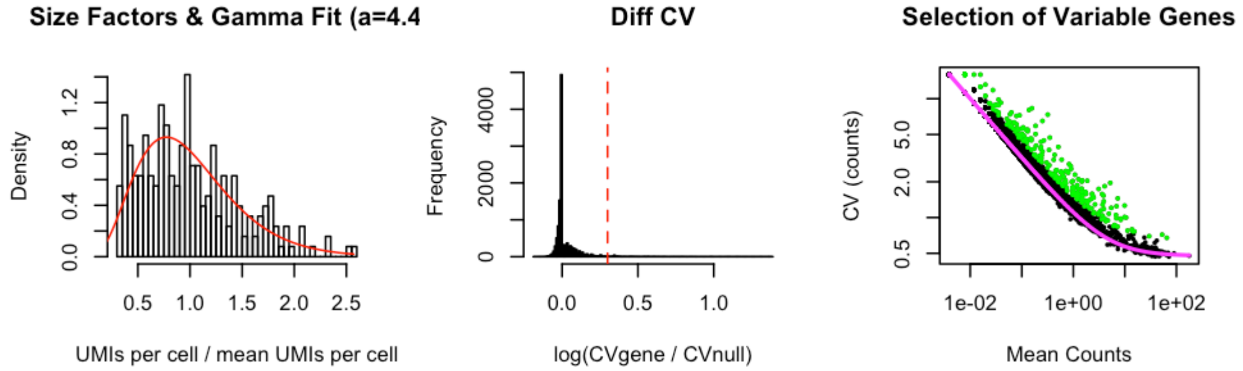

## 07-36h

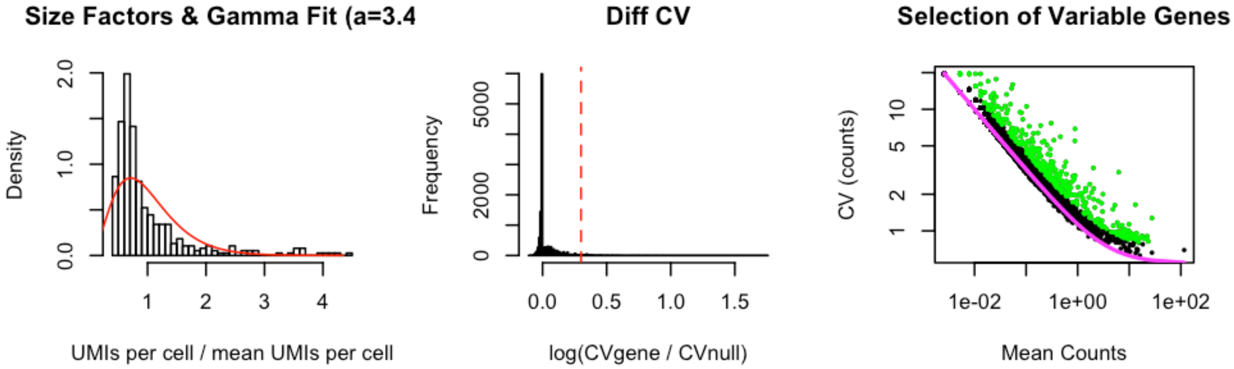

## 08-2d

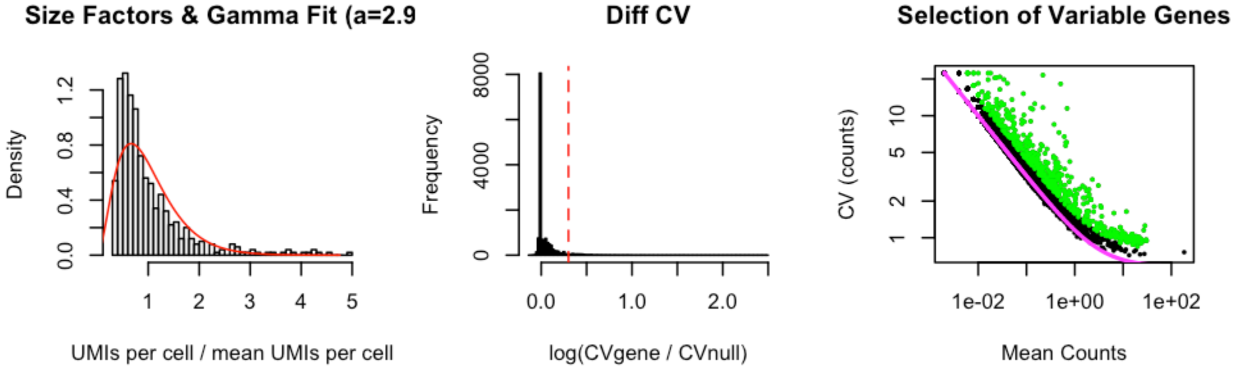

## 09-3d

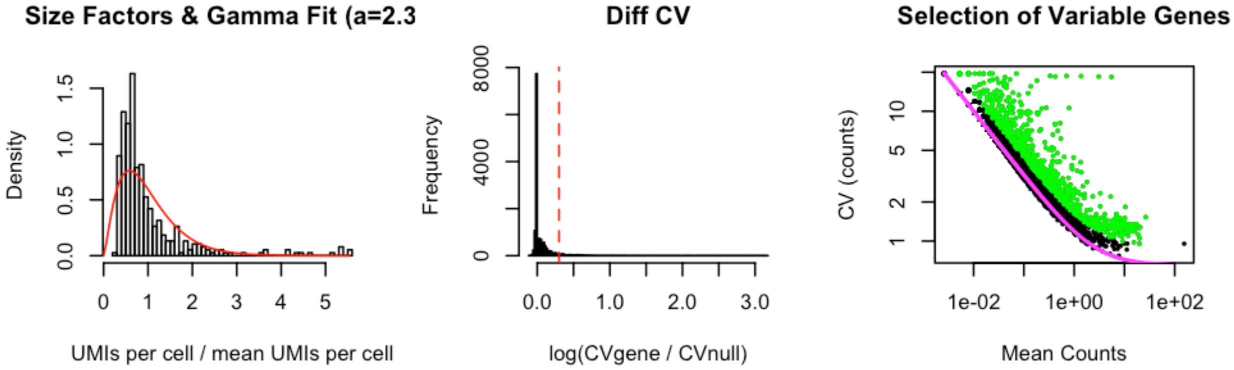

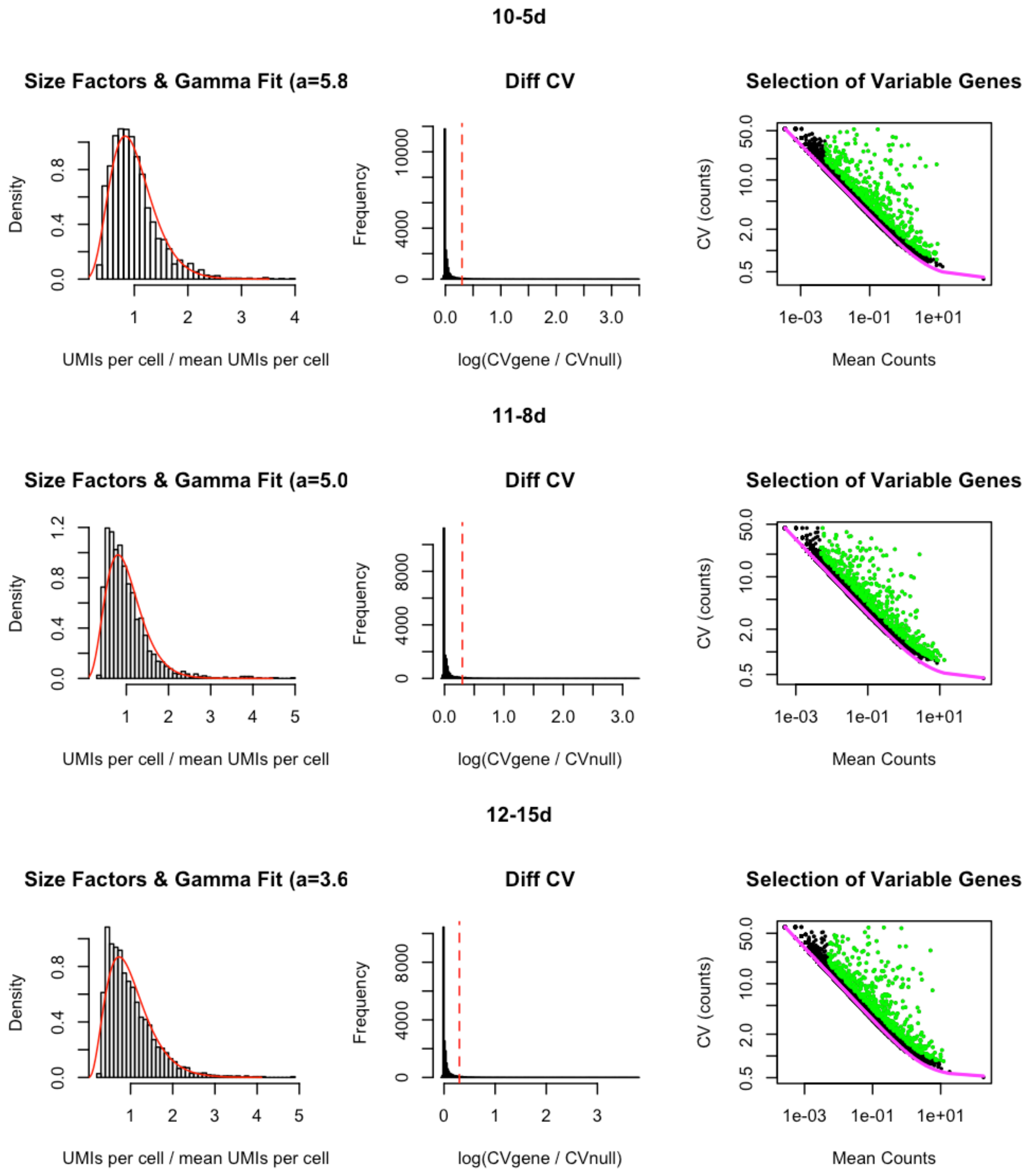

```r
names(var.genes.by.stage) <- stages
var.genes <- sort(unique(unlist(var.genes.by.stage)))
print(paste0("Length of variable genes is ", length(var.genes)))
```

```
## [1] "Length of variable genes is 1783"
```

```r
var.genes.twice <- names(which(table(unlist(var.genes.by.stage)) >= 2))
print(paste0("Length of variable genes shared across at least 2 stages is ",
    length(var.genes.twice)))
```

```
## [1] "Length of variable genes shared across at least 2 stages is 957"
```

```
# Remove mitochondrial genes
var.mito <- grep("^mt-|^AC0", var.genes.twice, value = T)
# Remove ribosomal genes
var.ribo <- grep("^rps|^rpl", var.genes.twice, value = T)
# Remove hsp genes
var.hsp <- grep("^hsp", var.genes.twice, value = T)
# Remove genes with duplicates
var.dups <- grep("of many", var.genes.twice, value = T)
suburd@var.genes <- setdiff(var.genes.twice, c(var.mito, var.ribo, var.hsp,
    var.dups))
print(paste0("Length of final variable genes list (after removing mito, ribo, hsp genes) is ",
    length(suburd@var.genes)))
```

```
## [1] "Length of final variable genes list (after removing mito, ribo, hsp genes) is 856"
```

To prevent downstream problems, we also removed any cells from the data that had the exact same expression of the variable genes (i.e. cells with completely duplicated coordinates in the high-dimensional space we would use for analysis downstream).

```
# Check for duplicate data points - cells with exact same expression of
# variable genes
vg.dups <- duplicated(as.data.frame(as.matrix(t(suburd@logupx.data[suburd@var.genes,
    ]))))
if (length(which(vg.dups)) > 0) {
    print(paste("Removing", length(which(vg.dups)), "cell(s) with duplicated variable gene expression."))
    not.dup.cells <- colnames(suburd@logupx.data)[!vg.dups]
    suburd <- urdSubset(suburd, not.dup.cells)
}
```

```
## [1] "Removing 1 cell(s) with duplicated variable gene expression."
```

### Calculate KNN graph and remove outliers

We then calculated a k-nearest neighbor graph and removed cells that had unusual distance to their nearest neighbor, or unusual distance to their 20th nearest neighbor (given their distance to their nearest neighbor). These sorts of outliers often cause problems or skew diffusion maps (used downstream).

```
# Calculate k-nn
suburd <- calcKNN(suburd)
```

```
# Check what the outliers are
outliers <- knnOutliers(suburd, nn.1 = 1, nn.2 = 20, x.max = 40, slope.r = 1.1,
    int.r = 3, slope.b = 0.66, int.b = 11.5, title = "Identifying Outliers by k-NN Distance.")
```

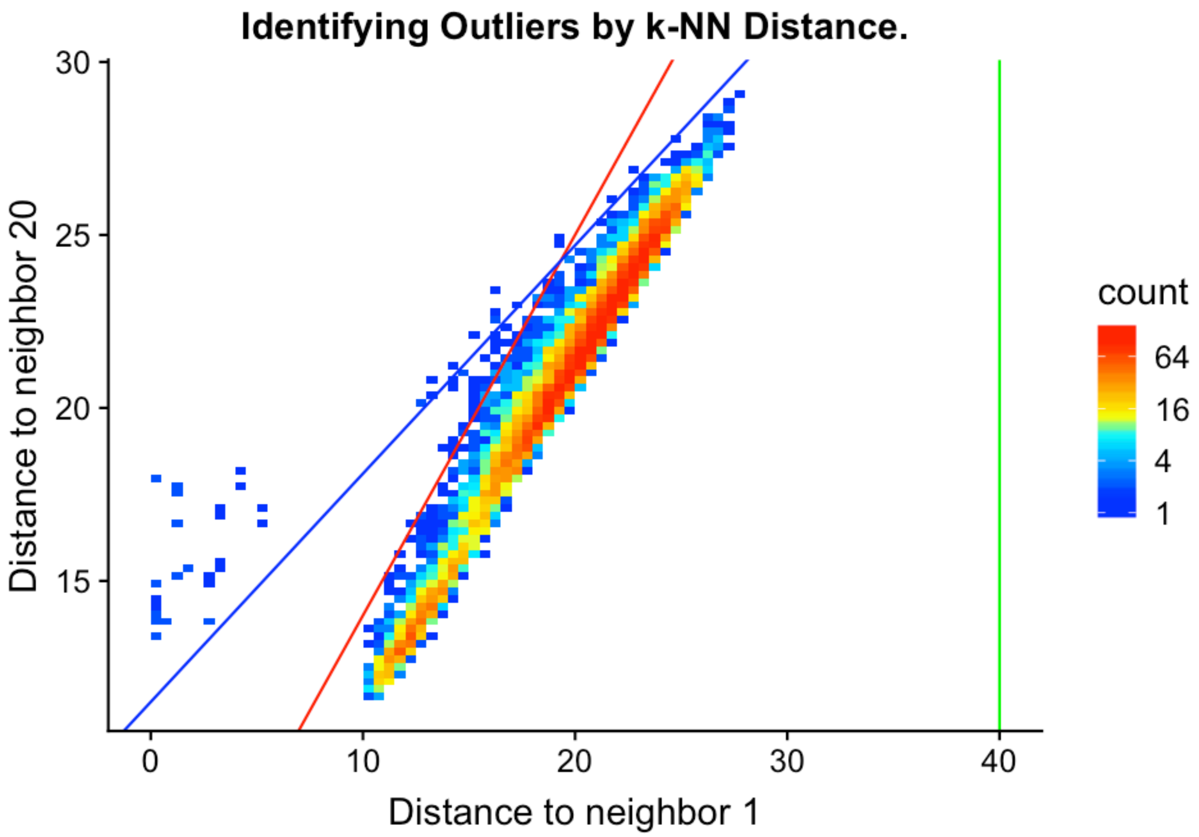

```
length(outliers)
```

```
## [1] 87
```

```
suburd <- urdSubset(suburd, cells.keep = setdiff(colnames(suburd@logupx.data),
    outliers))
```

### Remove cell type doublets

#### Add UMAP projection

While not strictly required, a UMAP projection can make it easier to assess the expression of NMF modules and whether thresholds for overlap are set correctly.

```
## add UMAP command

# Load pre-calculated UMAP
umap <- readRDS(paste0(base.path, "/umap/umap_hypo.rds"))

# Add projection to URD object
suburd@tsne.y <- umap
```

#### Load NMF results and import into object

NMF results were calculated by providing `suburd@logupx.data` to an external NMF pipeline written in Python. The output results are imported here, scaled, and added to the URD object.

```
# Load the NMF results
load(paste0(base.path, "/NMF/hypo/result_tbls.Robj"))

# The results object contains NMF runs for several K values.  k=28 was
# chosen for this tissue, so this extracts the results for that
# particular parameter
k.use <- "28"
nmf.cells <- result_obj[[paste0("K=", k.use)]][[1]]$C
rownames(nmf.cells) <- paste0("nmf", 1:nrow(nmf.cells))
colnames(nmf.cells) <- gsub("\\.", "-", colnames(nmf.cells))
nmf.genes <- result_obj[[paste0("K=", k.use)]][[1]]$G
colnames(nmf.genes) <- paste0("nmf", 1:nrow(nmf.cells))

# Scale NMF results 0-1
nmf.cells.scaled <- sweep(nmf.cells, 1, apply(nmf.cells, 1, max), "/")

# Add scaled NMF results to the URD object
suburd@nmf.c1 <- as(t(as.matrix(nmf.cells.scaled)), "dgCMatrix")
```

#### Select cell-type specific modules

Several NMF modules will be poor markers of cell types — these are often modules driven mostly be the expression of 1-2 genes (where the gene loading of the first gene is much greater than that of the fourth gene, for instance), or modules that don't exhibit any restriction in a tSNE or UMAP projection.

```
# Plot size parameters
plot.height = 6
plot.width = 16
dpi = 150

# Plot every module to determine which exhibit cell-type specificity
# This saves directly to the hard drive: two example plots are shown
# below.

# for (n in colnames(suburd@nmf.c1)) { png(paste0(path, '/doublets/',
# subset, '-plots/', n, '.png'), width=dpi*plot.width,
# height=dpi*plot.height) plot(plotDim(suburd, n)) dev.off() }

gridExtra::grid.arrange(grobs = list(plotDim(suburd, "nmf2", plot.title = "nmf2: exhibits poor restriction"),
    plotDim(suburd, "nmf27", plot.title = "nmf27: exhibits good cell-type")),
    ncol = 2)
```

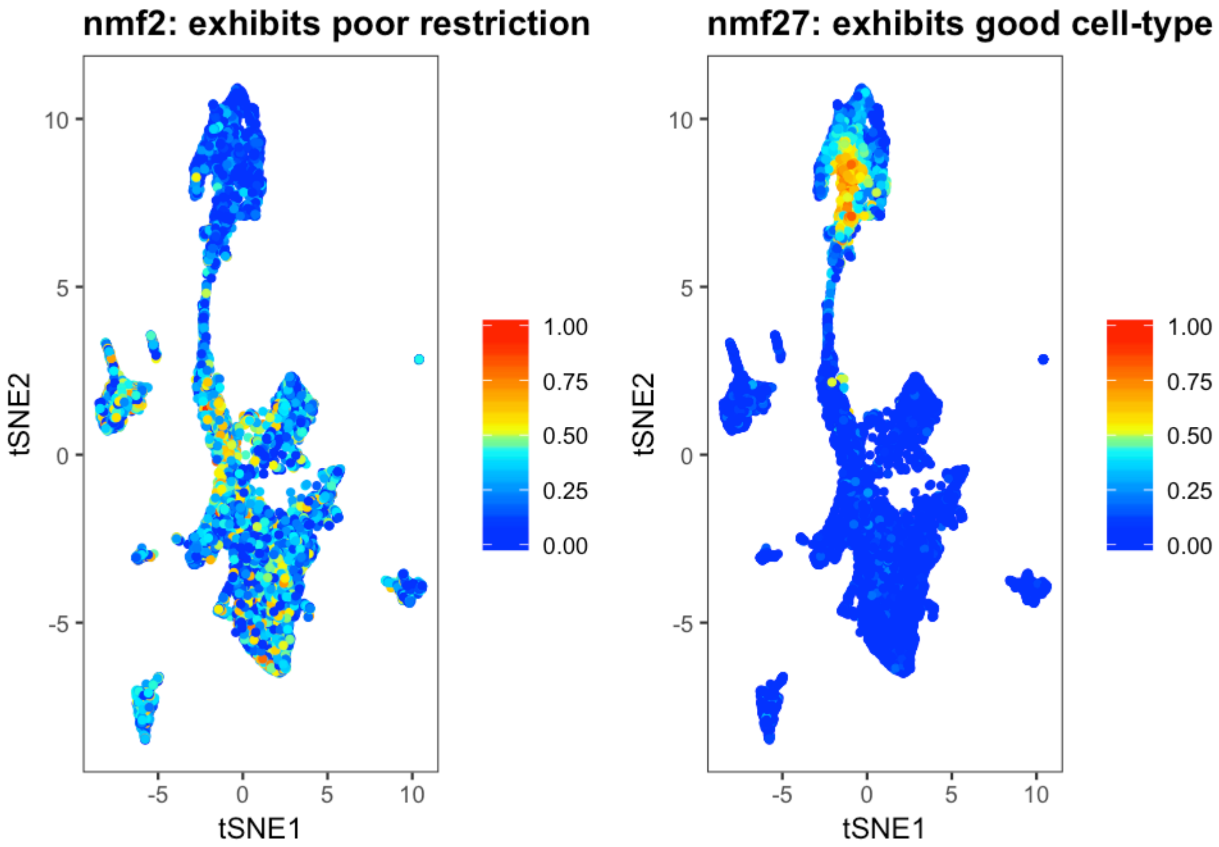

```
# Module Gene 1 : Gene 4 Ratios
top.genes <- result_obj[[paste0("K=", k.use)]][[1]]$top30genes
top.weights <- top.genes[, grep("Weights", colnames(top.genes), value = T)]
colnames(top.weights) <- paste0("nmf", 1:nrow(nmf.cells))
top.weights.ratio <- top.weights[1, ]/top.weights[4, ]

# Which modules exhibit cell-type restriction?
modules.bad.ratio <- names(top.weights.ratio)[which(top.weights.ratio >
    3)]
unrestricted.modules <- paste0("nmf", c("12", "19", "24", "28"))
good.modules <- setdiff(colnames(suburd@nmf.c1), c(modules.bad.ratio, unrestricted.modules))
```

#### Determine which module pairs to use for doublet removal

We consider NMF modules pairwise and only use those pairs that don't are non-overlapping in the data. (In other words, NMF modules that are mutually exclusive in the majority of the data.) Here, we determine thresholds for selecting those module pairs.

```
# Determine overlaps between module pairs
nmf.doublet.combos <- NMFDoubletsDefineModules(suburd, modules.use = good.modules,
    module.thresh.high = 0.4, module.thresh.low = 0.15)

# Determine thresholds for NMF modules
frac.overlap.max = 0.03
frac.overlap.diff.max = 0.11
```

```
module.expressed.thresh = 0.33

# Determine which module pairs to use for doublets
NMFDoubletsPlotModuleThresholds(nmf.doublet.combos, frac.overlap.max = frac.overlap.max,
    frac.overlap.diff.max = frac.overlap.diff.max)
```

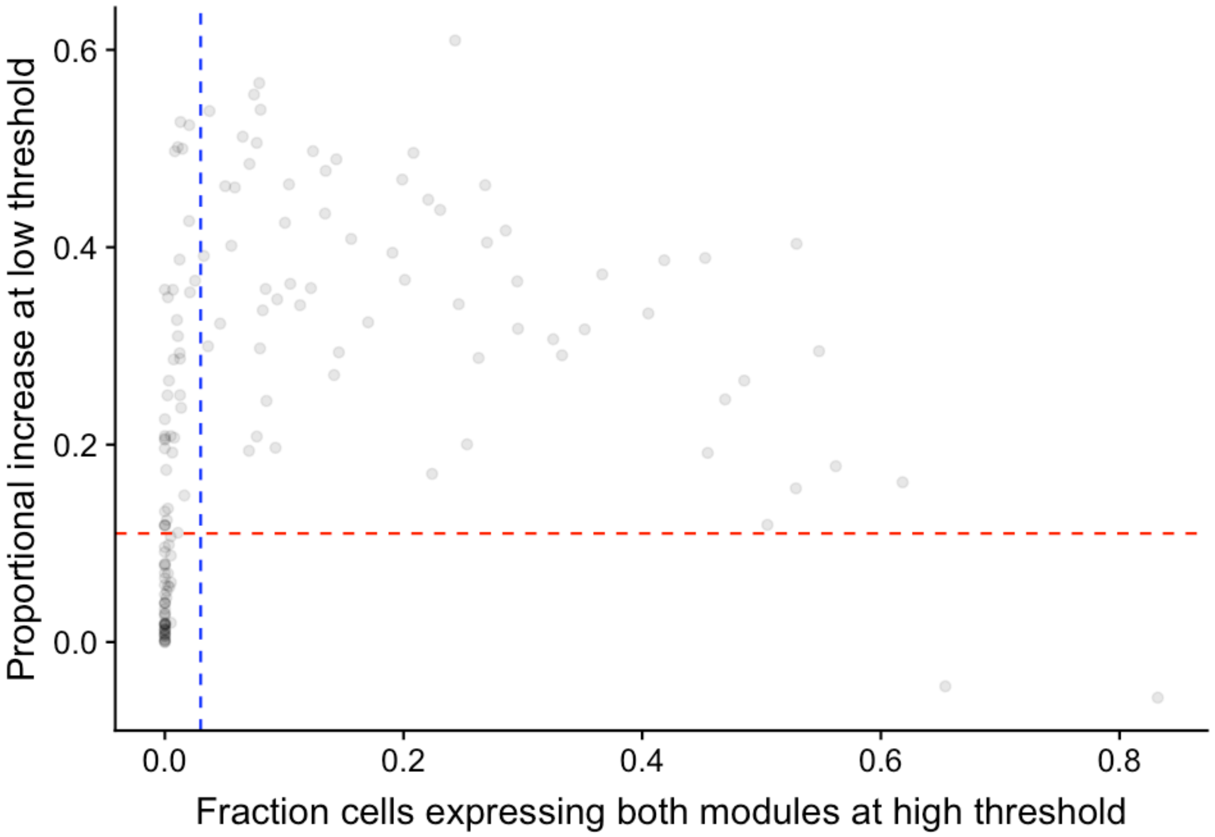

```
# These commands save plots directly to the hard-drive.

# Make plots to see how your thresholds are
NMFDoubletsPlotModuleCombos(suburd, path = paste0(path, "/doublets/", subset,
    "-doublet-combos/"), module.combos = nmf.doublet.combos, module.expressed.thresh = module.expressed.thresh,
    frac.overlap.max = frac.overlap.max, frac.overlap.diff.max = frac.overlap.diff.max,
    boundary = "pass", sort = "near", n.plots = 25)
NMFDoubletsPlotModuleCombos(suburd, path = paste0(path, "/doublets/", subset,
    "-ok-combos/"), module.combos = nmf.doublet.combos, module.expressed.thresh = module.expressed.thresh,
    frac.overlap.max = frac.overlap.max, frac.overlap.diff.max = frac.overlap.diff.max,
    boundary = "discarded", sort = "near", n.plots = 25)
```

```
# Define doublet cells
nmf.doublets <- NMFDoubletsDetermineCells(suburd, nmf.doublet.combos, module.expressed.thresh = module.expressed.
    frac.overlap.max = frac.overlap.max, frac.overlap.diff.max = frac.overlap.diff.max)  # 49 cells / 11307 cells

# Plot doublet cells on the UMAP
suburd <- groupFromCells(suburd, "nmf.doublets", cells = nmf.doublets)
plot(plotDimHighlight(suburd, clustering = "nmf.doublets", cluster = "TRUE",
    plot.title = paste0("NMF doublets: ", length(nmf.doublets), " cells"),
```

```
    point.size = 2, highlight.color = "blue"))
```

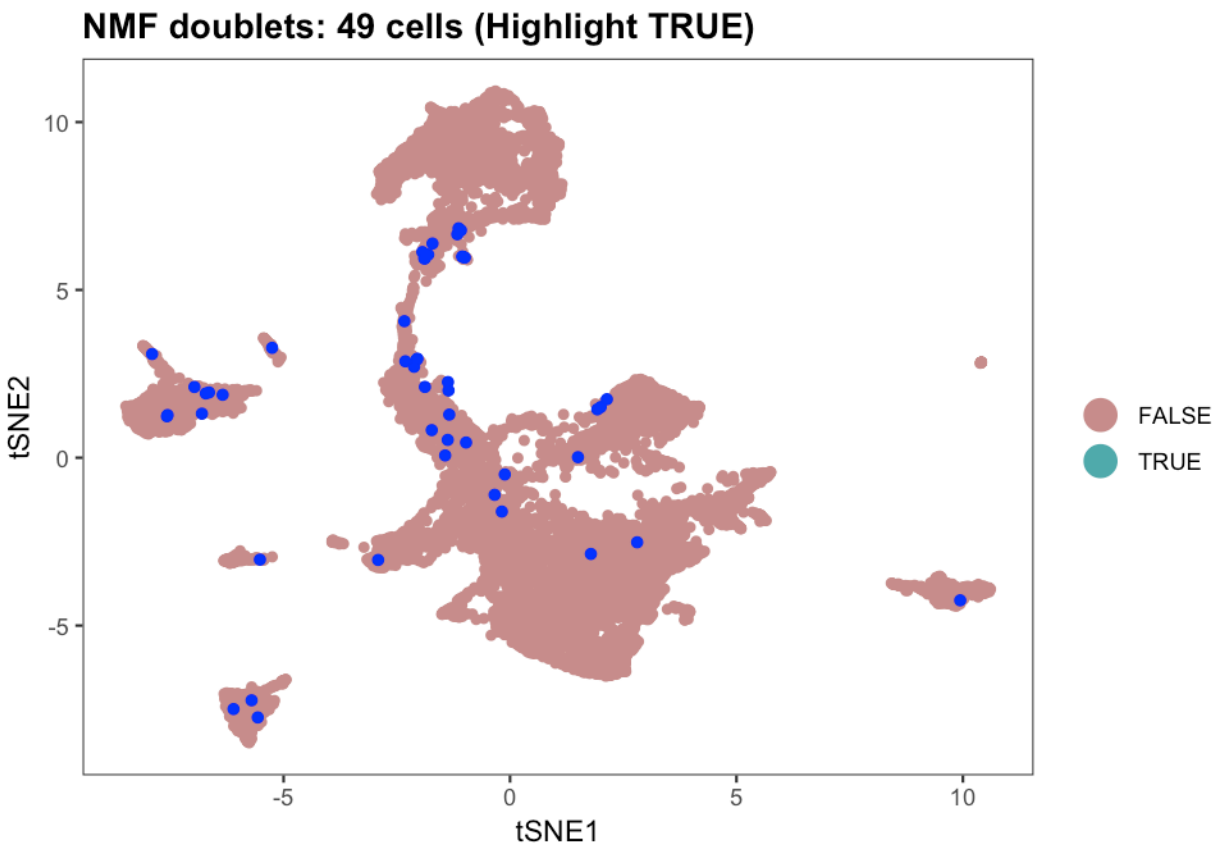

```
# Crop object to exclude doublets
suburd.cropped <- urdSubset(suburd, cells.keep = setdiff(colnames(suburd@logupx.data),
    nmf.doublets))
```

And then save the completed object for use downstream in building a tree using URD.

```
saveRDS(suburd.cropped, file = paste0(base.path, "/obj/URD_hypo_ND.rds"))
```

### Hypothalamus: 2 - URD tree

*Jeff Farrell*

*9/07/2019*

Build the URD tree 12

Save the URD tree 13

#### Load data

```
suppressPackageStartupMessages(library(URD))
suppressPackageStartupMessages(library(Seurat))
```

```
base.path <- "~/urd-cluster-bushra/"

# Load procesed URD object
object <- readRDS(paste0(base.path, "obj/URD_hypo_ND.rds"))
```

#### Processed on the cluster

Most of the following steps were run on a computing cluster. These individual tissue subsets can be run on a modern, well-equipped laptop. The use of a computing cluster allows multiple parameter choices to be tried in parallel, and also allows further parallelization of the random walk procedure, speeding it up. Below, we should the commands that one would run on their laptop, and then generally load the pre-processed results from the cluster that were used in the paper. If you want to parallelize your own processing on a compute cluster, the scripts we used will be available at http://github.com/farrellja/URD/cluster/

### Calculate diffusion map and pseudotime

These two steps are run in the cluster script URD-DM-PT.R.

#### Calculate diffusion map

```
# To run locally: Calculate a diffusion map projection
object <- calcDM(object, knn = 100, sigma.use = 8)
```

```
# Or: Load a pre-computed diffusion map projection
dm <- readRDS(paste0(base.path, "dm/dm_hypoND_knn-100_sigma-8.rds"))
object <- importDM(object, dm)
```

```
# Plot diffusion maps
stage.colors <- c("antiquewhite", "#FFCCCC", "#99CC00", "#33CC00", "cyan3",
    "gold", "goldenrod", "darkorange", "indianred1", "plum", "deepskyblue2",
    "lightgrey")

# Plot by stage
plotDimArray(object = object, reduction.use = "dm", dims.to.plot = 1:18,
    label = "stage", plot.title = "", outer.title = "Diffusion map labeled by Stage",
    legend = T, alpha = 0.45, discrete.colors = stage.colors)
```

Diffusion map labeled by Stage

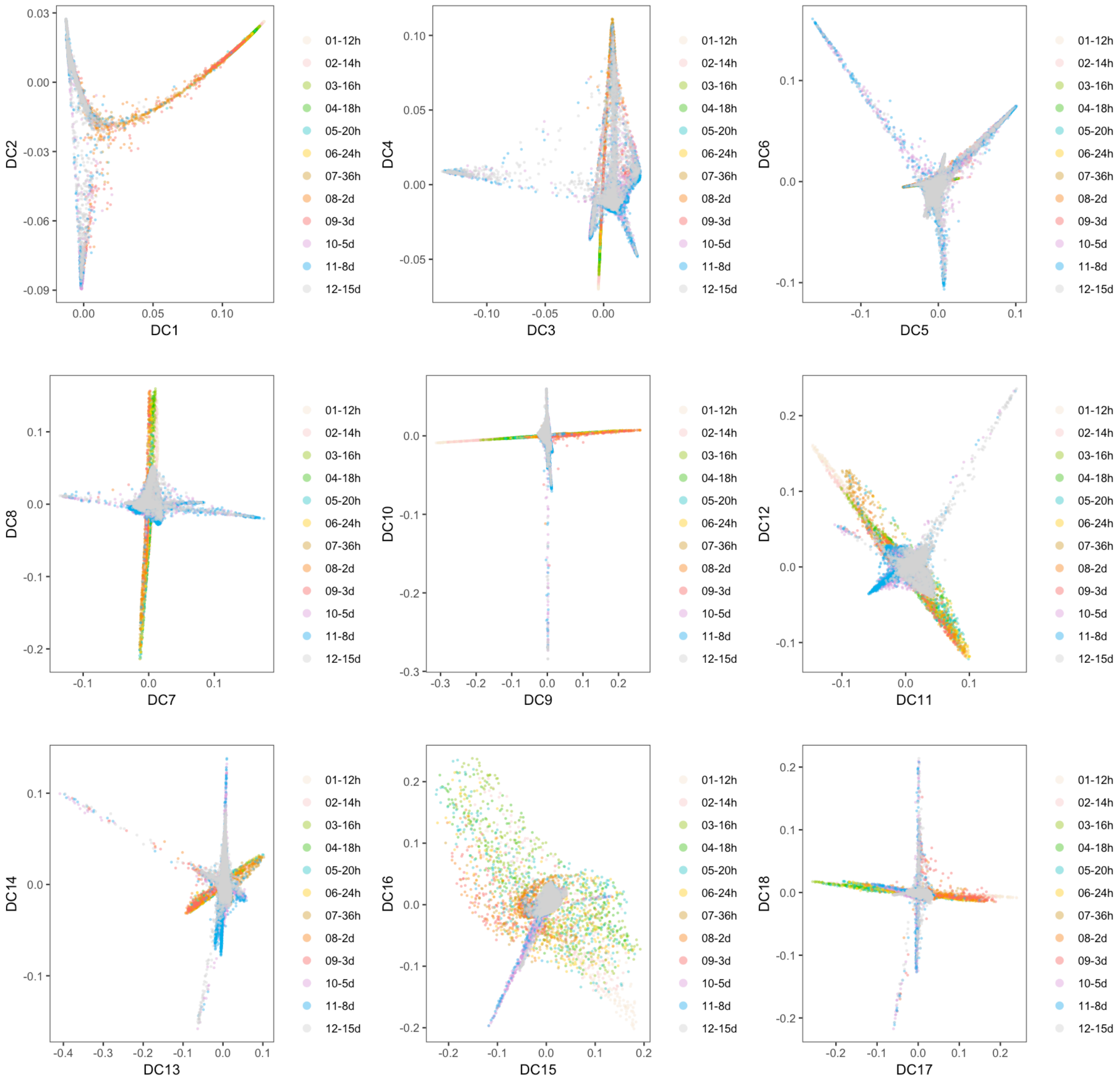

```r
# Plot with final cell types labeled
object@group.ids$final.cluster <- NA
object@group.ids[cellsInCluster(object, "stage", "12-15d"), "final.cluster"] <- object@group.ids[cellsInCluster(c
    "stage", "12-15d"), "res.5"]
plotDimArray(object = object, reduction.use = "dm", dims.to.plot = 1:18,
    label = "final.cluster", plot.title = "", outer.title = "Diffusion map with final clusters",
    legend = T, alpha = 0.6)
```

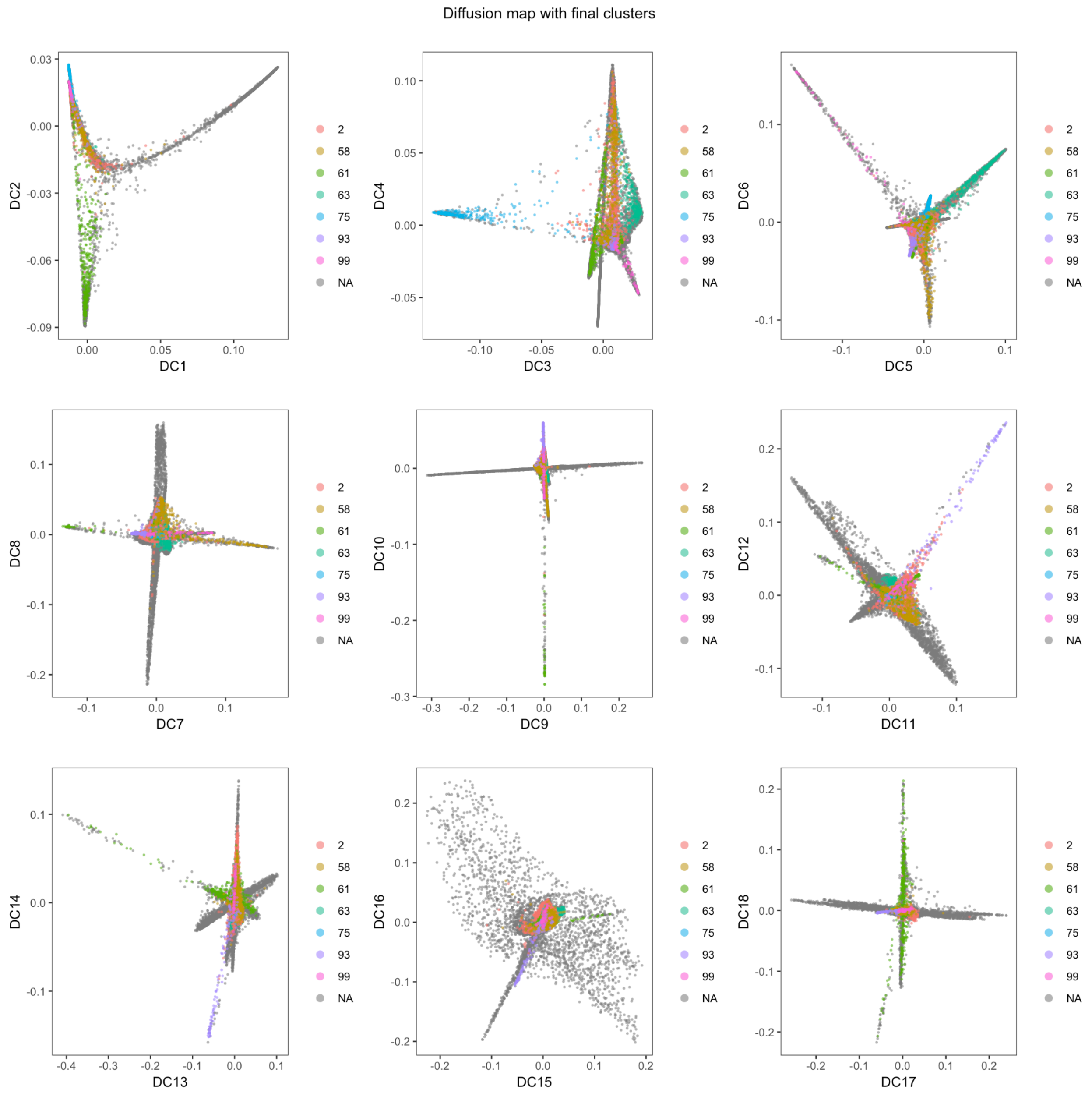

#### Calculate pseudotime

URD requires a starting point or 'root' for determining pseudotime. Here, we used all cells from the first timepoint (i.e. 12 hpf) as the root.

```
# Here, we used all cells from the first timepoint (i.e. 12 hours) as
# the root.
root.cells <- cellsInCluster(object, "stage", "01-12h")
plotDimHighlight(object, "stage", "01-12h", plot.title = "Root is 12 hpf cells")
```

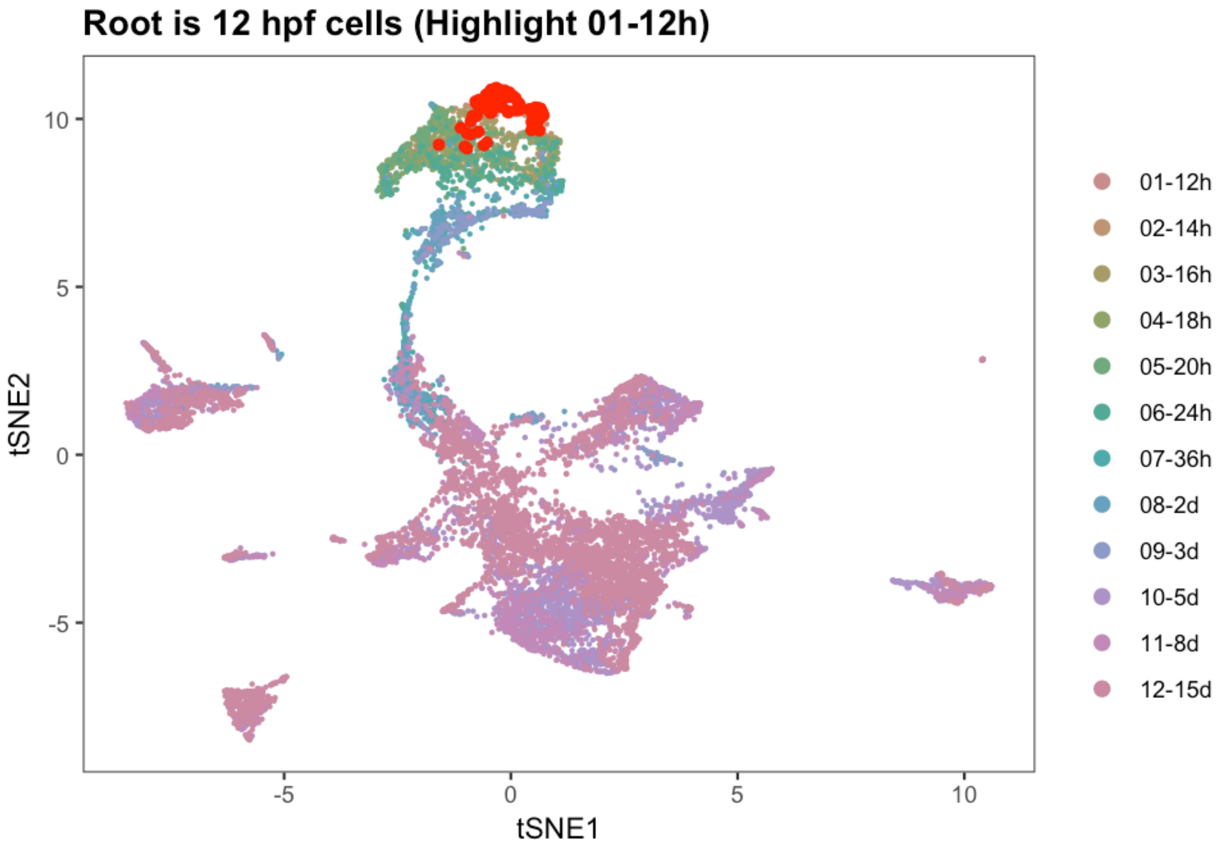

```r
# To run locally: Run graph-search simulations to determine pseudotime
flood.result <- floodPseudotime(object, root.cells = root.cells, n = 100,
    minimum.cells.flooded = 2, verbose = T)
```

```r
# Or load a pre-computed graph-search simulation result
flood.result <- readRDS(paste0(base.path, "flood/flood_hypoND_knn-100_sigma-8.rds"))
```

```r
# Process the graph-search simulations to determine the pseudotime of
# each cell
object <- floodPseudotimeProcess(object, flood.result, floods.name = "pseudotime",
    max.frac.NA = 0.4, pseudotime.fun = mean, stability.div = 10)

# If enough simulations have been run, then as additional simulations
# are added, the overall change in pseudotime of cells should reach an
# asymptote. If it does not, then floodPseudotime should be run with a
# higher n.
pseudotimePlotStabilityOverall(object)
```

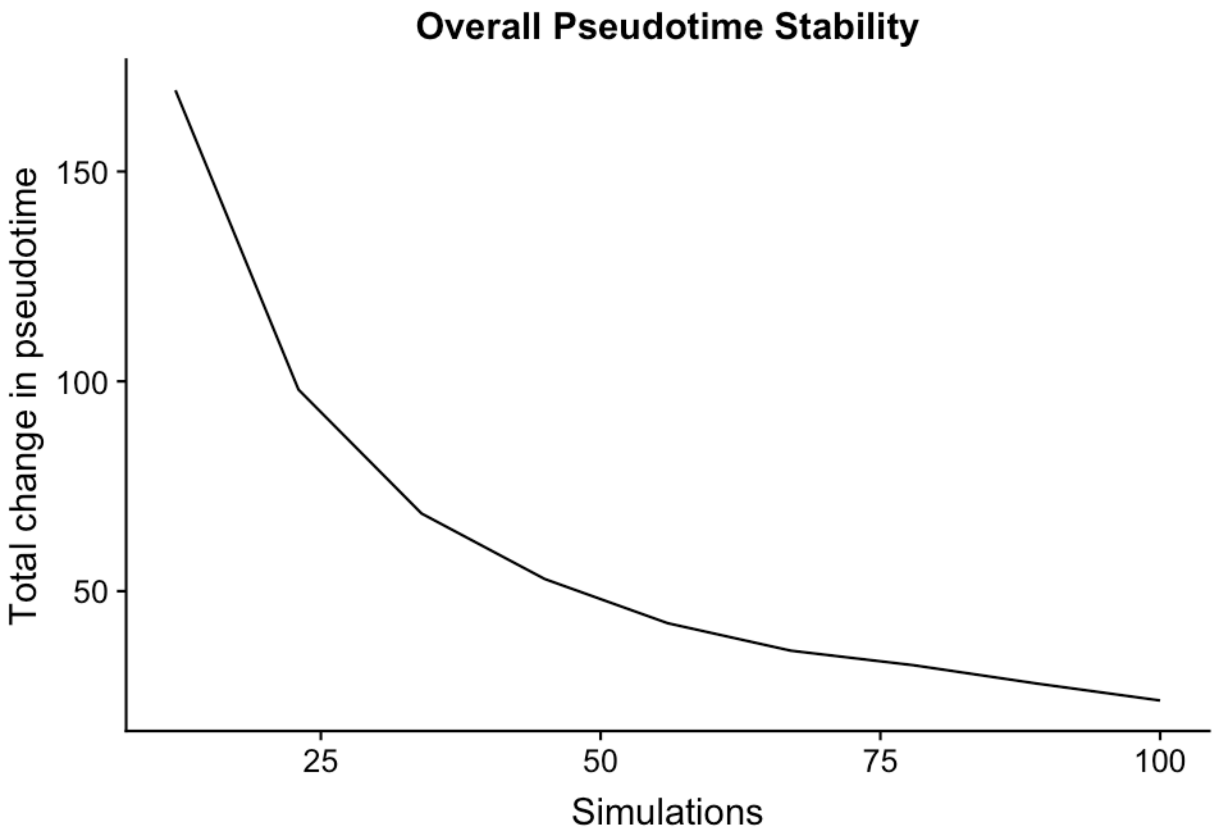

```
plotDimArray(object = object, reduction.use = "dm", dims.to.plot = 1:18,
    label = "pseudotime", plot.title = "", outer.title = "Diffusion Map labeled by pseudotime",
    legend = F, alpha = 0.4)
```

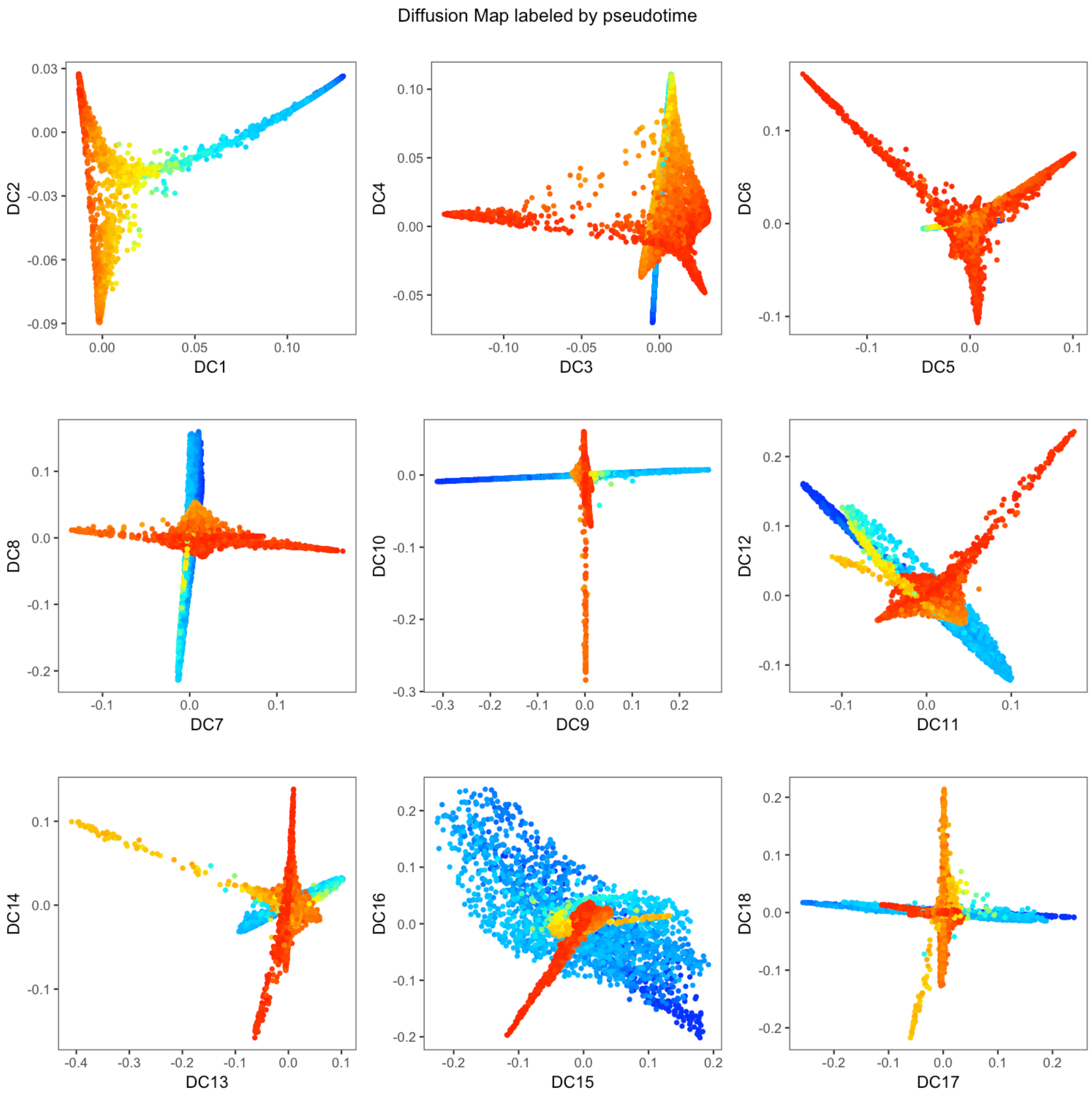

```r
plotDim(object, "pseudotime", plot.title = "UMAP projection colored by pseudotime")
```

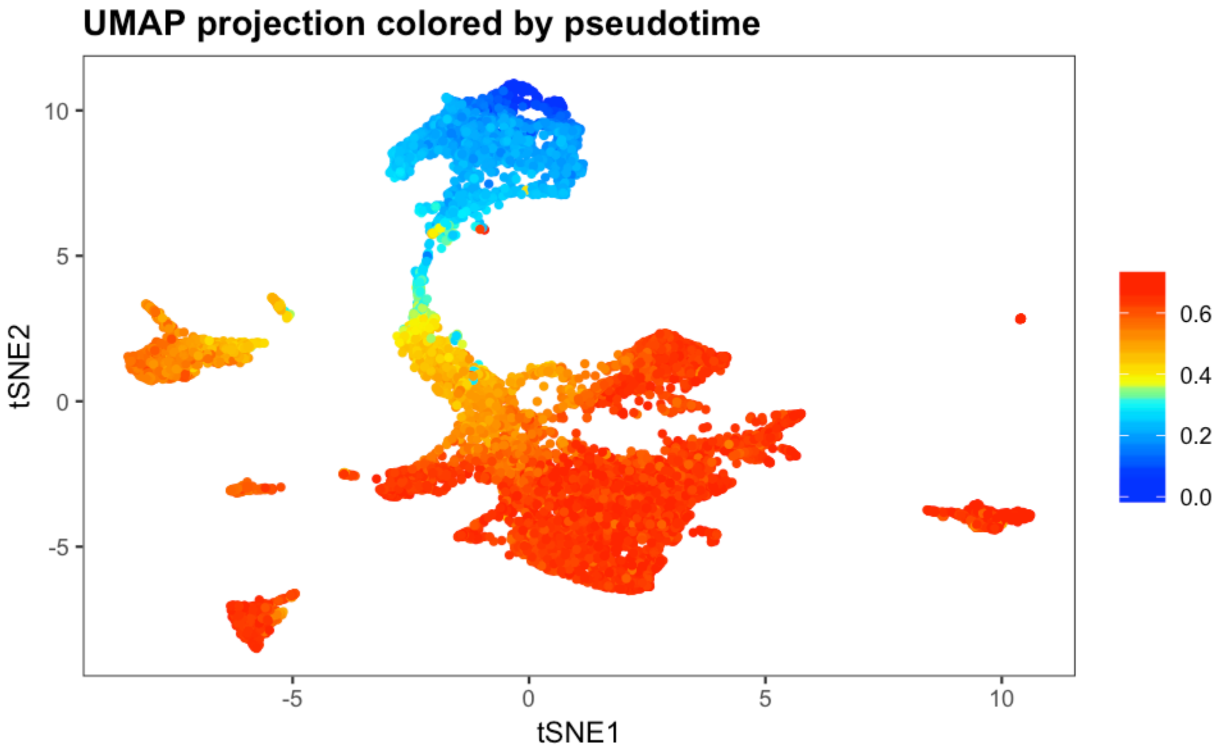

```r
plotDists(object, "pseudotime", "stage", plot.title = "Pseudotime by stage")
```

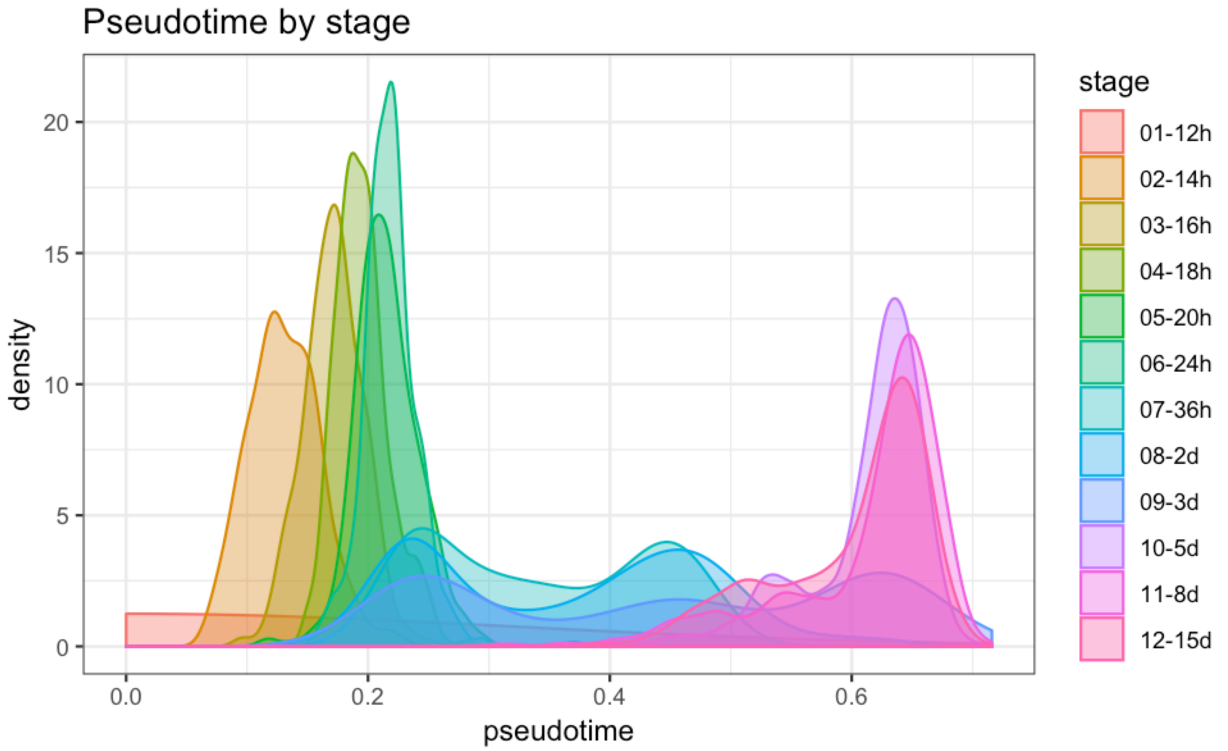

#### Calculate biased transition matrix

In order to perform biased random walks, we must first bias the transition matrix to ensure that walks proceed towards the root and do not turn into other differentiated cell types. This is performed in the cluster script **URD-TM.R**.

```
# Calculate parameters for biasing the transition matrix.
diffusion.logistic <- pseudotimeDetermineLogistic(object, "pseudotime",
    optimal.cells.forward = 40, max.cells.back = 80, pseudotime.direction = "<",
    do.plot = T, print.values = T)
```

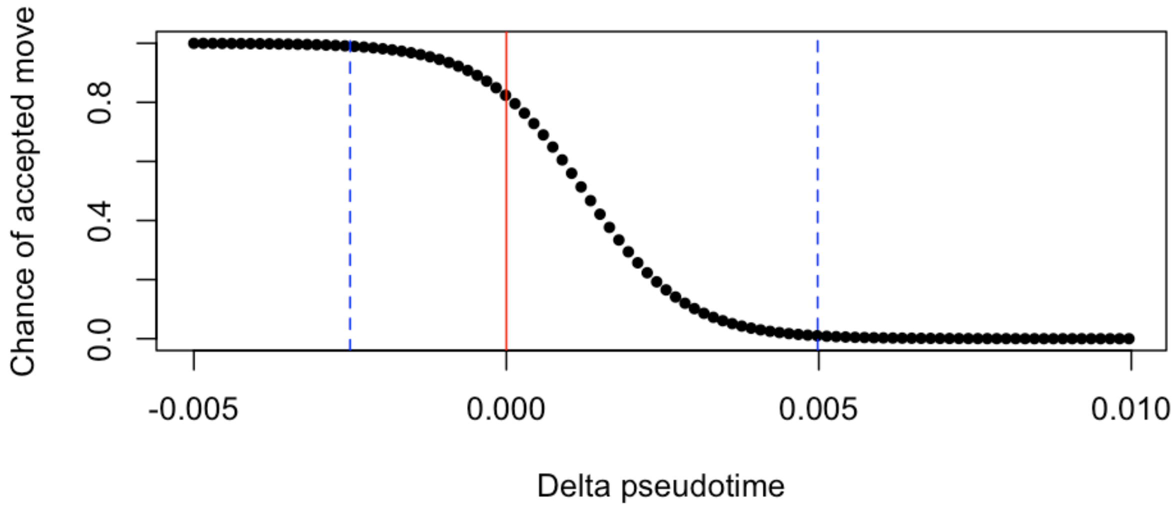

```
## [1] "Mean pseudotime back (~80 cells) 0.00498088811519173"
## [1] "Chance of accepted move to equal pseudotime is 0.821561374686937"
## [1] "Mean pseudotime forward (~40 cells) -0.00250030667341253"
```

```
# Calculate the biased matrix.
biased.tm <- pseudotimeWeightTransitionMatrix(object, pseudotime = "pseudotime",
    logistic.params = diffusion.logistic, pseudotime.direction = "<")
```

#### Perform biased random walks

Then, we perform biased walks starting from each tip. Visited cells are inferred to lie along the trajectory that connects the root to each cell type. This is performed in the cluster script **URD-Walk.R**.

##### Determine tips

We used clusters from 15 dpf as the tips for performing biased random walks. Here we define the cells belonging to each of those clusters.

```r
# All clusters at 15 days
clusters.15day <- unique(object@group.ids[grep("15d", object@group.ids$stage),
    "res.5"])
# All cells at 15 days
cells.15day <- rownames(object@group.ids)[grep("15d", object@group.ids$stage)]
# Cell lists of each cluster at 15dpf
cells.15dpf.clusters <- lapply(clusters.15day, function(clust) intersect(cells.15day,
    cellsInCluster(object, "res.5", clust)))
names(cells.15dpf.clusters) <- paste0("15d-", clusters.15day)
```

We also load a .csv file that contains information about the tips. It has four columns:

- id: Cluster ID for the tip
- use: Whether this cluster should be used when building the tree
- name: The name for this tip, which will be used on 2D plots
- short.name: The ‘short' name for this tip, which would be used on 3D plots (though we did not use that feature in this study).

```r
# Load CSV
tip.names <- read.csv(paste0(base.path, "tips/tip_names_hypoND.csv"), header = F,
    stringsAsFactors = F, colClasses = c("character", "logical", "character",
        "character"))

# Name columns and rows
names(tip.names) <- c("id", "use", "name", "short.name")
rownames(tip.names) <- gsub("_", "-", tip.names$id)

# Sort alphabetically
tip.names <- tip.names[order(rownames(tip.names)), ]
```

These are the tips that were considered during the construction of the retina URD tree (some were excluded during tree construction later in the `buildTree` command).

```r
# Define a 'tips' clustering
object@group.ids$tip <- NA
object@group.ids$tip.id <- NA
object@group.ids$tip.name <- NA

# If the tip will be used in the tree, define its cells in the
# clustering
for (i in 1:nrow(tip.names)) {
    tip.cells <- cells.15dpf.clusters[[rownames(tip.names)[i]]]
    object@group.ids[tip.cells, "tip"] <- as.character(i)
    object@group.ids[tip.cells, "tip.id"] <- rownames(tip.names)[i]
    object@group.ids[tip.cells, "tip.name"] <- as.character(tip.names[i,
        "name"])
}

# Plot the tips
plotDim(object, "tip.name")
```

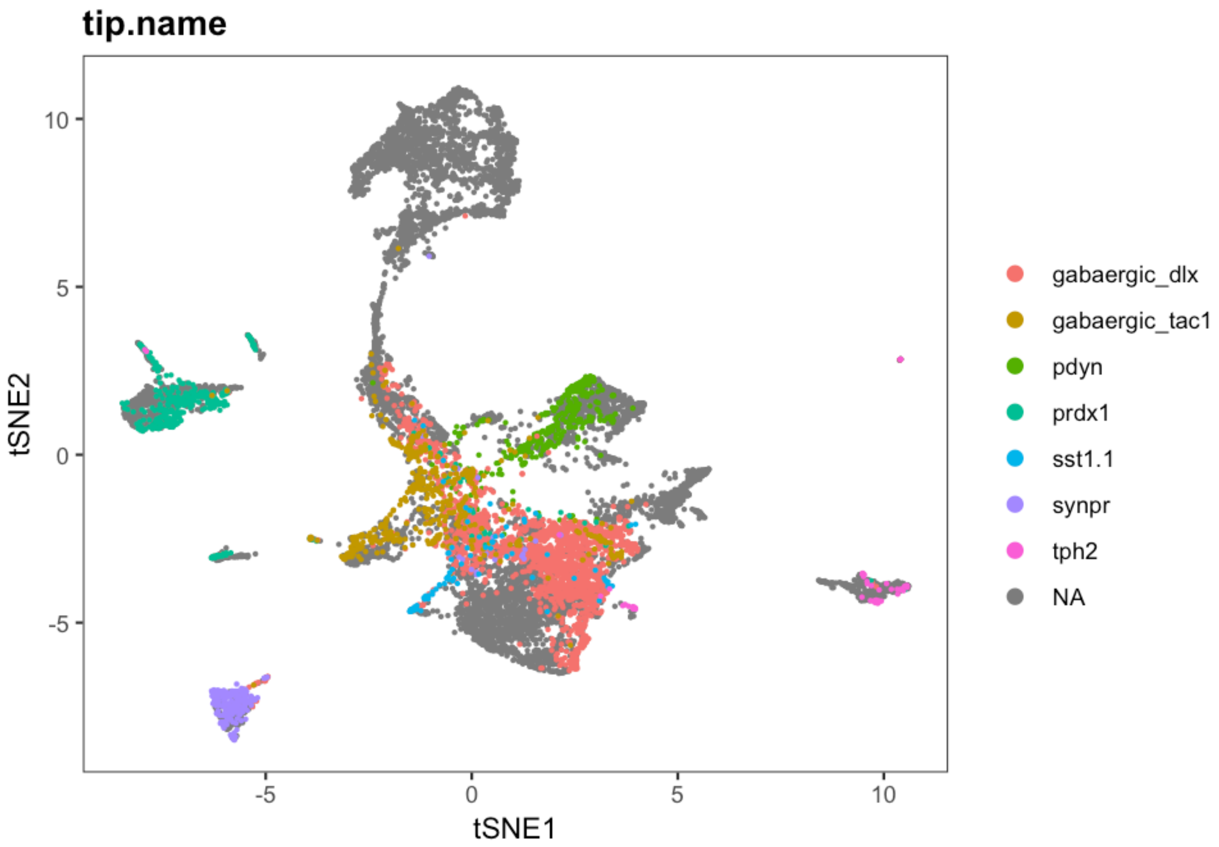

#### Perform the biased random walks

Biased random walks then need to be run starting from each tip. This can be performed on a laptop, but is an ideal candidate for parallelization on a cluster. (The walks from each tip can be run as a separate job.)

```
## IF RUNNING LOCALLY

# Loop through each cluster
walks <- lapply(rownames(tip.names), function(c) {
    # Exclude any tip cells that for whatever reason didn't end up in the
    # biased TM (e.g. maybe not assigned a pseudotime).
    tip.cells <- intersect(cells.15dpf.clusters[[c]], rownames(biased.tm))
    # Perform the random walk simulation
    this.walk <- simulateRandomWalk(start.cells = tip.cells, transition.matrix = biased.tm,
        end.cells = root.cells, n = 50000, end.visits = 1, verbose.freq = 1000,
        max.steps = 5000)
    return(this.walk)
})
names(walks) <- rownames(tip.names)

# Alternatively, this loop is automated by the function
# simulateRandomWalksFromTips
```

Alternatively, a set of pre-calculated walks can be loaded. Since the walks are a simulation (and therefore not deterministic), this is particularly crucial for reproducing results.

```
## IF LOADING PRE-CALCULATED WALKS

# Get list of files in the walks directory
walks.files <- list.files(paste0(base.path, "/walks/hypoND/"), pattern = ".rds")

# Load the walks previously performed for each cluster
walks <- lapply(rownames(tip.names), function(c) {
    walk.file <- grep(pattern = paste0("_tip-", c, "_"), x = walks.files,
        value = T)[1]
    return(readRDS(paste0(base.path, "/walks/hypoND/", walk.file)))
})
names(walks) <- rownames(tip.names)
```

#### Process the random walks

The walks are then converted to visitation frequency by importing them into the URD object.

```
for (i in 1:nrow(tip.names)) {
    # Load the individual walk visitation frequencies into the object
    object <- processRandomWalks(object, walks = walks[[i]], walks.name = i,
        n.subsample = 1, verbose = F)
}
```

### Build the URD tree

Then, a branching tree is constructed, by joining trajectories in an agglomerative fashion when cells are highly visited by walks from multiple tips. The following steps were performed in the cluster script URD-Tree.R.

```
# Tree building is destructive, so create a copy of the object
object.tree <- object

# Load tip cells
object.tree <- loadTipCells(object.tree, "tip")

# Determine tips to use
tips.to.use <- which(tip.names$use)
```

```
# Build the tree
object.tree <- buildTree(object.tree, pseudotime = "pseudotime", divergence.method = "ks",
    cells.per.pseudotime.bin = 40, bins.per.pseudotime.window = 5, save.all.breakpoint.info = T,
    p.thresh = 1e-04, verbose = F, tips.use = as.character(tips.to.use))
```

```
# Name the tips of the tree
object.tree <- nameSegments(object.tree, segments = tips.to.use, segment.names = as.character(tip.names[tips.to.u[illegible]
    "name"], short.names = as.character(tip.names[tips.to.use, "short.name"])))
```

```
plotTree(object.tree, "stage", discrete.colors = stage.colors, label.segments = T)
```

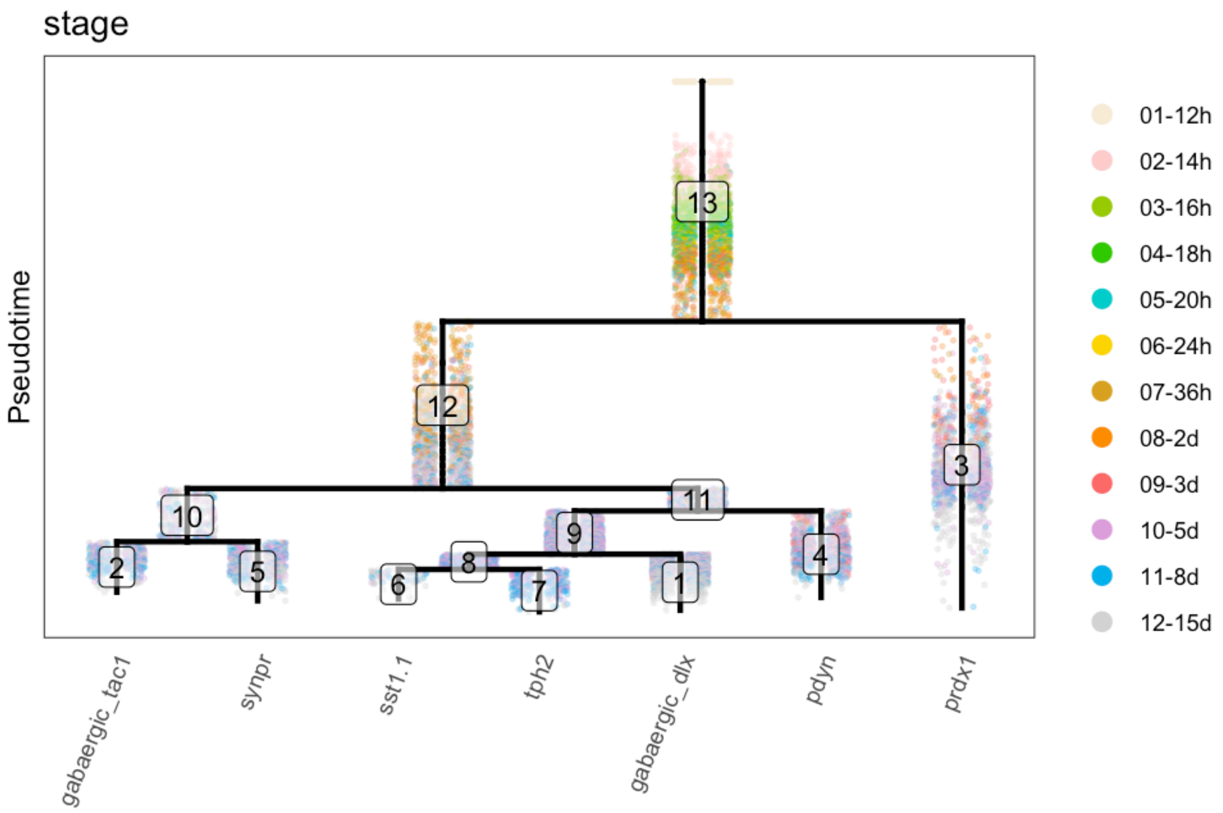

#### Save the URD tree

The tree is then saved for use in downstream analysis, and can easily be loaded for further perusal.

```
saveRDS(object.tree, file = paste0(base.path, "tree/URD-Tree-Hypo.rds"))
```

### Hypothalamus: 3 - URD Cascades and Figures

*Jeff Farrell*

*10/08/2019*

### Load data

```r
# Load URD
library(URD)
```

```
## Loading required package: ggplot2
```

```
## Warning: package 'ggplot2' was built under R version 3.4.4
```

```
## Loading required package: Matrix
```

```
## Warning: package 'Matrix' was built under R version 3.4.4
```

```r
# Basic location
base.path <- "~/urd-cluster-bushra/"

# Load completed hypothalamus tree object
obj.path <- paste0(base.path, "tree/hypoND/tree-hypoND_knn-100_sigma-8_40F-80B_NO-_ks_0001.rds")
obj <- readRDS(obj.path)
```

### Plot gene expression on the tree

#### Plot tree by stage

```r
stage.colors <- c("antiquewhite", "#FFCCCC", "#99CC00", "#33CC00", "cyan3",
    "gold", "goldenrod", "darkorange", "indianred1", "plum", "deepskyblue2",
    "lightgrey")

plotTree(obj, "stage", label.type = "group", discrete.colors = stage.colors)
```

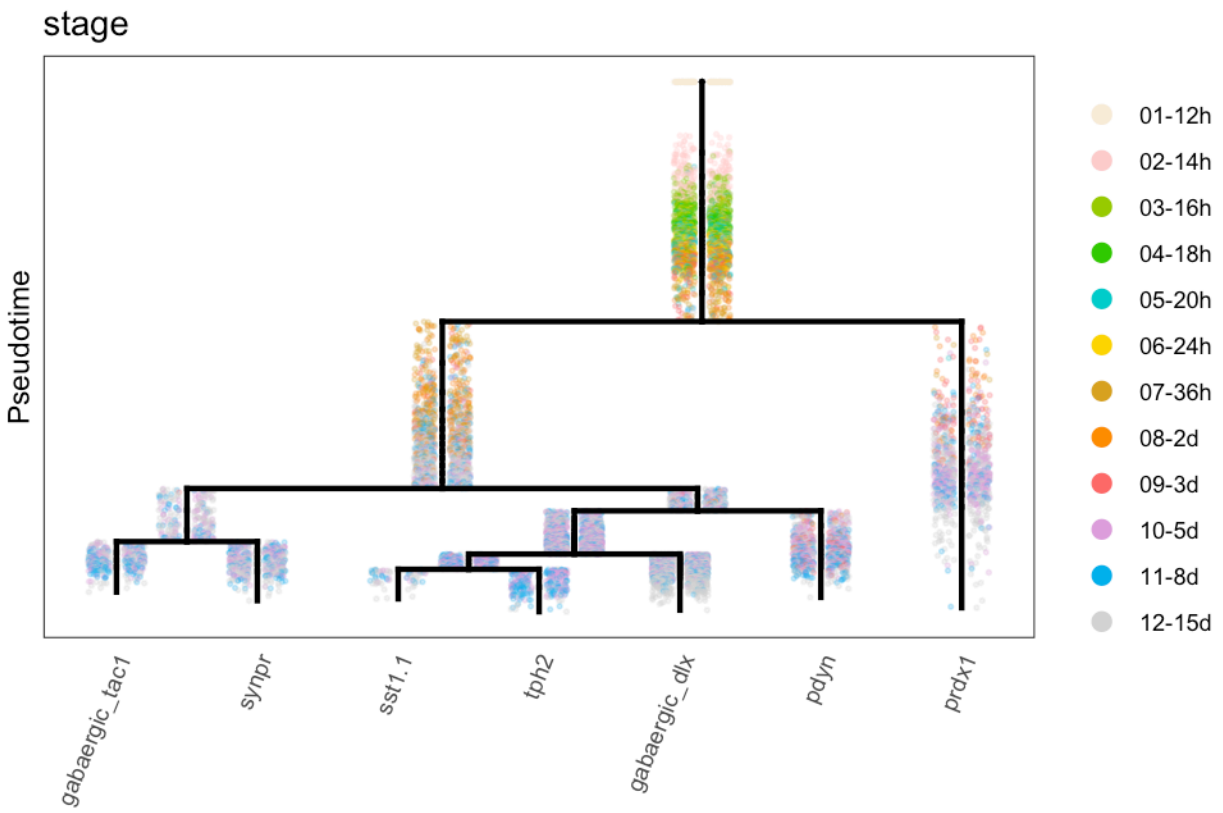

#### Plot tree with gene expression: main figures

```
gridExtra::grid.arrange(grobs = lapply(c("shha", "pdyn", "rx3", "nrgna"),
    plotTree, object = obj, label.x = F, plot.cells = F), ncol = 2)
```

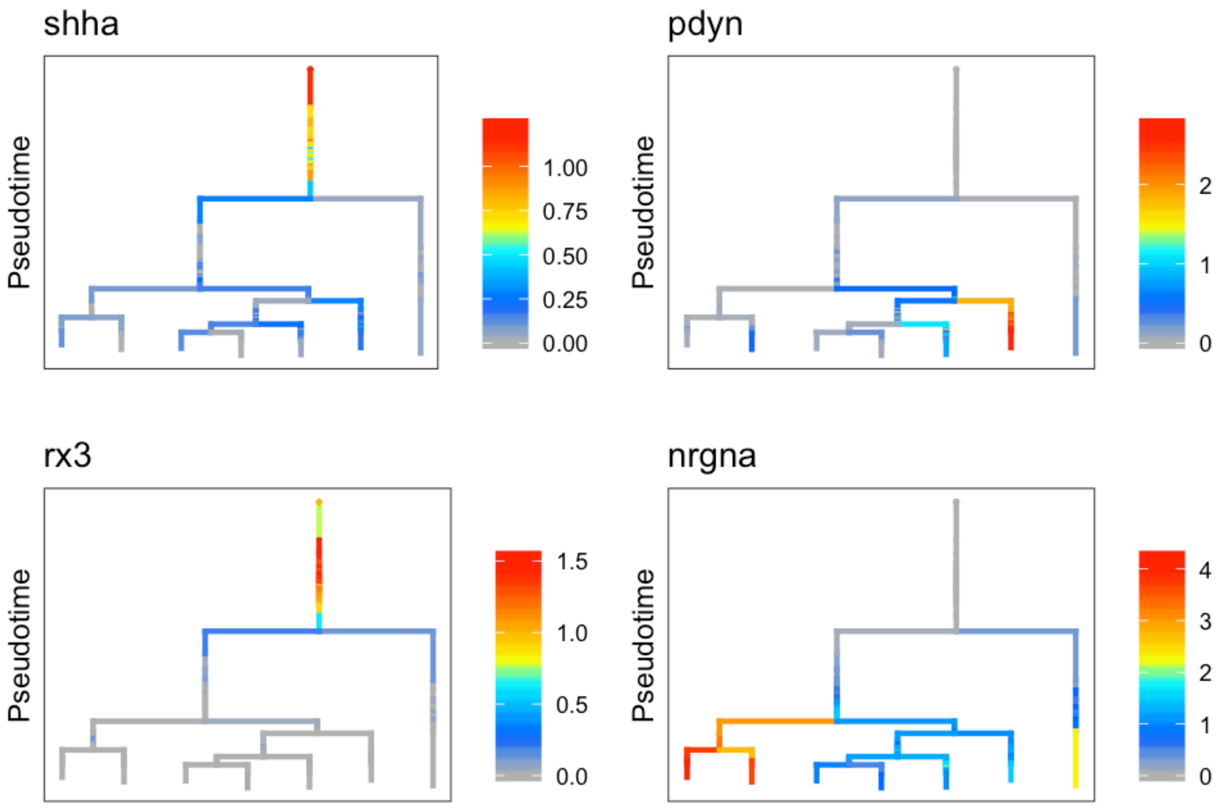

```
gridExtra::grid.arrange(grobs = lapply(c("dlx5a", "dlx6a", "vax1"), plotTree,
    object = obj, label.x = F, plot.cells = F), ncol = 2)
```

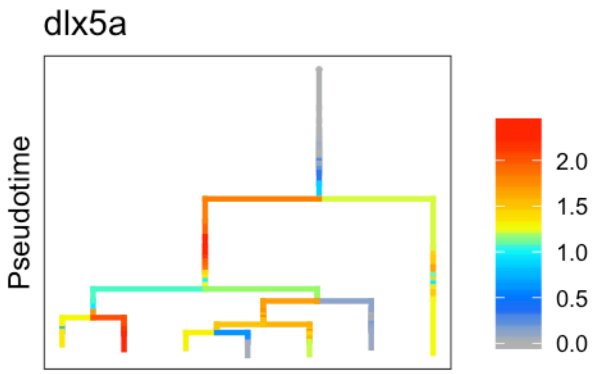

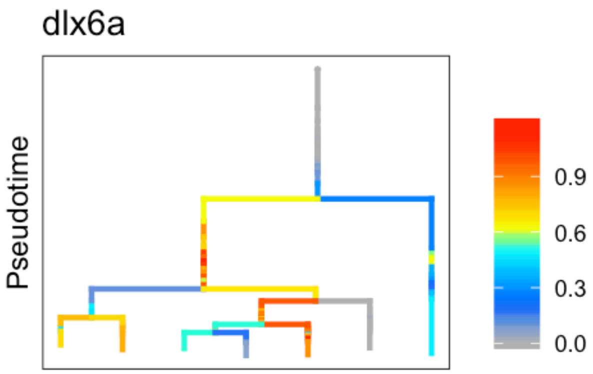

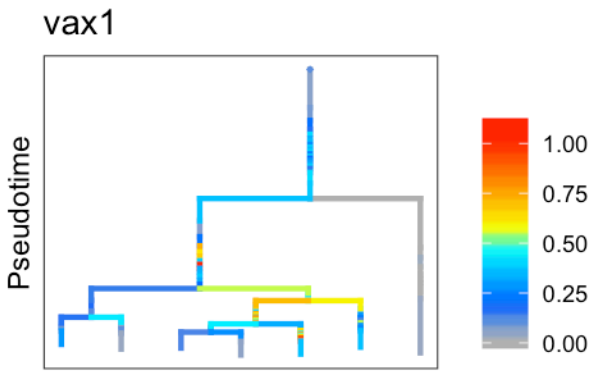

#### Plot tree with gene expression: supplemental figures

```
gridExtra::grid.arrange(grobs = lapply(c("nkx2.4b", "ascl1a", "insm1a",
    "tubb5", "scg2b", "dlx2a", "nkx2.4a", "nrgna", "tac1", "synpr", "sp8a",
    "gad1b", "npy", "sst1.1", "tph2", "fezf1", "pdyn", "slc17a6b", "prdx1",
    "pou3f1"), plotTree, object = obj, label.x = F, plot.cells = F), ncol = 4)
```

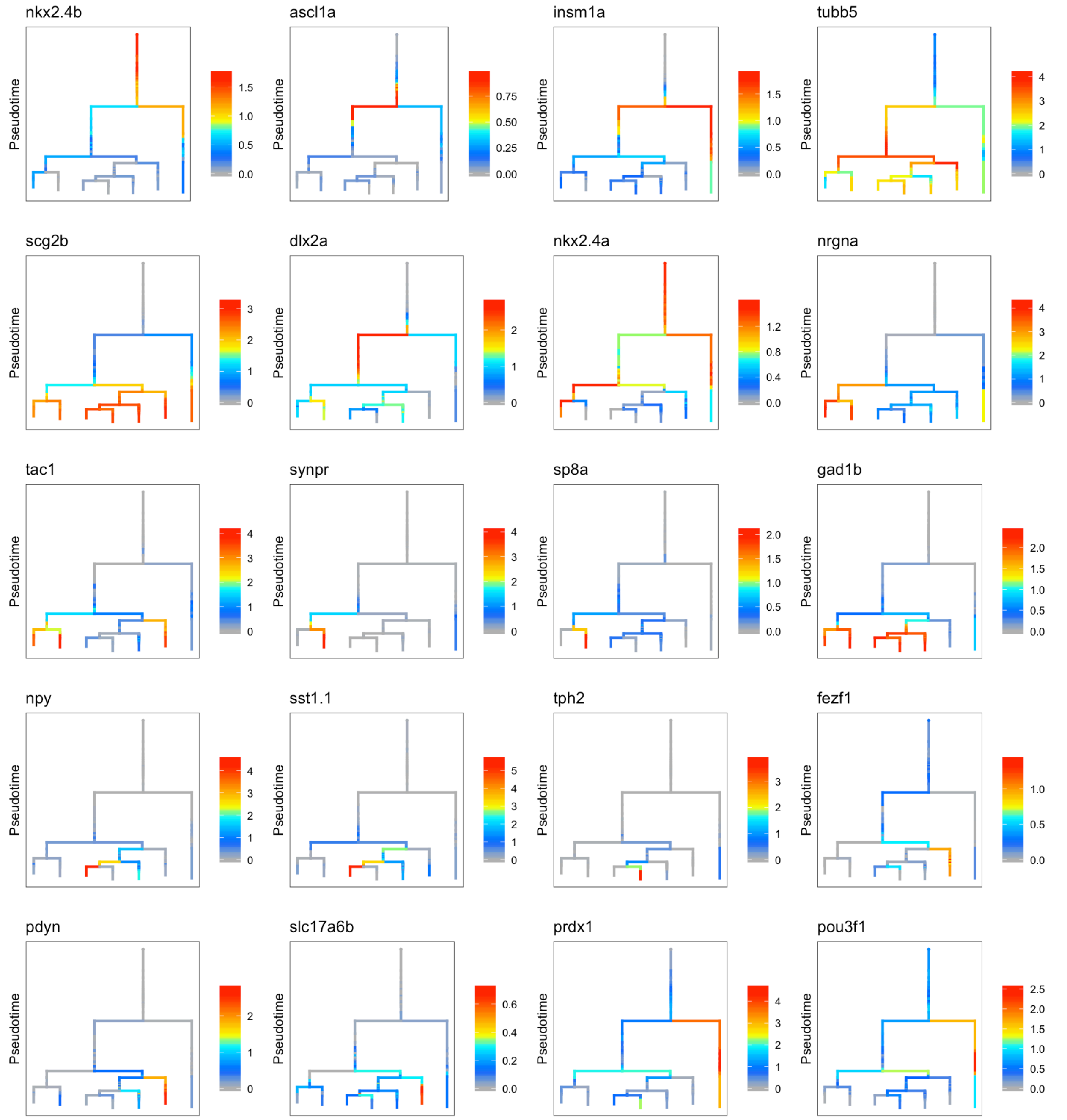

### Determine genes enriched in trajectories to particular cell types

#### Comparison between major cell types

We took each major group ("clade") of branches from the end of the tree as a single entity (i.e. prdx1+ neurons, pdyn+ neurons, GABAergic dlx+ neurons, nrgna+ neurons) and compared them against each other to look for differentially expressed genes.

```r
# Get the parent segment of each clade to consider as a group
combined.tips <- c("3", "4", "9", "10")

# Get the cells in that segment and all child segments
cells.combined.tips <- lapply(combined.tips, function(t) whichCells(obj,
    label = "segment", value = segChildrenAll(obj, t, include.self = T)))
names(cells.combined.tips) <- combined.tips

# Loop through each of these clades and look for differentially
# expressed genes
combined.markers <- lapply(combined.tips, function(tip) {
    # Find all of the other clades
    opposing.tips <- setdiff(combined.tips, tip)
    # Perform pairwise comparisons to each other clade
    m.o <- lapply(opposing.tips, function(tip.opposing) {
        # message(paste0(Sys.time(), ': Comparing tip ', tip, ' to ',
        # tip.opposing, '.')) Find differentially expressed genes between the
        # pair of clades
        ma <- markersAUCPR(object = obj, cells.1 = cells.combined.tips[[tip]],
            cells.2 = cells.combined.tips[[tip.opposing]], effect.size = 0.5,
            auc.factor = 1.1)
        # In order to facilitate combining all of the results later, add
        # columns about which two clades were compared and also a duplicate
        # entry of the name of each gene that's recovered.
        if (nrow(ma) > 0) {
            ma$gene <- rownames(ma)
            ma$tip1 <- tip
            ma$tip2 <- tip.opposing
        }
        return(ma)
    })
    names(m.o) <- opposing.tips
    return(m.o)
})
names(combined.markers) <- combined.tips

# Require that genes are markers against at least 2 other clades
combined.markers.beatmult <- lapply(combined.markers, function(m) {
    names(which(table(unlist(lapply(m, rownames))) >= 2))
})

# Since genes might be a marker in a comparison to several other
# clades, combine the results into a single table, where each gene is
# listed only once with the info from the pairwise comparison where it
# had the strongest differential expression.
combined.markers.best <- lapply(1:length(combined.markers.beatmult), function(i) {
    cm <- do.call("rbind", combined.markers[[i]])
    cm <- cm[cm$gene %in% combined.markers.beatmult[[i]], ]
    cmb <- do.call("rbind", lapply(combined.markers.beatmult[[i]], function(g) {
        cmr <- cm[cm$gene == g, ]
        return(cmr[which.max(cmr$AUCPR.ratio), ])
    }))
    rownames(cmb) <- cmb$gene
```

```
    if (!is.null(cmb)) {
        cmb <- cmb[order(cmb$AUCPR.ratio, decreasing = T), ]
        cmb$exp.global <- apply(obj@logupx.data[rownames(cmb), unlist(obj@tree$cells.in.segment)],
            1, mean.of.logs)
        cmb$exp.global.fc <- cmb$nTrans_1 - cmb$exp.global
    }
    return(cmb)
})
names(combined.markers.best) <- combined.tips
```

#### AUCPR along tree

We also used the AUCPRTestAlongTree function to ask for genes that are differential markers of a lineage using URD's tree structure. This makes a comparison at each branchpoint from a particular cell type up to the root.

```
# Get all of the tips from the tree
tips.in.tree <- as.character(obj@tree$tips)

# Tree segments to use as root
roots <- rep("12", length(tips.in.tree))
names(roots) <- tips.in.tree
roots["3"] <- "13"

# Define parameters to use for calculation Used more permissive values
# in the sst1.1+ / tph2+ / gabaergic dlx+ neuronal comparisons due to
# the small number of cells in these populations
auc.use <- rep(1.2, length(tips.in.tree))
names(auc.use) <- tips.in.tree
auc.use[c("1", "6", "7")] <- 1.15
log.effect.use <- rep(0.8, length(tips.in.tree))
names(log.effect.use) <- tips.in.tree
log.effect.use[c("1", "6", "7")] <- 0.6

# Perform a loop of tests with each tip.
markers <- lapply(tips.in.tree, function(t) {
    this.root <- roots[t]
    this.auc <- auc.use[t]
    this.log <- log.effect.use[t]
    # message(paste0(Sys.time(), ': Starting tip ', t, ' and root ',
    # this.root, ' with params ', this.auc, ' AUC and ', this.log, ' effect
    # size.'))
    these.markers <- aucprTestAlongTree(obj, pseudotime = "pseudotime",
        tips = as.character(t), genes.use = NULL, must.beat.sibs = 0.6,
        report.debug = F, root = this.root, auc.factor = this.auc, log.effect.size = this.log)
    these.markers$gene <- rownames(these.markers)
    these.markers$tip <- t
    return(these.markers)
})
names(markers) <- tips.in.tree
```

#### Markers of the prdx1- neuron clade

```
# Calculate from segment 12 against segment 3 specifically
nonprdx.markers <- markersAUCPR(obj, clust.1 = "12", clust.2 = "3", clustering = "segment",
    effect.size = 0.8, auc.factor = 1.2)
# Also look at segment 12 vs. rest of the hypothalamus with lower
# thresholds
nonprdx.markers.global <- markersAUCPR(obj, clust.1 = "12", clust.2 = as.character(c(1:11,
    13)), clustering = "segment", effect.size = 0.4, auc.factor = 1.1)
```

```
## Warning in names(genes.data)[4:7] <- paste(c("posFrac", "posFrac",
## "nTrans", : number of items to replace is not a multiple of replacement
## length
```

### Functions for curating differential expression results

We further curated those differentially expressed genes using the following functions:

#### threshold.tree.markers

Function to threshold markers from a markersAUCPRAlongTree test with additional criteria

- markers: list of results from markersAUCPRAlongTree tests
- tip: which tip (or element of the list to pursue)
- global.fc: fold.change that gene must have along the trajectory pursued vs. rest of the data
- aucpr.ratio.all: classifier score that gene must exhibit along trajectory test vs. rest of the data
- branch.fc: fold.change that gene must have (in best case) vs. the opposing branch at any branchpoint along the trajectory.
- Returns `markers` with only a subset of rows retained.

```
threshold.tree.markers <- function(markers, tip, global.fc = 0.1, branch.fc = 0.4,
    aucpr.ratio.all = 1.03) {
    m <- markers[[tip]]
    # First off -- lose global FC < x
    bye.globalfc <- rownames(m)[m$expfc.all < global.fc]
    # Second -- get rid of branch FC < x
    bye.branchfc <- rownames(m)[m$expfc.maxBranch < branch.fc]
    # Third -- get rid of stuff essentially worse than random
    # classification on global level
    bye.badglobalaucpr <- rownames(m)[m$AUCPR.ratio.all < aucpr.ratio.all]
    bye.all <- unique(c(bye.globalfc, bye.branchfc, bye.badglobalaucpr))
    m.return <- m[setdiff(rownames(m), bye.all), ]
    return(m.return)
}
```

#### divide.branches

Function to compare genes between two branches. Use this on a compiled list of markers to do a final selection of genes that are specific to one branch or another or markers of both (i.e. when making photoreceptor heatmap, use to divide into photoreceptor, cone, and rod markers)

- **object**: An URD object
- **genes**: (Character vector) Genes to test
- **clust.1**: (Character) Cluster 1
- **clust.2**: (Character) Cluster 2
- **clustering**: (Character) Clustering to pull from
- **exp.fc**: (Numeric) Minimum expression fold-change between branches to consider different
- **exp.thresh**: (Numeric) Minimum fraction of cells in order to consider gene expressed in a branch
- **exp.diff**: (Numeric) Minimum difference in fraction of cells expressing to consider gene differential
- Returns list of gene names ("specific.1" = specific to clust.1, "specific.2" = specific to clust.2, "markers" = all genes tested)

```
divide.branches <- function(object, genes, clust.1, clust.2, clustering = "segment",
    exp.fc = 0.4, exp.thresh = 0.1, exp.diff = 0.1) {
    # Double check which markers are unique to one or the other population
    mcomp <- markersAUCPR(object, clust.1 = clust.1, clust.2 = clust.2,
        clustering = clustering, effect.size = -Inf, auc.factor = 0, genes.use = genes,
        frac.min.diff = 0, frac.must.express = 0)
    specific.b <- rownames(mcomp)[abs(mcomp$exp.fc) > exp.fc & mcomp[,
        4] < exp.thresh & mcomp[, 5] > pmin((mcomp[, 4] + exp.diff), 1)]
    specific.a <- rownames(mcomp)[abs(mcomp$exp.fc) > exp.fc & mcomp[,
        5] < exp.thresh & mcomp[, 4] > pmin((mcomp[, 5] + exp.diff), 1)]
    r <- list(specific.a, specific.b, mcomp)
    names(r) <- c("specific.1", "specific.2", "markers")
    return(r)
}
```

#### divide.branches.triple

Function to compare genes between three branches. Use this on a compiled list of markers to do a final selection of genes that are specific to one branch or another or markers of both (i.e. when making gad2+ heatmap, use to divide into general, dlx+, sst+, and tph2+ markers).

- **object**: An URD object
- **genes**: (Character vector) Genes to test
- **clust.1**: (Character) Cluster 1
- **clust.2**: (Character) Cluster 2
- **clust.3**: (Character) Cluster 3
- **clustering**: (Character) Clustering to pull from
- **exp.fc**: (Numeric) Minimum expression fold-change between branches to consider different
- **exp.thresh**: (Numeric) Minimum fraction of cells in order to consider gene expressed in a branch
- **exp.diff**: (Numeric) Minimum difference in fraction of cells expressing to consider gene differential
- Returns list of gene names ("nonspecific" = genes not in specific.1/2/3, "specific.1" = specific to clust.1, "specific.2" = specific to clust.2, "specific.3" = specific to clust.3, each pairwise comparison, and "markers" = all genes tested)

```
divide.branches.triple <- function(object, genes, clust.1, clust.2, clust.3,
    clustering = "segment", exp.fc = 0.4, exp.thresh = 0.1, exp.diff = 0.1) {
    # Double check which markers are unique to one or the other population
    mcomp12 <- markersAUCPR(object, clust.1 = clust.1, clust.2 = clust.2,
        clustering = clustering, effect.size = -Inf, auc.factor = 0, genes.use = genes,
```

```r
        frac.min.diff = 0, frac.must.express = 0)
    mcomp23 <- markersAUCPR(object, clust.1 = clust.2, clust.2 = clust.3,
        clustering = clustering, effect.size = -Inf, auc.factor = 0, genes.use = genes,
        frac.min.diff = 0, frac.must.express = 0)
    mcomp13 <- markersAUCPR(object, clust.1 = clust.1, clust.2 = clust.3,
        clustering = clustering, effect.size = -Inf, auc.factor = 0, genes.use = genes,
        frac.min.diff = 0, frac.must.express = 0)

    specific.1v2 <- rownames(mcomp12)[abs(mcomp12$exp.fc) > exp.fc & mcomp12[,
        5] < exp.thresh & mcomp12[, 4] > pmin((mcomp12[, 5] + exp.diff),
        1)]
    specific.2v1 <- rownames(mcomp12)[abs(mcomp12$exp.fc) > exp.fc & mcomp12[,
        4] < exp.thresh & mcomp12[, 5] > pmin((mcomp12[, 4] + exp.diff),
        1)]

    specific.2v3 <- rownames(mcomp23)[abs(mcomp23$exp.fc) > exp.fc & mcomp23[,
        5] < exp.thresh & mcomp23[, 4] > pmin((mcomp23[, 5] + exp.diff),
        1)]
    specific.3v2 <- rownames(mcomp23)[abs(mcomp23$exp.fc) > exp.fc & mcomp23[,
        4] < exp.thresh & mcomp23[, 5] > pmin((mcomp23[, 4] + exp.diff),
        1)]

    specific.1v3 <- rownames(mcomp13)[abs(mcomp13$exp.fc) > exp.fc & mcomp13[,
        5] < exp.thresh & mcomp13[, 4] > pmin((mcomp13[, 5] + exp.diff),
        1)]
    specific.3v1 <- rownames(mcomp13)[abs(mcomp13$exp.fc) > exp.fc & mcomp13[,
        4] < exp.thresh & mcomp13[, 5] > pmin((mcomp13[, 4] + exp.diff),
        1)]

    specific.1 <- unique(setdiff(c(specific.1v2, specific.1v3), c(specific.2v3,
        specific.3v2)))
    specific.2 <- unique(setdiff(c(specific.2v1, specific.2v3), c(specific.1v3,
        specific.3v1)))
    specific.3 <- unique(setdiff(c(specific.3v2, specific.3v1), c(specific.2v1,
        specific.1v2)))
    nonspecific <- setdiff(genes, c(specific.1, specific.2, specific.3))

    markers.comp <- list(mcomp12, mcomp13, mcomp23)
    names(markers.comp) <- c("1v2", "1v3", "2v3")

    r <- list(nonspecific, specific.1, specific.2, specific.3, specific.1v2,
        specific.1v3, specific.2v1, specific.2v3, specific.3v1, specific.3v2,
        markers.comp)
    names(r) <- c("nonspecific", "specific.1", "specific.2", "specific.3",
        "specific.1v2", "specific.1v3", "specific.2v1", "specific.2v3",
        "specific.3v1", "specific.3v2", "markers")
    return(r)
}
```

### Functions for heatmap generation

These functions were used in the production of heatmaps:

#### Color scale

Generate color scale to use with heatmaps.

```
cols <- (scales::gradient_n_pal(RColorBrewer::brewer.pal(9, "YlOrRd")))(seq(0,
    1, length.out = 50))
```

#### determine.timing

**Determines order to plot genes in heatmap.** "Expression" is defined as 20% higher expression than the minimum observed value. "Peak" expression is defined as 50% higher expression than minimum observed value. The two longest stretches of "peak" expression are found, and then the later one is used. The onset time of the stretch of expression that contains that peak is also determined. Genes are then ordered by the pseudotime at which they enter "peak" expression, leave "peak" expression, start "expression", and leave "expression".

- **s**: result from `geneSmoothFit`
- **genes**: genes to order; default is all genes that were fit.
- Returns `s` but with an additional list entry (`$timing`) of the order to plot genes

```
determine.timing <- function(s, genes = rownames(s$mean.expression)) {
    s$timing <- as.data.frame(do.call("rbind", lapply(genes, function(g) {
        sv <- as.numeric(s$scaled.smooth[g, ])
        pt <- as.numeric(colnames(s$scaled.smooth))
        # Figure out baseline expression & threshold for finding peaks
        min.val <- max(min(sv), 0)
        peak.val <- ((1 - min.val)/2) + min.val
        exp.val <- ((1 - min.val)/5) + min.val
        # Run-length encoding of above/below the peak-threshold
        peak.rle <- rle(sv >= peak.val)
        peak.rle <- data.frame(lengths = peak.rle$lengths, values = peak.rle$values)
        peak.rle$end <- cumsum(peak.rle$lengths)
        peak.rle$start <- head(c(0, peak.rle$end) + 1, -1)
        # Run-length encoding of above/below the expressed-threshold
        exp.rle <- rle(sv >= exp.val)
        exp.rle <- data.frame(lengths = exp.rle$lengths, values = exp.rle$values)
        exp.rle$end <- cumsum(exp.rle$lengths)
        exp.rle$start <- head(c(0, exp.rle$end) + 1, -1)
        # Take top-two longest peak RLE & select later one.  Find stretches
        # that are above peak value
        peak <- which(peak.rle$values)
        # Order by length and take 1 or 2 longest ones
        peak <- peak[order(peak.rle[peak, "lengths"], decreasing = T)][1:min(2,
            length(peak))]
        # Order by start and take latest one.
        peak <- peak[order(peak.rle[peak, "start"], decreasing = T)][1]
        # Identify the actual peak value within that stretch
        peak <- which.max(sv[peak.rle[peak, "start"]:peak.rle[peak, "end"]]) +
```

```
            peak.rle[peak, "start"] - 1
        # Identify the start and stop of the expressed stretch that contains
        # the peak
        exp.start <- exp.rle[which(exp.rle$end >= peak & exp.rle$start <=
            peak), "start"]
        exp.end <- exp.rle[which(exp.rle$end >= peak & exp.rle$start <=
            peak), "end"]
        # Identify values of expression at start and stop
        smooth.start <- sv[exp.start]
        smooth.end <- sv[exp.end]
        # Convert to pseudotime?
        exp.start <- pt[exp.start]
        exp.end <- pt[exp.end]
        peak <- pt[peak]
        # Return a vector
        v <- c(exp.start, peak, exp.end, smooth.start, smooth.end)
        names(v) <- c("pt.start", "pt.peak", "pt.end", "exp.start", "exp.end")
        return(v)
    })))
    rownames(s$timing) <- genes

    # Decide on ordering of genes
    s$gene.order <- rownames(s$timing)[order(s$timing$pt.peak, s$timing$pt.start,
        s$timing$pt.end, s$timing$exp.end, decreasing = c(F, F, F, T),
        method = "radix")]

    return(s)
}
```

##### filter.heatmap.genes

Removes undesired (mitochondrial, ribosomal, tandem duplicated genes) from heatmaps for presentation purposes.

- genes: (Character vector) genes to check
- Returns `genes` with undesired genes removed.

```
filter.heatmap.genes <- function(genes) {
    mt.genes <- grep("^mt-", ignore.case = T, genes, value = T)
    many.genes <- grep("\\(1 of many\\)", ignore.case = T, genes, value = T)
    ribo.genes <- grep("^rpl|^rps", ignore.case = T, genes, value = T)
    cox.genes <- grep("^cox", ignore.case = T, genes, value = T)
    hsp.genes <- grep("^hsp", ignore.case = T, genes, value = T)
    return(setdiff(genes, c(mt.genes, many.genes, ribo.genes, cox.genes,
        hsp.genes)))
}
```

#### Heatmaps of gene cascades

Using the genes that were determined as differentially expressed along the way to particular cell types, we generated expression cascades and plotted them as heatmaps.

#### Pdyn+ neurons

##### Prepare cascade

```
## Pdyn+ neurons: Seg 4

# Get markers from the two approaches
t <- combined.markers.best[["4"]]  # pdyn+ markers from above the combined clades
t <- t[t$exp.global.fc >= 0.8, ]  # limited to those with good global parameters
m <- threshold.tree.markers(markers, "4", global.fc = 0.6)  # Pdyn+ Cell markers from aucprTestAlongTree
pdyn.markers <- unique(c(rownames(t), rownames(m)))

# Just want to plot part of cells from upstream segment 12, which is
# very long. Going to use cells from segments 4, 11, and from segment
# 12 with pseudotime > 0.23
cells.plot <- unique(c(intersect(cellsInCluster(obj, "segment", "12"),
    whichCells(obj, "pseudotime", c(0.45, Inf))), cellsInCluster(obj, "segment",
    c("11", "4"))))

# Calculate spline curve
spline.plot <- geneSmoothFit(obj, pseudotime = "pseudotime", cells = cells.plot,
    genes = pdyn.markers, method = "spline", moving.window = 5, cells.per.window = 25,
    pseudotime.per.window = 0.005, spar = 0.5, verbose = F)

# Calculate gene expression timing for ordering rows
spline.plot <- determine.timing(s = spline.plot)
order.plot <- filter.heatmap.genes(spline.plot$gene.order)

# Output gene table
table.save <- data.frame(gene = order.plot, stringsAsFactors = F)
table.save$clade.AUCPR.ratio <- t[table.save$gene, "AUCPR.ratio"]
table.save$clade.exp.fc <- t[table.save$gene, "exp.fc"]
table.save$clade.exp.fc.global <- t[table.save$gene, "exp.global.fc"]
table.save$pdyn.AUCPR.ratio.all <- m[table.save$gene, "AUCPR.ratio.all"]
table.save$pdyn.AUCPR.ratio.maxBranch <- m[table.save$gene, "AUCPR.ratio.maxBranch"]
table.save$pdyn.exp.fc.all <- m[table.save$gene, "expfc.all"]
table.save$pdyn.exp.fc.best <- m[table.save$gene, "expfc.maxBranch"]
write.csv(table.save, quote = F, file = paste0(base.path, "/heatmaps/hypo-pdyn.csv"))
```

##### Generate heatmap: all genes

```
## Generate heatmap Make sure any values <0 in the spline curves get set
## to 0 so that the heatmap scale doesn't get messed up.
spline.plot$scaled.smooth[spline.plot$scaled.smooth < 0] <- 0
# Open a PDF and generate the heatmap pdf(paste0(base.path,
# '/heatmaps/hypo-pdyn.pdf'), width=6, height=10)
gplots::heatmap.2(x = as.matrix(spline.plot$scaled.smooth[order.plot, ]),
    Rowv = F, Colv = F, dendrogram = "none", col = cols, trace = "none",
    density.info = "none", key = F, cexCol = 0.8, cexRow = 0.6, margins = c(8,
        8), lwid = c(0.3, 4), lhei = c(0.3, 4), labCol = NA)
title(main = "pdyn+ Neurons")
```

#### pdyn+ Neurons

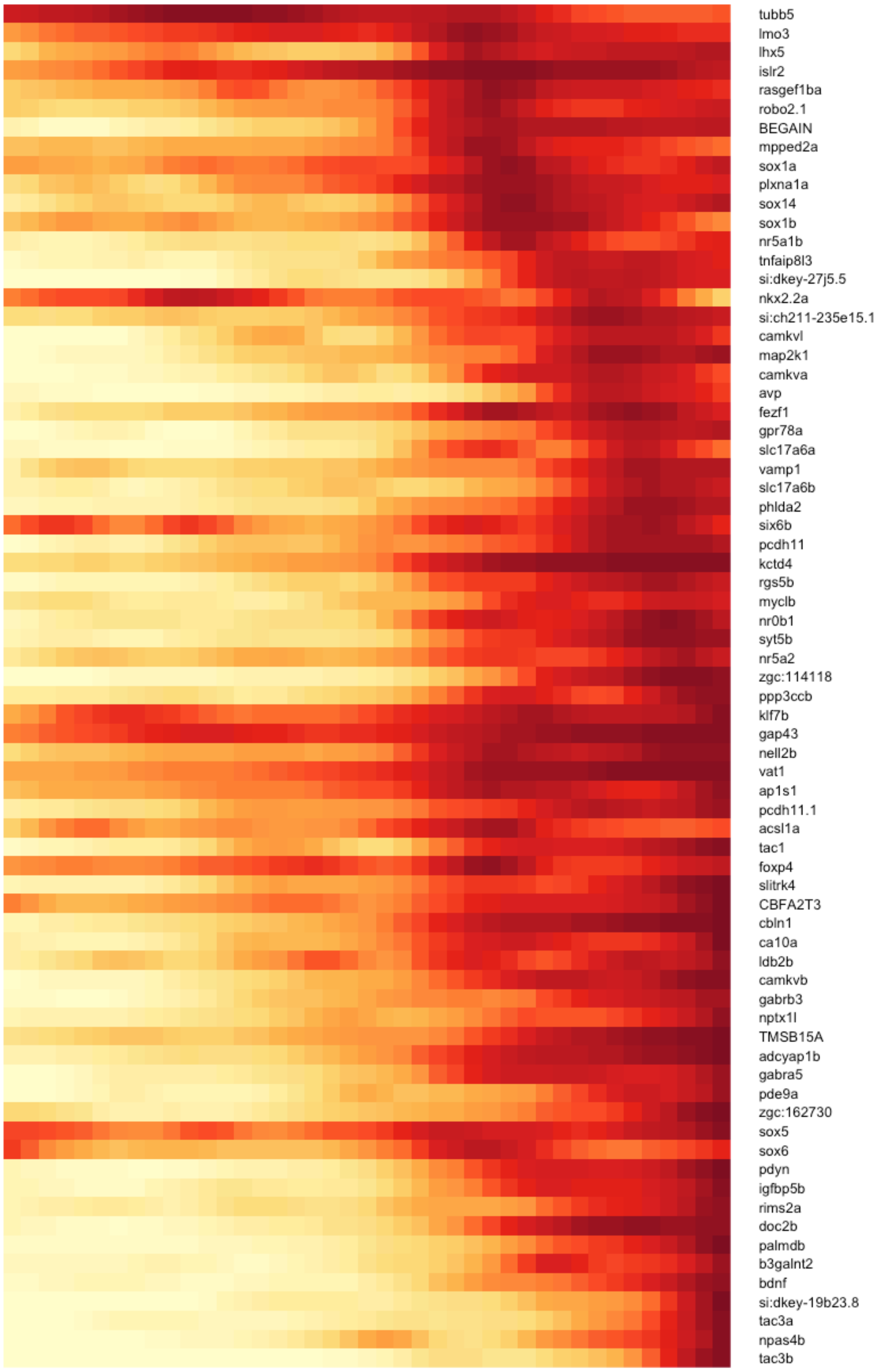

```
# dev.off()
```

#### Prdx1+ neurons vs. other neurons

##### Prepare cascade

```
## Prdx1+ neurons: Seg 3

# Get markers from the two approaches
t <- combined.markers.best[["3"]]  # prdx1+ markers from above the combined clades
t <- t[t$exp.global.fc >= 0.8, ]  # limited to those with good global parameters
m <- threshold.tree.markers(markers, "3", global.fc = 0.6)  # prdx1+ Cell markers from aucprTestAlongTree
prdx.markers <- unique(c(rownames(t), rownames(m)))

# Get markers for the opposing segment
opposing.prdx.markers <- intersect(rownames(nonprdx.markers), rownames(nonprdx.markers.global))

prdx.hm.markers <- unique(c(prdx.markers, opposing.prdx.markers))

# Calculate spline curves Using segments 13 and 12 or 3.
spline.12 <- geneSmoothFit(obj, pseudotime = "pseudotime", cells = cellsInCluster(obj,
    "segment", c("13", "12")), genes = prdx.hm.markers, method = "spline",
    moving.window = 5, cells.per.window = 25, pseudotime.per.window = 0.005,
    spar = 0.5, verbose = F)
spline.3 <- geneSmoothFit(obj, pseudotime = "pseudotime", cells = cellsInCluster(obj,
    "segment", c("13", "3")), genes = prdx.hm.markers, method = "spline",
    moving.window = 5, cells.per.window = 25, pseudotime.per.window = 0.005,
    spar = 0.5, verbose = F)
spline.123 <- geneSmoothFit(obj, pseudotime = "pseudotime", cells = cellsInCluster(obj,
    "segment", c("13", "12", "3")), genes = prdx.hm.markers, method = "spline",
    moving.window = 5, cells.per.window = 25, pseudotime.per.window = 0.005,
    spar = 0.5, verbose = F)

# Want to plot a heatmap that shows expression in most upstream
# progenitors and then each branch (i.e. prdx1- vs. prdx1+ neurons) as
# separate columns. Going to crop each spline fit to the correct
# pseudotime range and then combine them into a single one that can be
# plotted as a three-column heatmap.

pt.12v3 <- obj@tree$segment.pseudotime.limits["3", "start"]  # pseudotime where the crop should happen
splines.prdx <- list(cropSmoothFit(spline.123, pt.min = -Inf, pt.max = pt.12v3),
    cropSmoothFit(spline.12, pt.min = pt.12v3, pt.max = Inf), cropSmoothFit(spline.3,
        pt.min = pt.12v3, pt.max = Inf))
names(splines.prdx) <- c("Hypo Precursors", "Prdx1-", "Prdx1+")
splines.prdx.hm <- combineSmoothFit(splines.prdx)  # Combine into a single one

# Calculate gene expression timing for ordering rows
spline.12 <- determine.timing(s = spline.12)
spline.3 <- determine.timing(s = spline.3)
spline.123 <- determine.timing(s = spline.123)
```

```
# Decide which markers are specific to one cell type or both
d12v3 <- divide.branches(obj, prdx.hm.markers, clust.1 = "12", clust.2 = "3",
    exp.fc = 0.4, exp.thresh = 0.2, exp.diff = 0.1)

# Generate gene ordering based on timing & specificity
order.123 <- filter.heatmap.genes(setdiff(spline.123$gene.order, c(d12v3$specific.1,
    d12v3$specific.2)))
order.12 <- filter.heatmap.genes(intersect(spline.12$gene.order, d12v3$specific.1))
order.3 <- filter.heatmap.genes(intersect(spline.3$gene.order, d12v3$specific.2))
gene.order <- c(order.123, order.12, order.3)

# Output gene table
table.save <- data.frame(gene = gene.order, marks = c(rep("both", length(order.123)),
    rep("prdx1-", length(order.12)), rep("prdx1+", length(order.3))), stringsAsFactors = F)
table.save$clade.AUCPR.ratio <- t[table.save$gene, "AUCPR.ratio"]
table.save$clade.exp.fc <- t[table.save$gene, "exp.fc"]
table.save$clade.exp.fc.global <- t[table.save$gene, "exp.global.fc"]
table.save$non.AUCPR.ratio <- nonprdx.markers[table.save$gene, "AUCPR.ratio"]
table.save$non.exp.fc <- nonprdx.markers[table.save$gene, "exp.fc"]
table.save$non.exp.fc.global <- nonprdx.markers.global[table.save$gene,
    "exp.fc"]
table.save$prdx.AUCPR.ratio.all <- m[table.save$gene, "AUCPR.ratio.all"]
table.save$prdx.AUCPR.ratio.maxBranch <- m[table.save$gene, "AUCPR.ratio.maxBranch"]
table.save$prdx.exp.fc.all <- m[table.save$gene, "expfc.all"]
table.save$prdx.exp.fc.best <- m[table.save$gene, "expfc.maxBranch"]
write.csv(table.save, quote = F, file = paste0(base.path, "/heatmaps/hypo-prdx1-vs-non.csv"))
```

##### Generate heatmap: all genes

```
# Make sure any values <0 in the spline curves get set to 0 so that the
# heatmap scale doesn't get messed up.
splines.prdx.hm$scaled.smooth[splines.prdx.hm$scaled.smooth < 0] <- 0
# Determine where to place column separators (i.e. how many columns
# will each cell type occupy in the heatmap )
colsep <- cumsum(as.numeric(head(unlist(lapply(splines.prdx, function(x) ncol(x$scaled.smooth))),
    -1)))
# Determine where to place row separators (i.e. how many common
# markers, and markers are specific to each cell type)
rowsep <- cumsum(c(length(order.123), length(order.12)))
# Open a PDF and generate the heatmap pdf(paste0(base.path,
# '/heatmaps/hypo-prdx-vs-non.pdf'), width=6, height=10)
gplots::heatmap.2(x = as.matrix(splines.prdx.hm$scaled.smooth[gene.order,
    ]), Rowv = F, Colv = F, dendrogram = "none", col = cols, trace = "none",
    density.info = "none", key = F, cexCol = 0.8, cexRow = 0.35, margins = c(8,
        8), lwid = c(0.3, 4), lhei = c(0.3, 4), labCol = NA, colsep = colsep,
    rowsep = rowsep, sepwidth = c(0.05, 0.2))
title(main = "Precursors", line = -41, adj = 0)
title(main = "prdx1 -", line = -41, adj = 0.425)
title(main = "prdx1 +", line = -41, adj = 0.725)
```

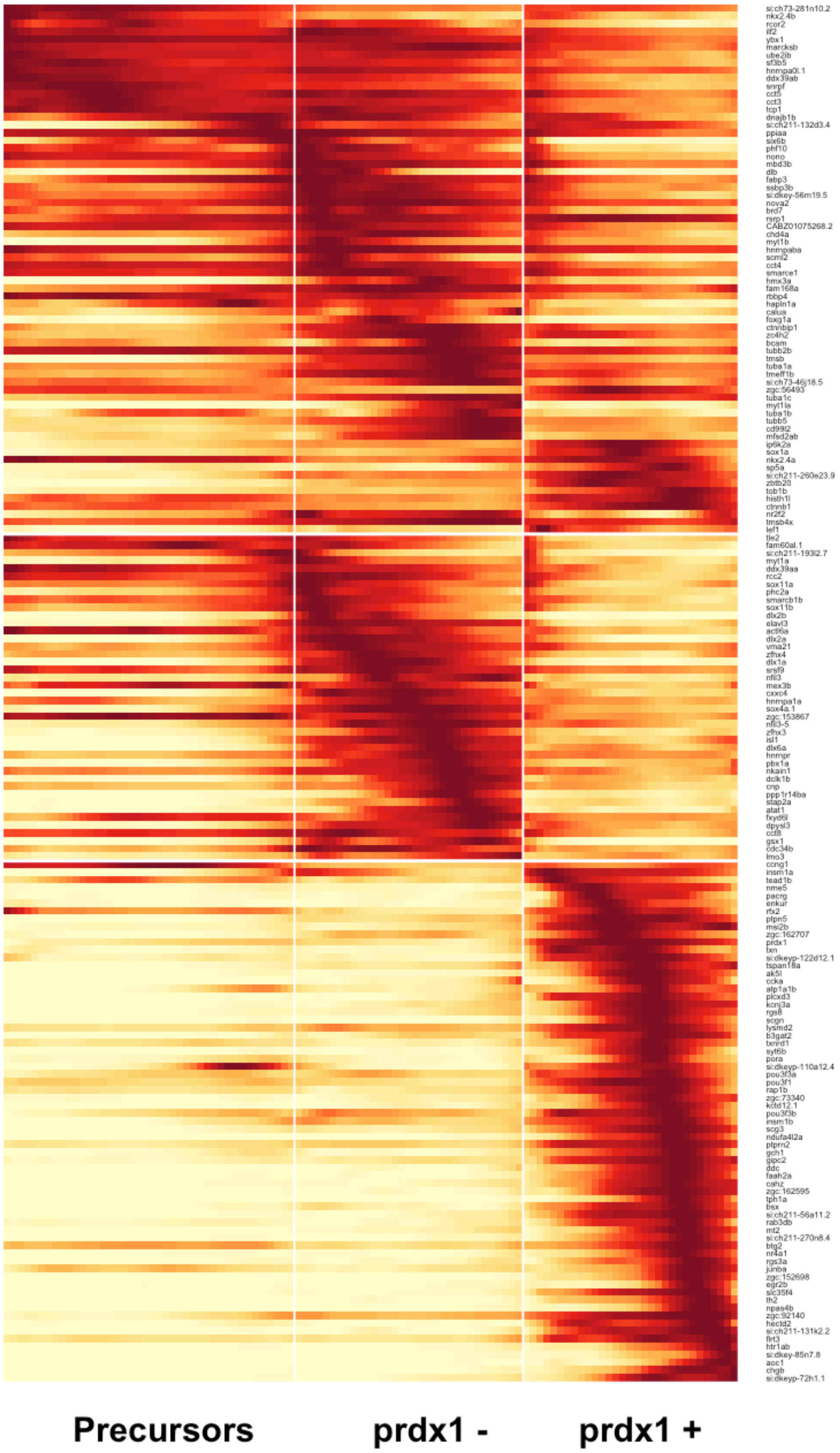
Precursors
prdx1 -
prdx1 +

```
# dev.off()
```

#### nrgna+ neurons

##### Prepare cascade

```
# Get markers from the two approaches
t <- combined.markers.best[["10"]]  # Markers from above the combined clades
t <- t[t$exp.global.fc >= 0.8, ]  # limited to those with good global parameters
m2 <- threshold.tree.markers(markers, "2", global.fc = 0.6)  # Markers from aucprTestAlongTree
m5 <- threshold.tree.markers(markers, "5", global.fc = 0.6)  # Markers from aucprTestAlongTree
tacsyn.markers <- unique(c(rownames(t), rownames(m2), rownames(m5)))

# Just want to plot part of cells from upstream segment 12, which is
# very long. Going to use cells from segments 2 or 5, 10, and from
# segment 12 with pseudotime > 0.23
cells.plot.2 <- unique(c(intersect(cellsInCluster(obj, "segment", "12"),
    whichCells(obj, "pseudotime", c(0.45, Inf))), cellsInCluster(obj, "segment",
    c("10", "2"))))
cells.plot.5 <- unique(c(intersect(cellsInCluster(obj, "segment", "12"),
    whichCells(obj, "pseudotime", c(0.45, Inf))), cellsInCluster(obj, "segment",
    c("10", "5"))))
cells.plot.25 <- unique(c(intersect(cellsInCluster(obj, "segment", "12"),
    whichCells(obj, "pseudotime", c(0.45, Inf))), cellsInCluster(obj, "segment",
    c("10", "2", "5"))))

# Calculate spline curves
spline.2 <- geneSmoothFit(obj, pseudotime = "pseudotime", cells = cells.plot.2,
    genes = tacsyn.markers, method = "spline", moving.window = 5, cells.per.window = 25,
    pseudotime.per.window = 0.005, spar = 0.5, verbose = F)
spline.5 <- geneSmoothFit(obj, pseudotime = "pseudotime", cells = cells.plot.5,
    genes = tacsyn.markers, method = "spline", moving.window = 5, cells.per.window = 25,
    pseudotime.per.window = 0.005, spar = 0.5, verbose = F)
spline.25 <- geneSmoothFit(obj, pseudotime = "pseudotime", cells = cells.plot.25,
    genes = tacsyn.markers, method = "spline", moving.window = 5, cells.per.window = 25,
    pseudotime.per.window = 0.005, spar = 0.5, verbose = F)

# Want to plot a heatmap that shows expression in upstream progenitors
# and then each branch (i.e. gabaergic_tac1 vs. synpr+) as separate
# columns. Going to crop each spline fit to the correct pseudotime
# range and then combine them into a single one that can be plotted as
# a three-column heatmap.

pt.2v5 <- obj@tree$segment.pseudotime.limits["2", "start"]  # pseudotime where the crop should happen
splines.tacsyn <- list(cropSmoothFit(spline.25, pt.min = -Inf, pt.max = pt.2v5),
    cropSmoothFit(spline.2, pt.min = pt.2v5, pt.max = Inf), cropSmoothFit(spline.5,
        pt.min = pt.2v5, pt.max = Inf))
names(splines.tacsyn) <- c("Precursors", "Synpr-", "Synpr+")
splines.tacsyn.hm <- combineSmoothFit(splines.tacsyn)  # Combine into a single one

# Calculate gene expression timing for ordering rows
```

```r
spline.2 <- determine.timing(s = spline.2)
spline.5 <- determine.timing(s = spline.5)
spline.25 <- determine.timing(s = spline.25)

# Decide which markers are specific to one cell type or both
d2v5 <- divide.branches(obj, tacsyn.markers, clust.1 = "2", clust.2 = "5",
    exp.fc = 0.4, exp.thresh = 0.2, exp.diff = 0.1)

# Generate gene ordering based on timing & specificity
order.25 <- filter.heatmap.genes(setdiff(spline.25$gene.order, c(d2v5$specific.1,
    d2v5$specific.2)))
order.2 <- filter.heatmap.genes(intersect(spline.2$gene.order, d2v5$specific.1))
order.5 <- filter.heatmap.genes(intersect(spline.5$gene.order, d2v5$specific.2))
gene.order <- c(order.25, order.2, order.5)

# Output gene table
table.save <- data.frame(gene = gene.order, marks = c(rep("both", length(order.25)),
    rep("synpr-", length(order.2)), rep("synpr+", length(order.5))), stringsAsFactors = F)
table.save$clade.AUCPR.ratio <- t[table.save$gene, "AUCPR.ratio"]
table.save$clade.exp.fc <- t[table.save$gene, "exp.fc"]
table.save$clade.exp.fc.global <- t[table.save$gene, "exp.global.fc"]
table.save$nonsynpr.AUCPR.ratio.all <- m2[table.save$gene, "AUCPR.ratio.all"]
table.save$nonsynpr.AUCPR.ratio.maxBranch <- m2[table.save$gene, "AUCPR.ratio.maxBranch"]
table.save$nonsynpr.exp.fc.all <- m2[table.save$gene, "expfc.all"]
table.save$nonsynpr.exp.fc.best <- m2[table.save$gene, "expfc.maxBranch"]
table.save$synpr.AUCPR.ratio.all <- m5[table.save$gene, "AUCPR.ratio.all"]
table.save$synpr.AUCPR.ratio.maxBranch <- m5[table.save$gene, "AUCPR.ratio.maxBranch"]
table.save$synpr.exp.fc.all <- m5[table.save$gene, "expfc.all"]
table.save$synpr.exp.fc.best <- m5[table.save$gene, "expfc.maxBranch"]
write.csv(table.save, quote = F, file = paste0(base.path, "/heatmaps/hypo-nrgna.csv"))
```

##### Generate heatmap: all genes

```r
# Make sure any values <0 in the spline curves get set to 0 so that the
# heatmap scale doesn't get messed up.
splines.tacsyn.hm$scaled.smooth[splines.tacsyn.hm$scaled.smooth < 0] <- 0
# Determine where to place column separators (i.e. how many columns
# will each cell type occupy in the heatmap )
colsep <- cumsum(as.numeric(head(unlist(lapply(splines.tacsyn, function(x) ncol(x$scaled.smooth))),
    -1)))
# Determine where to place row separators (i.e. how many common
# markers, and markers are specific to each cell type)
rowsep <- cumsum(c(length(order.25), length(order.2)))
# Open a PDF and generate the heatmap pdf(paste0(base.path,
# '/heatmaps/hypo-nrgna.pdf'), width=6, height=10)
gplots::heatmap.2(x = as.matrix(splines.tacsyn.hm$scaled.smooth[gene.order,
    ]), Rowv = F, Colv = F, dendrogram = "none", col = cols, trace = "none",
    density.info = "none", key = F, cexCol = 0.8, cexRow = 0.35, margins = c(8,
        8), lwid = c(0.3, 4), lhei = c(0.3, 4), labCol = NA, colsep = colsep,
    rowsep = rowsep, sepwidth = c(0.05, 0.2))
title(main = "synpr-", line = -41, adj = 0.535)
```

```r
title(main = "synpr+", line = -41, adj = 0.75)
```

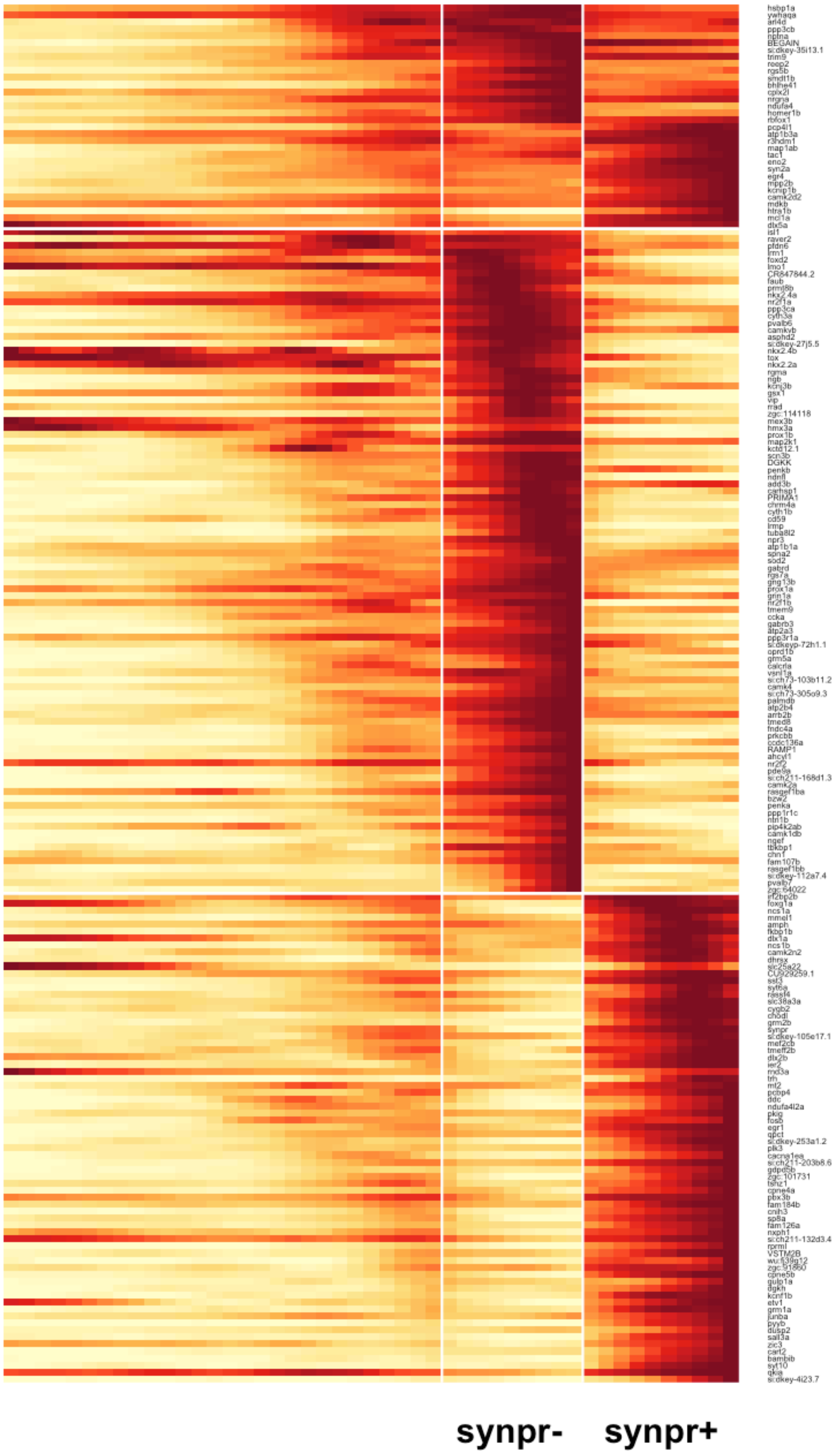
synpr-
synpr+

```
# dev.off()
```

Generate heatmap: main figure

```
# Plot heatmap with only certain genes labeled for main figure

genes.to.plot <- c("rgs5b", "dlx5a", "isl1", "nkx2.4a", "nkx2.2a", "hmx3a",
    "dlx1a", "dlx2b", "sp8a", "pbx3b")
rownames.to.plot <- gene.order
rtp <- rownames.to.plot %in% genes.to.plot
rownames.to.plot[!rtp] <- ""
rownames.to.plot[rtp] <- paste0("- ", rownames.to.plot[rtp])

# Open a PDF and generate the heatmap pdf(paste0(base.path,
# '/heatmaps/hypo-nrgna-mainfig.pdf'), width=6, height=10)
gplots::heatmap.2(x = as.matrix(splines.tacsyn.hm$scaled.smooth[gene.order,
    ]), Rowv = F, Colv = F, dendrogram = "none", col = cols, trace = "none",
    density.info = "none", key = F, cexCol = 0.8, cexRow = 1.8, margins = c(8,
        8), lwid = c(0.3, 4), lhei = c(0.3, 4), labCol = NA, colsep = colsep,
    rowsep = rowsep, sepwidth = c(0.05, 0.2), labRow = rownames.to.plot)
title(main = "synpr-", line = -41, adj = 0.535)
title(main = "synpr+", line = -41, adj = 0.75)
```

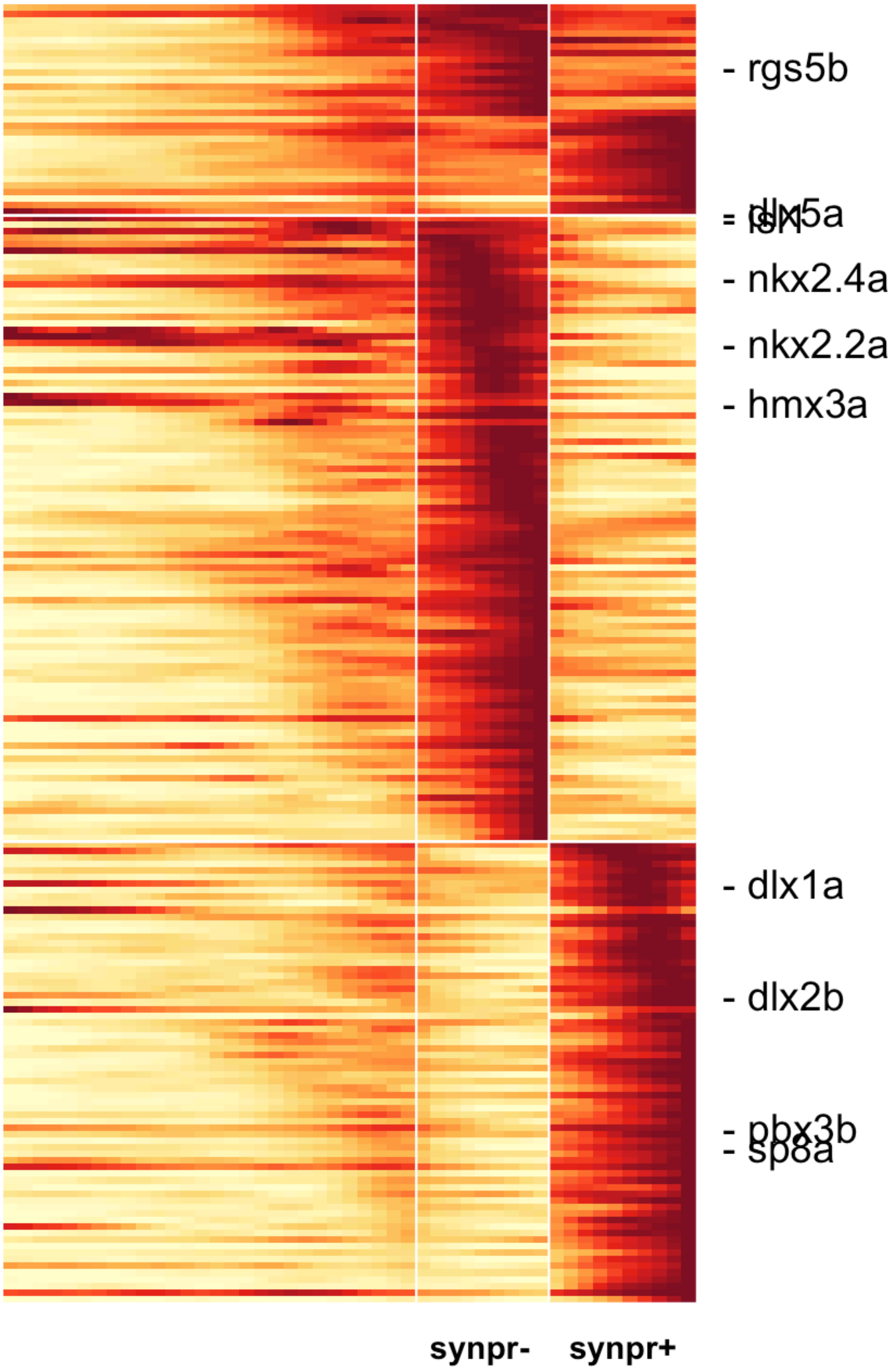
- rgs5b
- dlx5a
- isl1
- nkx2.4a
- nkx2.2a
- hmx3a
- dlx1a
- dlx2b
- pbx3b
- sp8a
synpr-
synpr+

```
# dev.off()
```

#### GABAergic neurons

##### Prepare cascade

```
# Get markers from the two approaches
t <- combined.markers.best[["9"]]  # Markers from above the combined clades
m1 <- threshold.tree.markers(markers, "1", global.fc = 0.6)  # Markers from aucprTestAlongTree
m6 <- threshold.tree.markers(markers, "6", global.fc = 0.6)  # Markers from aucprTestAlongTree
m7 <- threshold.tree.markers(markers, "7", global.fc = 0.6)  # Markers from aucprTestAlongTree
gaba.markers <- unique(c(rownames(t), rownames(m1), rownames(m6), rownames(m7)))

# Defining cell populations to use in the heatmap Just want to use a
# tiny bit of segment 12 and then the rest of the gabaergic clade
cells.plot.1 <- unique(c(intersect(cellsInCluster(obj, "segment", "12"),
    whichCells(obj, "pseudotime", c(0.5, Inf))), cellsInCluster(obj, "segment",
    c("11", "9", "1"))))
cells.plot.6 <- unique(c(intersect(cellsInCluster(obj, "segment", "12"),
    whichCells(obj, "pseudotime", c(0.5, Inf))), cellsInCluster(obj, "segment",
    c("11", "9", "8", "6"))))
cells.plot.7 <- unique(c(intersect(cellsInCluster(obj, "segment", "12"),
    whichCells(obj, "pseudotime", c(0.5, Inf))), cellsInCluster(obj, "segment",
    c("11", "9", "8", "7"))))
cells.plot.167 <- unique(c(intersect(cellsInCluster(obj, "segment", "12"),
    whichCells(obj, "pseudotime", c(0.5, Inf))), cellsInCluster(obj, "segment",
    c("11", "9", "8", "1", "6", "7"))))

# Calculate spline curves
spline.1 <- geneSmoothFit(obj, pseudotime = "pseudotime", cells = cells.plot.1,
    genes = gaba.markers, method = "spline", moving.window = 5, cells.per.window = 25,
    pseudotime.per.window = 0.005, spar = 0.5, verbose = F)
spline.6 <- geneSmoothFit(obj, pseudotime = "pseudotime", cells = cells.plot.6,
    genes = gaba.markers, method = "spline", moving.window = 5, cells.per.window = 25,
    pseudotime.per.window = 0.005, spar = 0.5, verbose = F)
spline.7 <- geneSmoothFit(obj, pseudotime = "pseudotime", cells = cells.plot.7,
    genes = gaba.markers, method = "spline", moving.window = 5, cells.per.window = 25,
    pseudotime.per.window = 0.005, spar = 0.5, verbose = F)
spline.167 <- geneSmoothFit(obj, pseudotime = "pseudotime", cells = cells.plot.167,
    genes = gaba.markers, method = "spline", moving.window = 5, cells.per.window = 25,
    pseudotime.per.window = 0.005, spar = 0.5, verbose = F)

# Want to plot a heatmap that shows expression in upstream progenitors
# and then each branch as separate columns. Going to crop each spline
# fit to the correct pseudotime range and then combine them into a
# single one that can be plotted as a three-column heatmap.

pt.1 <- obj@tree$segment.pseudotime.limits["1", "start"]  # pseudotime where the crop should happen
splines.gaba <- list(cropSmoothFit(spline.167, pt.min = -Inf, pt.max = pt.1),
    cropSmoothFit(spline.1, pt.min = pt.1, pt.max = Inf), cropSmoothFit(spline.6,
        pt.min = pt.1, pt.max = Inf), cropSmoothFit(spline.7, pt.min = pt.1,
```

```
        pt.max = Inf))
names(splines.gaba) <- c("Precursors", "dlx+", "sst1.1+", "tph2+")
splines.gaba.hm <- combineSmoothFit(splines.gaba)  # Combine into a single one

# Calculate gene expression timing for ordering rows
spline.1 <- determine.timing(s = spline.1)
spline.6 <- determine.timing(s = spline.6)
spline.7 <- determine.timing(s = spline.7)
spline.167 <- determine.timing(s = spline.167, genes = setdiff(gaba.markers,
    "sst1.2"))

# Decide which markers are specific to one cell type or not specific
d <- divide.branches.triple(obj, gaba.markers, clust.1 = "1", clust.2 = "6",
    clust.3 = "7", exp.fc = 0.4, exp.thresh = 0.2, exp.diff = 0.1)

# Generate gene ordering based on timing & specificity
order.167 <- filter.heatmap.genes(intersect(spline.167$gene.order, d$nonspecific))
order.1 <- filter.heatmap.genes(intersect(spline.1$gene.order, d$specific.1))
order.6 <- filter.heatmap.genes(intersect(spline.6$gene.order, d$specific.2))
order.7 <- filter.heatmap.genes(intersect(spline.7$gene.order, d$specific.3))
gene.order <- c(order.167, order.1, order.6, order.7)

# Output gene table
table.save <- data.frame(gene = gene.order, marks = c(rep("multiple", length(order.167)),
    rep("dlx+", length(order.1)), rep("sst+", length(order.6)), rep("tph2+",
        length(order.7))), stringsAsFactors = F)
table.save$clade.AUCPR.ratio <- t[table.save$gene, "AUCPR.ratio"]
table.save$clade.exp.fc <- t[table.save$gene, "exp.fc"]
table.save$clade.exp.fc.global <- t[table.save$gene, "exp.global.fc"]
table.save$dlx.AUCPR.ratio.all <- m1[table.save$gene, "AUCPR.ratio.all"]
table.save$dlx.AUCPR.ratio.maxBranch <- m1[table.save$gene, "AUCPR.ratio.maxBranch"]
table.save$dlx.exp.fc.all <- m1[table.save$gene, "expfc.all"]
table.save$dlx.exp.fc.best <- m1[table.save$gene, "expfc.maxBranch"]
table.save$sst.AUCPR.ratio.all <- m6[table.save$gene, "AUCPR.ratio.all"]
table.save$sst.AUCPR.ratio.maxBranch <- m6[table.save$gene, "AUCPR.ratio.maxBranch"]
table.save$sst.exp.fc.all <- m6[table.save$gene, "expfc.all"]
table.save$sst.exp.fc.best <- m6[table.save$gene, "expfc.maxBranch"]
table.save$tph.AUCPR.ratio.all <- m7[table.save$gene, "AUCPR.ratio.all"]
table.save$tph.AUCPR.ratio.maxBranch <- m7[table.save$gene, "AUCPR.ratio.maxBranch"]
table.save$tph.exp.fc.all <- m7[table.save$gene, "expfc.all"]
table.save$tph.exp.fc.best <- m7[table.save$gene, "expfc.maxBranch"]
write.csv(table.save, quote = F, file = paste0(base.path, "/heatmaps/hypo-gaba.csv"))
```

##### Generate heatmap: all genes

```
# Make sure any values <0 in the spline curves get set to 0 so that the
# heatmap scale doesn't get messed up.
splines.gaba.hm$scaled.smooth[splines.gaba.hm$scaled.smooth < 0] <- 0
# Determine where to place column separators (i.e. how many columns
# will each cell type occupy in the heatmap )
colsep <- cumsum(as.numeric(head(unlist(lapply(splines.gaba, function(x) ncol(x$scaled.smooth))),
```

```r
    -1)))
# Determine where to place row separators (i.e. how many common
# markers, and markers are specific to each cell type)
rowsep <- cumsum(c(length(order.167), length(order.1), length(order.6)))
# Open a PDF and generate the heatmap pdf(paste0(base.path,
# '/heatmaps/hypo-gaba.pdf'), width=6, height=10)
gplots::heatmap.2(x = as.matrix(splines.gaba.hm$scaled.smooth[gene.order,
    ]), Rowv = F, Colv = F, dendrogram = "none", col = cols, trace = "none",
    density.info = "none", key = F, cexCol = 0.8, cexRow = 1, margins = c(8,
        8), lwid = c(0.3, 4), lhei = c(0.3, 4), labCol = NA, colsep = colsep,
    rowsep = rowsep, sepwidth = c(0.05, 0.2))
title(main = "dlx+", line = -41, adj = 0.45)
title(main = "sst+", line = -41, adj = 0.61)
title(main = "tph2+", line = -41, adj = 0.75)
```

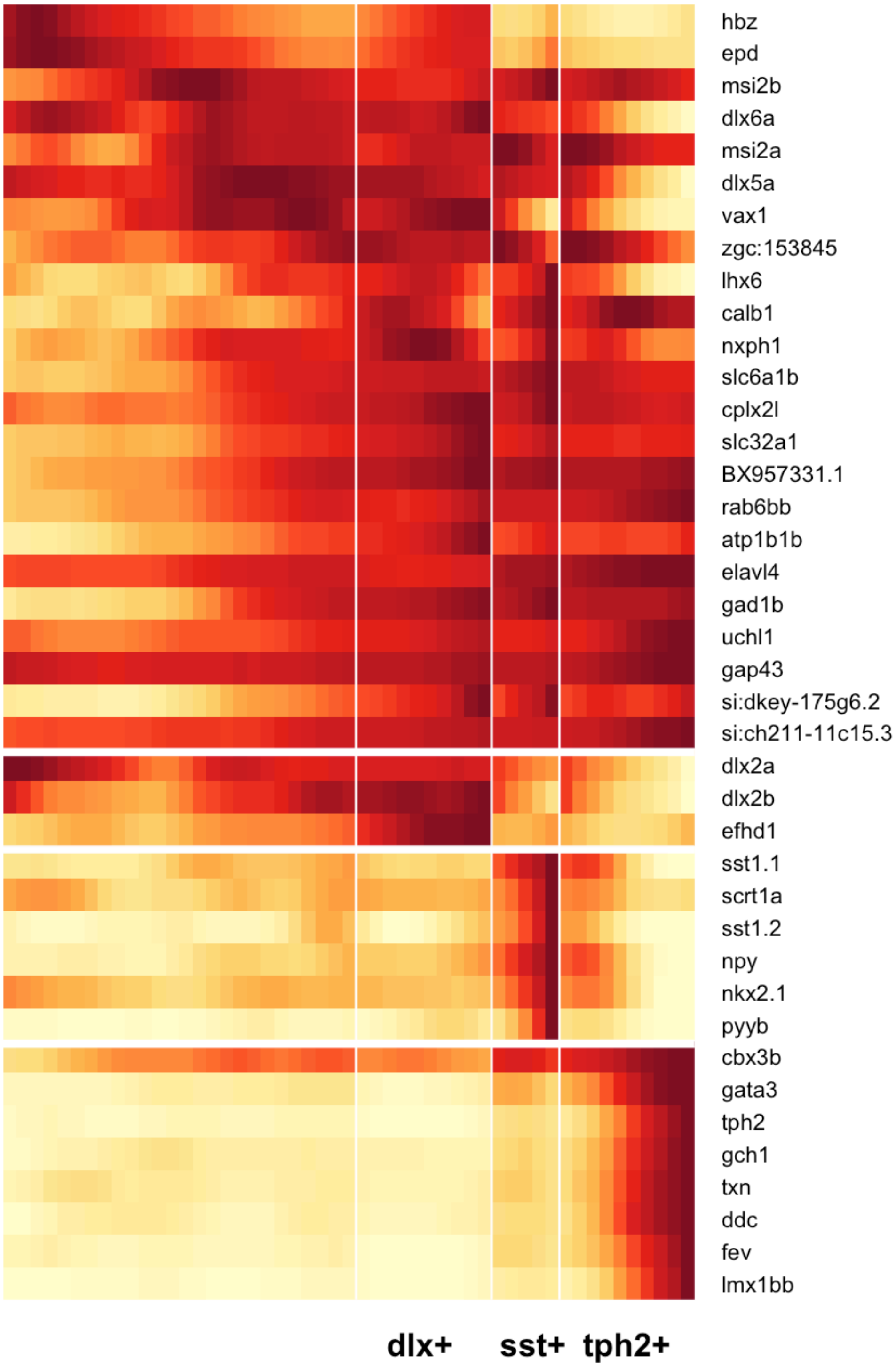
hbz
epd
msi2b
dlx6a
msi2a
dlx5a
vax1
zgc:153845
lhx6
calb1
nxph1
slc6a1b
cplx2l
slc32a1
BX957331.1
rab6bb
atp1b1b
elavl4
gad1b
uchl1
gap43
si:dkey-175g6.2
si:ch211-11c15.3
dlx2a
dlx2b
efhd1
sst1.1
scrt1a
sst1.2
npy
nkx2.1
pyyb
cbx3b
gata3
tph2
gch1
txn
ddc
fev
lmx1bb
dlx+
sst+
tph2+

```r
# dev.off()
```

#### Long-term undifferentiated states

We wanted to look for differences in the transcriptional states of long-term precursors between the retina and the hypothalamus. In the retina, cells persist in progenitor states, while in the hypothalamus, the progenitor state is short-lived, but cells persist in a neural precursor state.

##### GABA dlx+ is long-term

A cluster of neurons from 15 dpf that are GABAergic and dlx+ are found spanning many molecular states spanning the differentiation and maturation process of this cell type.

```r
plotTreeHighlight(obj, label.name = "clus.orig", label.value = "12-15d_2",
    highlight.size = 1, title = "15 dpf Cluster 2: GABA dlx+ cells")
```

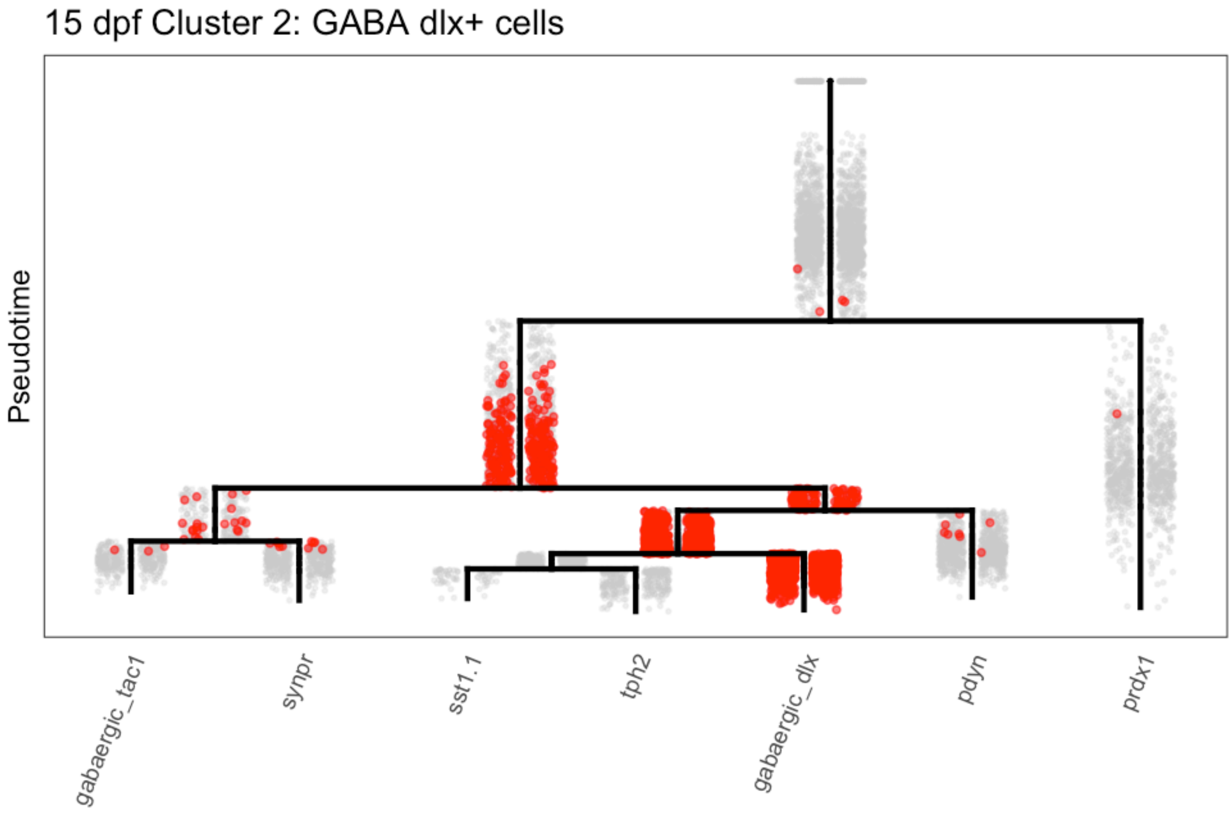

##### Identify populations to compare

We defined progenitor / precursor / neuron populations based on their location in the tree, cross-referenced with the expression of markers of each of these types.

```r
obj@group.ids$precursor.group <- NA

cells.s13 <- intersect(cellsInCluster(obj, "segment", "13"), whichCells(obj,
    "pseudotime", c(0.05, 1)))
obj@group.ids[cells.s13, "precursor.group"] <- "1_progenitors"

obj@group.ids[cellsInCluster(obj, "segment", "12"), "precursor.group"] <- "2_precursors"

obj@group.ids[cellsInCluster(obj, "segment", c("9", "1")), "precursor.group"] <- "3a_neurons"

obj@group.ids[cellsInCluster(obj, "segment", c("6", "7")), "precursor.group"] <- "3b_neurons_matured"
```

```r
# Colors for ggplot
stage.colors <- RColorBrewer::brewer.pal(12, "Paired")[c(9, 7, 1, 2)]

# Plot tree to show where the groups are
plotTree(obj, "precursor.group", discrete.colors = stage.colors)
```

```
## Warning: Removed 3944 rows containing missing values (geom_point).
```

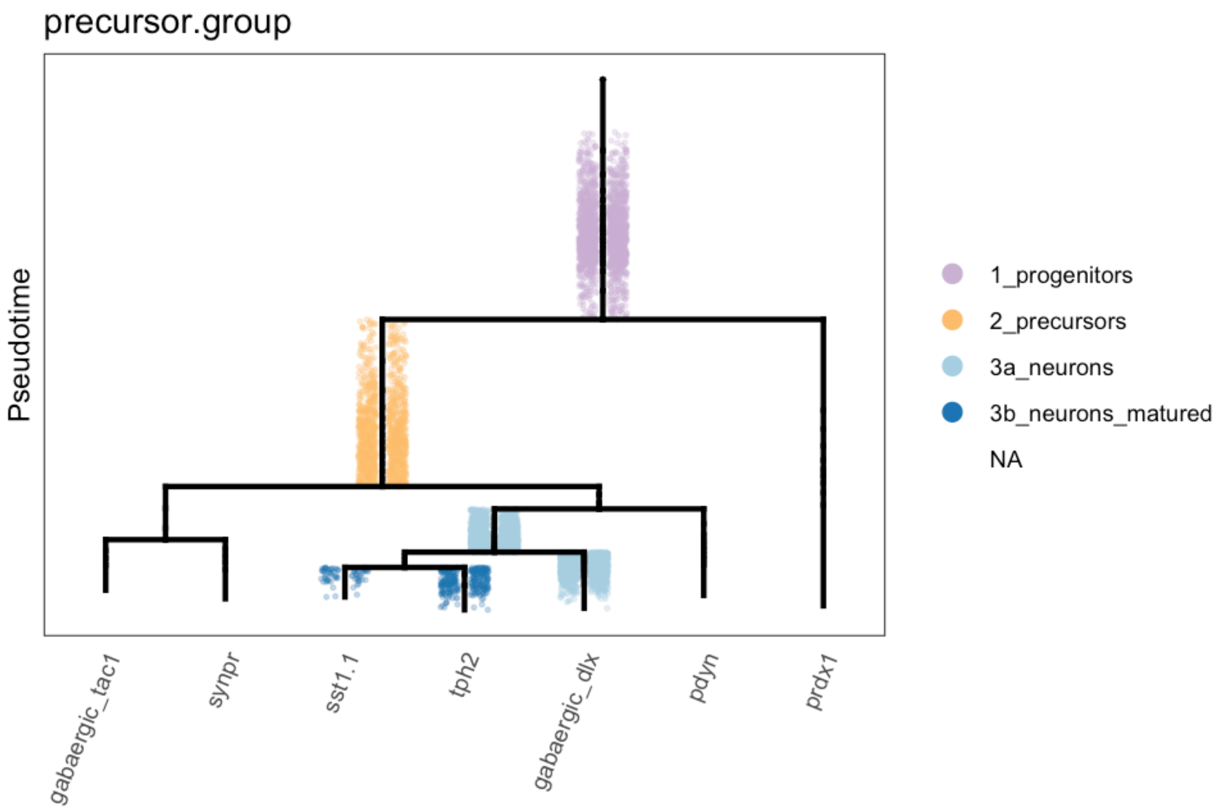

```r
# Plot genes in each group
plotDot(obj, genes = c("rx3", "shha", "nkx2.4b", "ascl1a", "scrt2", "scg2b",
    "elavl3", "elavl4"), clustering = "precursor.group", scale.by = "area") +
    theme_bw()
```

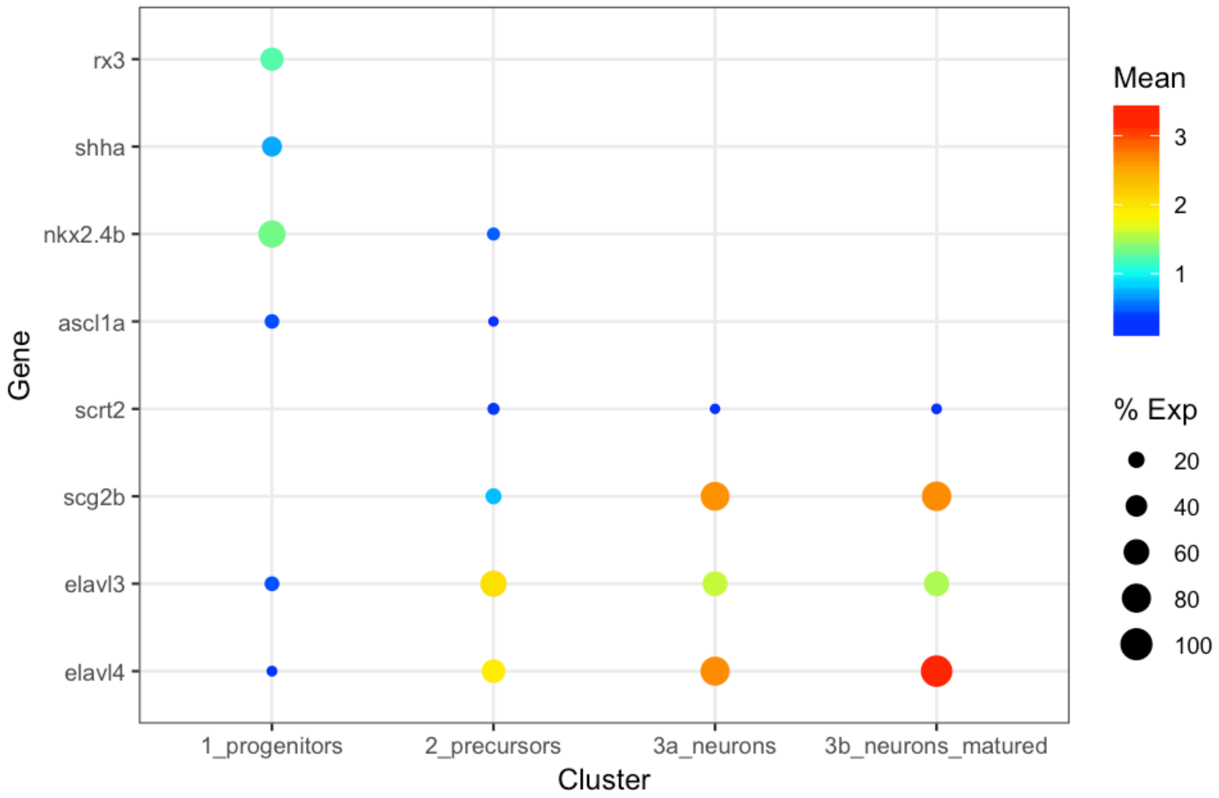

#### Determine proportion of cells in each state

We then determined the proportion of cells from different stages that fell into each of these transcriptional states.

```
# Combine stages to reduce number of plots
obj@group.ids$stage.collapsed <- plyr::mapvalues(x = obj@group.ids$stage,
    from = c("01-12h", "02-14h", "03-16h", "04-18h", "05-20h", "06-24h",
        "07-36h", "08-2d", "09-3d", "10-5d", "11-8d", "12-15d"), to = c(rep("01-12h-24h",
        6), rep("02-36h-3d", 3), rep("03-5d-15d", 3)))

# Count number of cells from each stage group in each precursor group
stage.group.count <- plyr::count(obj@group.ids, vars = c("stage.collapsed",
    "precursor.group"))

# Remove cells that weren't part of a precursor group
stage.group.count <- stage.group.count[complete.cases(stage.group.count),
    ]

# Cast into a data frame and convert NA to 0 (no cells of that type
# observed)
stage.group.df <- reshape2::dcast(stage.group.count, formula = stage.collapsed ~
    precursor.group)
```

```
## Using freq as value column: use value.var to override.
```

```r
stage.group.df[is.na(stage.group.df)] <- 0

# Normalize by the number of precursors from each stage group
stage.group.df[, 2:5] <- sweep(stage.group.df[, 2:5], 1, rowSums(stage.group.df[,
    2:5]), "/")

# Melt for ggplot
stage.group.df.melt <- reshape2::melt(stage.group.df, id.vars = "stage.collapsed")

# Plot proportions
ggplot(stage.group.df.melt, aes(x = variable, y = value, group = stage.collapsed,
    fill = variable)) + geom_bar(stat = "identity") + facet_wrap(~stage.collapsed) +
    theme_bw() + scale_fill_manual(values = stage.colors) + theme(axis.title.x = element_blank(),
    axis.text.x = element_blank(), axis.ticks.x = element_blank())
```

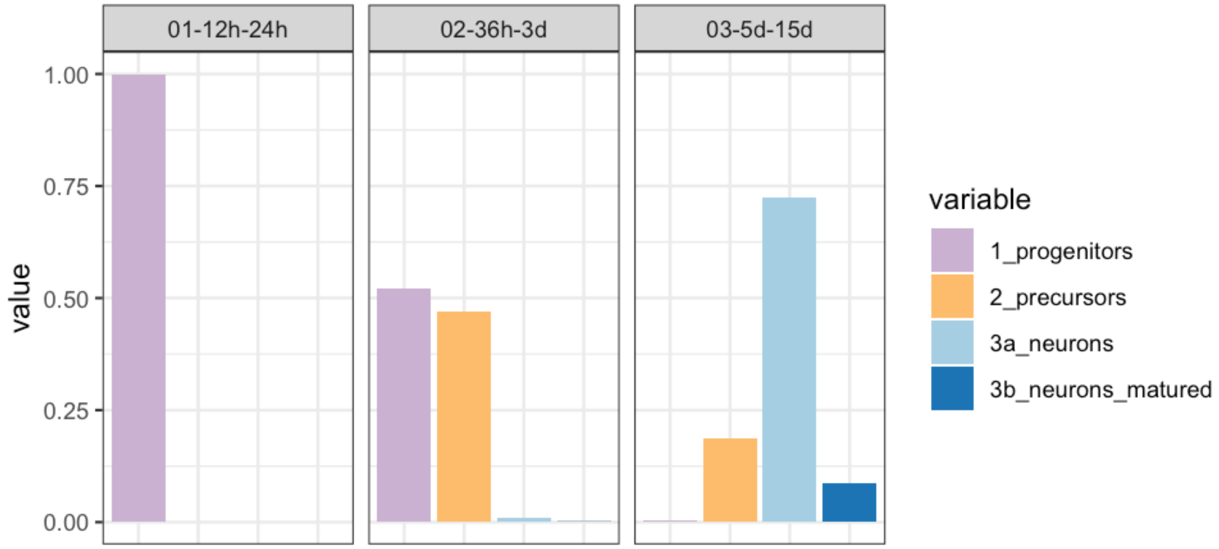
